# Speciation in kleptoparasites of oak gall wasps often correlates with a shift into a new tree habitat, tree organ, or gall morphospace

**DOI:** 10.1101/2023.09.07.556376

**Authors:** Anna K.G. Ward, Y. Miles Zhang, Guerin E. Brown, Alaine C. Hippee, Kirsten M. Prior, Shannon Rollins, Nicolas Sierra, Sofia I. Sheikh, Carly M. Tribull, Andrew A. Forbes

## Abstract

Host shifts to new plants can drive speciation for plant-feeding insects, but how commonly do host shifts also drive diversification for the parasites of those same insects? Oak gall wasps induce galls on oak trees and shifts to novel tree hosts and new tree organs have been implicated as drivers of oak gall wasp speciation. Gall wasps are themselves attacked by many insect parasites, which must find their hosts on the correct tree species and organ, but which also must navigate the morphologically variable galls with which they interact. Thus, we ask whether host shifts to new trees, organs, or gall morphologies correlate with gall parasite diversification. We delimit species and infer phylogenies for two genera of gall kleptoparasites, *Synergus* and *Ceroptres*, reared from a variety of North American oak galls. We find that most species were reared from galls induced by just one gall wasp species, and no parasite species was reared from galls of more than four species. Most kleptoparasite divergence events correlate with shifts to non-ancestral galls. These shifts often involved changes in tree habitat, gall location, and gall morphology. Host shifts are thus implicated in driving diversification for both oak gall wasps and their kleptoparasitic associates.

## Introduction

Natural selection in different environments acts on organismal traits, resulting in divergent adaptation to those environments (Lack 1947, Moodie 1972, Via 1990). But whether – and how often – divergent environmental adaptations contribute to speciation remains a subject of controversy, and the answer may differ among animal clades (Drès and Mallet 2002, Rocha et al. 2005, Rundle and Nosil 2005, Anderson and Weir 2022). For specialist insects at least, there is now little doubt that adaptation to different environments can and does lead to divergence (Matsubayashi et al. 2010, Forbes et al. 2017, McCulloch et al. 2020). Most examples involve phytophagous insects shifting to a new plant host, with spatial, temporal, or other differences between the ancestral and derived plant environments resulting in the evolution of reproductive isolating barriers between populations (Diehl and Bush 1984, Carroll and Boyd 1992, Funk 1998, Itami et al. 1998, Wood et al. 1999, Filchak et al. 2000, Matsubayashi and Katakura 2009, Bendall et al. 2017, Bagley et al. 2023). But is “host-shift speciation” important only for plant-feeding insects, or does it also commonly generate diversity at the next trophic level: parasitic insects attacking those same plant-feeders? Some limited evidence of host-associated divergence “cascading” across trophic levels exists (Stireman et al. 2006; Forbes et al. 2009), but its ubiquity is not known.

Oak gall wasps and their parasites provide an ideal host system to address whether host shifts commonly beget new diversity across trophic levels. In the Nearctic, an estimated 700 species of oak gall wasp (Melika et al. 2021) induce galls on >150 oak species (Hipp et al. 2018, Manos and Hipp 2021). Each gall wasp induces galls of unique and diagnostic morphology on specific plant organs of one or a few closely related oak tree species (Abrahamson et al. 2003; Ronquist et al. 2015; Egan et al. 2018). Phylogenetic study of the Nearctic gall wasps reveals that they have shifted to new host tree species and organ tissues numerous times and that these shifts correlate with lineage divergence (Ward et al. 2022a). Similar studies in the Palearctic, where there are fewer oak species, recover fewer shifts between host trees (Stone et al. 2009) but shifts between galls on different organs are still frequent (Cook et al. 2002). Population genetic studies of single oak gall wasp species using multiple host trees also suggest a role for host association in gall wasp diversification, with use of different tree species particularly strongly implicated in divergence (Hood et al. 2019; Zhang et al. 2019, 2021; Ward et al. 2022a).

Just as oak trees present a large set of actual and potential hosts for oak gall wasps, the collective set of galls induced by oak gall wasps represent a diversity of potential hosts for their parasites. Most oak gall wasp species have both a sexual and an asexual generation, each which induces a gall of a specific type on a specific organ of one or a limited range of oak species (Pujade-Villar et al. 2001, Egan et al. 2018; Ward et al. 2022a). Already variable in their tree host and location on those trees, the galls themselves vary greatly in traits shown to play a role in defense against parasitic wasp attack (Bailey et al. 2009). Some galls are smooth, others may have a light pubescence, long hairs, or even spines. Some produce nectar, which can both trap potential parasites and attract beneficial predators (Seibert, 1993; Nicholls et al. 2017). Galls can be as small as 1-2 mm or more than 120 mm in diameter (Weld 1957, 1959, 1960; Russo 2021). Some have their larval cells protected by bark, while in others the cells are suspended by fibers or are free-rolling in hollow chambers, all traits which may make it more challenging for parasites to oviposit (Stone et al. 2002). A single oak tree species can be host to many temporally and spatially co-occurring galls induced by different wasp species (Weld 1957, 1959, 1960; Jones et al. 2022) and oak tree species in many regions are broadly sympatric. Gall morphology has also changed substantially across the gall wasp phylogeny (Ronquist et al. 2015; Ward et al. 2022a). In all, a multi-dimensional diversity of oak galls – potential hosts for parasites – exists across time and space.

Confronted by diverse and evolving gall habitats, parasitic insects associated with oak galls might respond – or not – in a variety of different ways. If the presence of diverse oak gall hosts does not often lead to reproductive isolation among parasites associated with different galls, then the species diversity of gall-associated parasitic insect clades may be low, with each species attacking a relatively large number of gall types. A second scenario is that when gall wasps speciate, their ancestral parasites diverge in concert. In this case of “phylogenetic tracking” (Russo et al. 2017), one would predict high levels of diversity within clades of gall-associated parasites, along with a strong signal of cophylogeny between gall wasp and parasite clades. In a third scenario, wherein specialist parasites diverge after one shifts to a new gall environment, we would again expect high parasite diversity, with each species attacking a narrow range of gall hosts, but with phylogenies of gall wasps and their parasites largely decoupled. Importantly, in this third scenario, some phylogenetic tracking might still occur, since a parasite that tracks a host-shifting gall wasp would still be moving into a new (non-ancestral) tree and/or organ habitat.

We focus on two genera of kleptoparasitic wasp associates. Often described as inquilines, wasps in the genera *Synergus* Hartig, 1840 (Hymenoptera: Cynipidae) and *Ceroptres* Hartig, 1840 (Hymenoptera: Cynipidae) form secondary chambers in their host gall (Askew 1961; Evans 1965; Ronquist 1994) and feed primarily on the tissue of their host gall and not on the immature gall wasp. Some species form gall chambers in the original chamber of the gall inducer, resulting in the death of the gall wasp, while other species form peripheral chambers that may allow the gall inducer’s continued development (Evans 1965; Pénzes et al. 2012). In either case their co-option of some or all of the gall wasp’s resource makes them more parasitic than strictly inquilinous and thus we use the term kleptoparasite. Though both are in family Cynipidae, *Ceroptres* and *Synergus* are distantly related and their kleptoparasitic strategies have evolved independently (Ronquist et al. 2015; Blaimer et al. 2020). In North America, any single oak gall wasp species may be attacked by one or more species of one or both genera (Ward et al. 2020, 2022b). While both genera parasitize some of the same galls, analyses of rearing data show that they are significantly less likely to co-occur with one another than predicted by chance, suggesting that *Synergus* and *Ceroptres* might generally be adapted to different types of galls (Brookfield 1972; Ward et al. 2022b).

If host shifts have often contributed to speciation, parasites should be species-rich and ecologically specialized and their phylogenies should not track those of their hosts. We first test whether *Ceroptres* and *Synergus* are both comprised of many species, and whether individual species tend to specialize on dimensions of the gall niche. We delimit species and then reduce gall traits to principal coordinates and ask whether different kleptoparasite species occupy different areas of gall trait space. Finding that both genera are composed of diverse, specialized species, we then evaluate whether these two kleptoparasite genera have often tracked the diversification of their ancestral gall wasp hosts or if instead they have tended to shift among gall wasp lineages. Finally, we ask whether lineage divergences in these two kleptoparasites tend to be commonly associated with changes in particular gall locations (tree or organ) and/or gall morphology.

## Methods

### Collections and Sample Selection

We used *Synergus* and *Ceroptres* wasps previously reared from Nearctic oak galls (Ward et al. 2022b). Briefly, galls were collected from trees, put into plastic deli containers with mesh covers, and held in an incubator that simulated seasonal temperatures and day lengths. For 1-3 years, each container was checked daily and any insects that had emerged from galls were immediately put into individually labeled microcentrifuge tubes with 95% ethanol. Importantly, because all insects were reared as opposed to collected in the field as adults, their host gall, host tree, and geographic location are all known. A phylogeny of many of the host gall wasps from these collections was previously published (Ward et al. 2022a) and thus no additional genetic work was performed for the host insects.

Recent analyses of other North American gall parasites (Sheikh et al. 2022, Zhang et al. 2022), and previous study of *Synergus* (Ward et al. 2020) led us to hypothesize that both *Ceroptres* and *Synergus* might contain several unnamed and morphologically cryptic species, and thus we decided not to limit sample selection based on wasp morphology alone. To increase the likelihood of sampling the greatest number of species, we selected samples that maximized different combinations of morphology, tree host, host gall, and geographic location. For *Synergus*, we used 46 samples whose DNA had previously been extracted and which had already been shown to belong to a diverse set of putative species based on a combination of ecological and genetic (mtCOI) data (Ward et al. 2020). Specifically, we sampled 24 of the 27 clades reported in Ward et al. (2020). To these, we added 16 new specimens for a total of 62 individual *Synergus* wasps used for genetic work (Table S1).

For *Ceroptres*, we had no pre-existing species IDs for our collections, and attempts to key specimens using the most recent taxonomic revision (Lobato-Vila and Pujade-Villar 2019) did not consistently produce clear matches to named species. Thus, we opted to eschew formal species names for the purpose of this work. We selected adult *Ceroptres* wasps representing a diverse set of host galls, geographic locations, and oak tree species to maximize potential species diversity in our sampled set. A total of 59 *Ceroptres* wasps were used for primary genetic work (sequencing of Ultraconserved Elements [UCEs] and part of the mitochondrial COI [mtCOI] gene), with an additional 65 *Ceroptres* wasps used only for mtCOI sequencing, including one wasp reared from a cecidomyiid midge gall (Table S2).

*Synergus* and *Ceroptres* wasps used for UCE sequencing work were photographed (lateral habitus and fore wing) and then destructively sampled (Ward et al. 2020, Figures S1 – S69). *Ceroptres* samples used only for mtCOI sequencing were non-destructively sampled using a QIAGEN DNeasy Blood and Tissue Kit (QIAGEN, Valencia, CA, USA) and deposited as vouchers (Table S2). For destructively extracted *Synergus* specimens, when a second individual from the same collection existed, or an individual that shared the same mtCOI haplotype, we deposited that individual as a likely representative of the same species (Table S1).

### UCE sequencing

For 62 *Synergus* and 59 *Ceroptres* samples, we prepared libraries using Kapa HyperPlus v5.19 Kits (Kapa Biosystems Inc., Wilmington, MA, USA). We performed enzymatic fragmentation of the bead-cleaned DNA using reagents from the KAPA HyperPlus Kit for 15 minutes to obtain a mean fragment distribution of 300-500 bp. We then used a double-sided size select bead clean on the amplified library to remove fragments outside of the 300-500 bp range (SPRIselect User Guide PN B24965AA protocol). First, we performed a 0.65x clean to eliminate small fragments, then a 0.5x clean to eliminate large fragments, and a final 1.3x bead clean to elute target fragments into TLE buffer. We verified fragment distributions using an Agilent Model 2100 Bioanalyzer (Agilent, Santa Clara, CA, USA). We then pooled libraries and hybridized them following the MyBaits protocol (ArborBiosciences, Ann Arbor, MI) using the Hym v2P bait set (Branstetter et al. 2017). We confirmed UCE enrichment for pooled samples using relative qPCR. We then sequenced the entire pooled library on one lane of a NovaSeq6000 SP flowcell (2x150bp; Illumina, Inc. San Diego, CA) at the University of Iowa Institute of Human Genetics.

### UCE phylogenies

We used the Phyluce v1.7.0 pipeline (Faircloth, 2016) to process UCE data. Adapters were trimmed using illumiprocessor (Faircloth, 2013) and trimmomatic (Bolger et al. 2014), and assembled using SPAdes v3.14.0 (Bankevich et al. 2012). The assemblies were aligned using MAFFT v7.490 (Katoh & Toh, 2008), and trimmed using gblocks (Castresana, 2000) using the following settings: b1 = 0.5, b2 = 0.5, b3 = 12, b4 = 7. We selected the 90% complete matrices for *Synergus* and *Ceroptres* with 993 and 1019 loci, respectively. We included several cynipid outgroups (*Synophrus pilulae*, *Rhoophilus lowei*, *Diastrophus turgidus*, *Andricus quercuspetiolicola*, and *Neuroterus floccosus*) from other published UCE studies (Blaimer et al. 2020, Ward et al. 2022a). One *Ceroptres* and three *Synergus* samples from Blaimer et al. (2020) were also included.

We built phylogenetic trees for *Synergus* and *Ceroptres* using IQ-TREE v2.20 (Minh et al. 2020), first by using tests of symmetry (Naser-Khdour et al. 2019) to remove loci that failed the two assumptions of stationarity and homogeneity to generate matrices with 710 loci for *Synergus* and 763 for *Ceroptres*. Best models for each locus were then chosen by ModelFinder (Kalyaanamoorthy et al. 2017), and 1000 ultrafast bootstrap replicates for nodal support with >95% being considered strongly supported (UFB, Hoang et al. 2017). Summary statistics were calculated using AMAS (Borowiec 2016).

### mtCOI sequencing

We relied on Ward et al. (2020) for all *Synergus* mtCOI data. For 106 *Ceroptres* samples, including 41 of those used for UCEs, we amplified a ∼658bp segment of mtCOI using barcoded HCO2198-LCO1490 primers (Folmer 1994). Amplicons were multiplexed into two libraries and run on two Flongle flow cells with R9.4.1 chemistry on a MinION sequencing device (Oxford Nanopore, Oxford, UK). We demultiplexed sequences and resolved sequence consensuses using minibarcoder (Srivathsan et al. 2018). We re-sequenced some sequences that were not well-resolved from the MinION run on an ABI 3500 Genetic Analyzer (Thermo Fisher Scientific, Waltham, MA, USA). To supplement our new sequences, we added existing COI sequences from Nylander (2004), Ács et al. (2010), and Weinersmith et al. (2020).

### Species delimitation

Our approach to delimiting putative species relied on integration of UCE and COI data. For *Synergus*, species assignments had previously been made based on a combination of mitochondrial COI sequence data, morphology, and ecology (Ward et al. 2020). We did not seek to repeat previous COI work here but built upon it with new UCE sequence data (below). For *Ceroptres*, we had no pre-existing mtCOI analysis, and so employed Assemble Species by Automatic Partitioning (ASAP; Puillandre et al. 2021) to sort COI sequences from individual wasps into putative species that we could compare with UCE results. We then used a quartet frequencies-based species delimitation method (SODA; Rabiee and Mirarab 2020) to estimate *Synergus* and *Ceroptres* species from the UCE data (our samples only, plus *Ceroptres cornigera* from Blaimer et al. (2020)). Individual gene trees were generated using the best models selected from ModelFinder using IQ-TREE, and SODA was performed without using a guide tree. We compared SODA results with ASAP results, and, in instances where they disagreed, used collection information, ecology, and morphology to finalize species assignments (see Results).

### Ecology of Ceroptres and Synergus species

To determine how *Synergus* and *Ceroptres* wasps interact with gall niche space, we constructed a matrix of traits for all gall hosts of any *Synergus* or *Ceroptres* used in this study, as well as for some additional galls from which *Synergus* wasps were reared in Ward et al. (2020), and for *Callirhytis quercuscornigera*, the host of *C. cornigera* from Blaimer et al. (2020) (Table S3). We then made a second interaction trait matrix of all *Ceroptres* and *Synergus* species with traits of each of their respective host galls. Though they were not included in our UCE sequencing, we also included representatives of three additional *Synergus* species from Ward et al. (2020) (Table S1: *S. villosus B*, *S. campanula C*, and *S. oneratus D*). We excluded the one midge gall and its associated *Ceroptres* species, and one unknown gall for which we did not have gall traits recorded (one of the hosts of *C. sp. 15-16*).

To make the trait matrices, for each gall host, we scored traits that are putatively defensive against parasites using our own observations and those recorded on a database (Gallformers.org, 2023). We scored the presence of gall traits that may impose defense against parasites: spatially (i.e., galls are clustered, or have multiple chambers - ‘polythalamous’); through internal tissue or structures (i.e., if galls are hollow, have radiating fibers, or have fleshy or woody tissue, or tissue that changes from fleshy to woody with time) or external structures (i.e., if galls are shaped like a bract, if they produce nectar, or if the external tissue is textured, wooly hairy or spiny). Finally, we scored galls as 1 if small (0.5 - 3.9 mm), 2 if medium (4.9 - 9 mm, or 3 if large (> 10 mm) (Zhang et al. 2022, Prior et al. 2023).

We calculated Gower’s dissimilarity index between all gall host species in the gall trait matrix (Laliberte and Legendre 2010), and then performed a principal coordinates analysis (PCoA) and created a biplot with the first two components to project gall host species in gall trait space. We calculated centroids of gall host species that kleptoparasites interact with, for each kleptoparasite species, in gall trait space.

Centroids were plotted in a biplot, representing the suite of gall host traits that each parasite species interacts with (or their gall niche space) (Dehling et al. 2016, Prior et al. 2023). In one biplot parasite species were colored by genera to visualize how genera differentiate across gall host trait space. In other biplots, we colored species by different types of putative defenses (spatial organization, internal traits, external traits, size), to visualize suites of defenses that influence interactions (Zhang et al. 2022, Prior et al. 2023).

We performed hierarchical clustering analyses using the “ward.D2” method (Kaufman & Rousseeuw 2009) that maximizes the dissimilarity between groups and minimizes the dissimilarity within groups, using a Euclidean distance matrix calculated from kleptoparasite species centroids plotted in gall trait space. We created a cluster dendrogram, with the optimal number of groups. We then performed a regression analysis between the matrix of gall host species traits and the primary cluster affiliations to uncover which suite of traits significantly influence clustering patterns (de Bello et al. 2021). Finally, we used Mantel tests within each kleptoparasite genus to test for correlation between genetic distances and trait distance matrices.

We used the “vegan” (Oksanen et al. 2022), “labdsv” (Roberts 2023), “FD” (Laliberte et al. 2022), and “NbClust” (Charrad et al. 2022) packages in R version .4.3.0) (R Core Team, 2023) to perform the niche space analysis.

### Cophylogeny and ancestral state reconstruction

To test whether *Synergus* and *Ceroptres* track the phylogeny of their host gall wasps, we used Procrustean Approach to Cophylogeny (PACo; Balbuena et al. 2013) implemented in R version 4.2.2 (R Core Team, 2022). PACo tests whether parasite clades are dependent on host clades by using shape superposition of a parasite phylogeny upon a host phylogeny. We used our newly-generated *Synergus* and *Ceroptres* UCE phylogenies as parasite trees and the North American gall wasp phylogeny from Ward et al. (2022a) as a host tree. PACo requires that all hosts have an associated parasite, so we pruned the galler and kleptoparasite trees as necessary to remove species with no interactions. Gall wasp species attacked by >1 parasite species and parasite species attacking >1 gall wasp species were retained in the analyses. We used the square roots of eigenvalues produced from superimposition of patristic distances (Hutchinson et al. 2017) and set r = 0 to allow testing against randomized interactions while preserving row frequencies and species richness. We also used the *symmetric* argument, which enforces the assumption that parasites respond to the evolution of their hosts, and not vice-versa. As a complement to PACo, we used ParaFit (Legendre et al. 2002) to test whether the position of species in the topology of the parasite tree is dependent on the topology of the host tree. ParaFit produces both a global evaluation of cophylogeny as well as tests of the individual contributions of host-parasite pairs to an inferred cophylogenetic pattern.

To infer discrete histories of kleptoparasite host tracking and host switching events, we reconciled the galler phylogeny with both kleptoparasite phylogenies using eMPRess (Santichaivekin et al. 2021). Unlike other cost-based maximum parsimony reconciliation (MPR) approaches which require costs of phylogenetic events to be assigned before analysis, eMPRess models several different costs at once for four different event types (cospeciations [host tracking], duplications, transfers [host-shifts], and losses) and then allows the user to explore “event cost space” across multiple possible solutions. We again limited the gall wasp input tree only to species from which either *Ceroptres* or *Synergus* had been reared, and conversely limited the two kleptoparasite input trees to species for which at least one host gall wasp was represented in the Ward et al. (2022a) phylogeny. Parasite tips cannot be linked to >1 host tips in eMPRess, but instead of removing those interactions we duplicated relevant parasite tips within species that parasitized multiple gall hosts. For example, because *Synergus laeviventris* B was associated with four gall hosts, we divided it into a clade with four tips, with each tip associated with one of the four hosts. We considered events resulting from divided tips in final reconstructions to be intraspecific changes and subtracted these from final event counts.

We considered the biology of the gall system when evaluating event cost space. eMPRess considers a “cospeciation” event to be a “null event” and sets its cost to zero (Santichaivekin et al. 2021). The program also sets parasite loss at a unitless value of 1.0, with costs of duplications and transfers being relative to that 1.0 value. In the gall system, a gall wasp speciation event might often involve that gall wasp shifting to a new host plant or organ (Ward et al. 2022a). Thus, regardless of whether a kleptoparasite cospeciates with a host-shifting gall wasp or colonizes a new gall wasp host, either event requires the kleptoparasite to adapt to a new habitat. Therefore, we decided only to consider cost space where the cost of transfers was between 0 (equal to the pre-set cost of cospeciation) and 1.5 (moderately higher than cospeciation). We also *a priori* assumed that duplication events would be unlikely in this system because a duplication would put sister parasite species in direct competition with one another on the same host (Hamerlinck et al. 2016). Thus, we only considered solutions where duplications had a relatively high cost (>2). Finally, we only considered solutions that were strongly time consistent, i.e., that relied on transfers of parasites between contemporaneous hosts.

To visualize changes in habitat that have occurred coincident with kleptoparasite divergence, we mapped traits to the *Ceroptres* and *Synergus* trees and performed Ancestral State Reconstruction (ASR). Traits used were oak tree subsection (Manos and Hipp, 2021), host organ (leaf, stem, bud, flower, fruit, petiole), and generation (galls of the sexual vs. asexual generation). To model evolution of use of different gall morphologies, we used each species’ assignment to one of eight clusters emergent from hierarchical clustering analysis (see results). Species reared from >1 galls with different traits were treated as polymorphic for those traits. We used a maximum likelihood approach in the R package ape v.5.7-1 (Paradis & Schliep 2019) and tested three unordered models of trait evolution: “equal rates”, “symmetrical, or “all rates different”. We performed multiple tests for each trait and chose the model with the lowest Akaike’s Information Criterion (AIC) value. An equal rates model was the most highly supported for ancestral state reconstruction for all traits tested, but the AIC for the symmetrical model was not strongly different so we chose this one to maximize model flexibility. Because we were modeling changes in discrete traits, we made trait changes independent of phylogenetic branch lengths by using a punctuated speciation model (kappa = 0; Pagel 1999). Ancestral character estimation was completed using joint ancestral state methods to reconstruct the complete history of changes by using a marginal = false argument (Holland et al. 2020). Lastly, polymorphic traits were proportionally separated into their respective discrete components following calculation of log likelihood of ancestral state.

We used ASRs to tally overall numbers of trait changes in each of the *Synergus* and *Ceroptres* phylogenies. We counted a node as coincident with a shift to a new trait (tree, organ, morphology, or generation) when the trait assigned the highest proportion at that node differed from the trait at the highest proportion at one or both daughter nodes. When no trait was clearly highest in proportion at a node (i.e., when a node was approximately evenly split between two or more traits), that node was only counted as a shift if different traits were at highest proportions in its two respective daughter nodes.

## Results

### Phylogenies and Species Delimitation

The filtered UCE 90% *Ceroptres* matrix included 65 taxa and 763 loci, with the total length being 534,960 bp, with 196,557 variable sites (36.7%), 69,712 parsimony-informative sites (13%), and 16.3% missing data. Similarly, the concatenated UCE 90% matrix for *Synergus* included 71 taxa and 710 loci, with the total length being 549,203 bp, with 254,143 variable sites (46.3%), 132,152 parsimony-informative sites (24.1%), and 17.7% missing data. The *Synergus* UCE phylogeny had >95% support at most nodes, with the exception of the node connecting the *S. laeviventris* clade from the rest of the tree (91%) and some within-species nodes (Figure S70). Most support values for the *Ceroptres* phylogeny were also >95%, but one large clade had several nodes with support <80% (Figure S71). The COI data for *Ceroptres* included 111 taxa at 658bp, with 277 variable sites (42.1%), 241 parsimony-informative sites (36.6%), and 0.03% missing data.

Previous work has suggested that *Synergus* may not be a monophyletic grouping (Blaimer et al. 2020, Lobato-Vila et al. 2022). While our UCE data from genera supposedly paraphyletic with *Synergus* were limited to one *Synorphus* individual (from Blaimer et al. 2020), we here recover genus *Synergus* as a single clade (Figure S70). We thus continue as if *Synergus* is monophyletic, but we note that with more data this may prove not to be the case.

SODA produced a hypothesis of 30 putative species of *Synergus* among the individuals for which UCEs were sequenced (Table S1). Synthesizing these estimates with COI species assignments from Ward et al. (2020), we find evidence for 28 putative species among those in our *Synergus* collections (Figure 1; Table S1). Our samples included individuals from 24 clades / putative species previously resolved using only COI (Ward et al. 2020), as well as several individuals that had not previously been analyzed. In each case of disagreement between COI and SODA, we adopted the more conservative estimate. Thus, in two instances, we followed the recommendation from SODA to combine clades 23 & 25 and clades 21 & 22 from the Ward et al. (2020) study. Similarly, in two cases (*Synergus magnus* and *Synergus laeviventris B* from Ward et al. 2020) where SODA suggested that single species estimated from COI data should be split, we did not split them. Retaining *Synergus magnus* as a single species seemed justified since all three samples were reared from the same host gall type in the same city and it thus seemed unlikely that they represent two reproductively isolated species in ecological and geographic sympatry. Similarly, we did not split *S. laeviventris B* because samples were reared from *Amphibolips* galls in Iowa versus Ohio/Pennsylvania, and we thought that COI haplotype differences between these samples likely to reflect isolation by distance and not the consequence of evolved reproductive isolating barriers. For all 18 remaining Ward et al. (2020) species, COI and UCE estimates agreed. Lastly, from among the new samples that had not been part of the previous study, SODA suggested six additional species. One of these matched the description of *Synergus ochreus*, while the others did not immediately fit an existing description, so we here call them *Synergus* sp. 4-8 (Figure 1).

**Figure 1.**
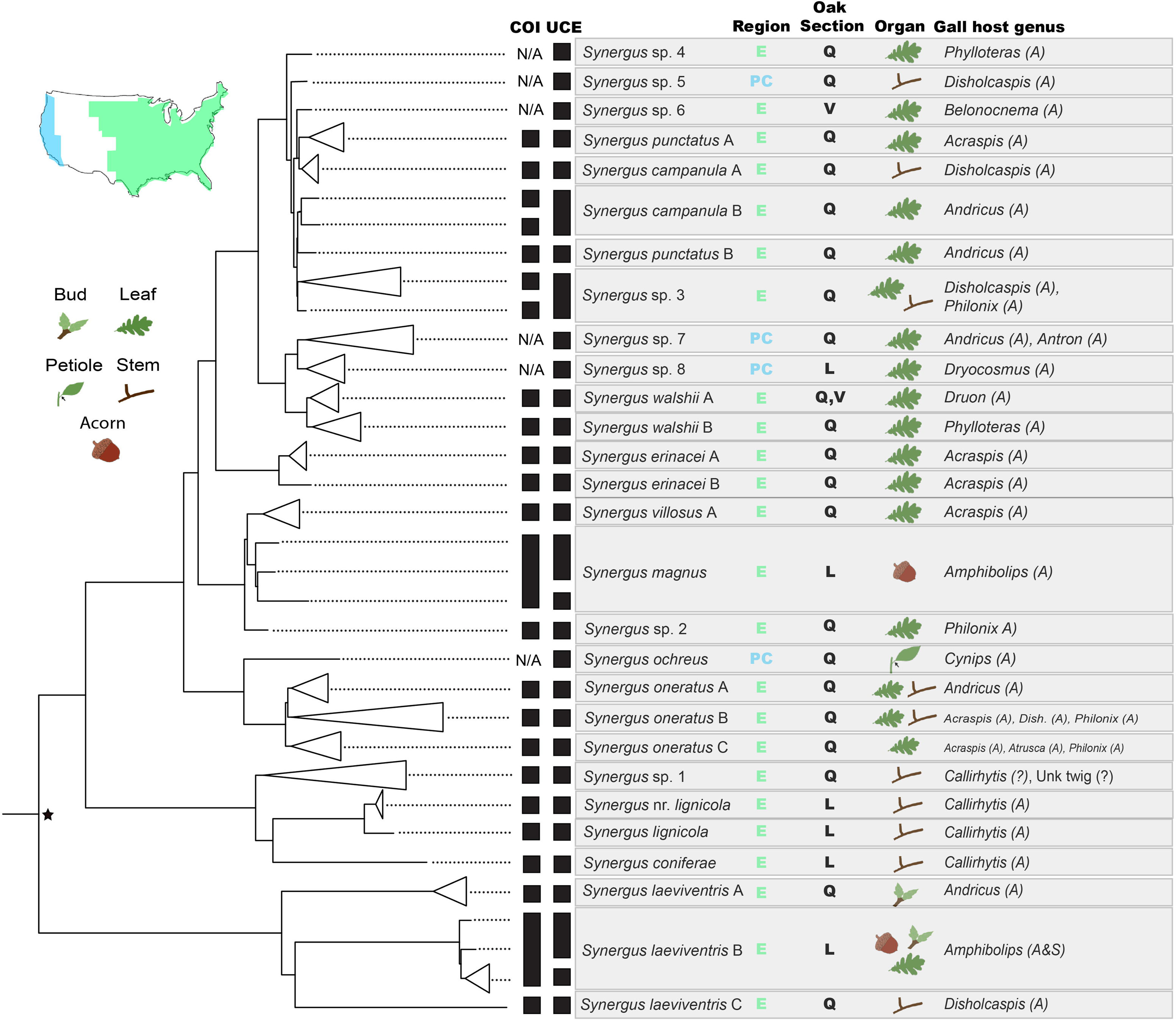
Summary of UCE phylogeny and molecular species delimitation for Nearctic Synergus wasps. Topology of the phylogeny comes from Figure S70, with outgroups and other samples from Blaimer et al. (2020) removed. Black star indicates support value <95% (Figure S70). Black bars indicate wasps assigned to the same species based on COI or UCE data. Gray boxes collect (from left to right) final species assignments, their geographic region, host oak section (Q=Quercus, L=Lobatae, V=Virentes), the organ(s) on which their host gall is located, the genus/genera of host gall inducers and the reproductive mode(s) of the gall wasp generation developing inside the host gall(s): A = gall of asexual generation, S = gall of sexual generation,= gall of unknown generation.

For *Ceroptres,* we relied primarily on molecular species hypotheses. SODA (based on the UCE phylogeny; Figure S71) suggested 36 species (numbered 0 - 35 in Table S2), while ASAP (based on COI sequences) suggested 47 species (numbered 1 - 47 in Table S2). In five cases, we lumped together UCE-based species based on shared ASAP assignments and/or because closely related individuals reared from the same hosts had been assigned to different species. We named these putative species by combining numbers from their SODA species assignments (*e.g., C.* sp. 7-8-9). In two cases, we also combined 2-3 species into one (*C*. sp. 26-27-28; *C*. sp. 15-16) based on their polyphyly on the UCE phylogeny and/or a shared gall host. In three other instances, (*Ceroptres* sp. 5, *C*. sp. 6, and *C*. sp. 30) COI/ASAP recommended that UCE-based *Ceroptres* species be split. Unlike for the instances of ASAP-SODA disagreements for *Synergus*, where in each case conspecific status was implied by a shared host gall, for instances of disagreement in *Ceroptres* the different ASAP assignments corresponded with differences in gall host. Because the host ranges of most other putative *Ceroptres* species were limited to one gall species or genus, we decided to follow the pattern emergent in the rest of our data and adopt the splits suggested by COI/ASAP. We discuss possible impacts of our decision to variously lump or split species in the Discussion. *Ceroptres* specimens that were not sequenced for UCEs but that were grouped as distinct COI/ASAP species were assigned to UCE-based species if they formed a well-supported haplotype clade with an individual from an UCE-based species and shared the same host gall or had been reared from the same collections. Eleven others were given distinct names (e.g., “*Ceroptres* sp. COI-A”) so they could be noted as potential additional species. The eleven species based only on COI sequences were not used in our cophylogenetic analyses and thus these decisions did not impact most conclusions in this study. However, all putative species were used in describing ecological dimensions of the gall host habitat.

### Ecology

Species from both kleptoparasite wasp genera were specialized on particular dimensions of their hosts. Across all *Synergus* species, 48.4% (13/31) were reared from galls of just one gall wasp species, and no *Synergus* species was reared from galls of more than four gall wasp species (Table S1). Only one species (*Synergus walshii* A) was reared from galls on two different oak sections, while all others were specialized on a single oak section. Most *Synergus* species (23/31; 74.2%) were associated with galls that shared the same plant organ, with only one *Synergus* species (*S. laeviventris* B) being reared from galls on three different plant organs. All but one *Synergus* species were reared from galls of the asexual generation. The exception was again *S. laeviventris* B, which was reared from both sexual and asexual galls (Figure 1).

*Ceroptres* species were similarly specialized on their respective gall hosts. Across all *Ceroptres* species (inclusive of COI-based species, but omitting the single species reared from a midge gall, 76.3% (29/38) of species were reared from galls of only one gall wasp species (Table S2). Only one (2.6%) *Ceroptres* species was reared from galls of more than three gall wasp species. All *Ceroptres* species were reared from galls formed on just one oak section and most (35/38; 92.1%) *Ceroptres* species were associated with galls on a single plant organ; just one species was reared from galls on three different organs (Figure 2). *Ceroptres* species were reared from either sexual or asexual galls, and two *Ceroptres* species were reared from galls of both sex and asex generations, though these were not of the same gall wasp species.

**Figure 2.**
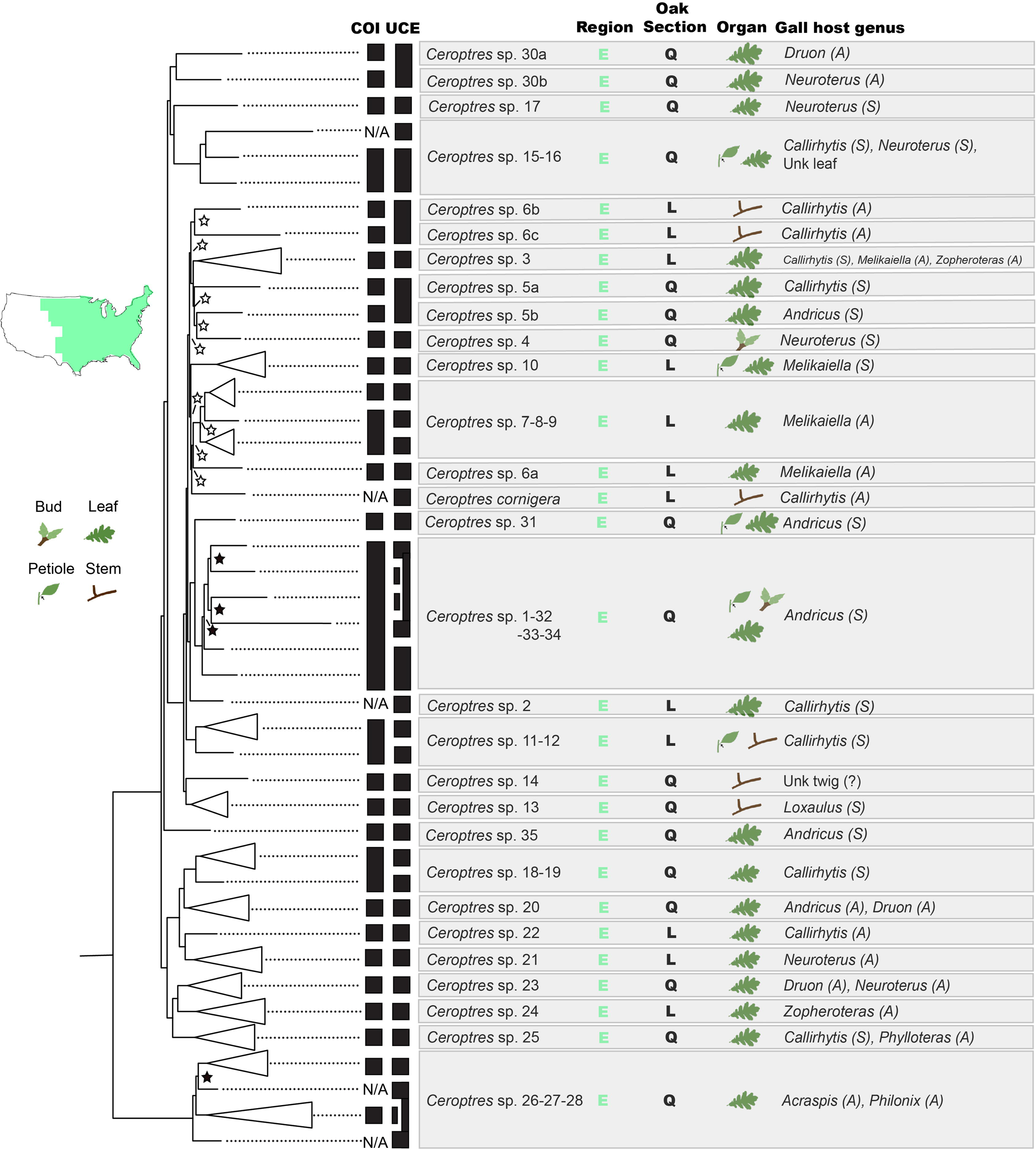
Summary of UCE phylogeny and molecular species delimitation for Nearctic Ceroptres wasps. Topology of the phylogeny comes from Figure S71, with outgroups removed. Black stars indicate support <95%, white stars support <80%. All other nodes have >95% support (Figure S71). C. cornigera was not collected in our study (Blaimer et al. 2020) but was included here because the host gall was known. Gray boxes collect (from left to right) final species assignments, their geographic region, host oak section (Q=Quercus, L=Lobatae), the organ(s) on which their host gall is located, the genus/genera of host gall inducers and the reproductive mode(s) of the gall wasp generation developing inside the host gall(s): A = gall of asexual generation, S = gall of sexual generation,= gall of unknown generation.

Hierarchical cluster analysis of *Ceroptres* and *Synergus* species mapped in gall trait space resolved two primary clusters (Figures 3, S72). One cluster (Cluster A) consisted of 21 *Ceroptres* species and 2 *Synergus* species. Kleptoparasite species in this cluster were associated primarily with medium-to-large polythalamous (multi-chambered) woody or fleshy galls with few external defenses. The two *Synergus* species in this cluster were *S. lignicola* (only reared from *Callirhytis quercuspunctata*, a woody, polythalamous gall) and *S. punctatus* A (reared only from *Acraspis erinacei*, a fleshy, polythalamous gall). Two other species, *Synergus erinacei* A and *S. oneratus* C were both also reared from *A. erinacei* but were also reared from other galls on leaves of white oaks. A second primary cluster (17 *Ceroptres* species, 29 *Synergus* species) were generally reared from unilocular galls that were single or in clusters that are medium to small, not woody, and with a variety of other features (Figures 3, S72). Secondary cluster analyses within Cluster A revealed six additional clusters representing groups interacting with either woody, large galls with some external defenses or fleshy, medium galls, with fewer external defenses. Cluster B was separated into two additional clusters interacting largely with unilocular clustered vs. single galls (Figure S72). A regression between kleptoparasite species in gall trait space and gall traits that took into account clusters suggested that eight variables significantly influence clustering (Table S4), with spatial traits (polythalamous / unilocular / clustered) explaining the most variation, followed by internal traits, size, and then external traits (Figure S73). Mantel tests showed no significant correlation between genetic distance and trait distance matrices for *Synergus* (r = 0.01968, sig = 0.389) or *Ceroptres* (r = −0.03342, sig = 0.547) species.

**Figure 3.**
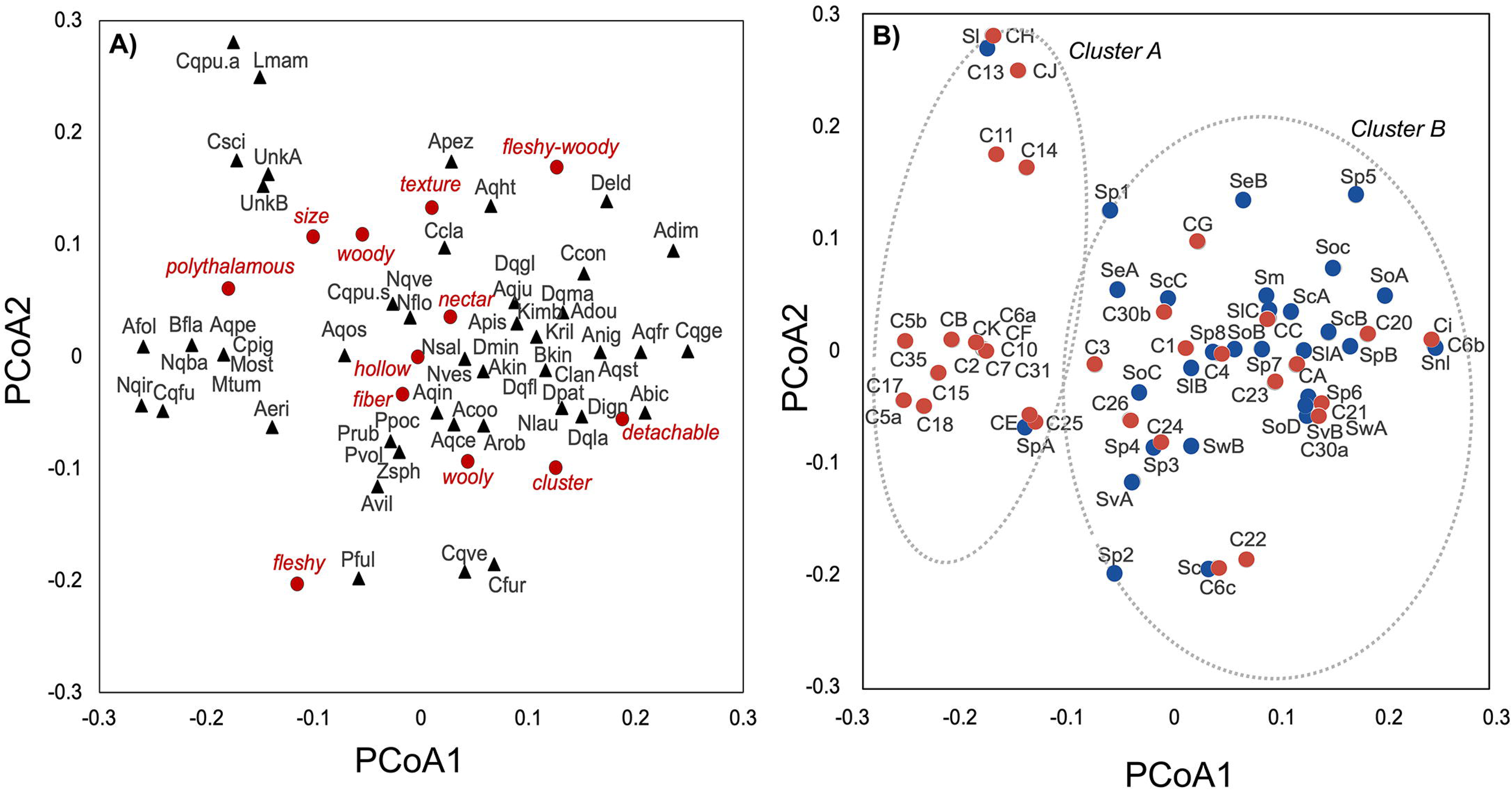
Results of principal coordinate analyses. A) Biplot of PCoA1 and PCoA2 for all host gall species (black triangles) plotted in gall trait space. Red circles and text denote trait loadings. B) Kleptoparasite species plotted in centroids of trait space of interacting host gall species. Blue circles = Synergus, Red circles = Ceroptres. Ellipses marked with dotted outlines denote the two primary clusters (A,B) identified via hierarchical clustering analyses. See Table S3 for host gall and kleptoparasite species abbreviations and figures S72 and S73 for additional clusters and information on traits of associated galls.

### Cophylogeny

Relationships between kleptoparasite and galler phylogenies were significantly different from random (Figures S74, S75). With PACo, we detected a significant signal that at least some part of the *Synergus* phylogeny’s ordination is constrained by the ordination of the gall wasp phylogeny (m^2^_xy_ = 0.650; P < 0.001). ParaFit also found a significant global relationship between the gall wasp and *Synergus* phylogenies (ParaFitGlobal statistic = 0.030, P = 0.023). However, only 6/34 interactions were implicated by ParaFit as contributing to the relationship between phylogenies (ParaFitLink1 and Link2 statistics, Table S5). Four of these six interactions involved the same *Synergus* species (*S. laeviventris* B) which was reared from all four *Amphibolips* species of gall wasps in the study. The contribution of these interactions to a signal of apparent cophylogeny thus reflects only that *S. laeviventris* attacks multiple, closely related host species and that they have not cospeciated with those hosts. Thus, only 2/34 interactions were identified as possibly contributing to a cophylogenetic pattern between the phylogenies. eMPRess rejected a hypothesis that gall wasp and *Synergus* phylogenies were concordant (P < 0.01).

PACo also detects a significant signal that at least some part of the *Ceroptres* phylogeny is constrained by the gall wasp phylogeny (m^2^_xy_ = 0.056; P < 0.001). However, ParaFit does not find evidence for a significant global relationship between these two phylogenies (ParaFitGlobal statistic = 0.010, P = 0.891) and only one interaction (*C.* sp. 30b and *Neuroterus quercusverrucarum*) was implicated as having a significant, potentially coevolutionary link (Table S6). eMPRess rejected a hypothesis that gall wasp and *Ceroptres* phylogenies were concordant (P < 0.01).

Within our predetermined ranges of event cost parameters, eMPRess produced just two solution sets for *Synergus*, one where the cost of a host shift event relative to the cost of a loss event ranged from 0-1, and the other where the relative cost of a host shift ranged from 1-2. Median solutions for both sets were strongly time consistent and were similar to one another in their reconstruction of events. After subtraction of intraspecific events (white shapes, Figure 4), one solution had 16 host shifts, four cospeciations (Figure 4), and no losses, while the other solution had 15 host shifts, five cospeciations, and one loss event (Figure S76). Neither solution had any duplication events. Thus, in these respective solution sets, 80.0% and 75% of all divergence events for *Synergus* were inferred to be coincident with host shifts.

**Figure 4.**
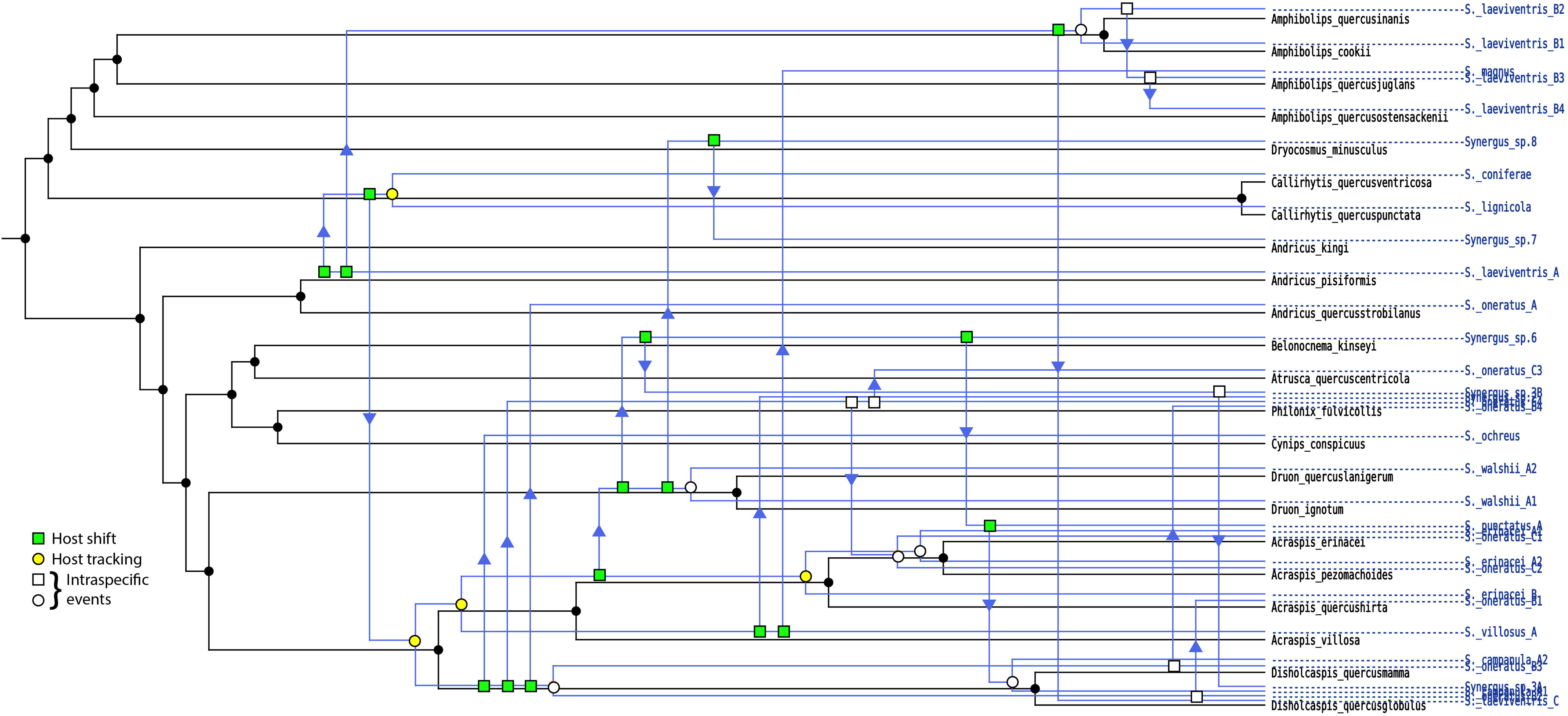
One of two median eMPRess solutions for the cophylogeny of Synergus parasites with their gall wasp hosts. Black phylogeny and labels are host gall wasps; blue phylogeny and labels are Synergus species. Green squares show inferred host shifts coincident with Synergus divergence events, while yellow circles show inferred host tracking events. White shapes represent intraspecific events. Event costs for this solution space were as follows: cospeciation (0), duplication (>2), transfer (0-1), loss (1). With intraspecific events subtracted, this solution produced four cospeciation events and 16 host shifting events.

For *Ceroptres*, eMPRess also produced two solution sets. In the first set the cost of a host shift event relative to the cost of a loss event ranged from 0-1, and in the second set the relative cost of a host shift ranged from 1-2. Median solutions for both sets were strongly time consistent and were similar to one another in reconstruction of events. After subtraction of intraspecific events, the median solution for the first set (Figure 5) had four cospeciations, two duplications, 16 host shifts, and no losses, while the median solution for the second set (Figure S77) had five cospeciations, two duplications, 15 host shifts, and one loss. Thus, in these respective solution sets, 72.7% and 68.2% of all divergence events for *Ceroptres* were inferred to be host shifts. The two duplication events resolved in both solutions were the split between *C*. sp. 1-32-33-34 and *C*. sp. 31 and the split between *C.* sp. 6a and *C.* sp. 7-8-9. These may be real duplications but could also be a signal of oversplitting in *Ceroptres* species assignments.

**Figure 5.**
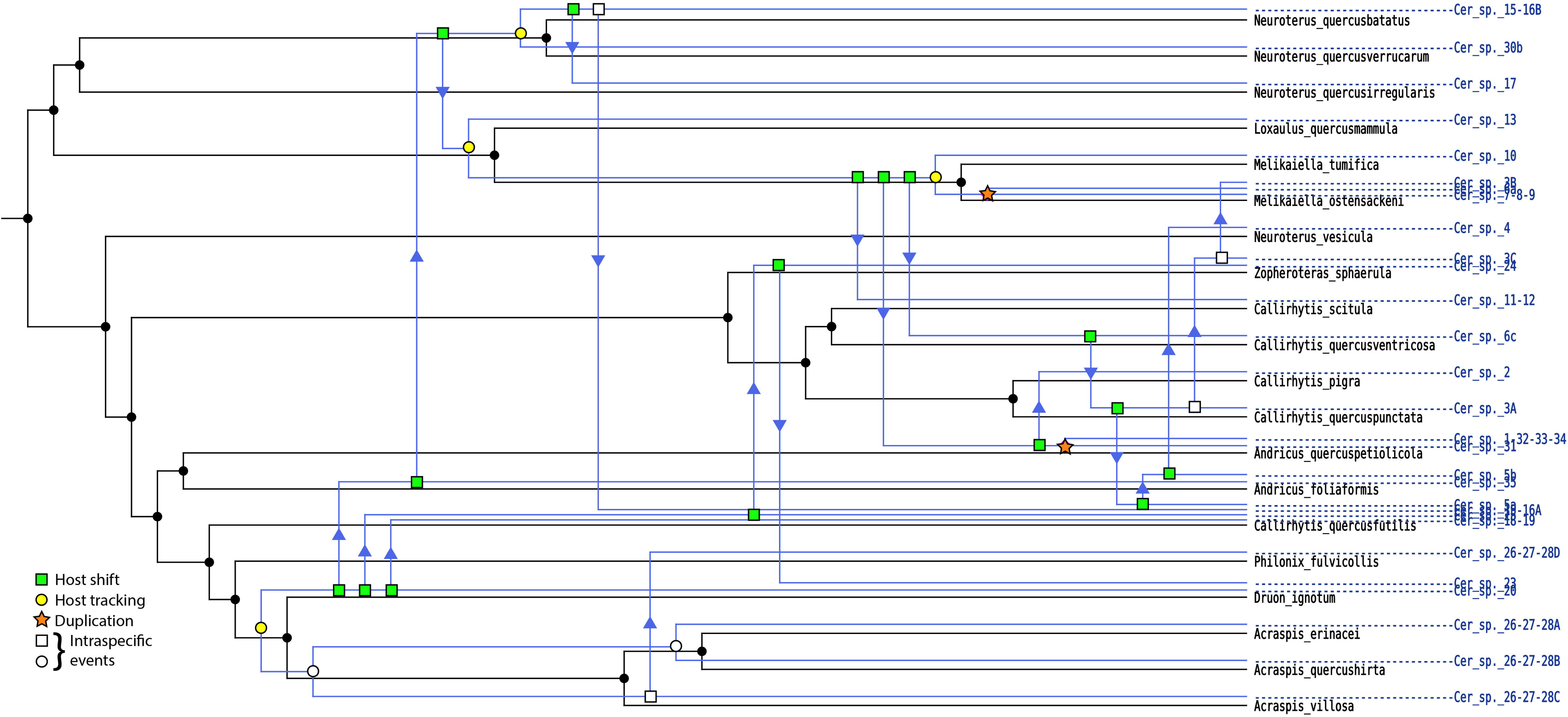
One of two median eMPRess solutions for the cophylogeny of Ceroptres parasites with their gall wasp hosts. Black phylogeny and labels are host gall wasps; blue phylogeny and labels are Ceroptres species. Green squares show inferred host shifts coincident with Ceroptres divergence events, while yellow circles show inferred host tracking events. Orange stars denote inferred duplications. White shapes represent intraspecific events. Event costs for this solution space were as follows: cospeciation (0), duplication (>2), transfer (0-1), loss (1). With intraspecific events subtracted, this solution produced four cospeciation events, 15 host shifting events, and two duplications.

Ancestral state reconstructions (ASRs) for *Synergus* show at least 14 shifts among oak subsections, six shifts among tree organs, and nine among morphogroups of galls defined by hierarchical trait clustering (Figure 6). Just six of 27 nodes showed no change in tree subsection, organ, or morphology. ASRs did not suggest any shifts in gall generation for *Synergus* – all species were associated with asexual galls, with only *S. laeviventris* B attacking both asexual and sexual generation galls.

**Figure 6.**
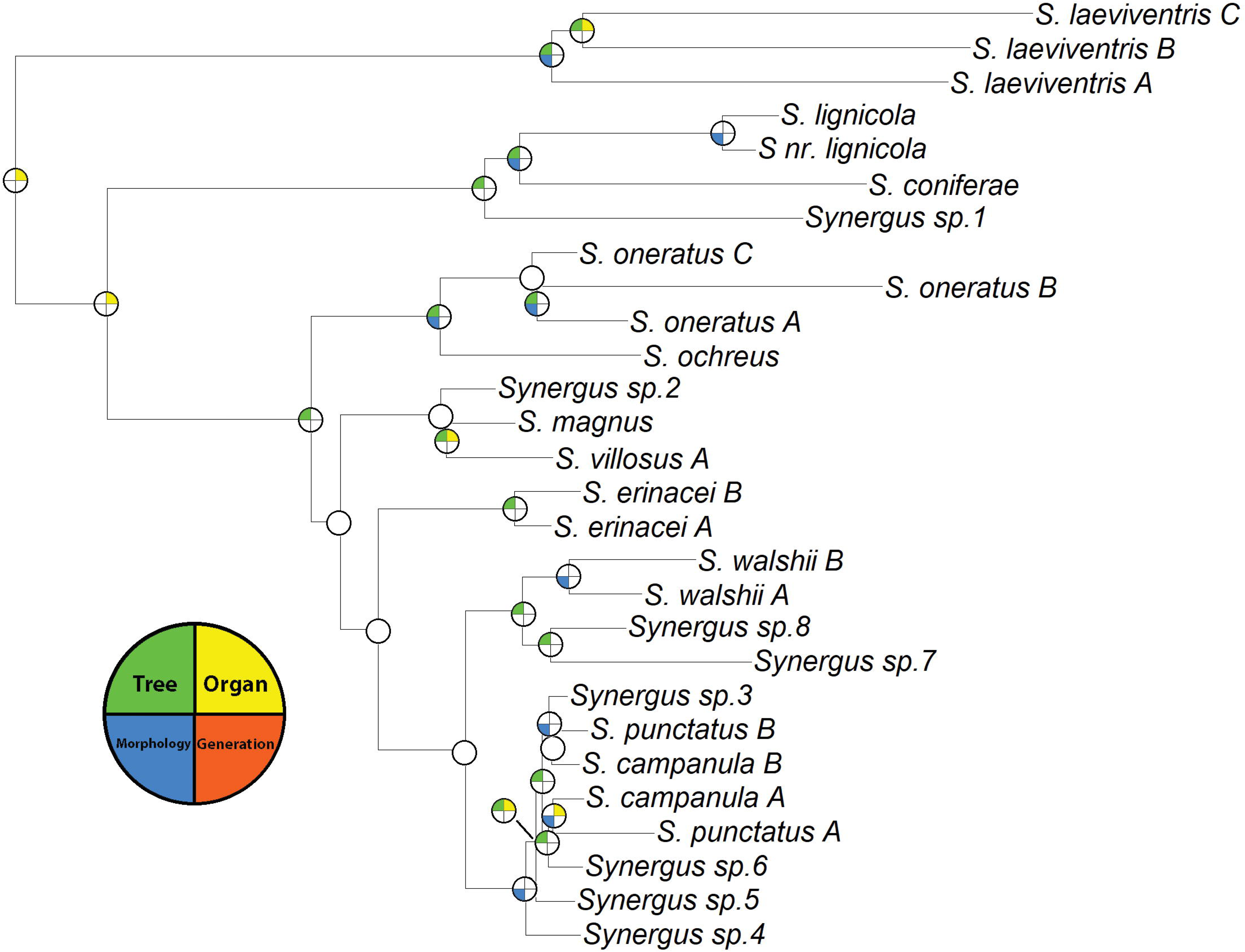
Summary of Ancestral State Reconstructions (ASRs) for Synergus wasps. Grey color in a quarter of a circle indicates change in state inferred at that node (see legend, bottom left). Circles not divided into quarters had no inferred changes in tree subsection, organ, morphology, or gall generation. See Figures S78-S81 for ASRs of individual traits.

ASRs for *Ceroptres* suggested 18 shifts among oak subsections, four shifts among tree organs, 16 among galls assigned to different gall morphogroups, and six shifts between galls of different generations (Figure 7). Four of 27 nodes showed no change in tree subsection, organ, morphology, or generation. Two of the nodes with no inferred changes were deep internal nodes. Of the other two, one was the node between *C.* sp.6a and *C*. sp. 7-8-9, while the other was the node between *C.* sp. 5a and the clade collecting *C*. sp. 5b and *C.* sp. 4. *Ceroptres* sp. 5 and *C*. sp. 4 were two of the three UCE-based species that we decided to split based on COI data, so these may again represent instances of over-splitting. The sister species *C.* sp.6a and *C*. sp. 7-8-9 both attack galls of *Melikaiella ostensackeni* on some of the same tree hosts, which makes oversplitting seem more likely.

**Figure 7.**
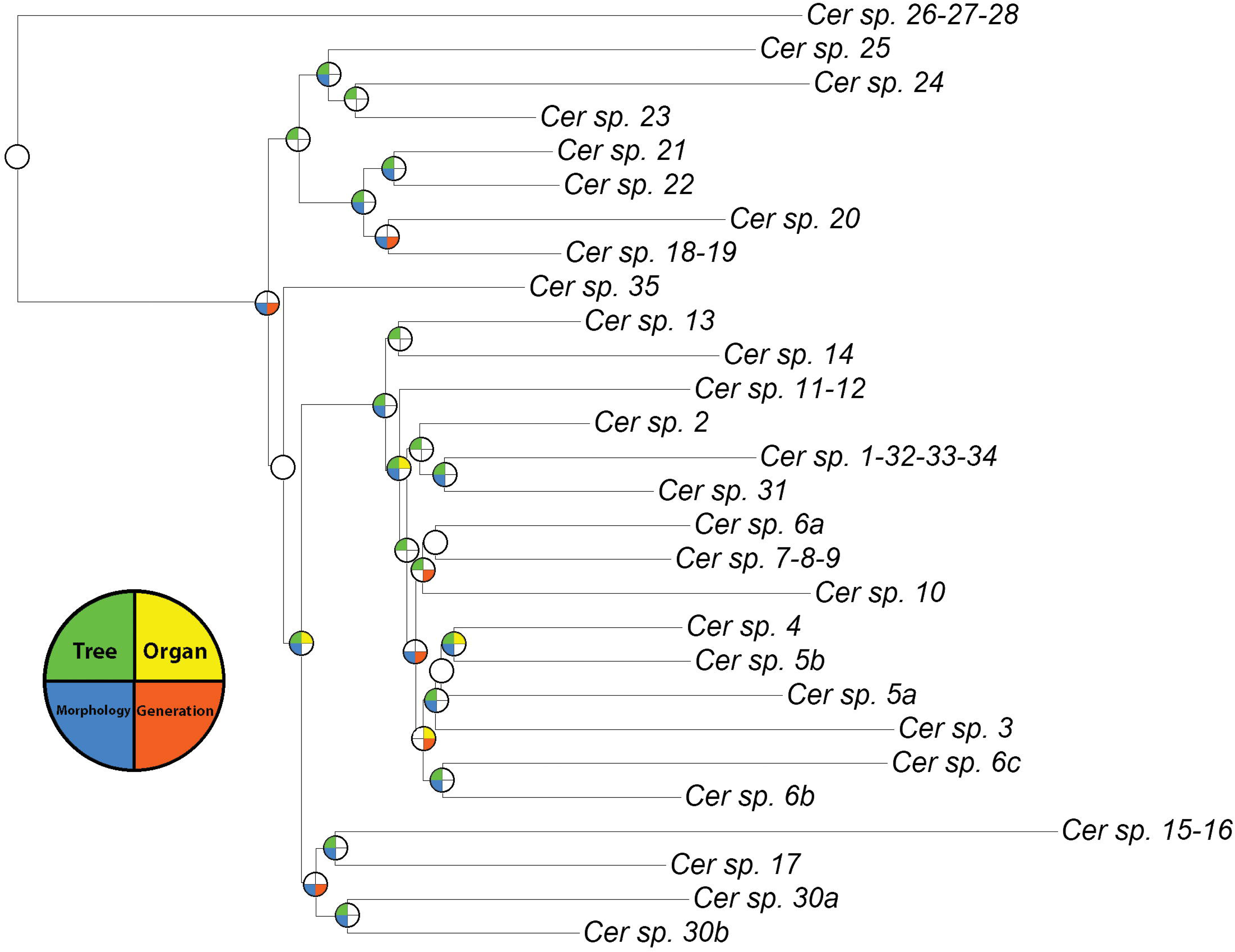
Summary of Ancestral State Reconstructions (ASRs) for Ceroptres wasps. Grey color in a quarter of a circle indicates change in state inferred at that node (see legend, bottom left). Circles not divided into quarters had no inferred changes in tree subsection, organ, morphology, or gall generation. See Figures S82-S85 for ASRs of individual traits.

## Discussion

### Nearctic Synergus and Ceroptres are species-rich host specialists

Shifts to new host galls may commonly result in gall parasite speciation. Underlying this hypothesis is the idea that parasites attacking galls of different types, and/or on different trees or organs, could experience divergent selection in the context of those different habitats, and that consequent adaptations may result in reproductive isolation. If this is true for parasites of galls, one would predict that each parasite species would specialize on only a limited number of host galls. And indeed, we find that *Ceroptres* and *Synergus*, the two kleptoparasitic wasp genera most commonly associated with Nearctic oak galls, are each comprised of a large number of species, with each species tending to be specialized on just one or a small number of oak gall types (Figures 1-2). While this pattern of host specialization had been suggested before for Nearctic *Synergus* based on COI data (Ward et al. 2020), we have now resolved the same patterns with a more robust, multi-locus genetic dataset for *Synergus*, as well as for a second genus, *Ceroptres*.

### Host shifting and sources of divergent selection

We also find that both kleptoparasite genera have frequently shifted to new host galls. Though we detected some signal of phylogenetic tracking between gall wasps and these parasites, these events were infrequent. Instead, lineage divergence in both kleptoparasite genera is commonly linked with shifts to new gall wasp associations. For *Synergus*, eMPRess reconstructions suggested >71% of divergence events were coincident with shifts to new hosts, while >68% of *Ceroptres* divergences were inferred to be host shifts (Figures 4, 5, Sx, Sy).

Frequent shifts to new hosts may portend a role for divergent ecological selection in speciation for *Synergus* and *Ceroptres*. In other insect systems, insects using different plants can develop strong assortative mating when plant habitats also act as mating sites (habitat isolation: Matsubayashi et al. 2010, Forbes et al. 2017). For kleptoparasites of oak galls, habitats relevant to the evolution of reproductive isolation might include the tree but could also include the tree organ or even the gall itself (Miller and Raman 2019). More than 62% (17/27) of divergence events for *Synergus* and >70% (19/27) for *Ceroptres* were correlated with a major change in the location of the gall (oak tree subsection, organ, or both). If changes in gall morphology (defined as association with one of eight morphogroups) are included, then >77% of *Synergus* and >85% of *Ceroptres* divergence events involve shifting to new habitats. Even in the relatively rare cases where a gall wasp and its kleptoparasite appear to have speciated in concert (“phylogenetic tracking” events), aspects of the gall environment often changed, such that the parasites often experienced a “habitat shift” even if they did not shift to a new gall wasp lineage. Shifts for both parasite genera were most common between oak subsections, followed by shifts among gall morphologies. Shifts between tree organs and generations were least common in both genera.

Any change in habitat (tree, organ, gall morphology) might result in reproductive isolation if mate choice for parasites is linked to the host gall. Mating for at least some *Synergus* species is known to occur at or near the site of female oviposition: male and female *Synergus pacificus* wasps parasitizing galls of the leaf gall *Besbicus mirabilis* mate on leaves of *Quercus garryana*, which are also the locations of their host galls (Evans et al. 1965). In a more extreme case of gall-associated mating, unidentified species of *Synergus* emerging from the same host gall mated upon emergence, and it was noted that they did not mate again for >1 week (Ikai and Hijii 2007). The maximum longevity for these *Synergus* was 10 days, such that natal-gall-associated mating may represent the only opportunities for sex in these wasps’ short lives. These accounts of host-gall associated mating are a strong indicator that changes in host could quickly translate into reproductive isolation, but studies of mating behavior for additional species in both genera are needed.

Reproductive isolating barriers emergent from host association need not be strictly due to habitat isolation. Temporal, or allochronic, isolation can result in greatly reduced gene flow between insects using different hosts (Taylor and Friesen 2017). This can occur when the host plants (or host insects) are susceptible to attack at different times during the year, such that divergent selection results in asymmetric evolution of parasite life history timing. For short-lived insects, even small temporal differences between populations can generate considerable allochronic isolation (Powell et al. 2014), and such isolation has recently been demonstrated both in oak gall wasps (Hood et al. 2019) and in their *Synergus* parasites (Zhang et al. 2019). Parasite shifts between galls induced by gall wasps of different generations could result in temporal isolation, as sexual generation galls tend to develop in the spring and early summer, while asexual generation galls develop in the late summer of fall. Temporal isolation in this manner may be more relevant to *Ceroptres*, which have shifted between gall generations multiple times (Figure 7), than to *Synergus,* which we have reared almost exclusively from asexual galls (Figure 1). Other potential ecological barriers might evolve from divergent adaptations to specific gall defenses, which could lead to ecological isolation between different host-associated populations via immigrant inviability or ecological inviability of hybrids. Both of these latter reproductive isolating barriers have been demonstrated to reduce gene flow between gall wasps using different tree hosts (Zhang et al. 2017, 2021b) but have not been measured in gall parasites.

### Caveats and Conclusions

Delimiting species based solely on molecular data has potential pitfalls, including introgression (Llopart et al. 2005) or incomplete lineage sorting (Hebert et al. 2013) leading to groupings that do not accurately identify reproductively isolated lineages. Our decisions to variously lump or split individuals into species could therefore have consequences for conclusions about specific evolutionary relationships. In general, our methods tended to use the most conservative of SODA or ASAP estimates, such that our primary concern would seem to be that we have lumped one or more specialized species into single, apparently oligophagous species. On the other hand, the resolution of an apparent duplication in eMPRess results for *Ceroptres* leads us to suspect that *C.* sp. 1-32-33-34 and *C.* sp. 31 are likely a single species that was split by our analyses. While erroneous lumping or splitting would affect the total number of inferred lineage splits in a genus and potentially influence specific species’ positions in gall space, inferences about ancestral changes in host associations, including how frequently ancestral nodes coincide with host shifts, should not be much affected. Resolution of species limits for these wasps will be a continuing process that ultimately requires an integrative taxonomic approach.

We do not assess gall wasp “escape” from parasites in this study, but this is another strong suggestion that there are areas of gall niche space that specific parasite genera do not (or only rarely) traverse (Price et al. 1987; Bailey et al. 2009; Ward et al. 2019). Previous work had suggested that *Synergus* and *Ceroptres* may not attack galls induced by the same gall-inducing wasp (Brookfield 1972). This is false in a strict sense: species from both genera were reared from galls induced by the same gall formers. However, we do show that *Ceroptres* and *Synergus* often occupy different gall niche space. Polythalamous galls were much more likely to have *Ceroptres* parasites than *Synergus* parasites, with only two *Synergus* species associated with the clusters of gall hosts principally defined by their polythalamy (Figure 3, clusters A1, A6). While we have not yet systematically studied intra-gall parasite interactions, rearing experiments and some preliminary micro-CT scans of galls suggest that *Ceroptres* may often take over existing gall wasp chambers (GEB and AAF, personal obs.). Polythalamous galls might thus be more attractive to *Ceroptres* than to *Synergus*, simply because they offer multiple chambers.

Though direct tests of the causes of reproductive isolation require broad experimentation at an intraspecific level, the current study and other recent results make it clear that host shifting is both common and correlated with diversification for both North American oak gall wasps (Ward et al. 2022a) and at least two of their associated kleptoparasite genera (Ward et al. 2020, this study). Other work shows that host associated genetic differentiation is associated with both temporal isolation and host association for some oak gall wasps and their *Synergus* parasites (Zhang et al. 2017, 2021b). Further, though comparisons with the gall-wasp phylogeny are pending, genetic studies of two other parasite genera that are broadly associated with North American oak galls, *Ormyrus* and *Sycophila*, have found similar patterns of species richness and ecological specialization (Sheikh et al. 2022; Zhang et al. 2022). Gall wasp shifts between oak organs (on the same or different trees) have also been frequent in the Western Palearctic (Cook et al. 2002) and genetic studies suggest that *Synergus* in the Western Palearctic are more specialized and species rich than morphology had previously implied (Ács et al. 2010). Thus, the species richness of both oak gall wasps and at least four of their associated parasite genera seem inextricably tied to the diversity of their respective hosts and habitats. Just as each oak tree species, with its many susceptible tissue types, represents myriad potential habitats for oak gall wasps, so does each oak gall represent a multi-dimensional habitat (varying in tree, organ, and/or morphology) for parasites of galls. Changes in any of these dimensions might stimulate host shifts and the subsequent evolution of reproductive isolation for one (or several) interacting insects. Thus, the effects of habitat variation on the origin of specialist parasite diversity in insect-plant gall systems may have broad importance across trophic levels.

## Author Contributions

Project idea: AKGW and AAF. Collections: AKGW, SIS, and AAF. Sequencing and analyses of UCE and mtCOI: AKGW, YMZ, GEB, ACH, SR, NS, SIS, CT, and AAF. Species delimitation: YMZ, SR, NS, CT, and AAF. Analyses of gall and inquiline niche space: KMP. Coevolutionary analyses: YMZ, GEB, and AAF. Writing and editing: all authors.

## Acknowledgements

The authors thank C. Davis, B. Foley, K. McElroy, D. McGarry, K. Neely, M. Neiman, S. Pelini, M. Shakally, E. Tvedte, J. Verry, H. Widmayer, and C. Wilson for help with gall collection and/or insect rearing. L. Nastasi helped with specimen vouchering. Funding for collections was provided to AAF by the University of Iowa and to AKGW by the Center for Global and Regional Environmental Research. Funding for library preparation and sequencing was provided to AKGW via an EECG award from the American Genetic Association. We acknowledge the Smithsonian Institution High Performance Cluster (https://doi.org/10.25572/SIHPC) for providing computational resources and support that have contributed to the research results reported in this publication. YMZ was funded by the European Union’s Horizon 2020 research and innovation program under the Marie Sklodowska-Curie grant agreement no. 101024056.

## Conflict of Interest

The authors declare no conflict of interest.

## Data Availability Statement

Mitochondrial COI sequences generated for this project are available on NCBI GenBank (OR371993 - OR372098). Raw UCE sequences are available on the NCBI Short Read Archive (BioProject PRJNA1001032). Raw trees, and files used in ASR analysis, and dataframes and codes used in trait analysis will be available on the Dryad Digital Repository and at Zenodo upon publication or may be requested from the corresponding author now.

## Notes

### Competing Interest Statement

The authors have declared no competing interest.

