## Supplemental Figures S1-S69 for "Speciation in kleptoparasites of oak gall wasps often correlates with a shift into a new tree habitat, tree organ, or gall morphospace"

**Supplemental File 1. Images of *Synergus* and *Ceroptres* wasps sequenced in this study. See Table S1 for collection details. Images for some Synergus in Table S1 were previously published in Ward et al., 2020, and those images are not replicated here.**

Figure S1. *Synergus* sp. 7 – 1524-124-1 – female from *Andricus kingi* on *Quercus lobata* in Westlake Villiage, CA.

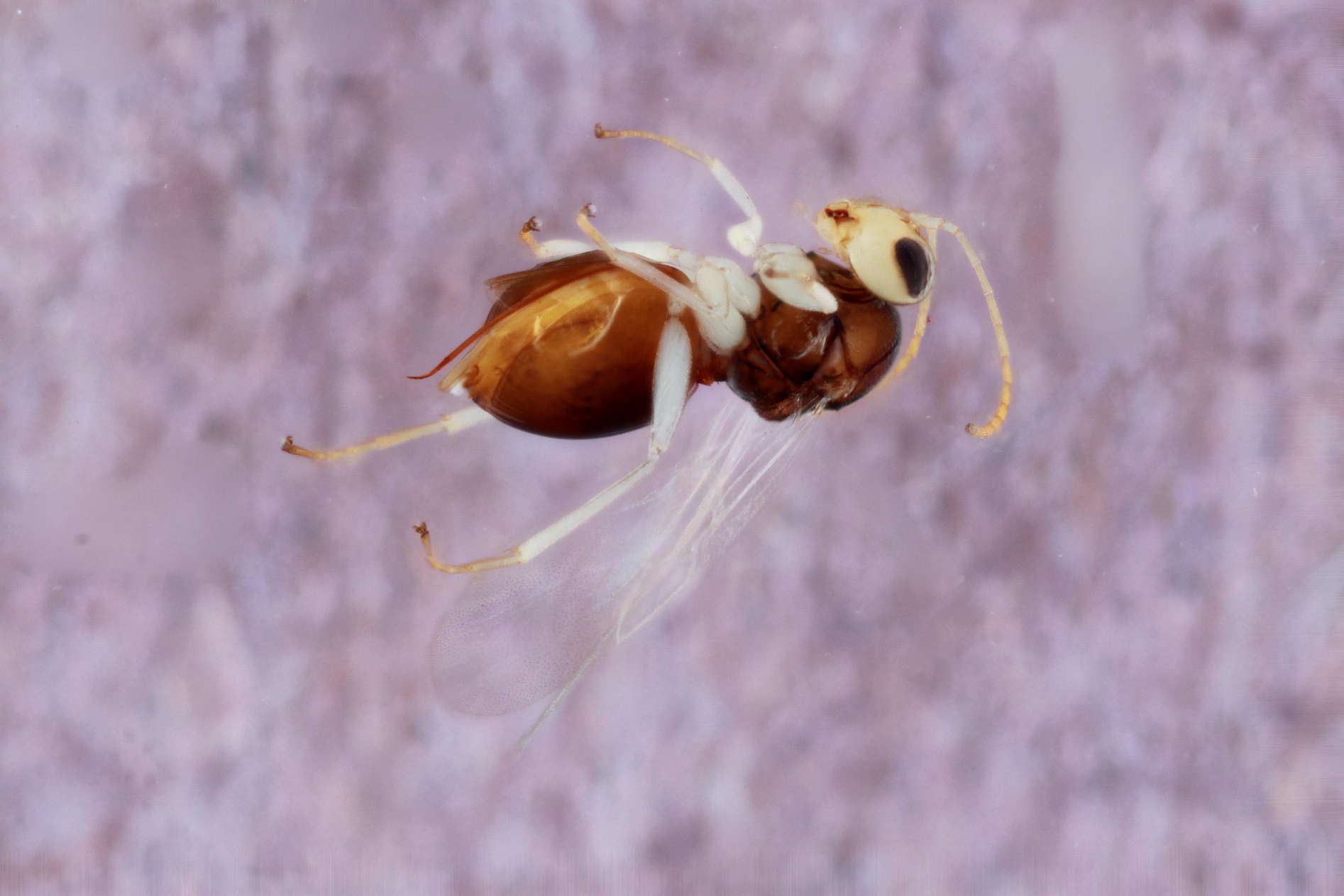

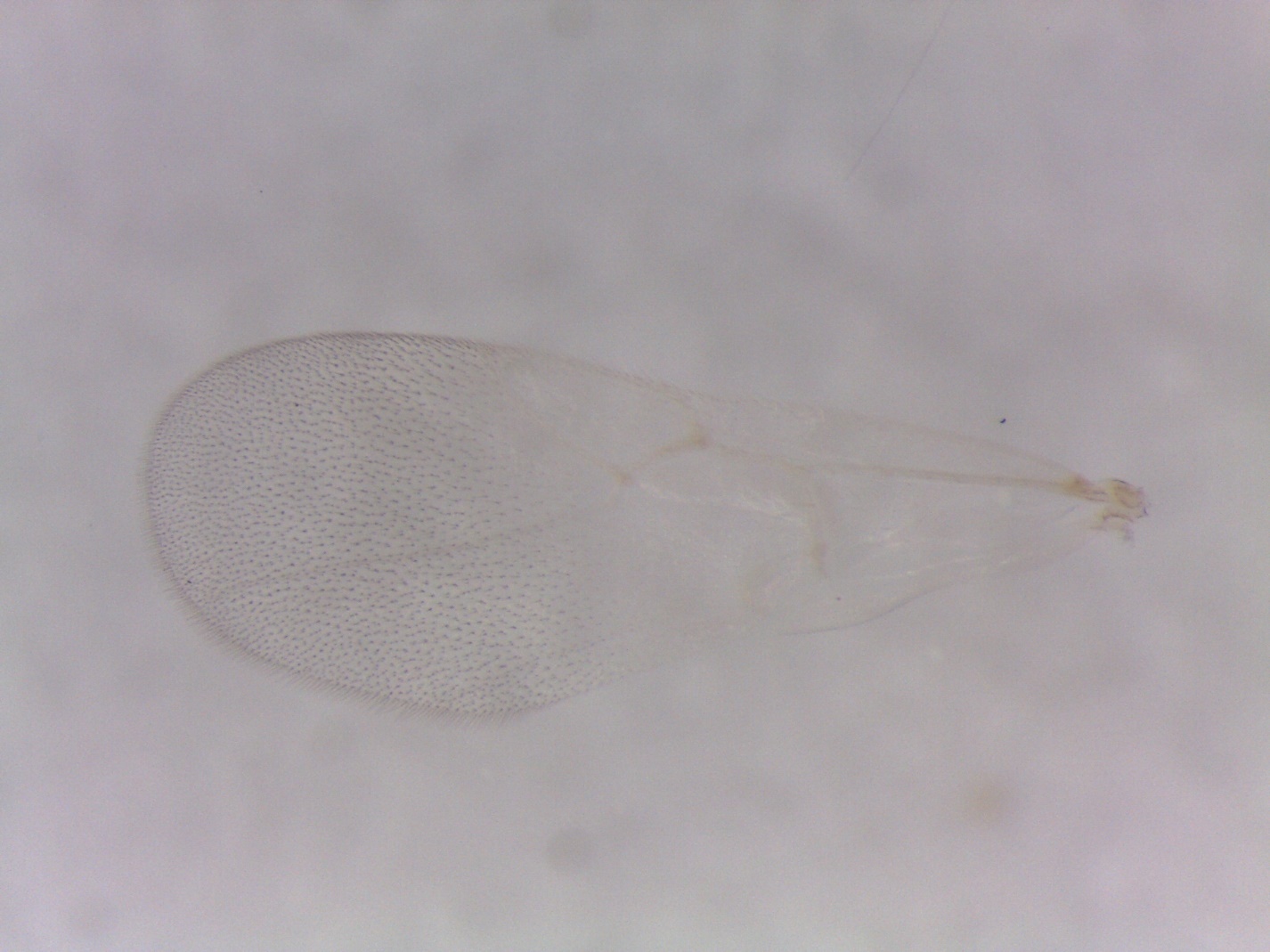

Figure S2. *Synergus* sp. 7 – 1426-126-1 – female from *Andricus douglasi* on *Quercus lobata* in Folsom, CA.

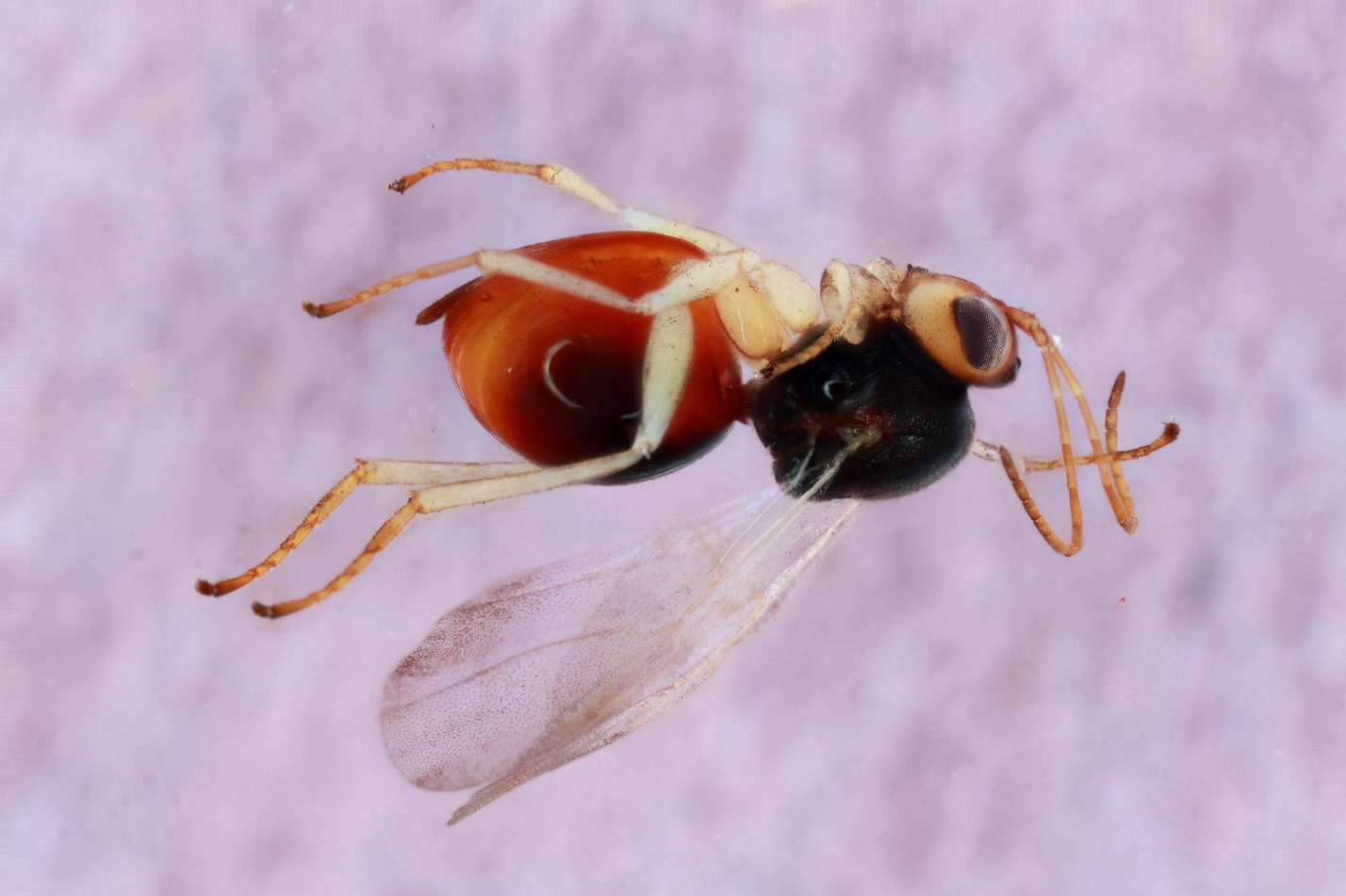

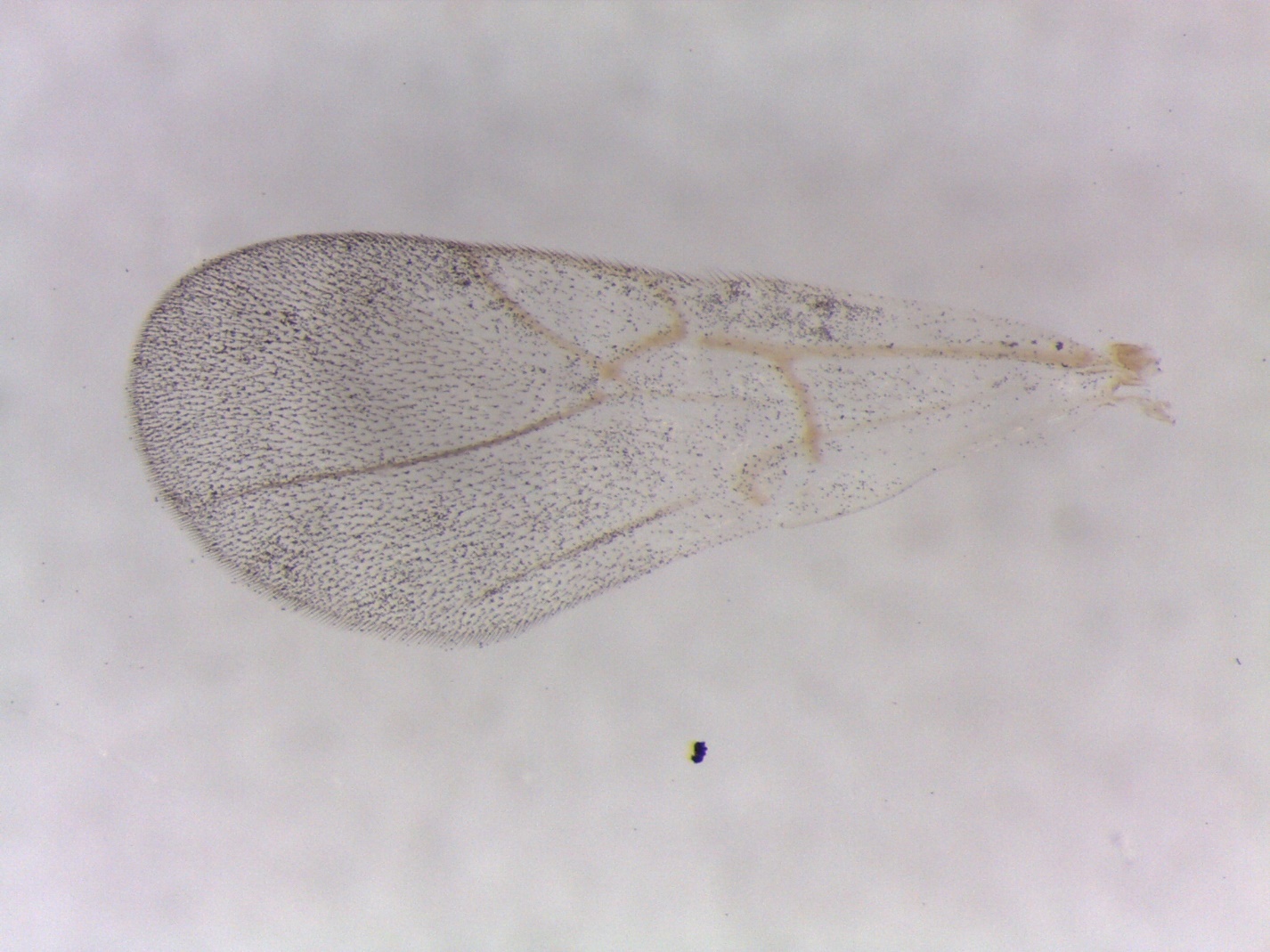

Figure S3. *Synergus* sp. 8 – 1497-141-3A – female from *Dryocosmus minusculus* on *Quercus agrifolia* in San Jose, CA.

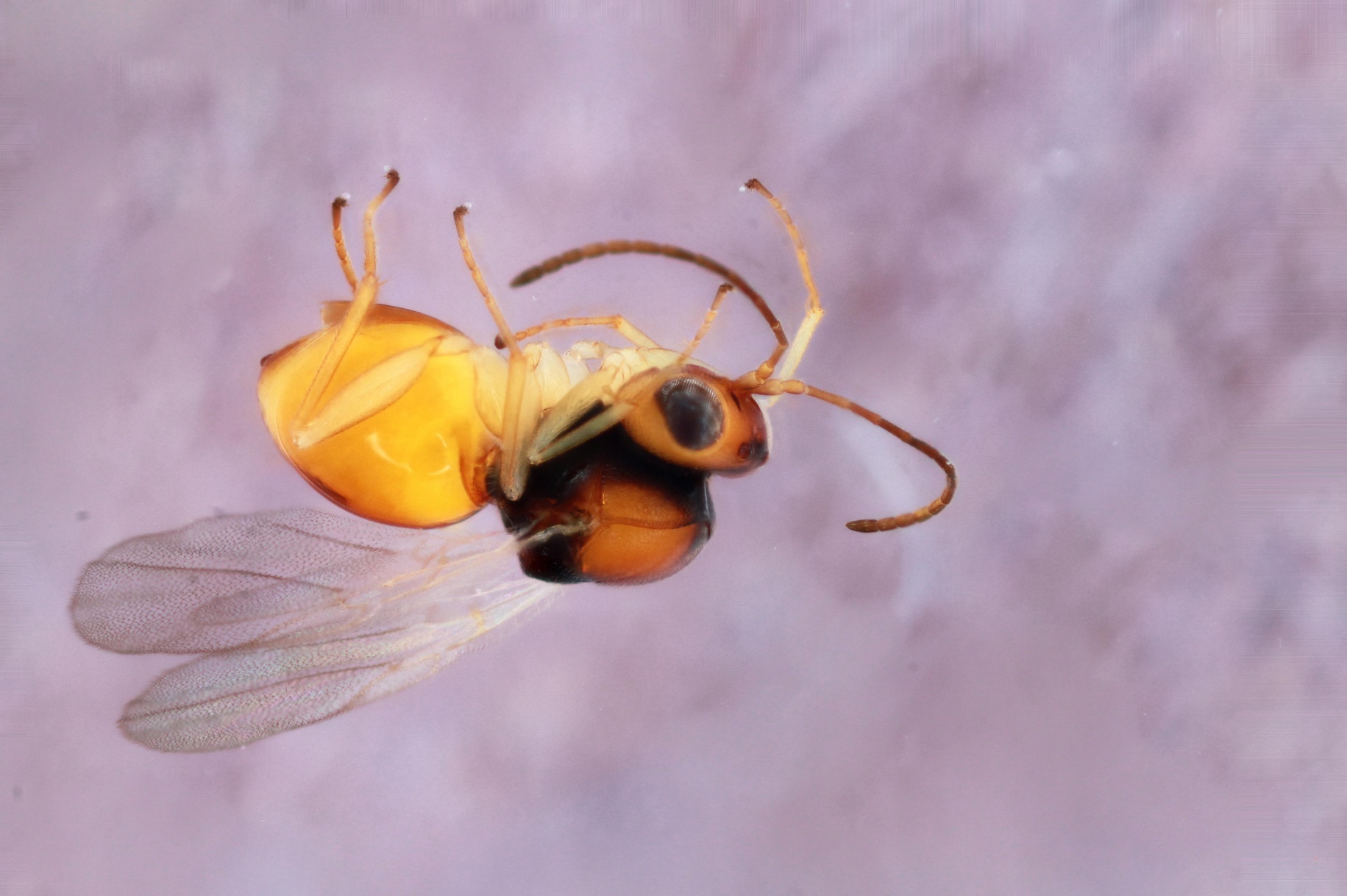

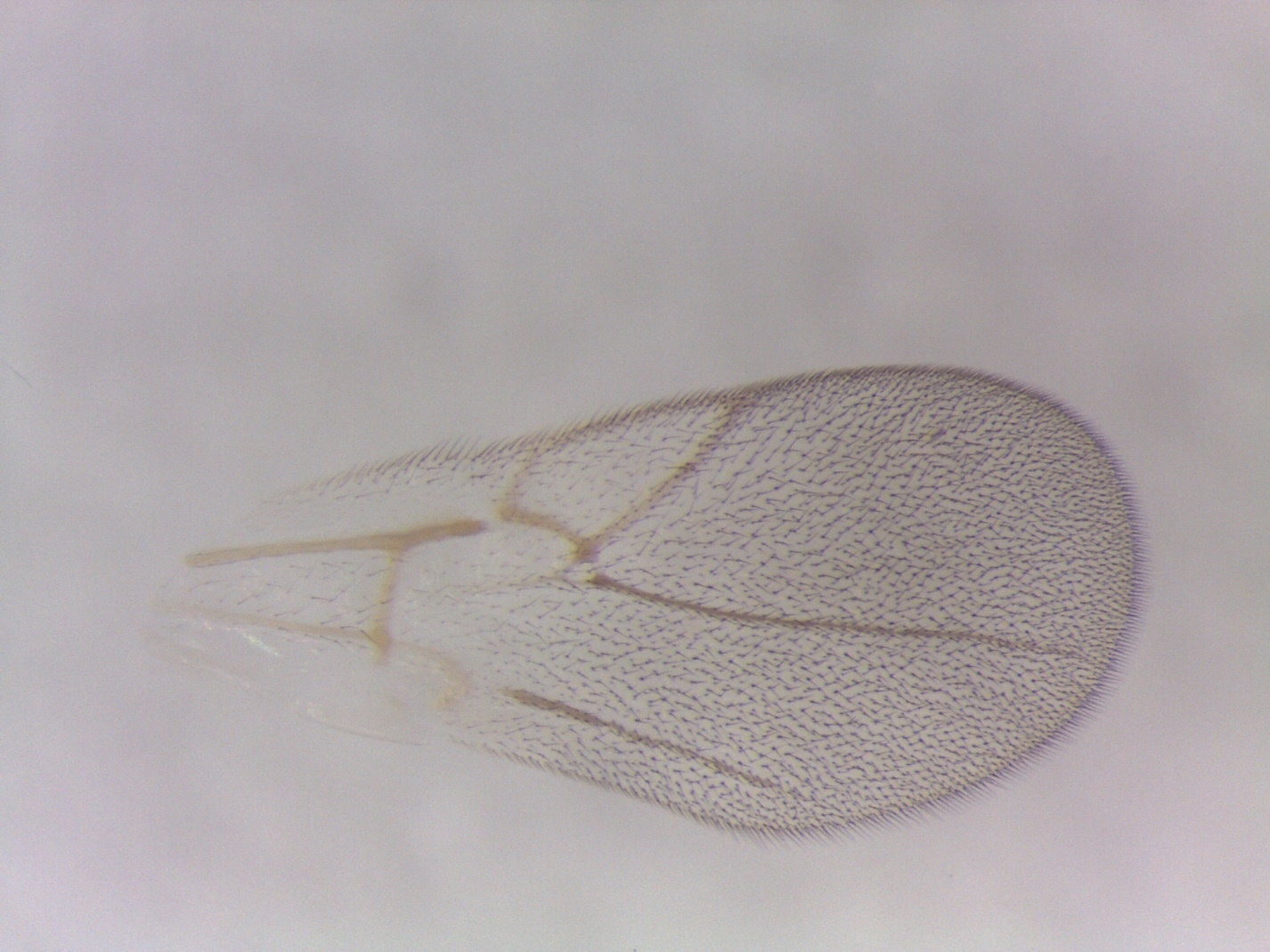

Figure S4. *Synergus* sp. 8 – 1497-141-4A – female from *Dryocosmus minusculus* on *Quercus agrifolia* in San Jose, CA.

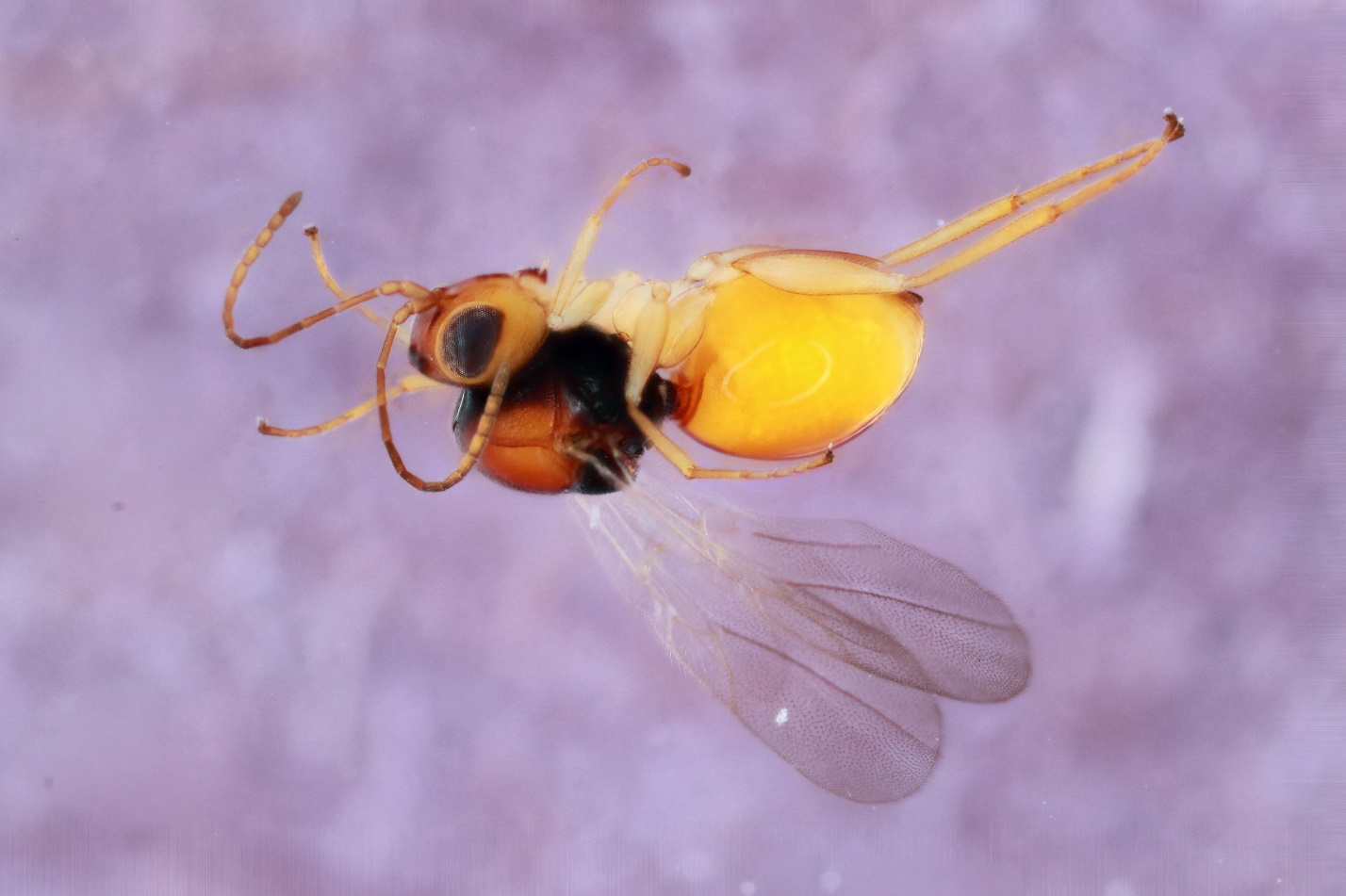

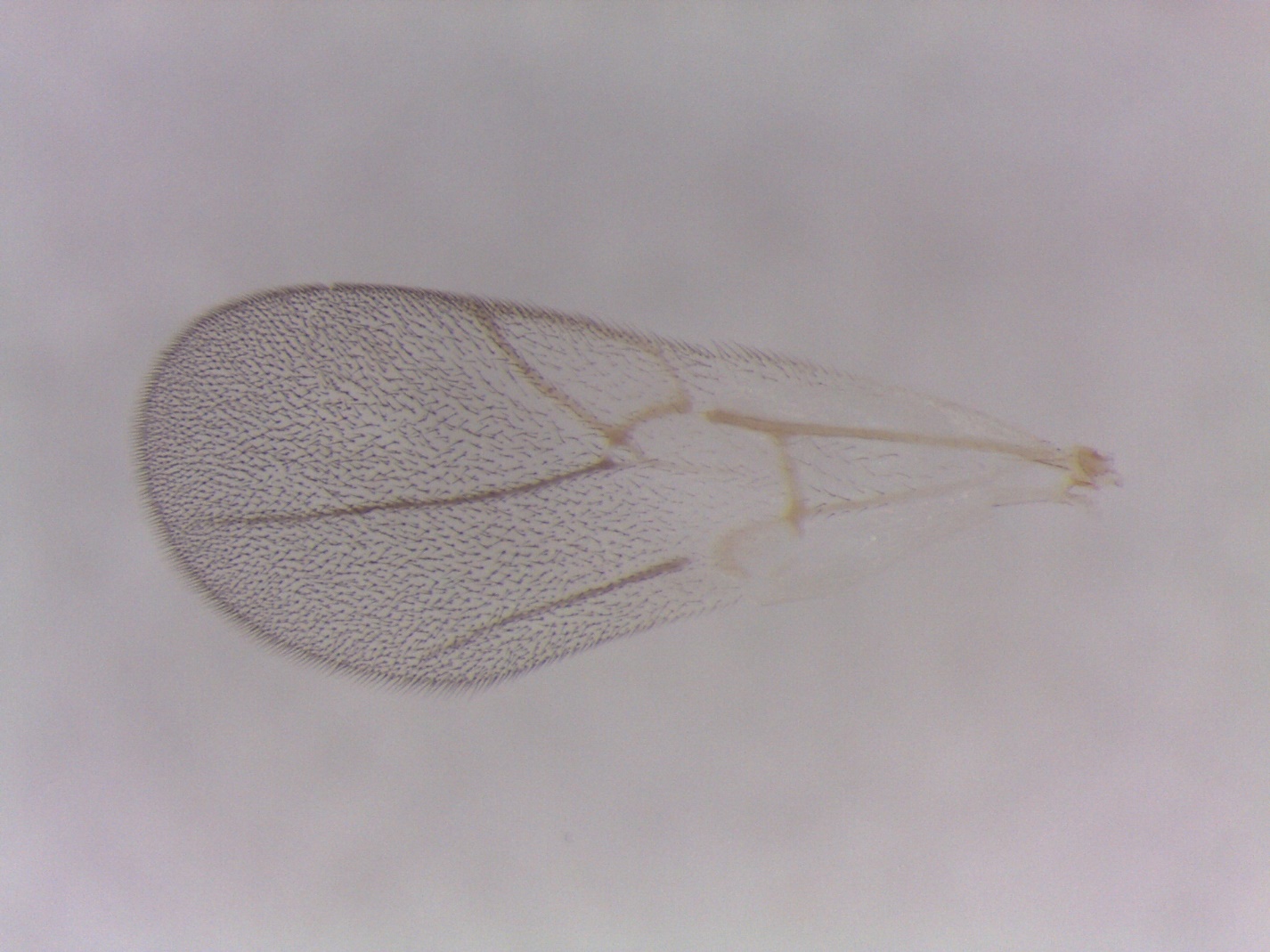

Figure S5. *Synergus* sp. 8 – 1515-141-2 – male from *Dryocosmus minusculus* on *Quercus agrifolia* in Buelton, CA.

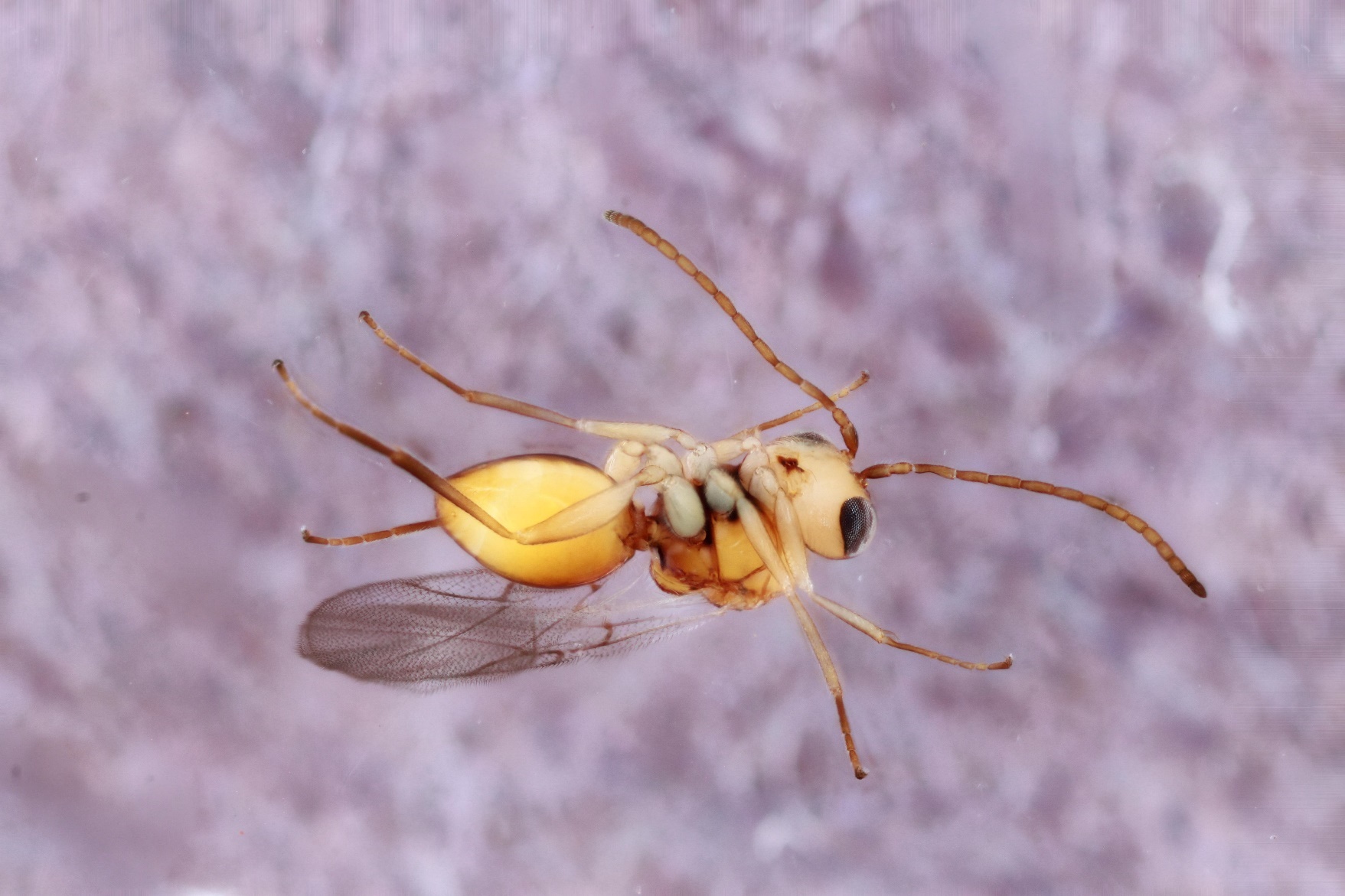

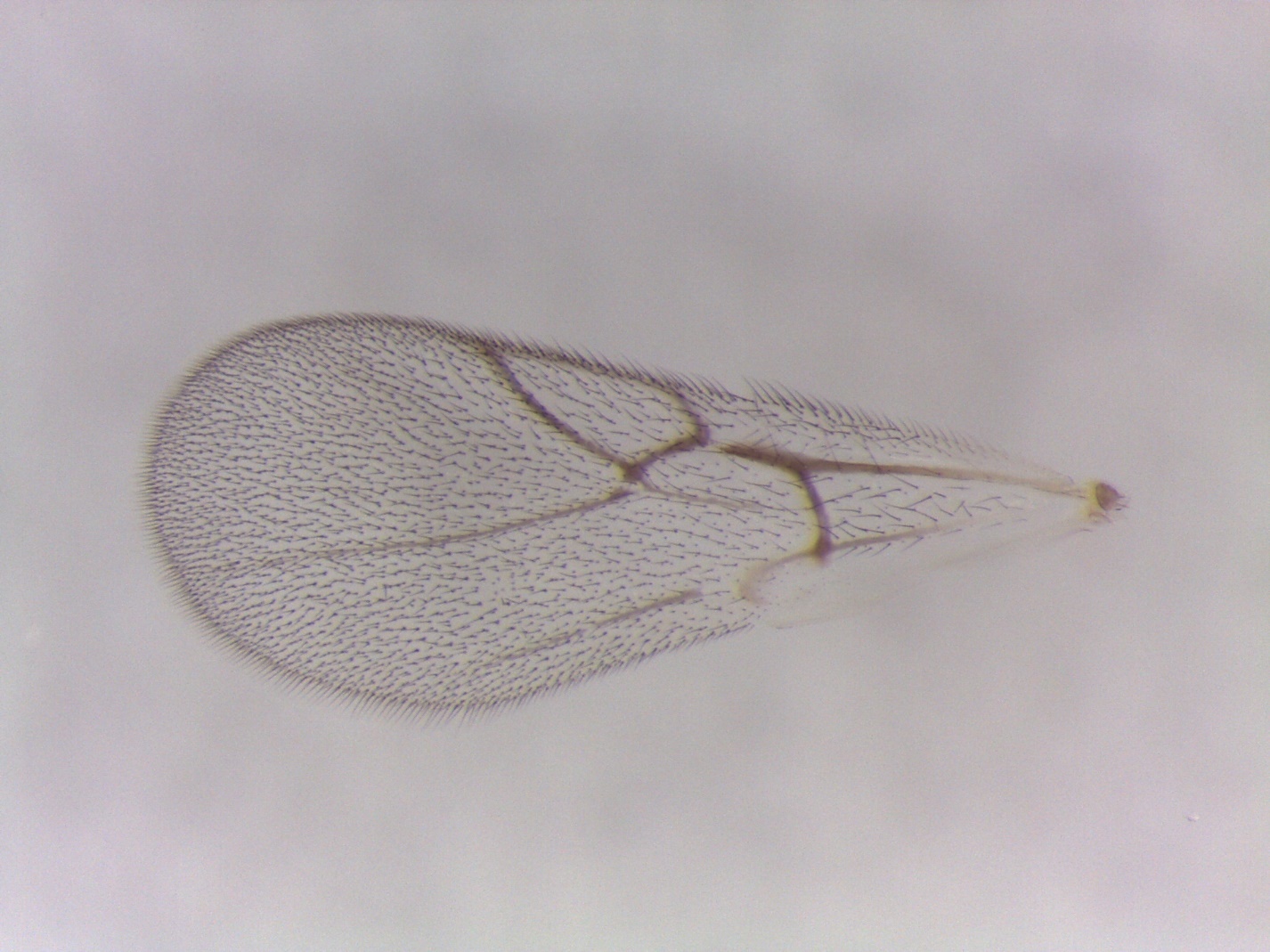

Figure S6. *Synergus* *ochreus* – 1467-125-1 – female from *Cynips conspicuus* on *Quercus lobata* in Folsom, CA.

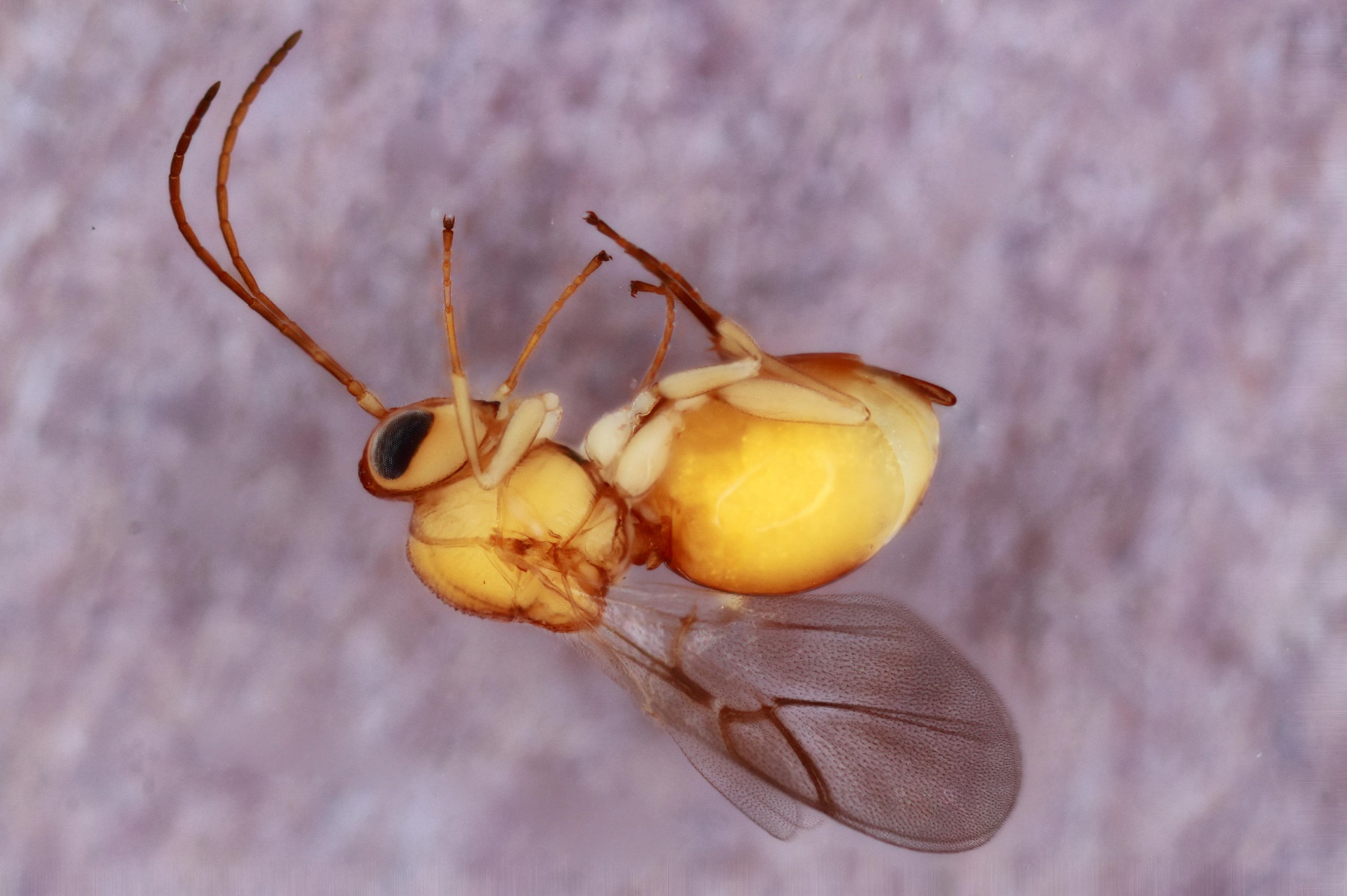

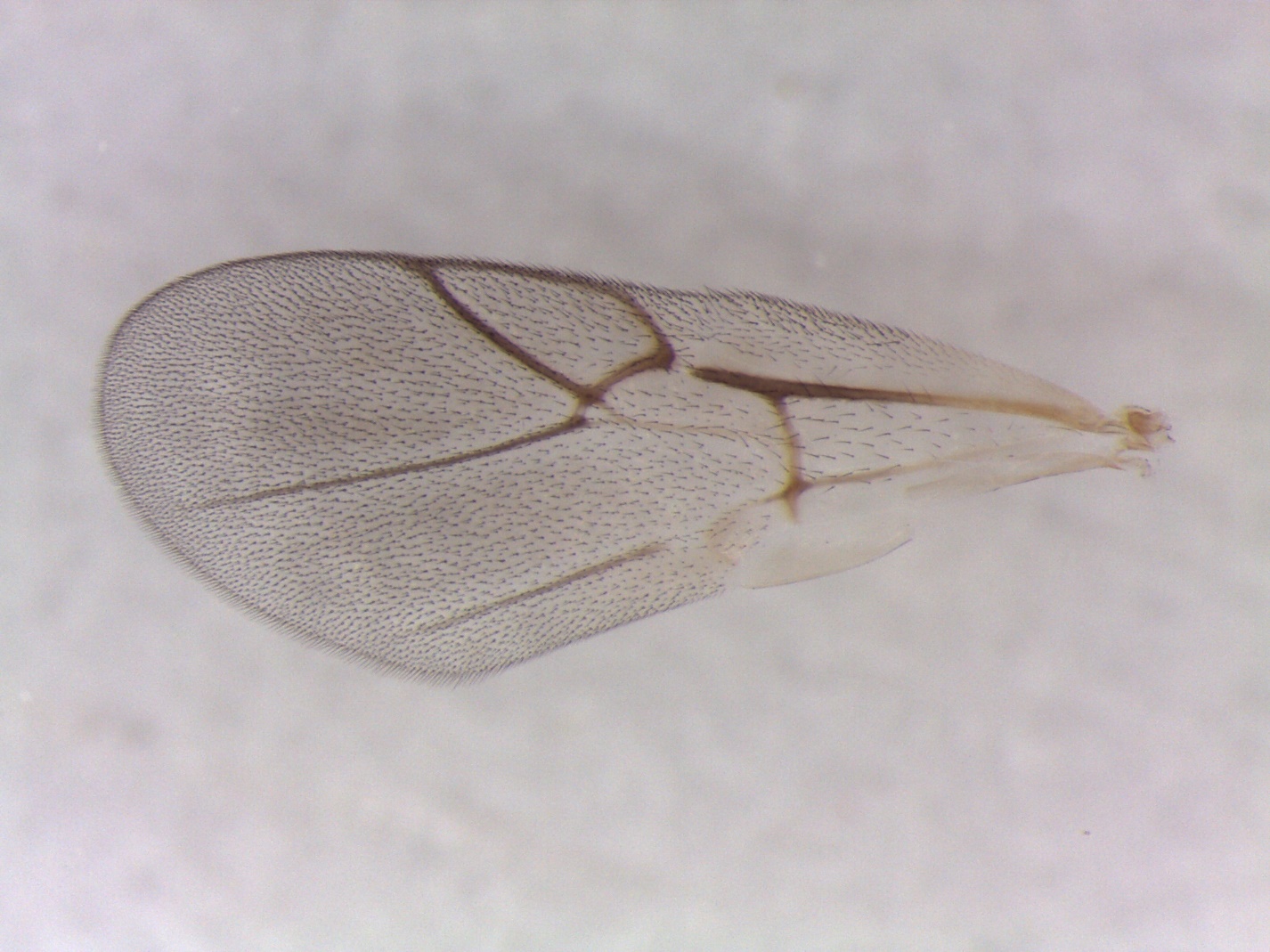

Figure S7. *Synergus* sp.1 – 1449-034-1 – male from *Callirhytis clavula* on *Quercus alba* in Dodgeville, WI.

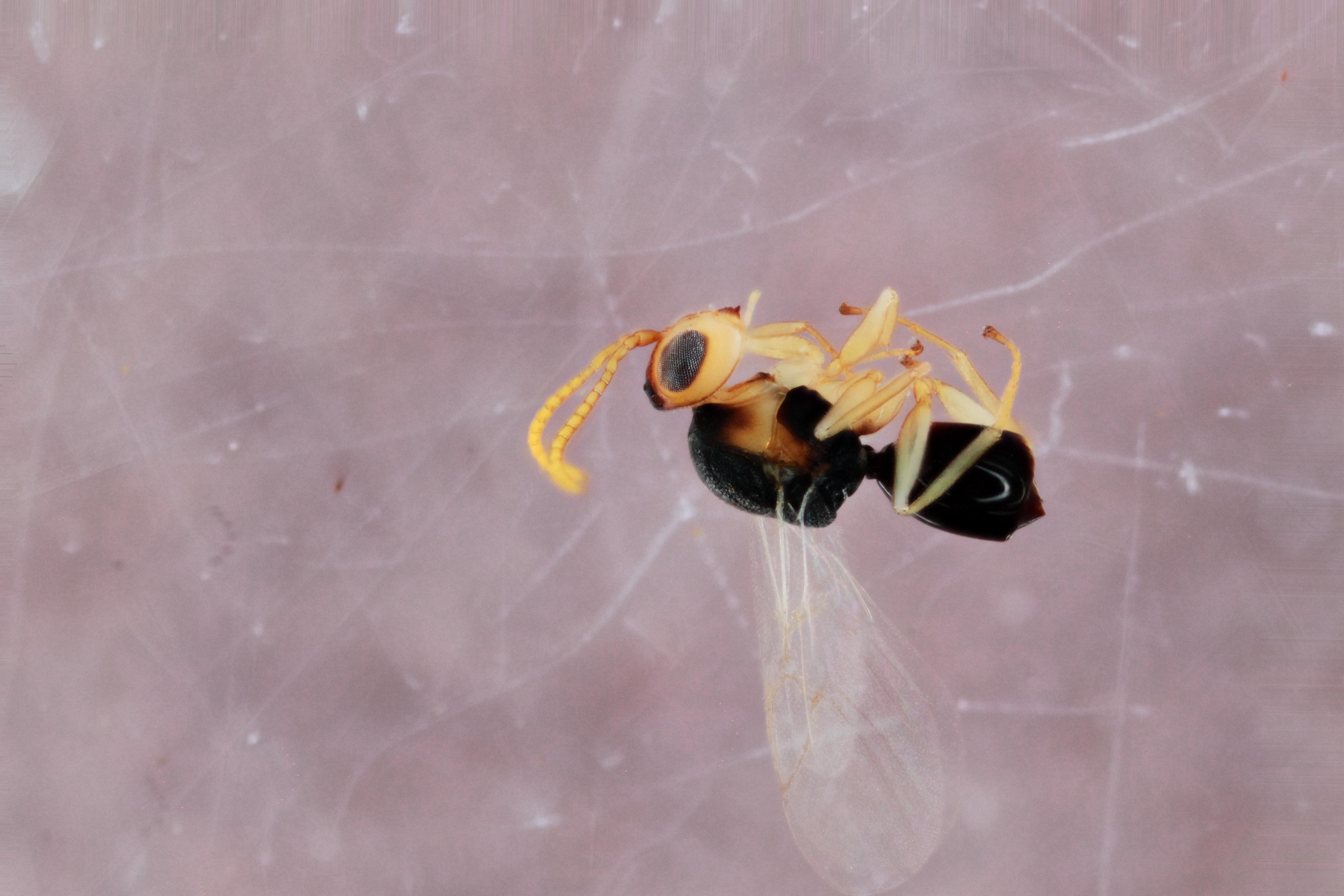

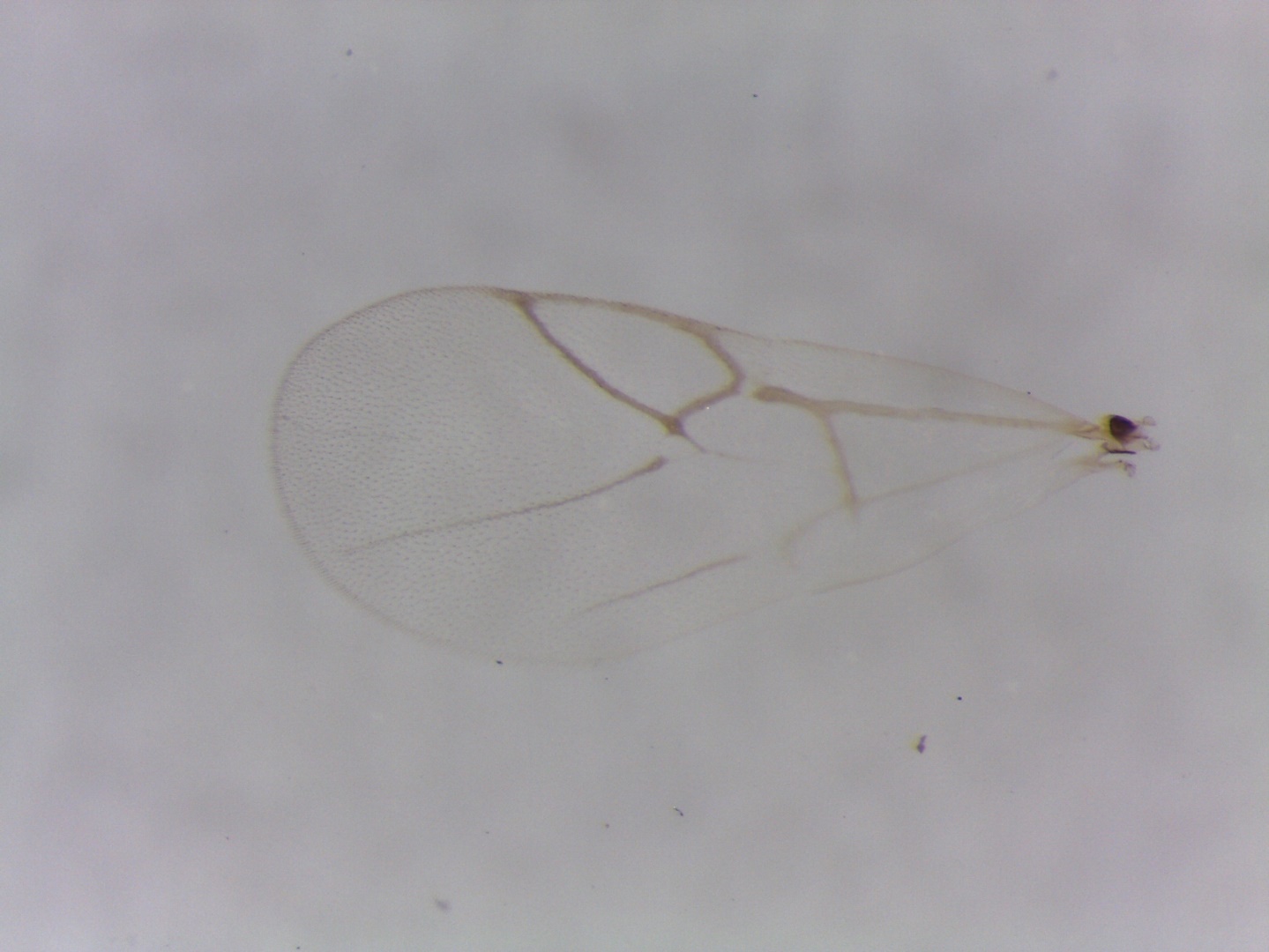

Figure S8. *Synergus* sp.4 – 1059-024-6 – female from *Phylloteras poculum* on *Quercus bicolor* in Coralville, IA.

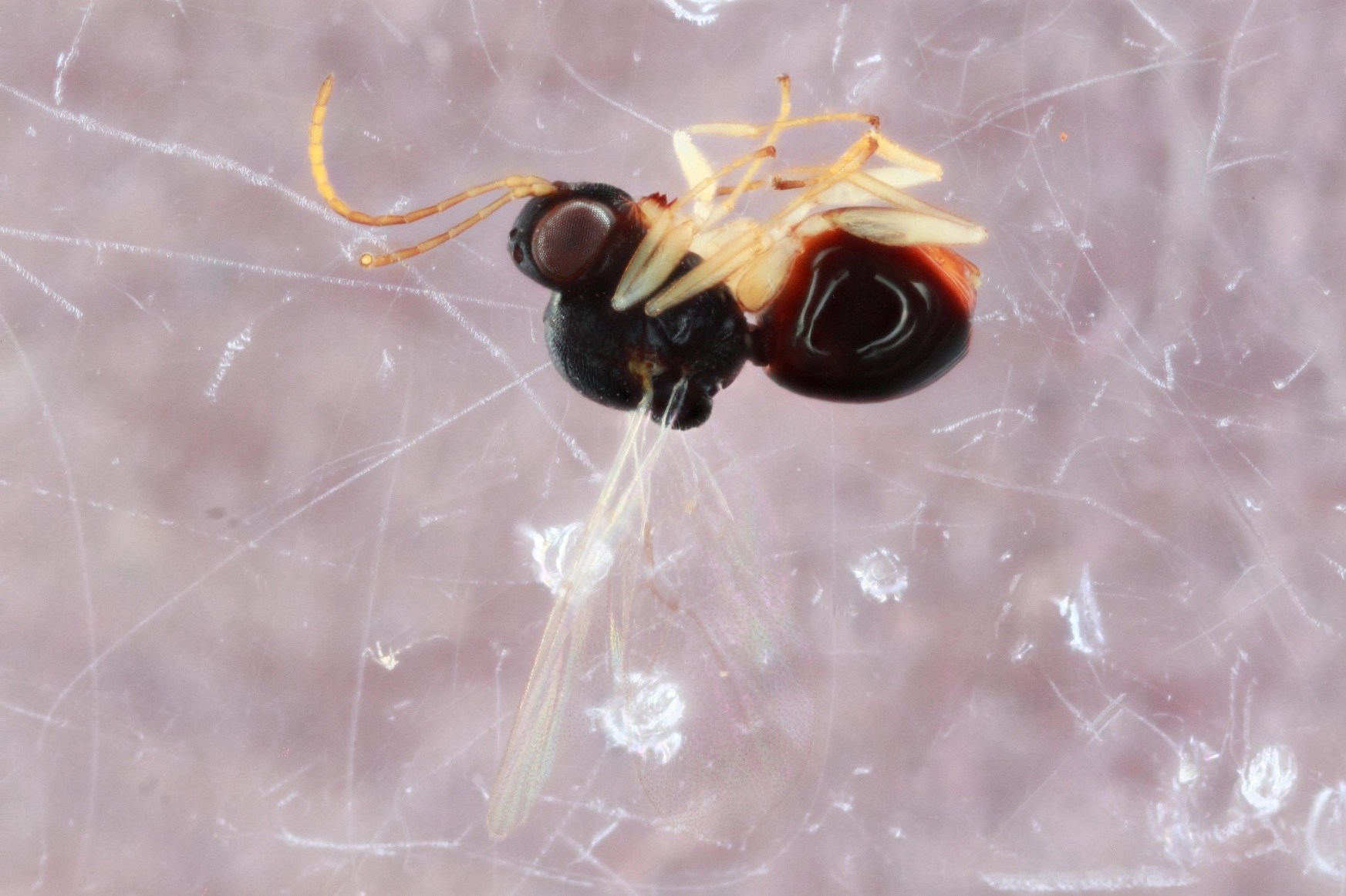

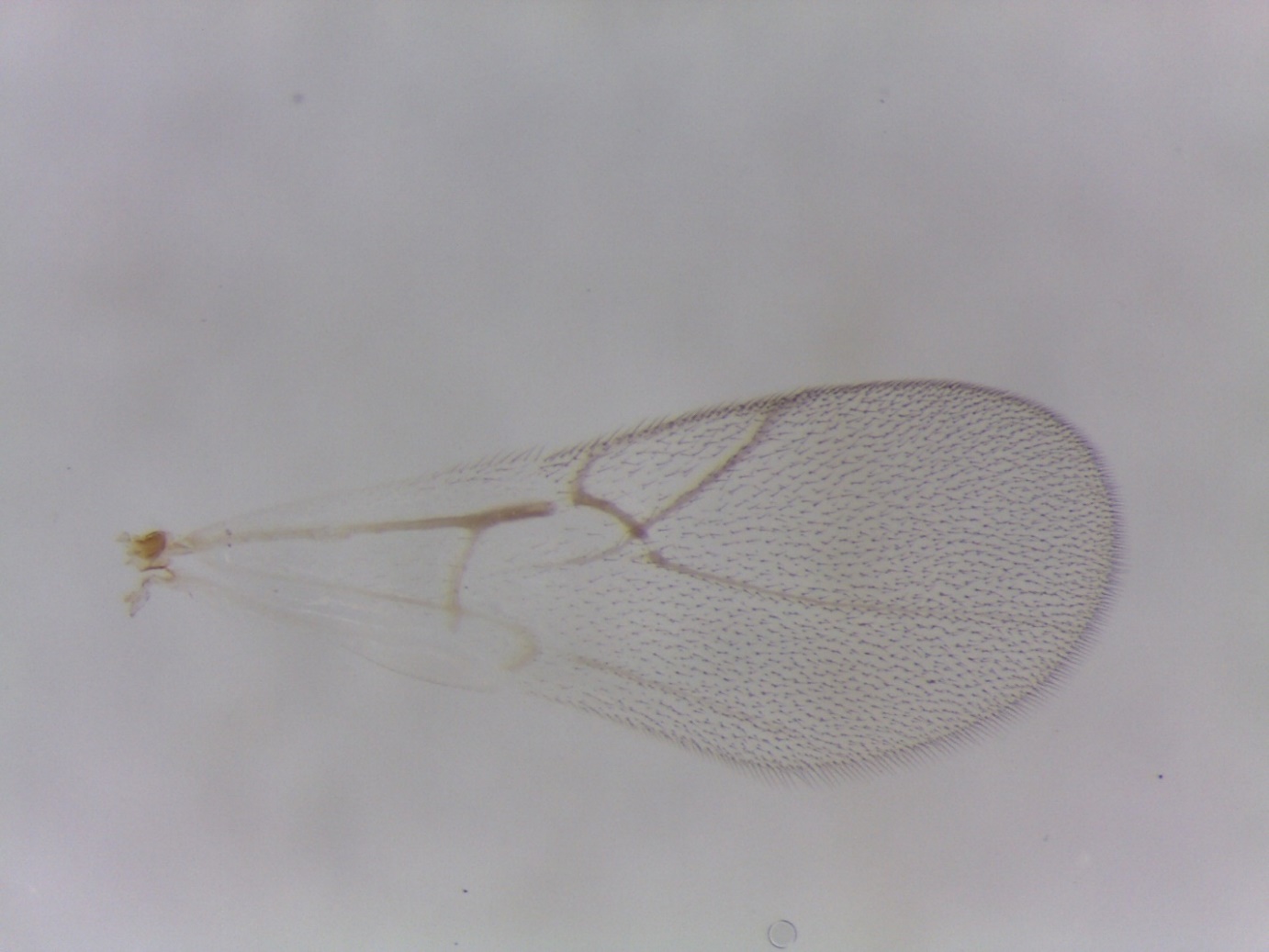

Figure S9. *Synergus* sp.5 – 1523-145-5 – female from *Disholcaspis eldoradensis* on *Quercus berberidifolia* in Borrego Springs, CA.

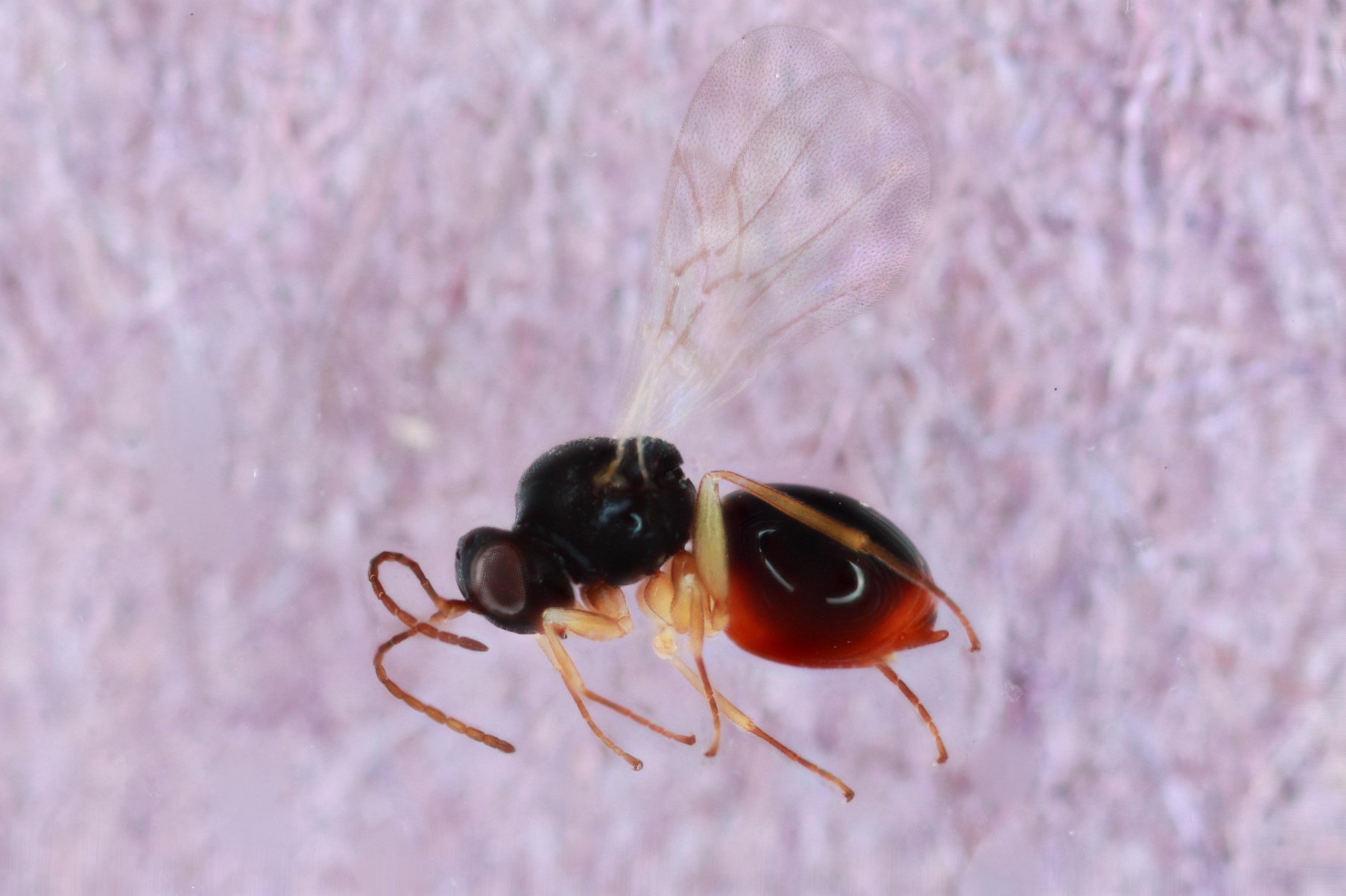

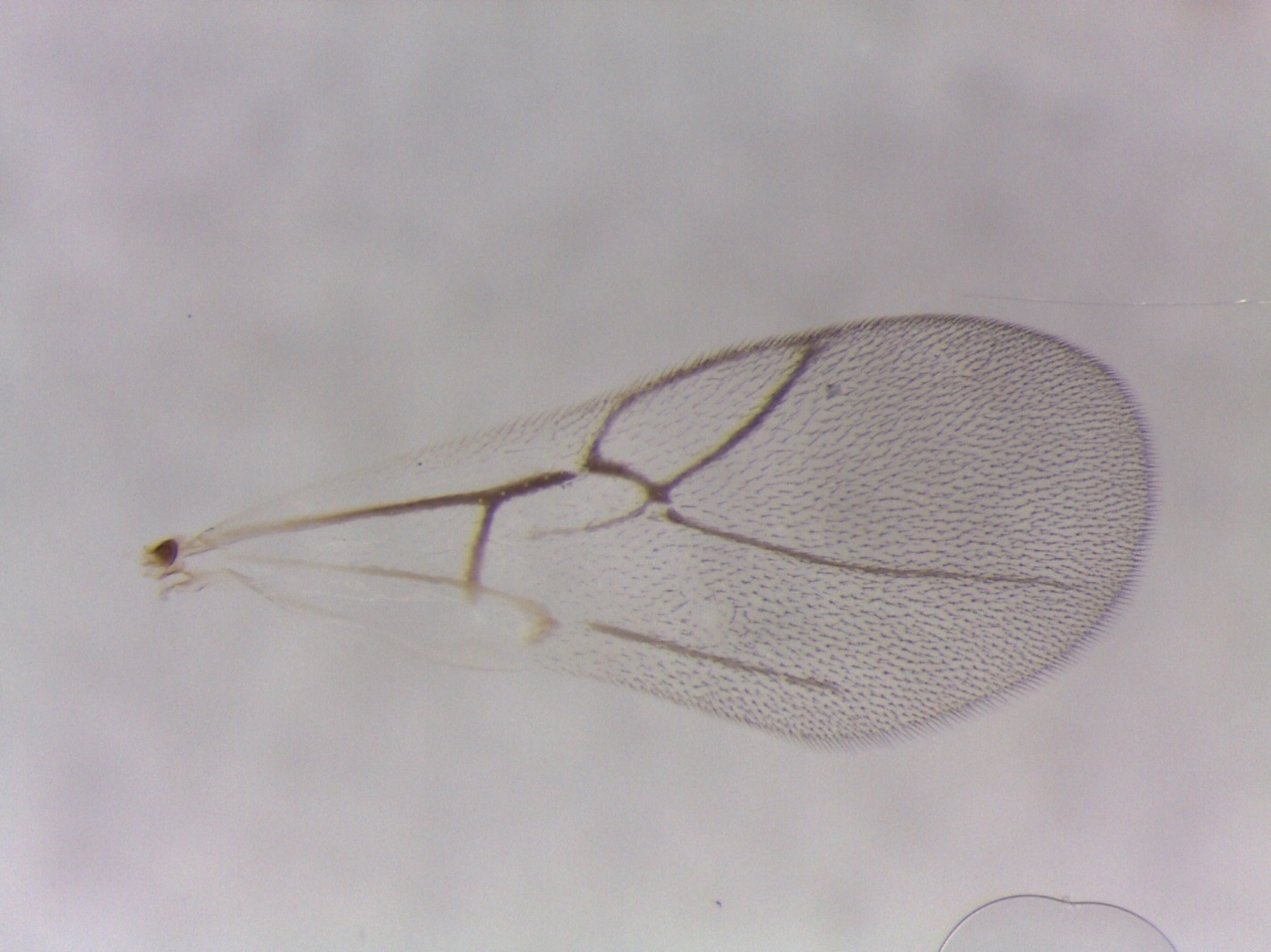

Figure S10. *Synergus* sp.6 – 1583-156-9 – female from *Belonocnema kinseyi* on *Quercus virginiana* in Austin, TX.

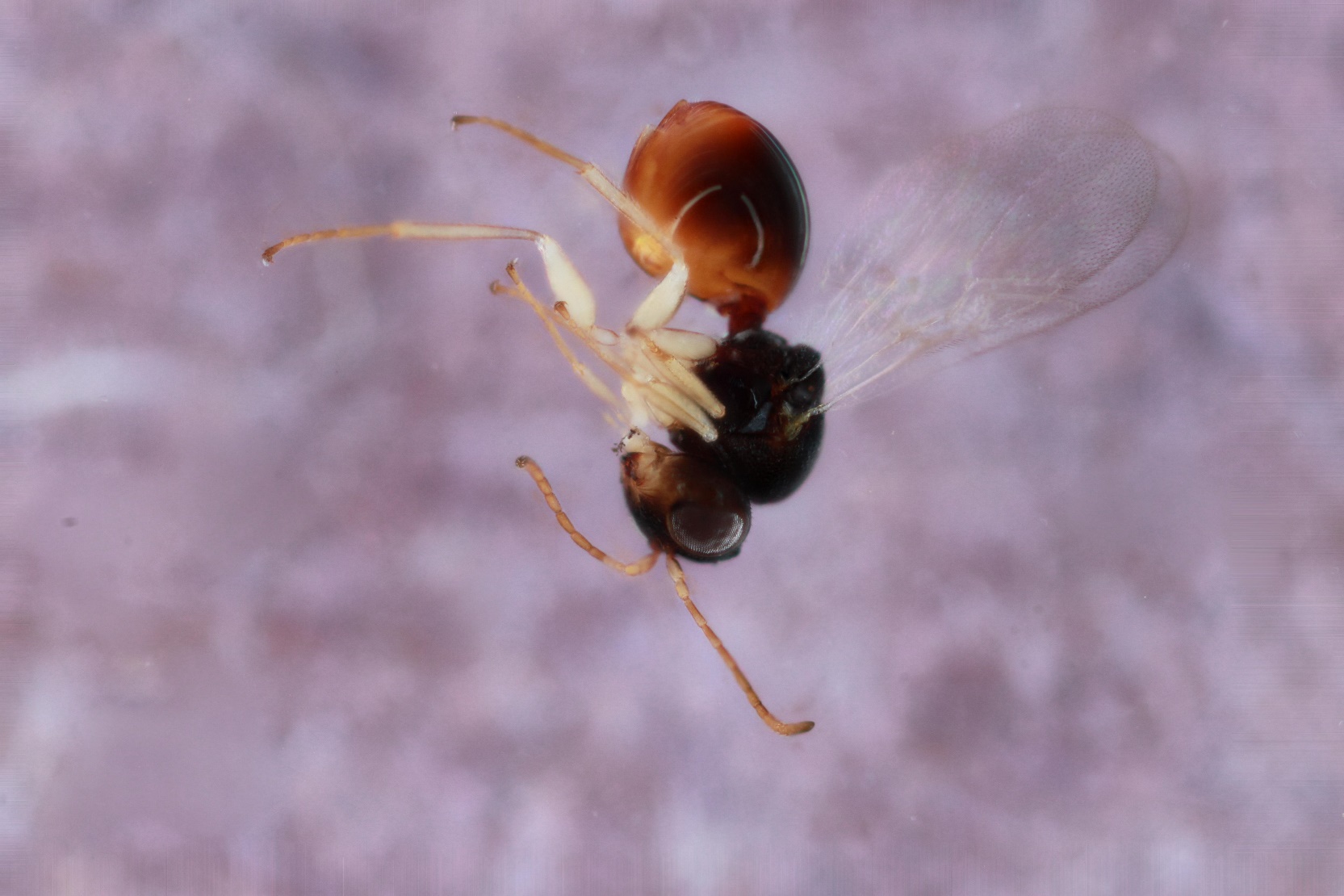

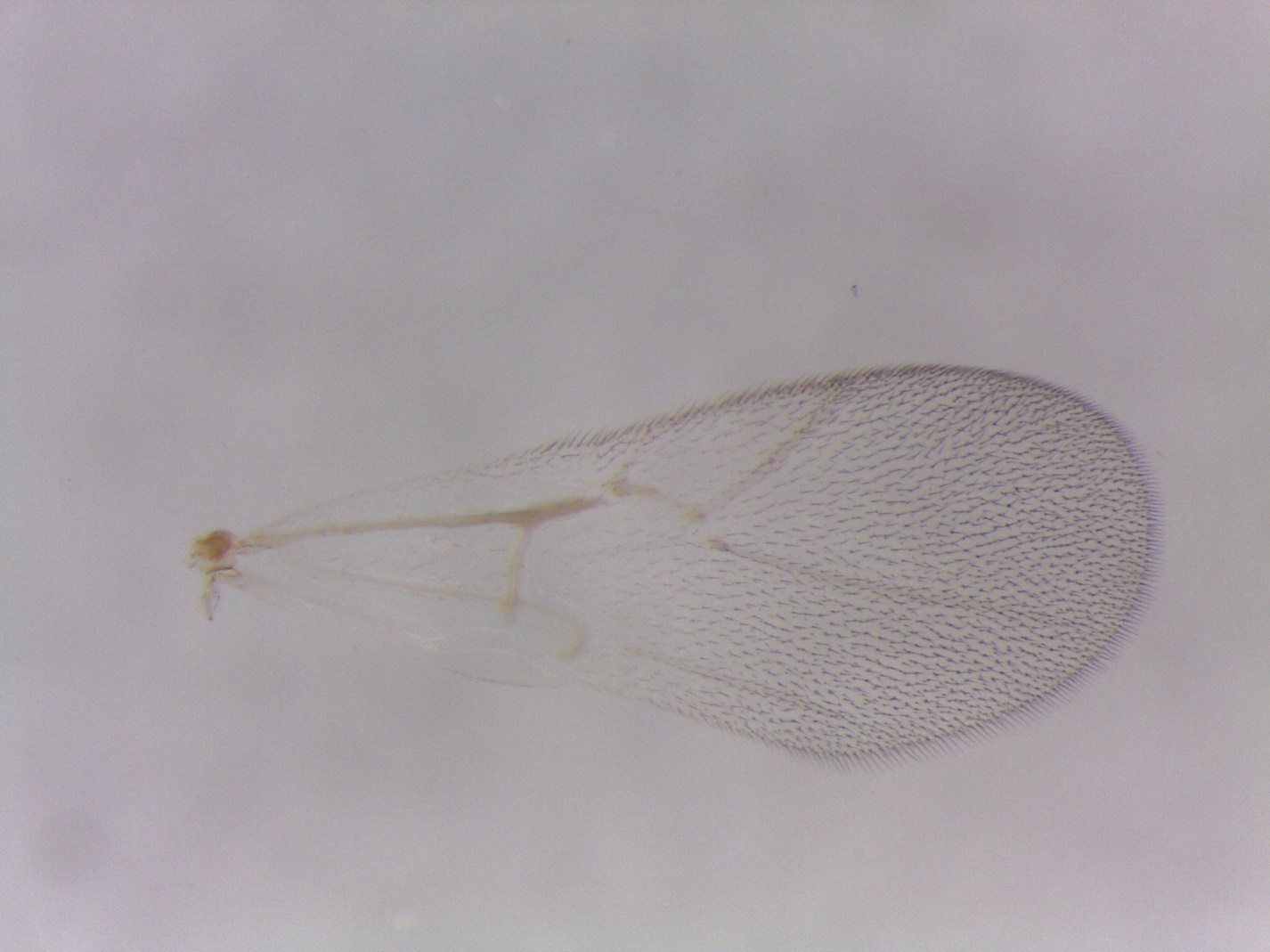

Figure S11. *Synergus* *walshii* A – 1589-160-36A – female from *Druon quercuslanigerum* on *Quercus virginiana* in Lake Kyle, TX.

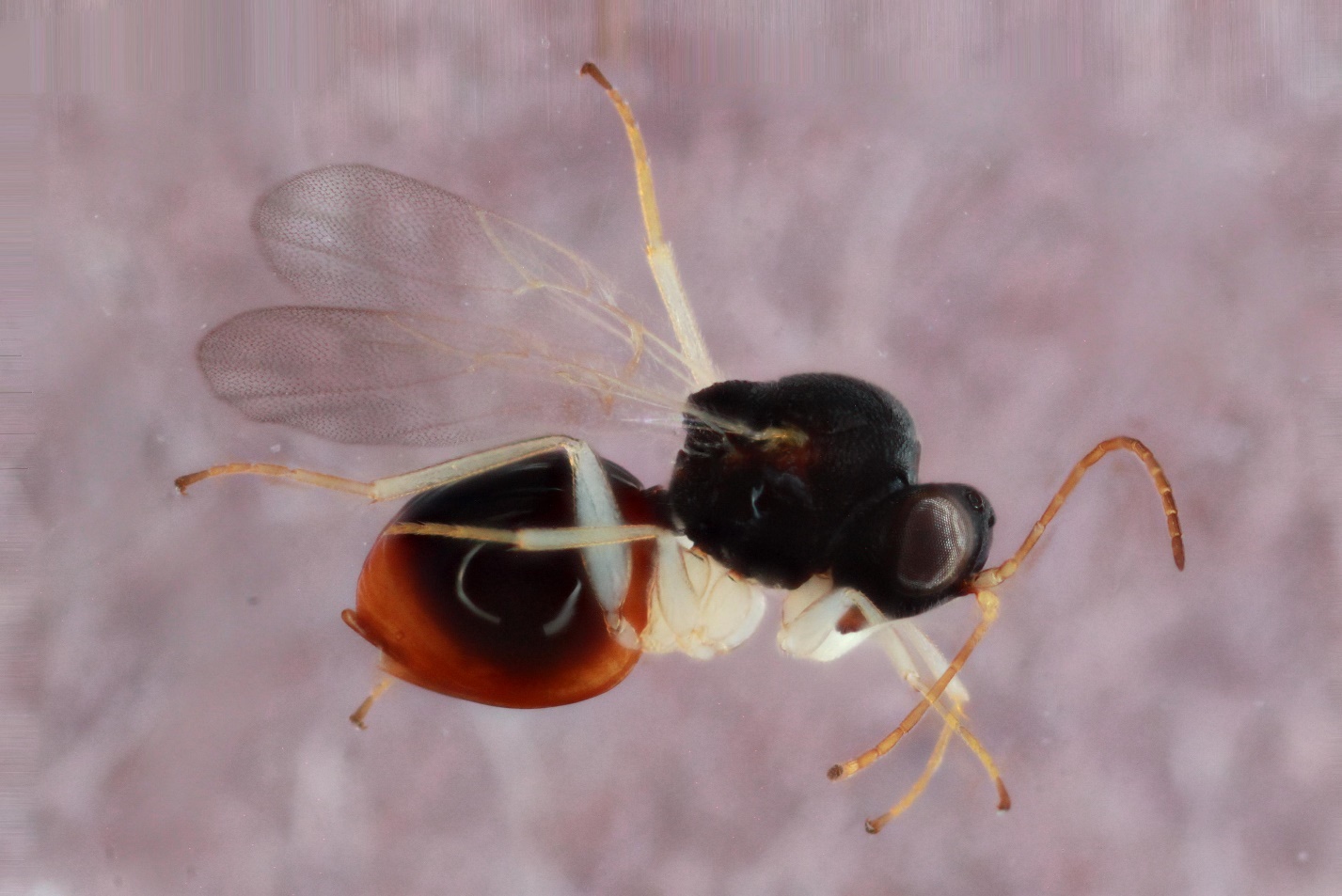

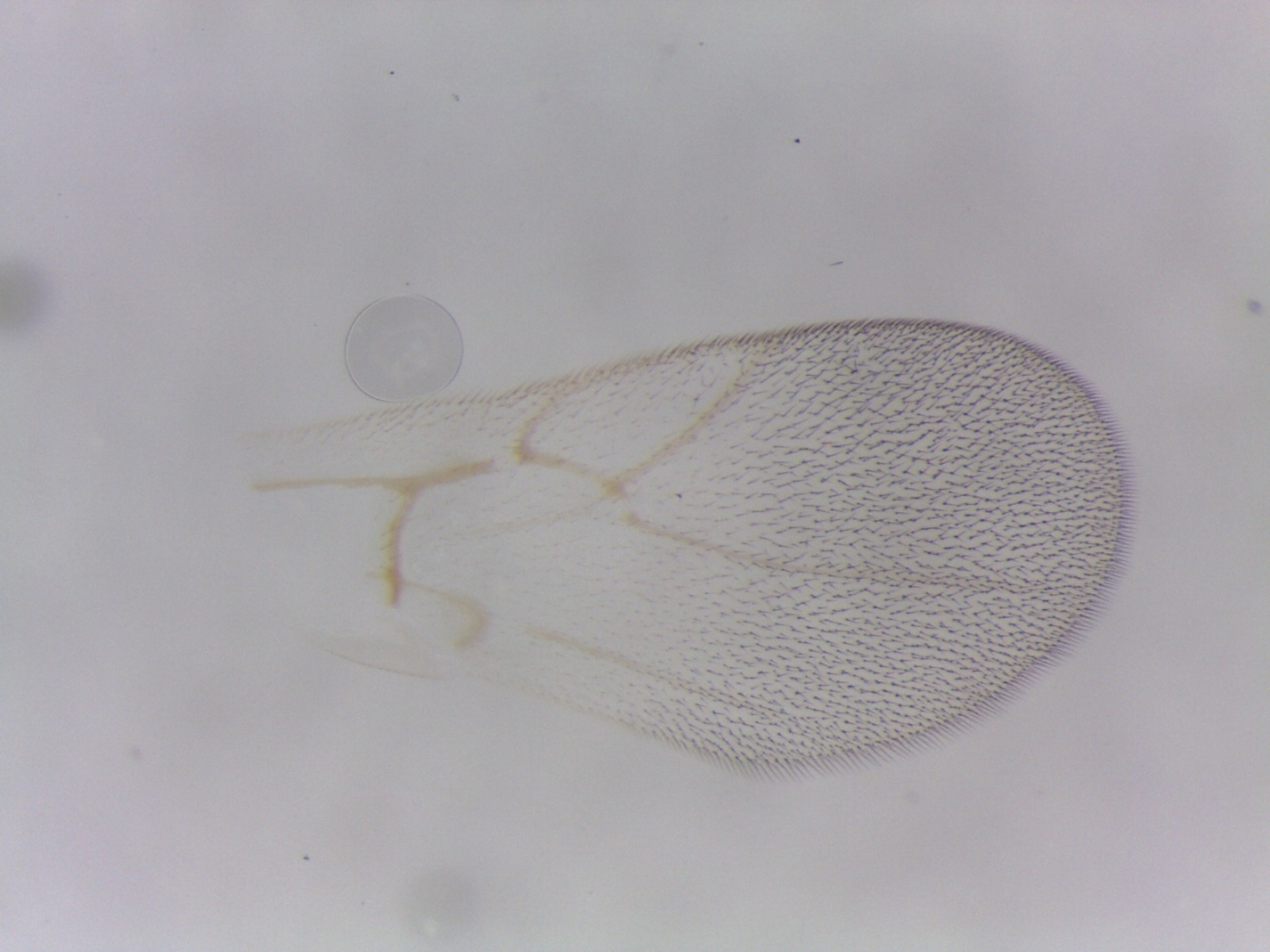

Figure S12. *Ceroptres* sp. 1-32-33-34 – 807-042-27 – female from *Andricus quercuspetiolicola* on *Quercus bicolor* in Tiffin, IA.

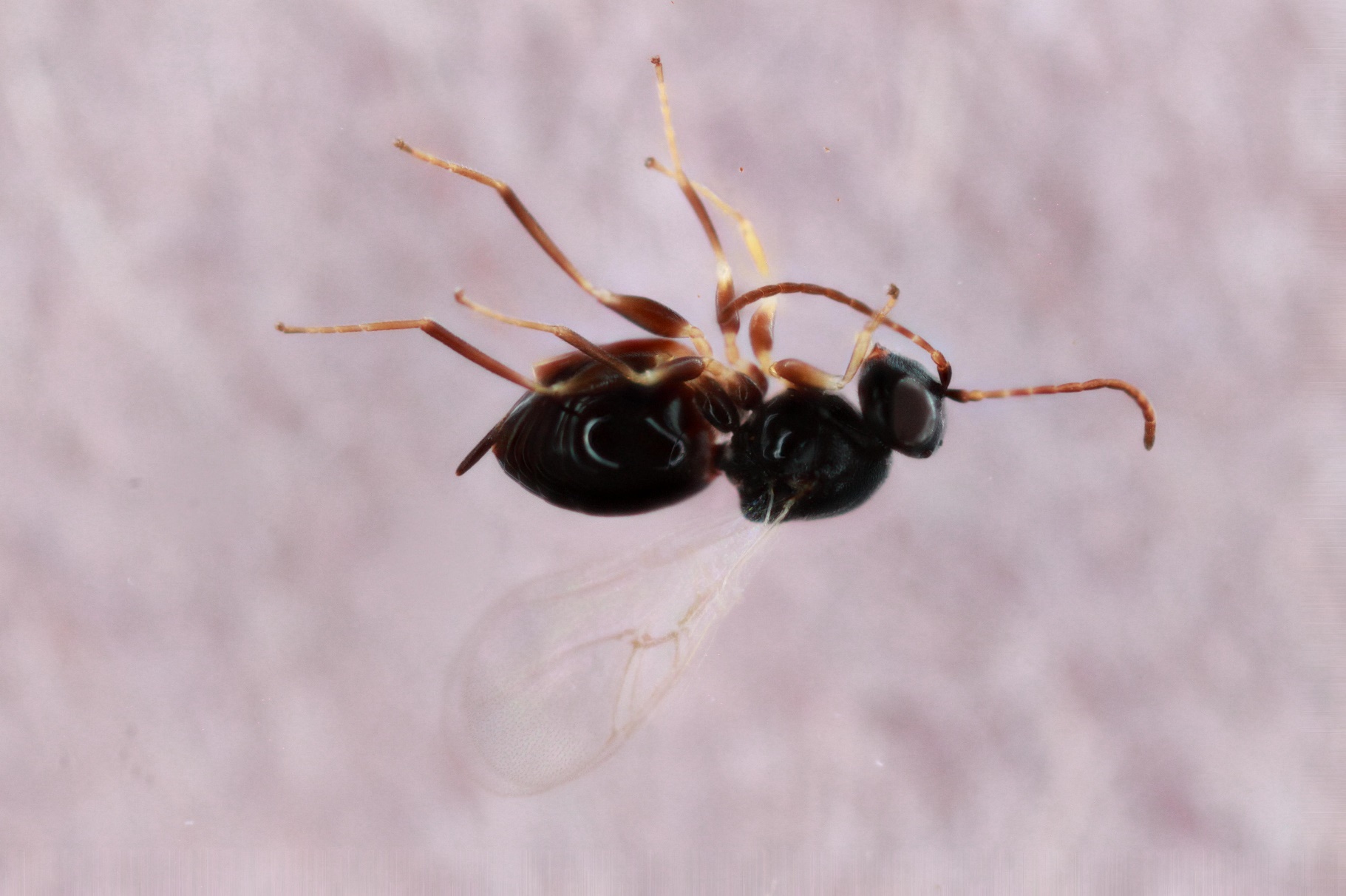

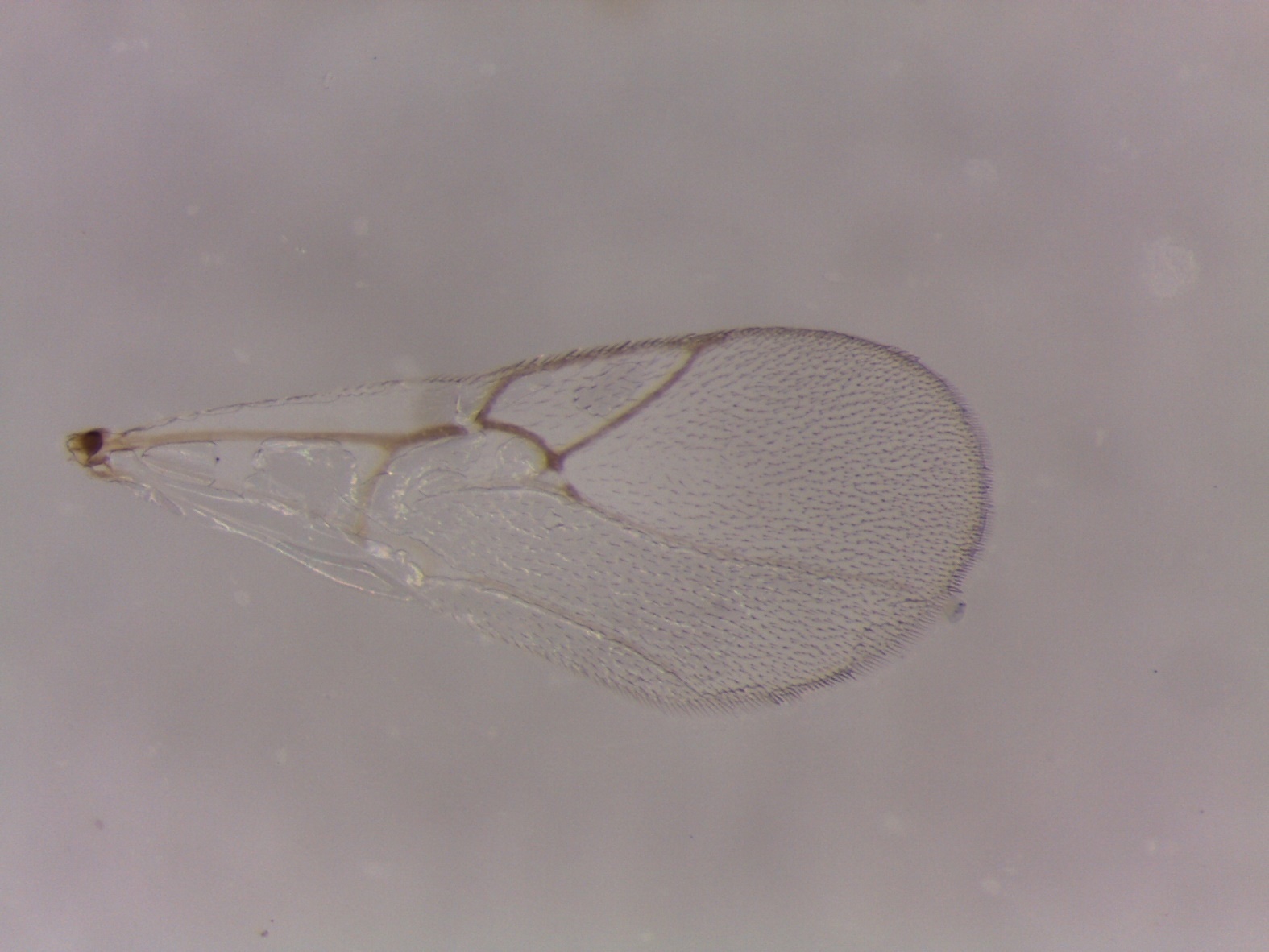

Figure S13. *Ceroptres* sp. 1-32-33-34 – 852-042-44 – female from *Andricus quercuspetiolicola* on *Quercus bicolor* in Coralville, IA.

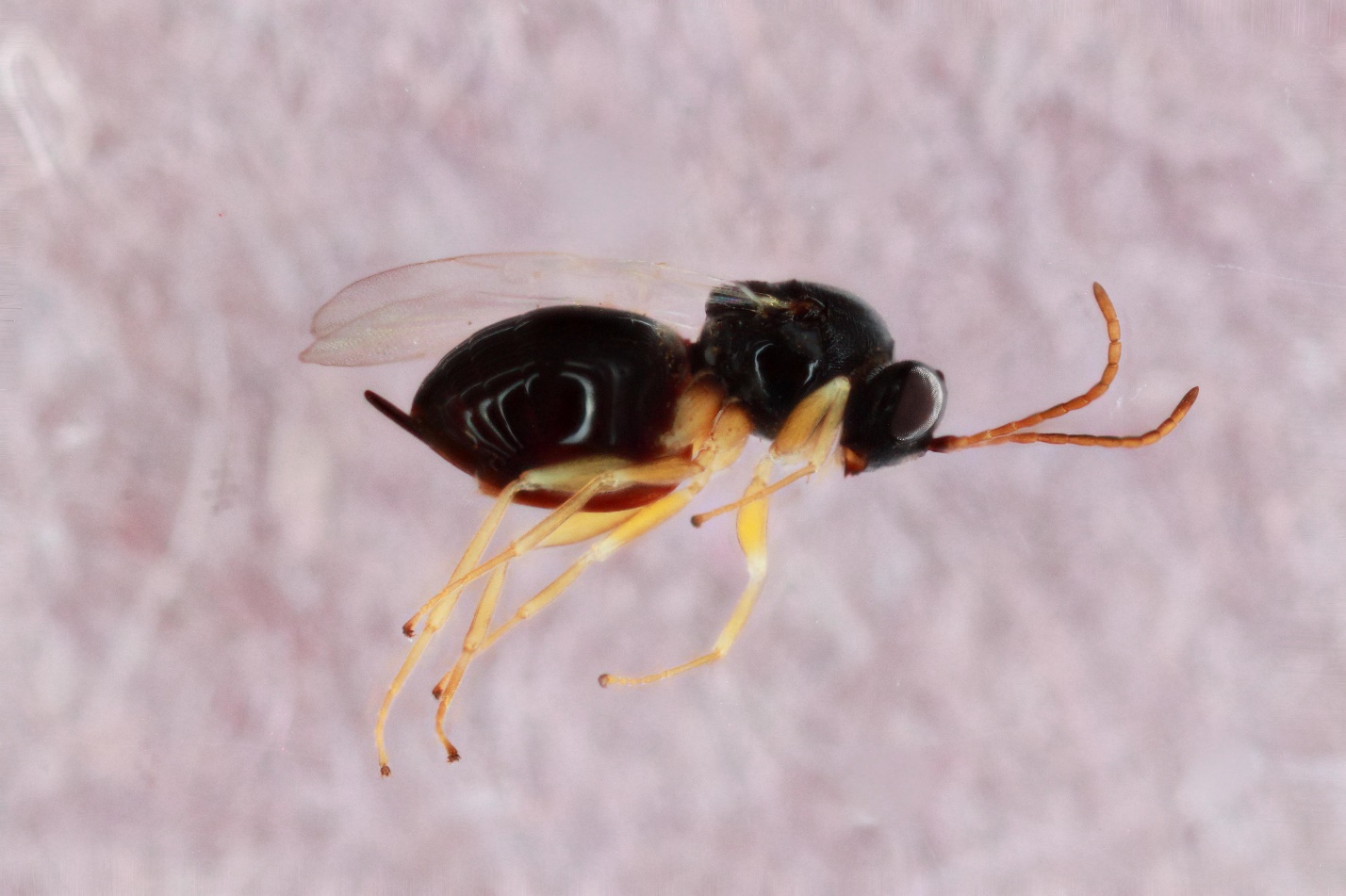

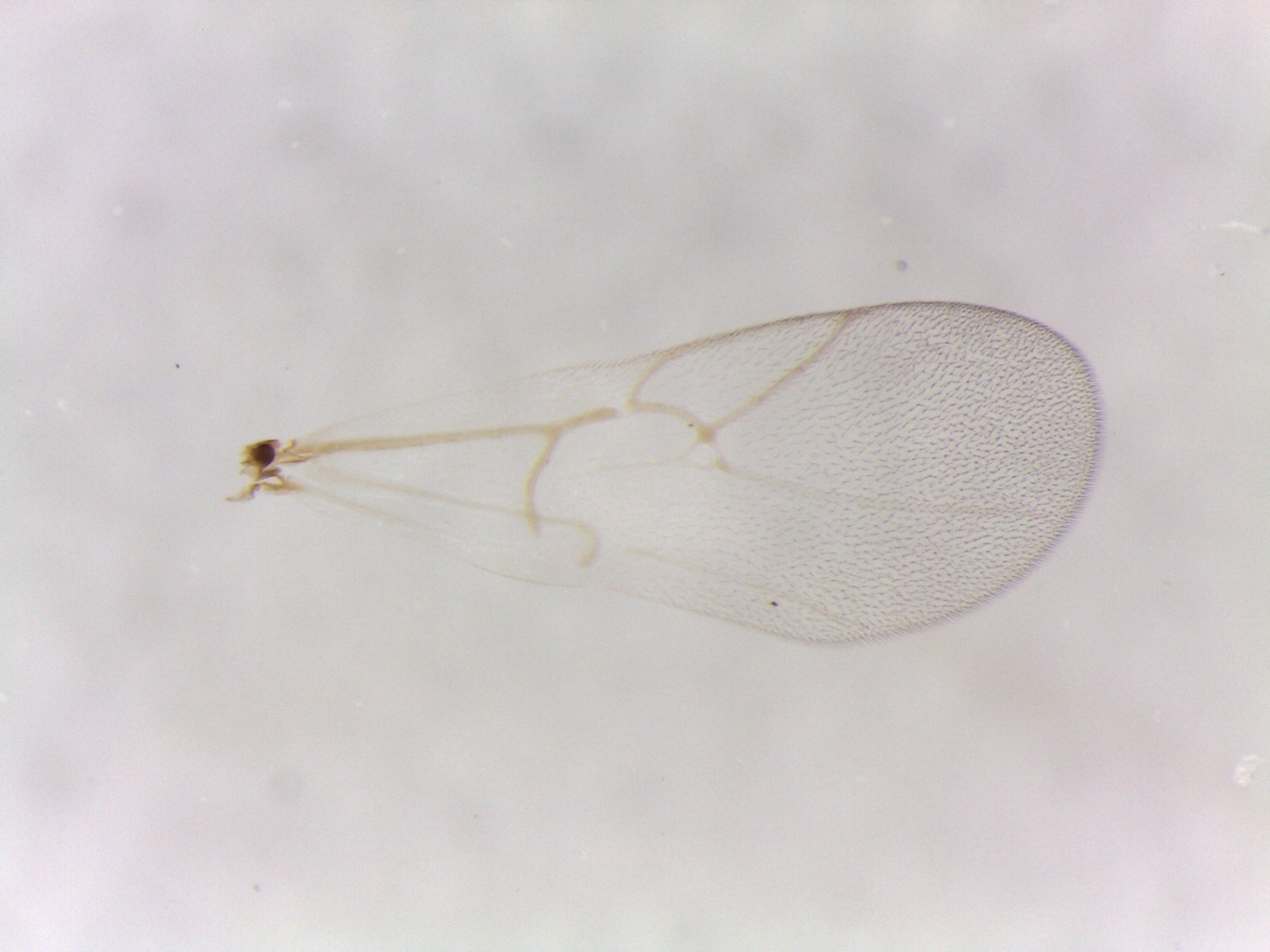

Figure S14. *Ceroptres* sp. 1-32-33-34 – 970-042-11 – female from *Andricus quercuspetiolicola* on *Quercus bicolor* in Iowa City, IA.
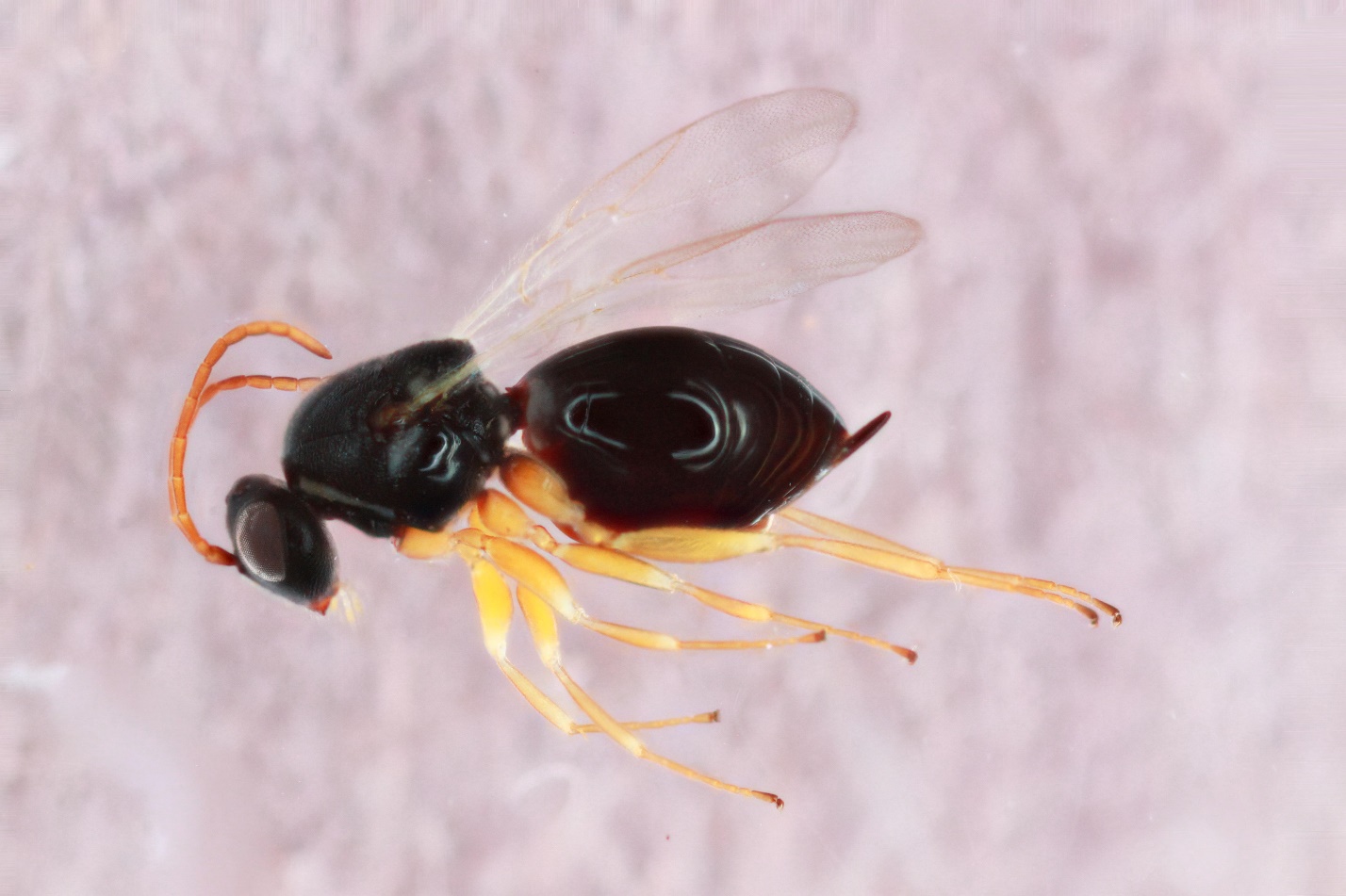

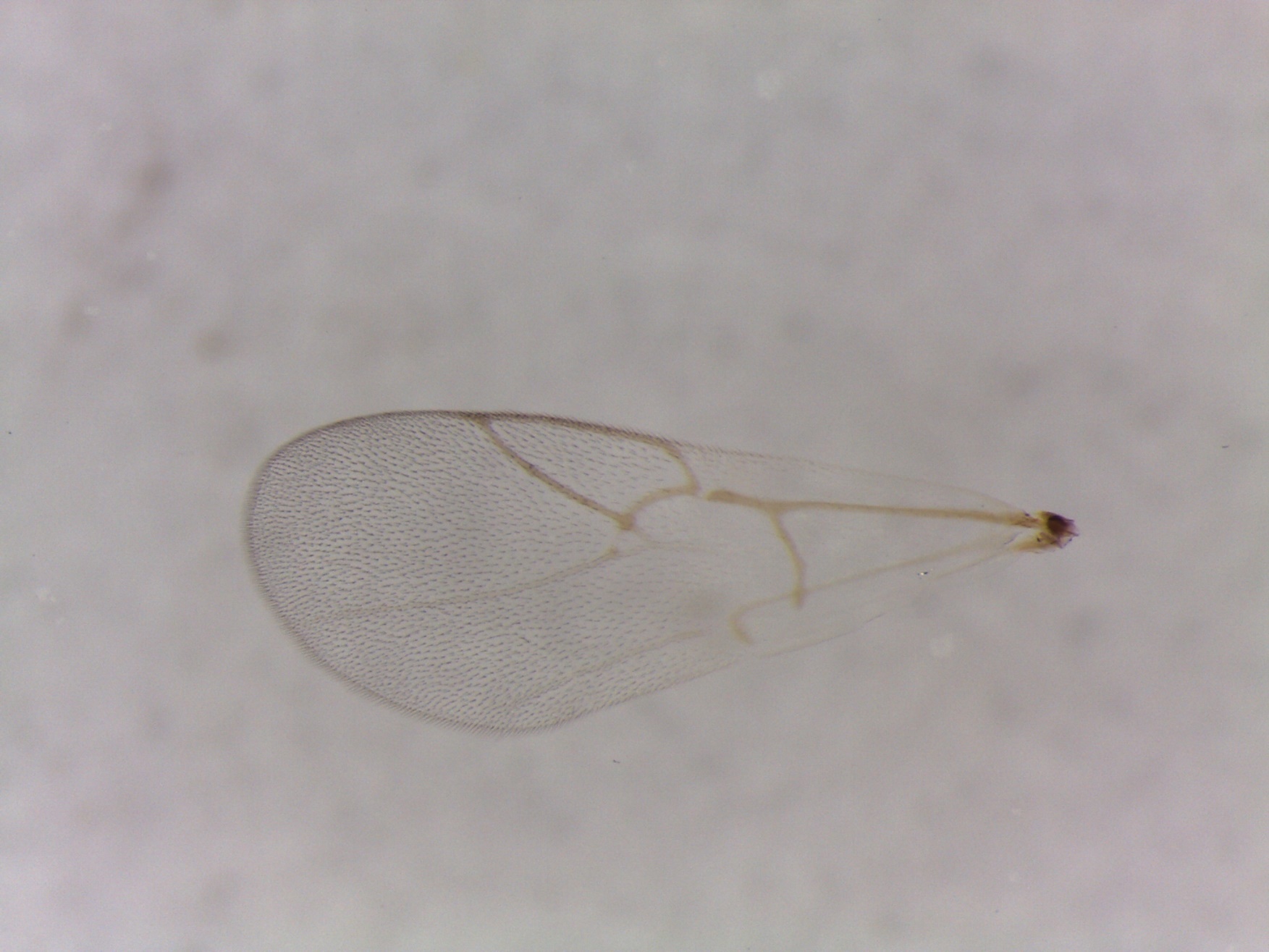

Figure S15. *Ceroptres* sp. 1-32-33-34 – 700-006-1 –male from *Andricus quercusfrondosus* on *Quercus bicolor* in Iowa City, IA.

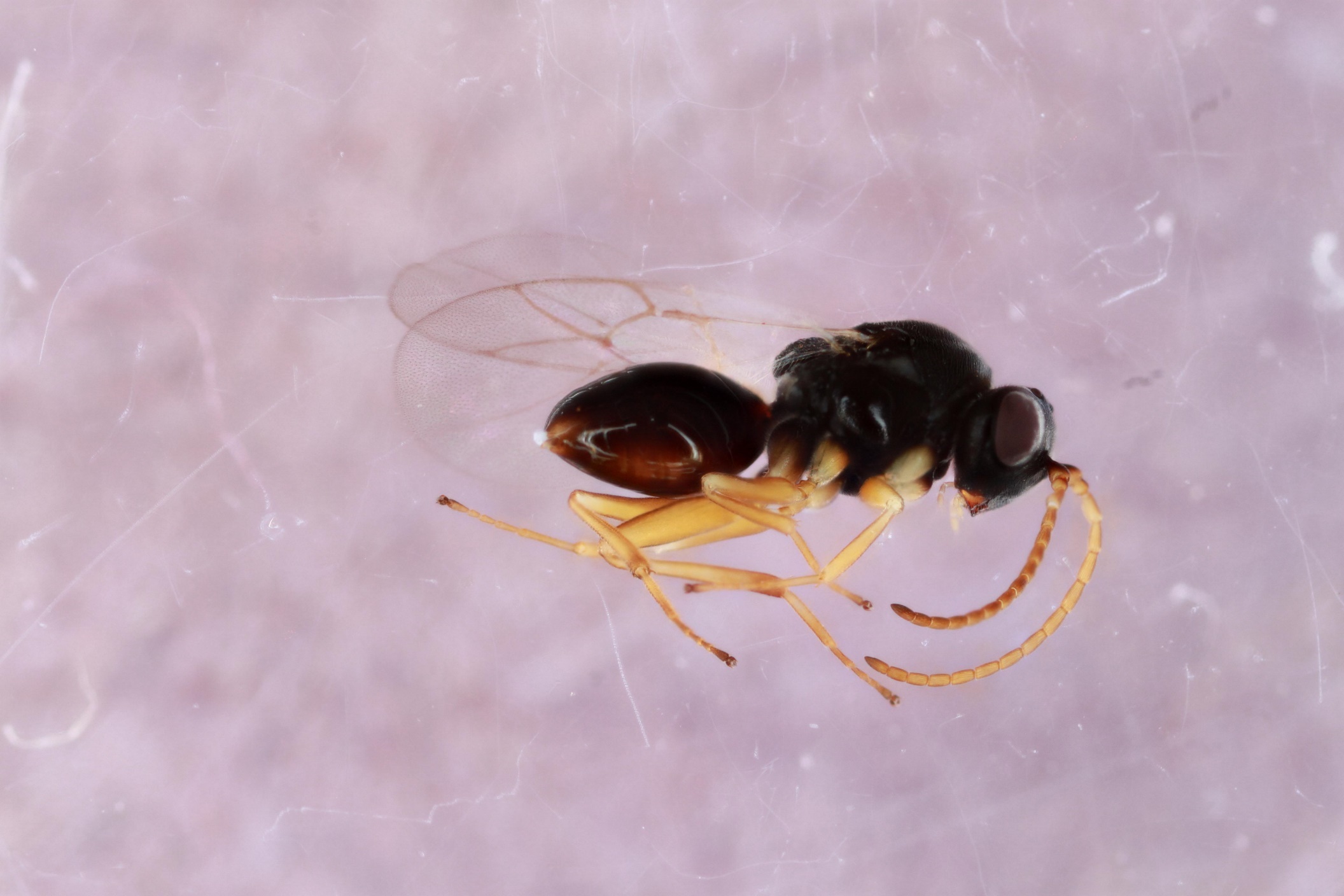

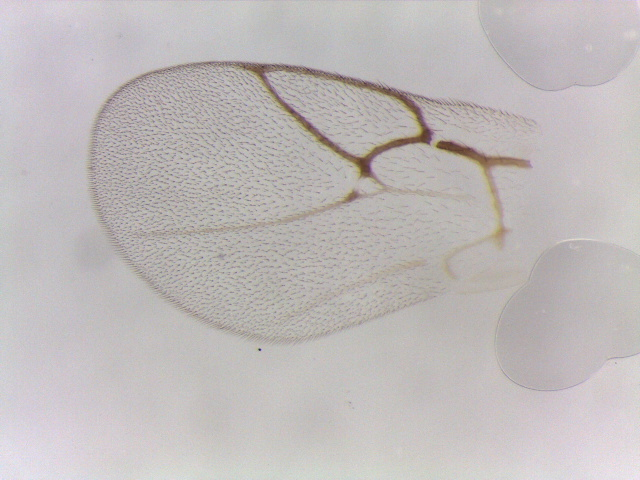

Figure S16. *Ceroptres* sp. 1-32-33-34 – 319-042-71 – female from *Andricus quercuspetiolicola* on *Quercus bicolor* in Tiffin, IA.

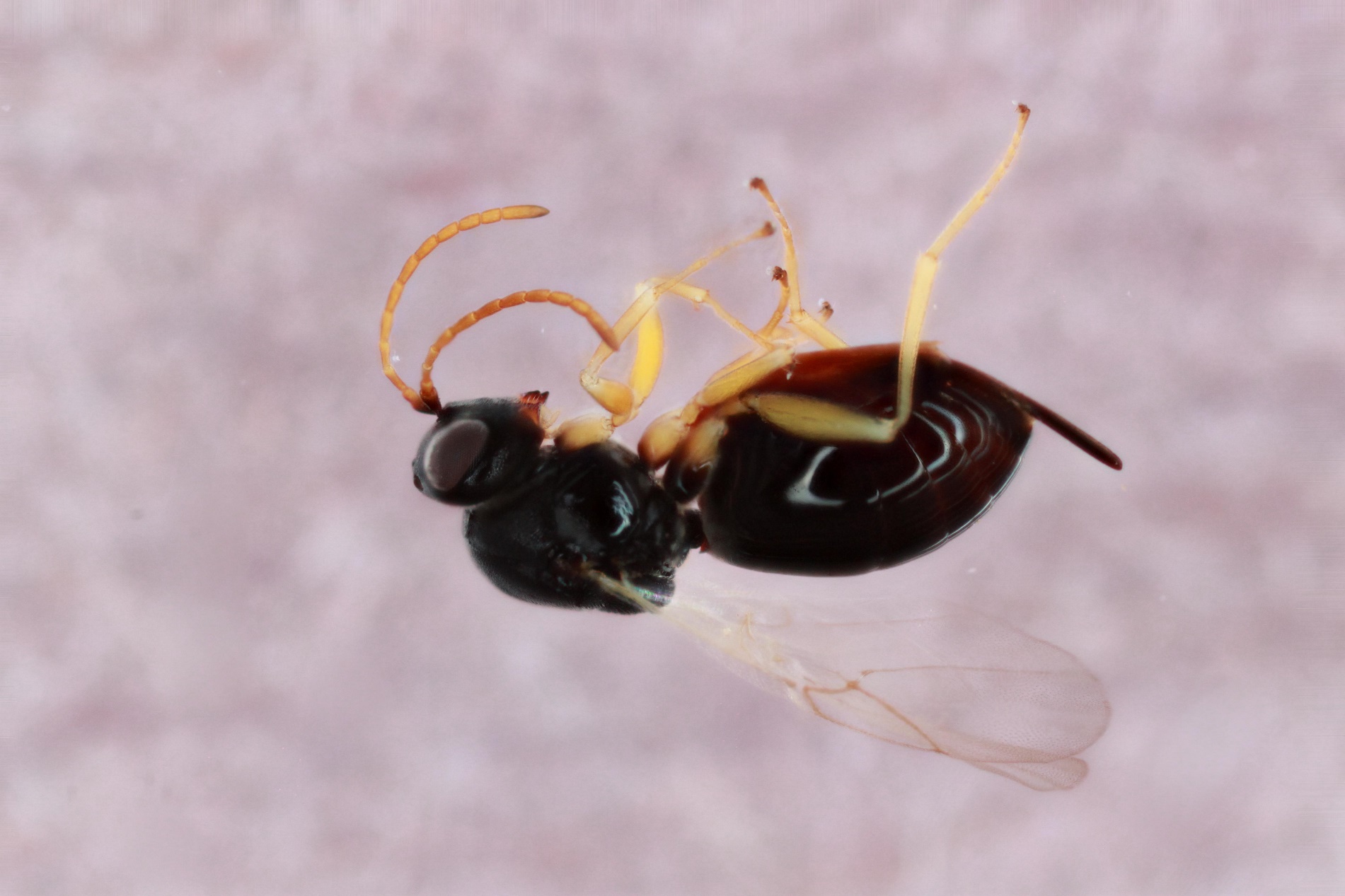

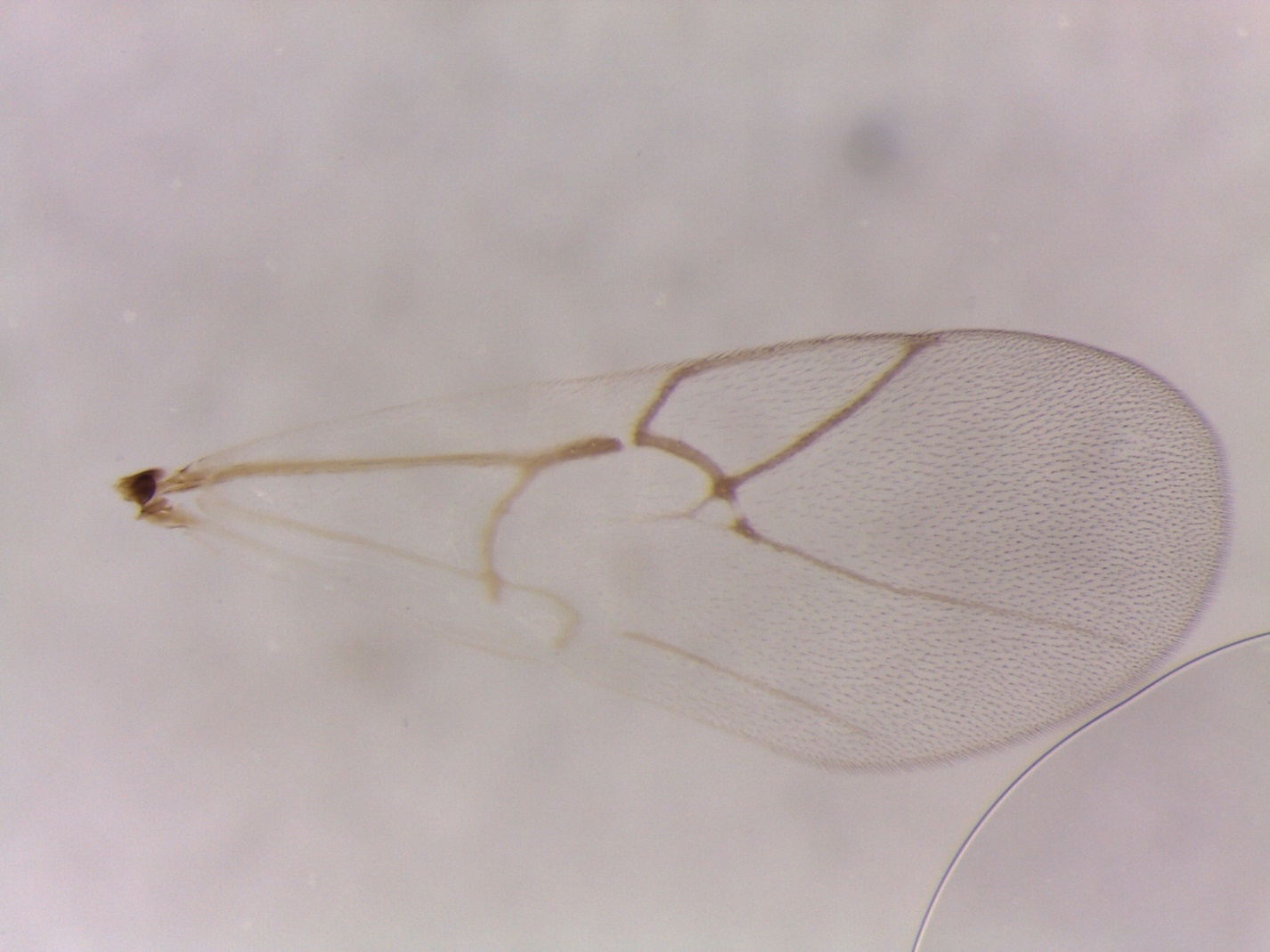

Figure S17. *Ceroptres* sp. 1-32-33-34 – 419-042-25 – female from *Andricus quercuspetiolicola* on *Quercus bicolor* in Iowa City, IA.

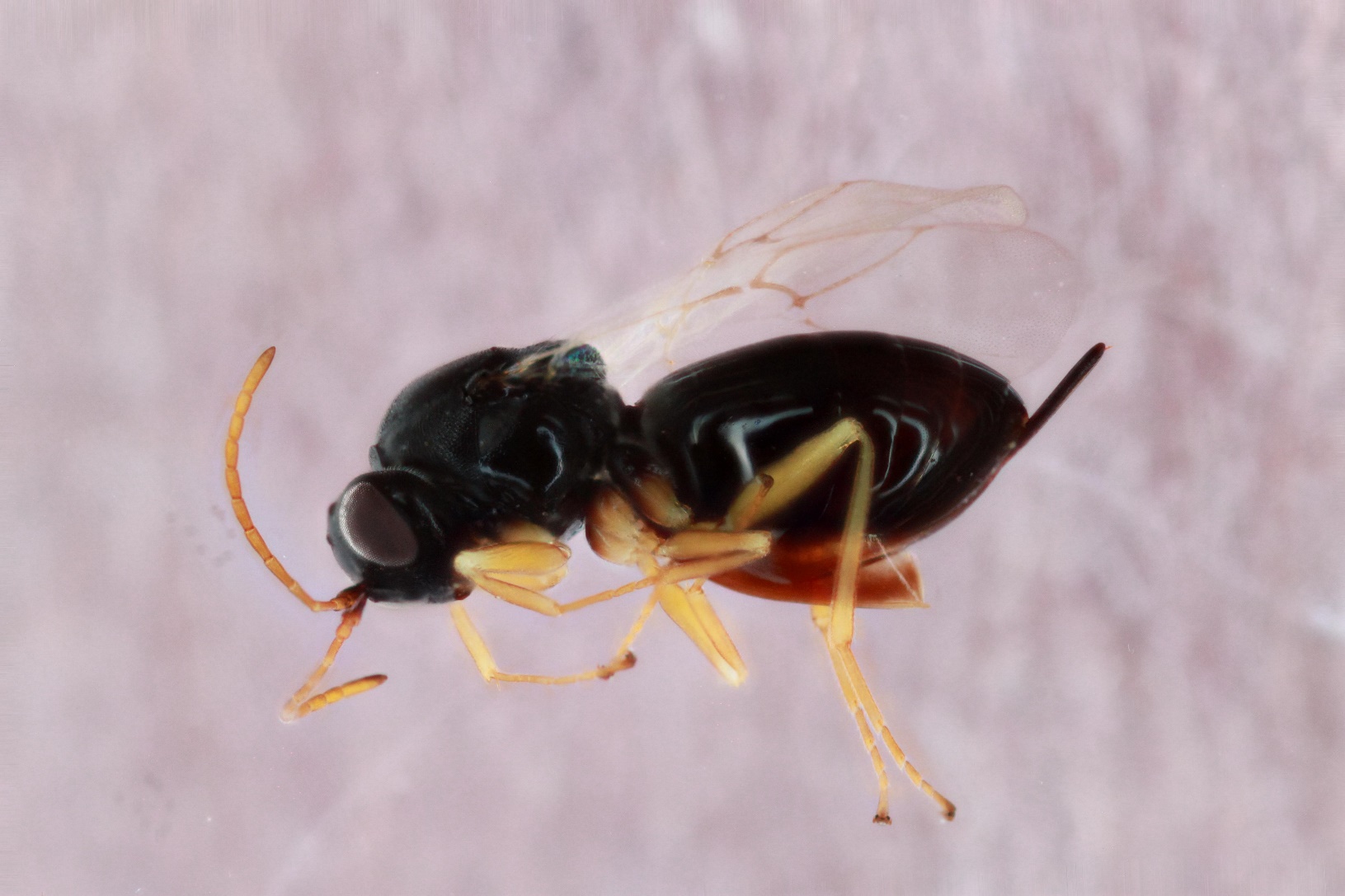

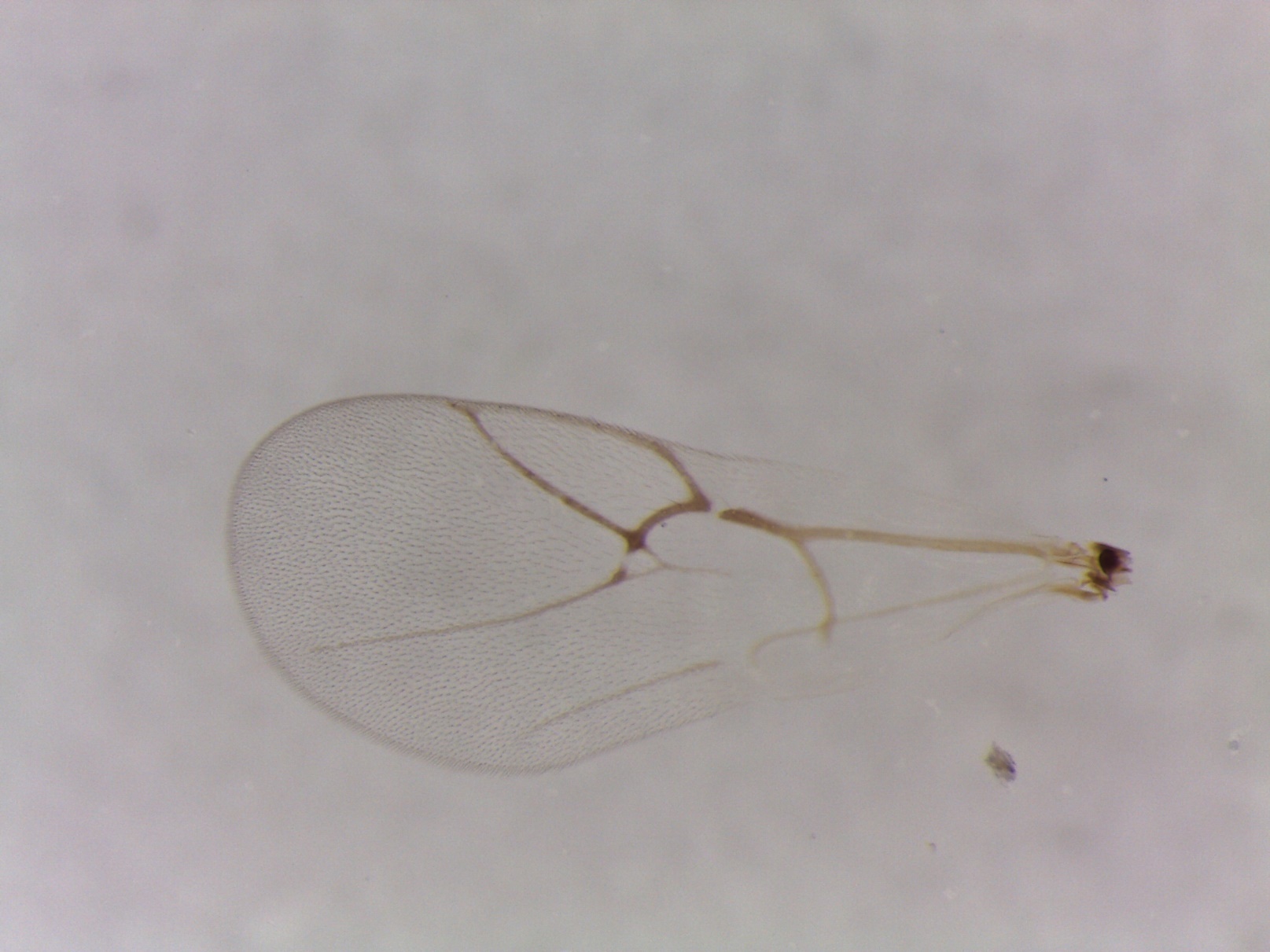

Figure S18. *Ceroptres* sp. 2 – 1574-104-15 – female from *Callirhytis pigra* on *Quercus velutina* in Vestal, NY.

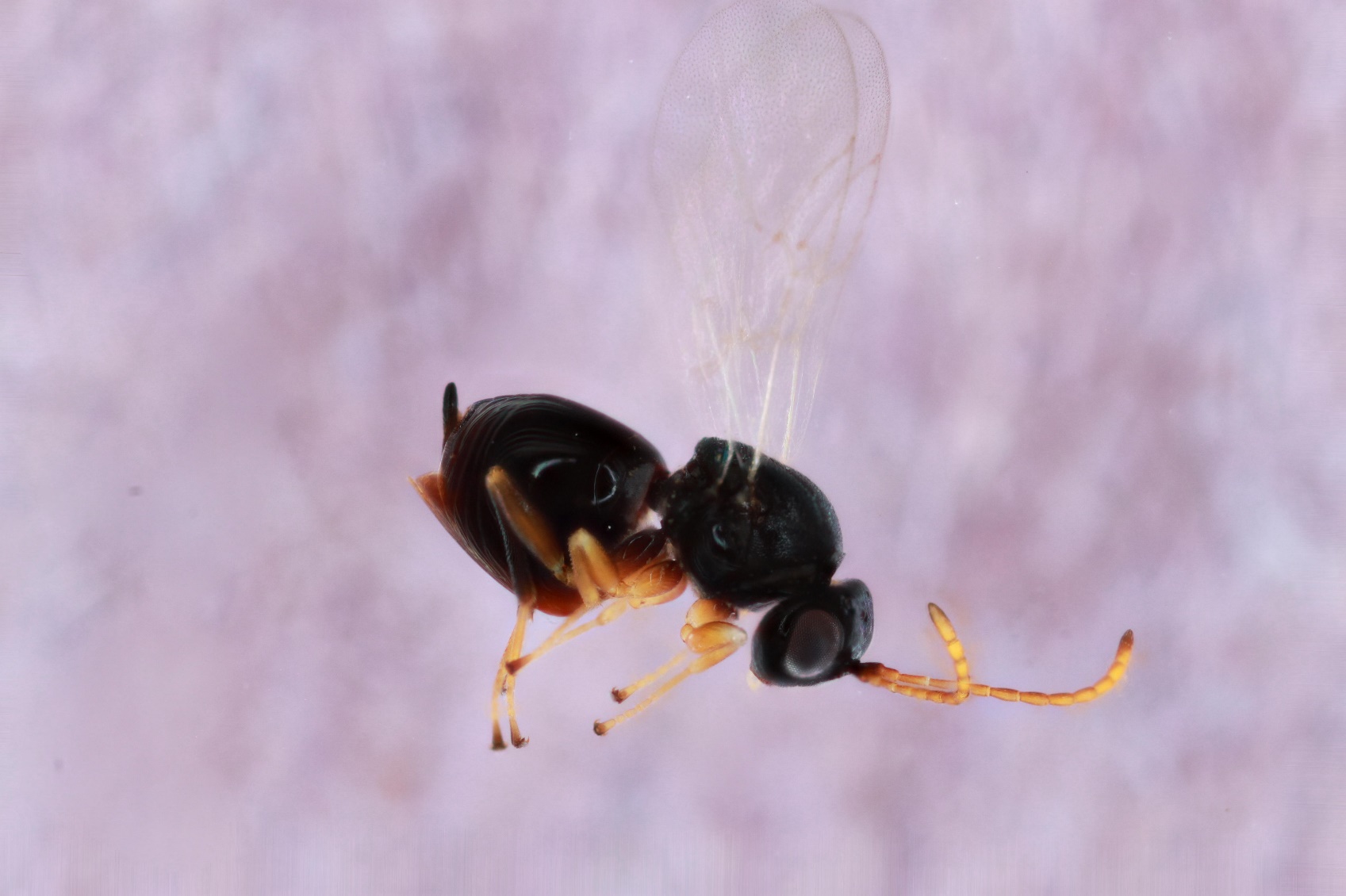

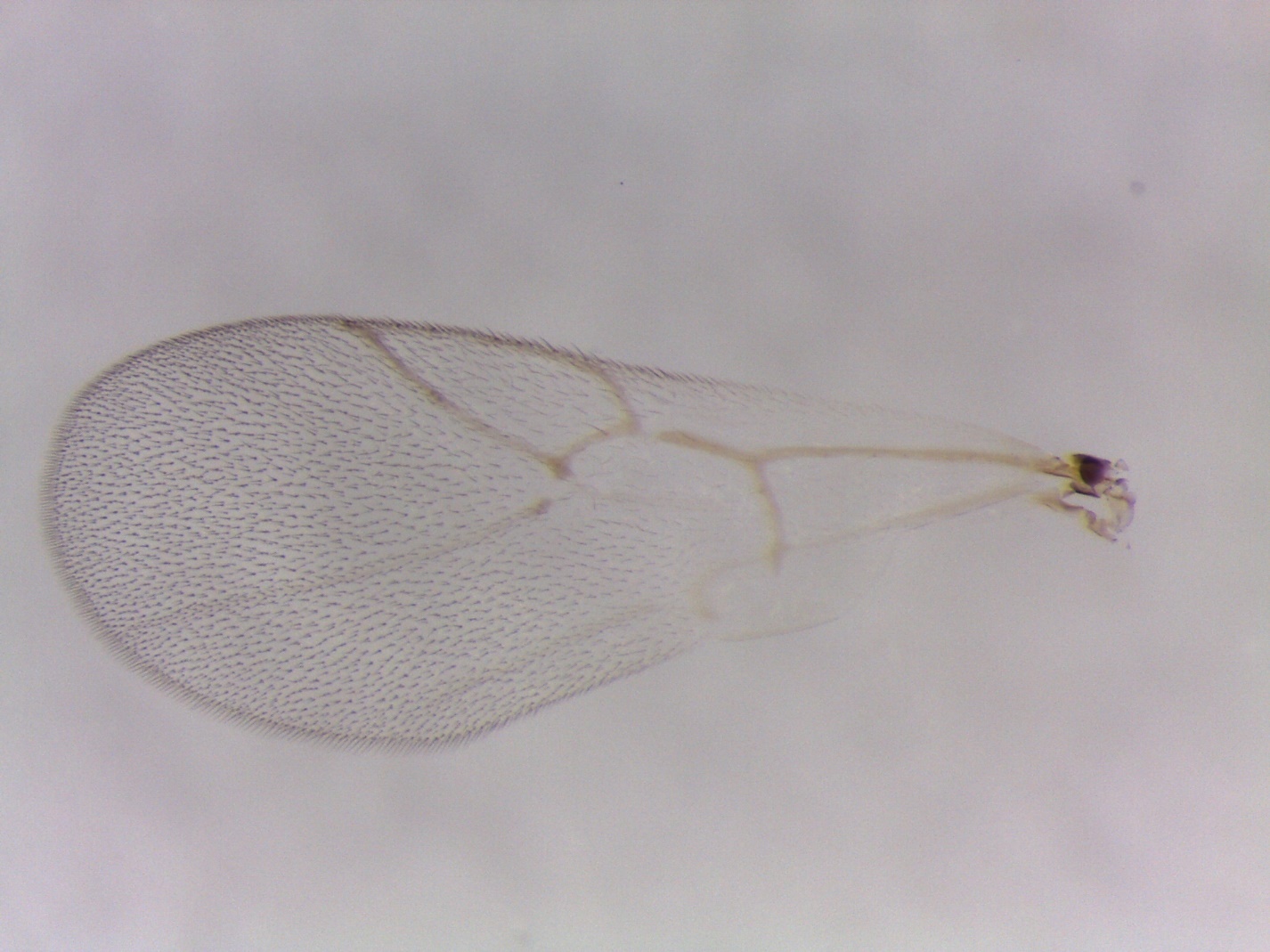

Figure S19. *Ceroptres* sp. 3 – 993-097-14 – female from *Melikaiella ostensackeni* on *Quercus palustris* in Coralville, IA.

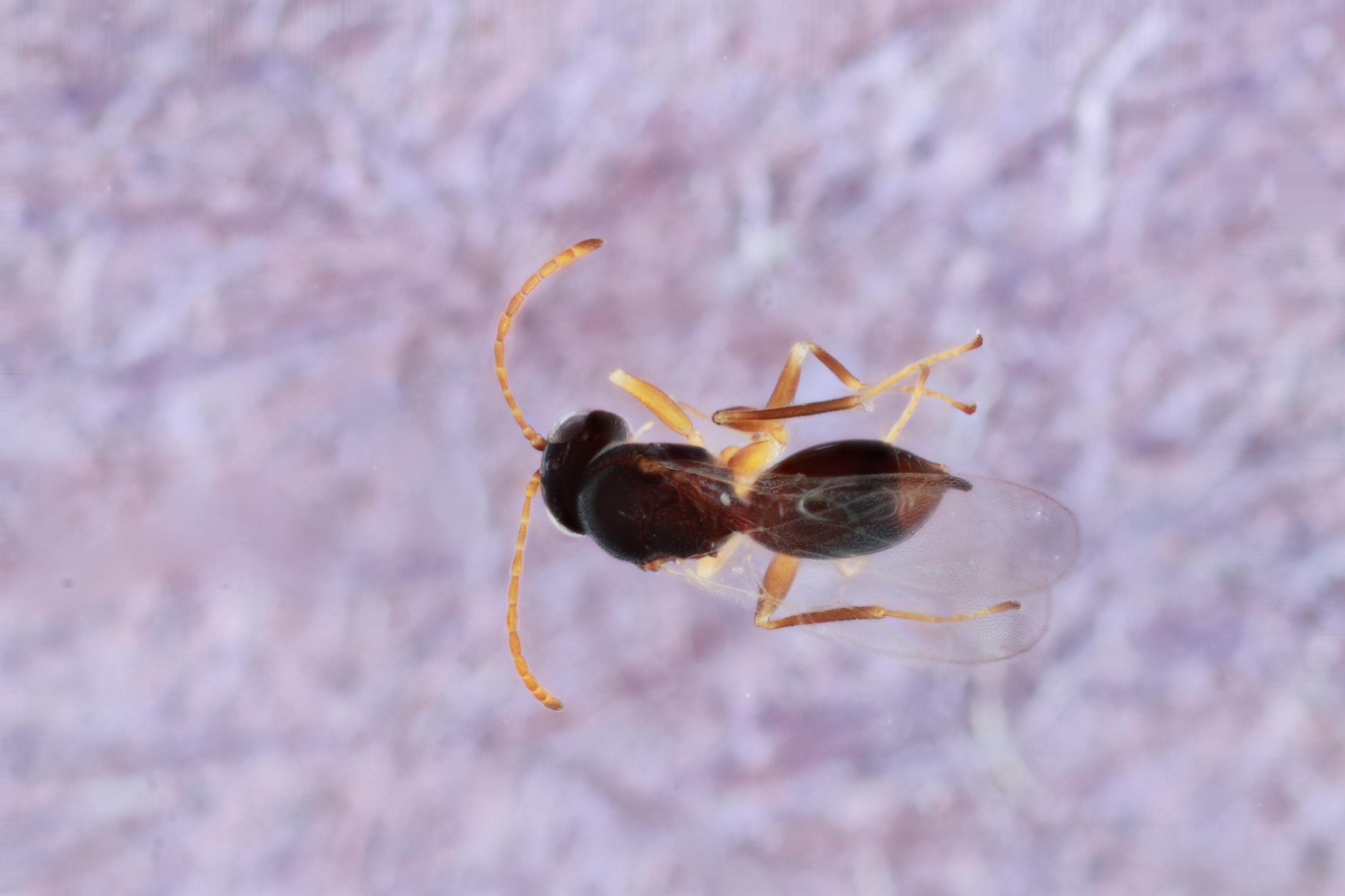

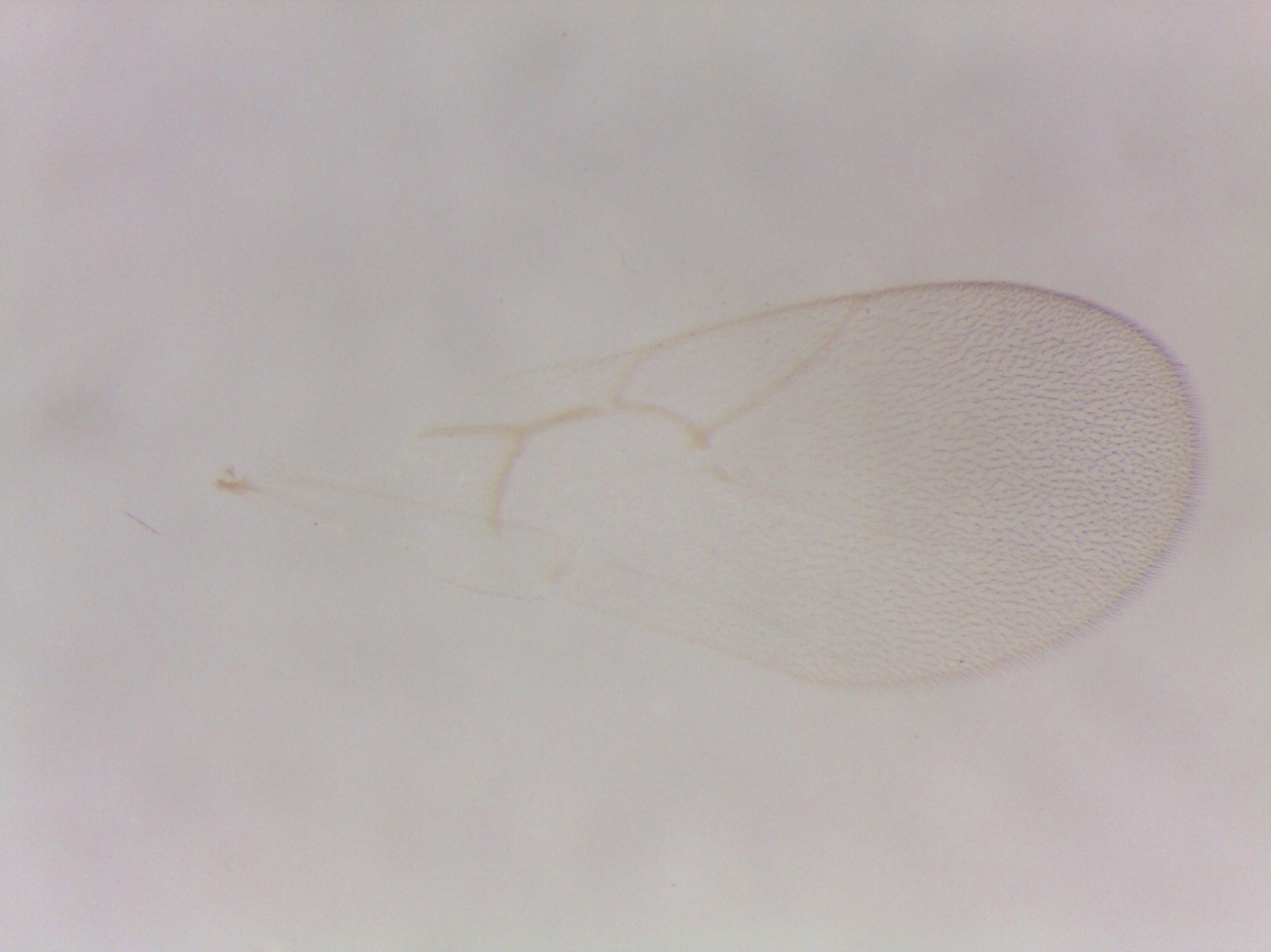

Figure S20. *Ceroptres* sp. 3 – 1563-062-5 – female from *Zopheroteras sphaerula* on *Quercus rubra* in Elkader, IA.

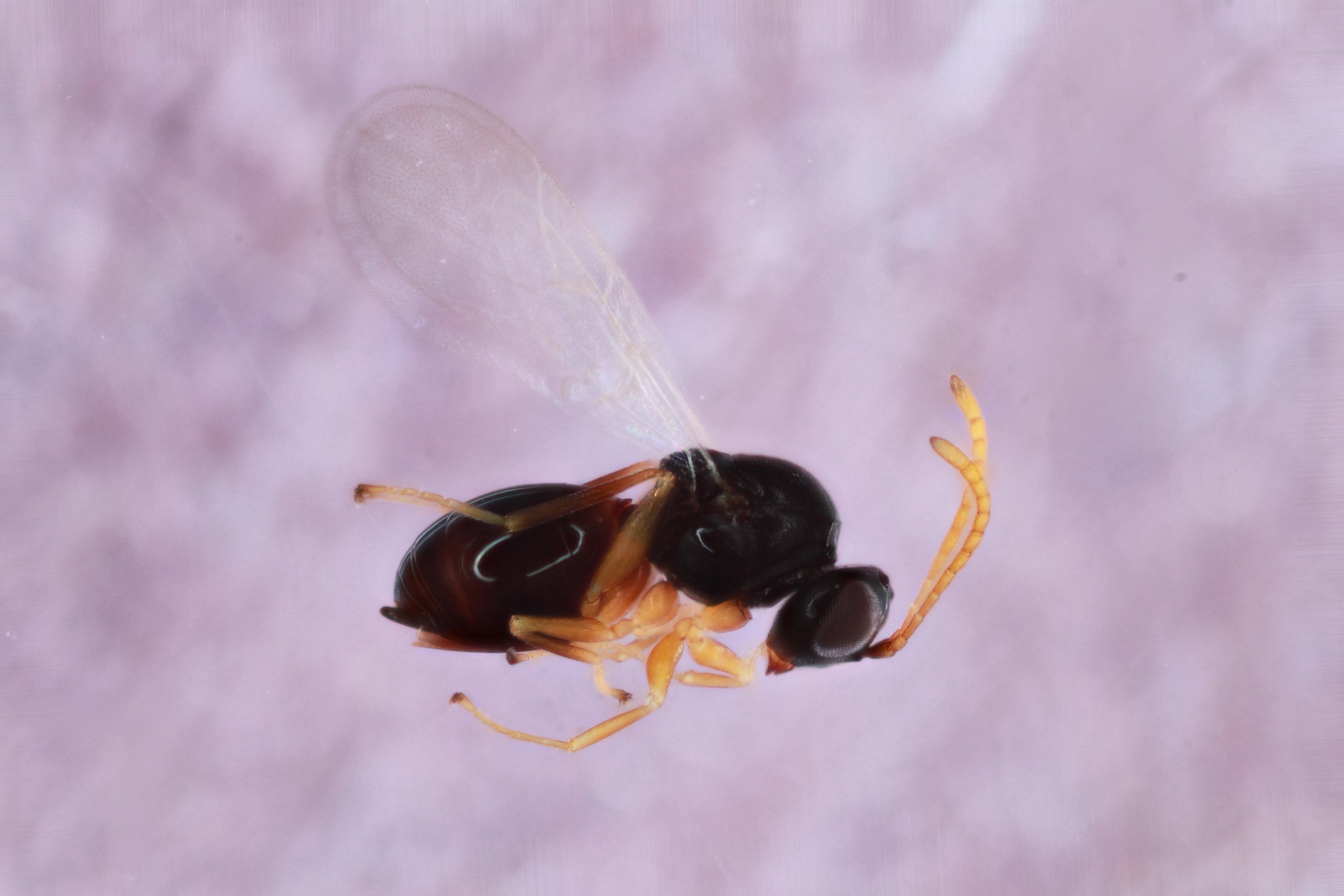

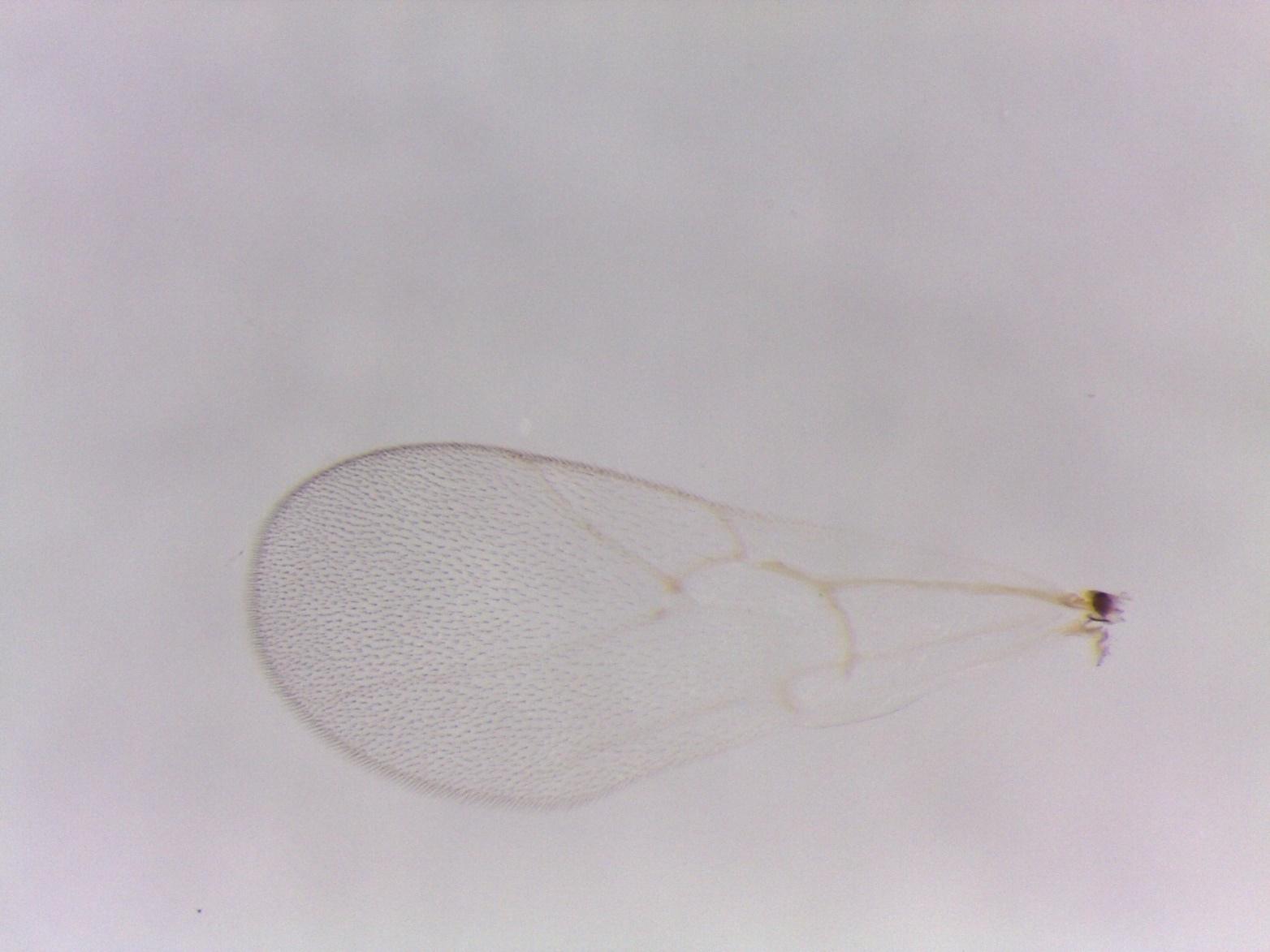

Figure S21. *Ceroptres* sp. 3 – 1078-102-1 – female from *Callirhytis quercuspunctata* (sexgen) on *Quercus rubra* in Traverse City, MI.

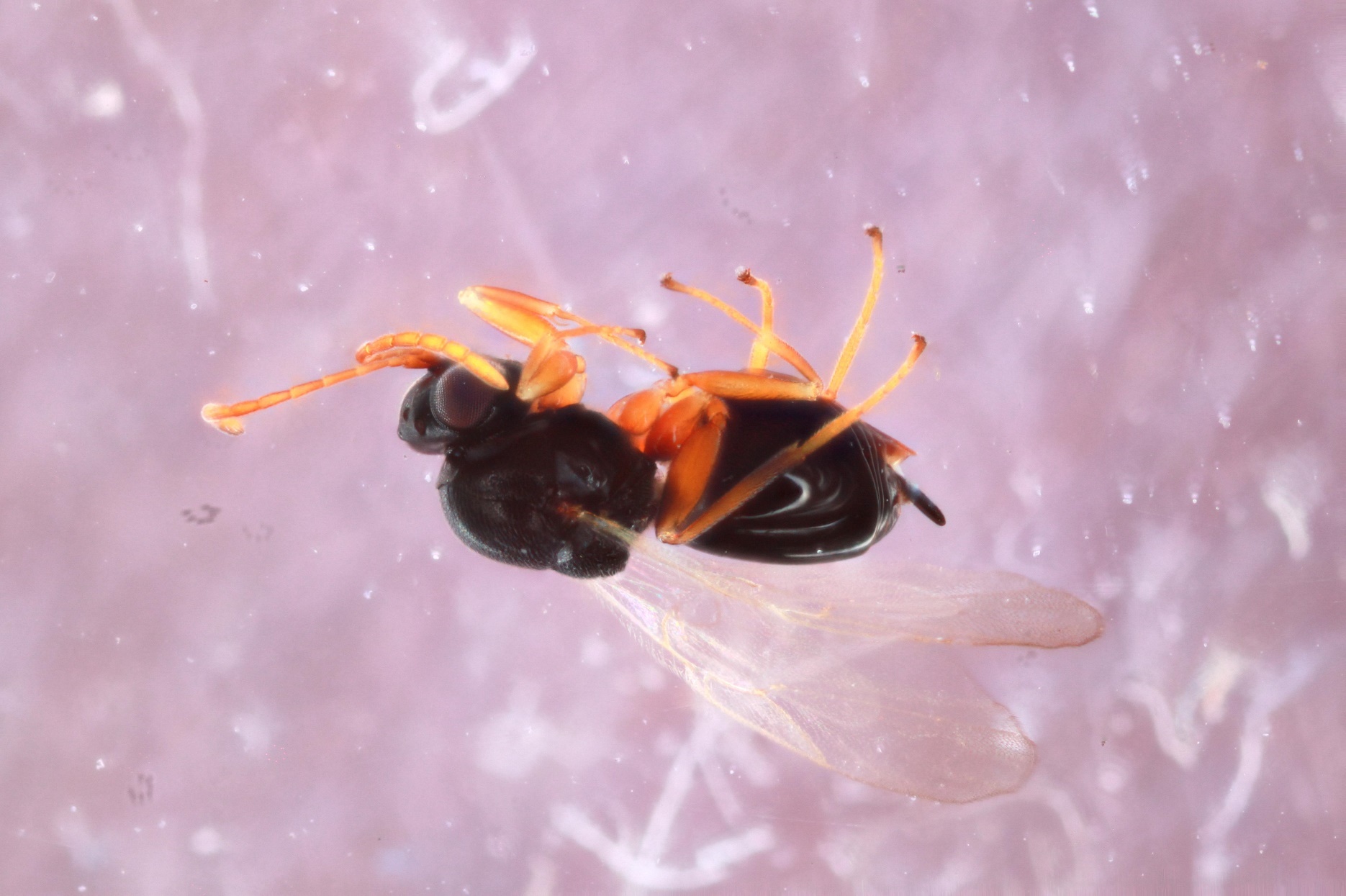

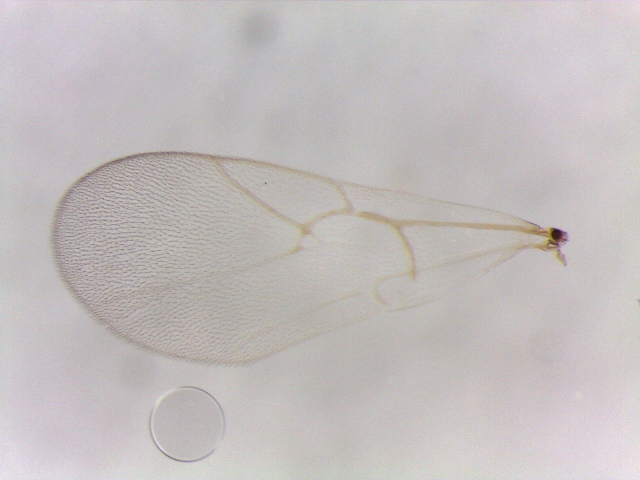

Figure S22. *Ceroptres* sp. 4 –694-033-3 – female from *Neuroterus vesicula* on *Quercus alba* in Iowa City, IA.

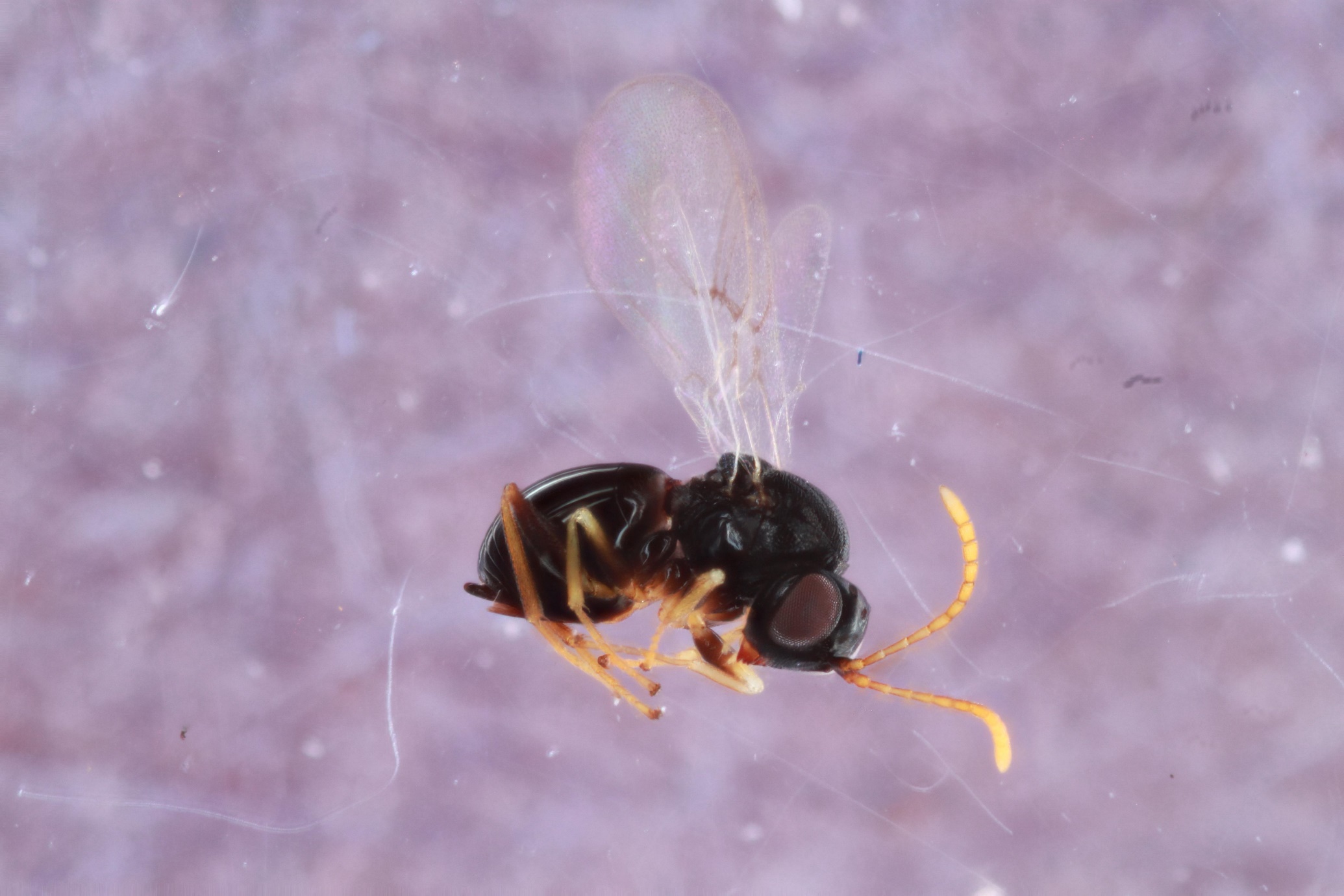

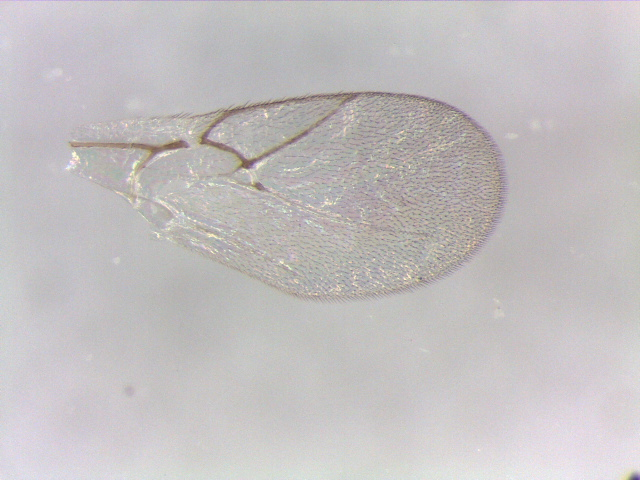

Figure S23. *Ceroptres* sp. 5a – 1450-051-8 – female from *Callirhytis quercusfutilis* on *Quercus alba* in Dodgeville, WI.

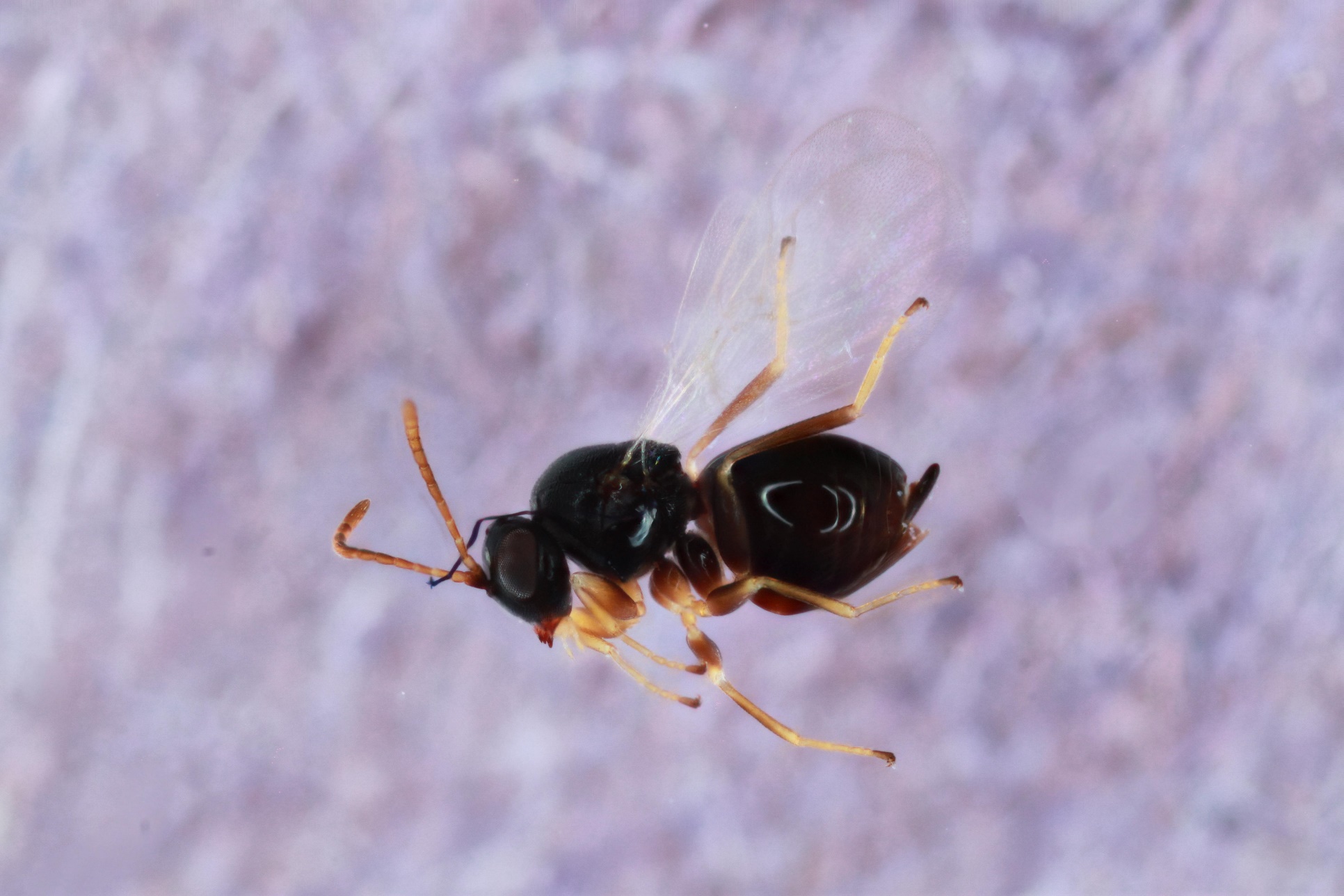

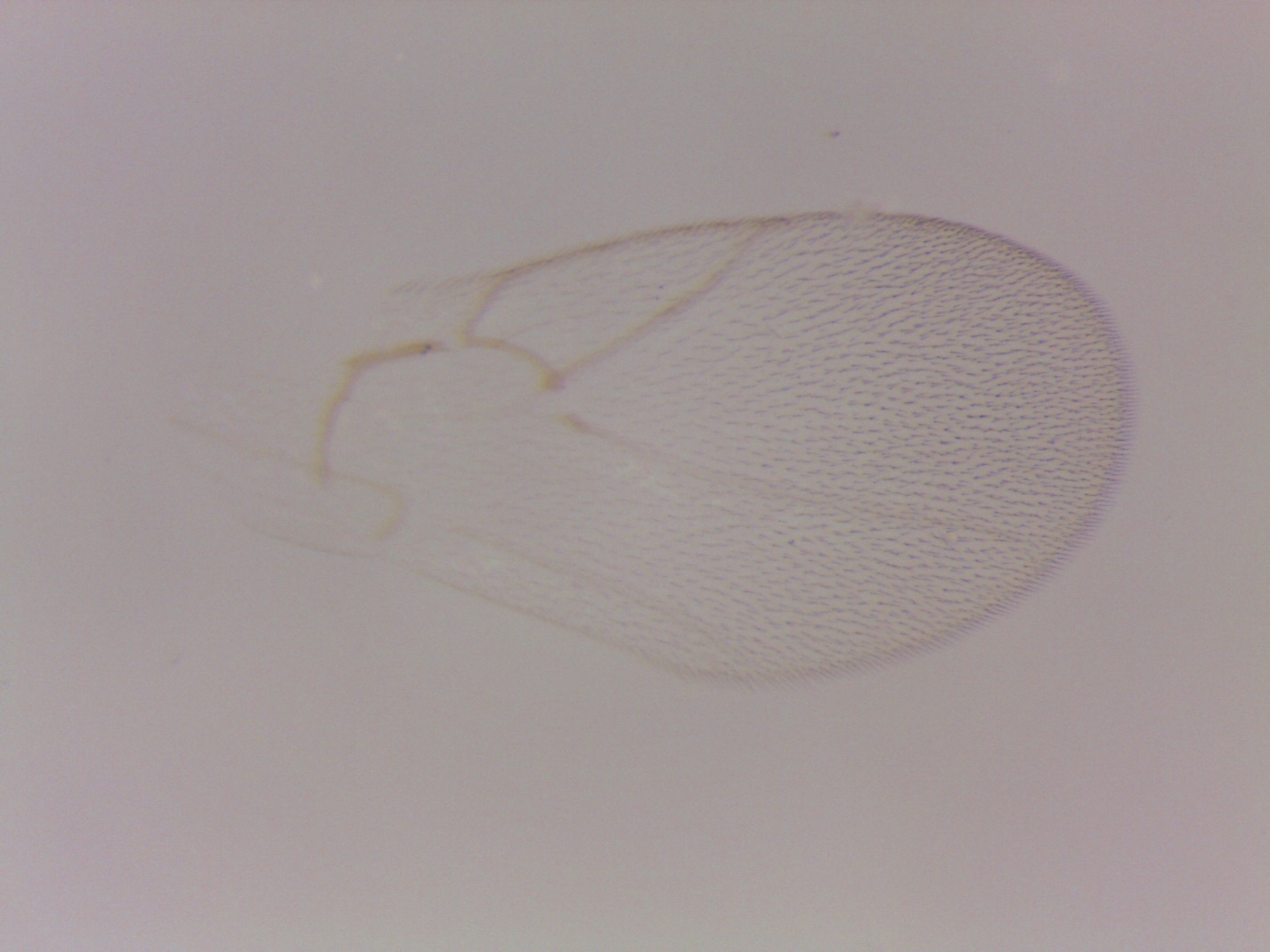

Figure S24. *Ceroptres* sp. 5b – 336-007-3 – female from *Andricus foliaformis* on *Quercus macrocarpa* in Iowa City, IA.

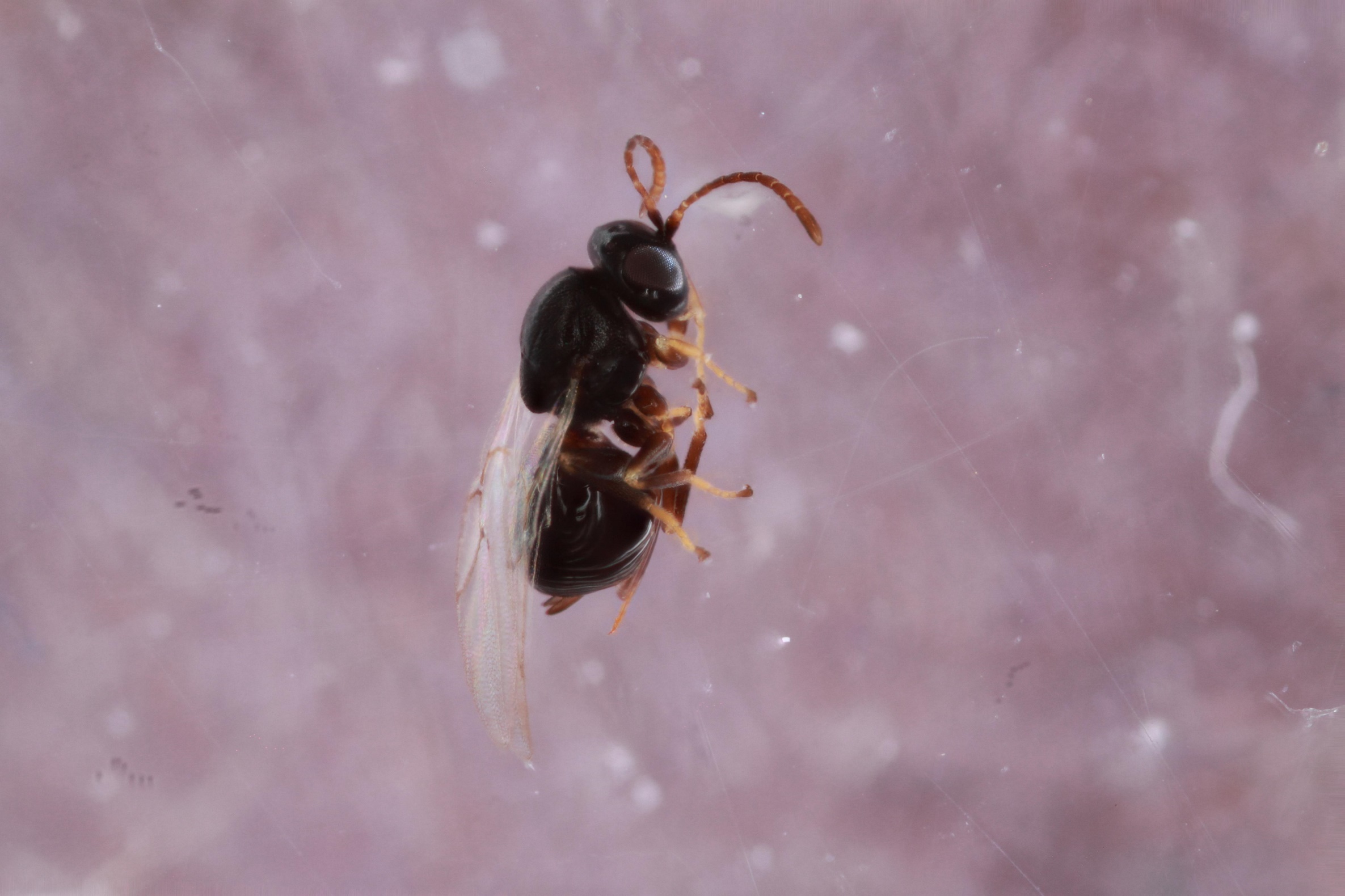

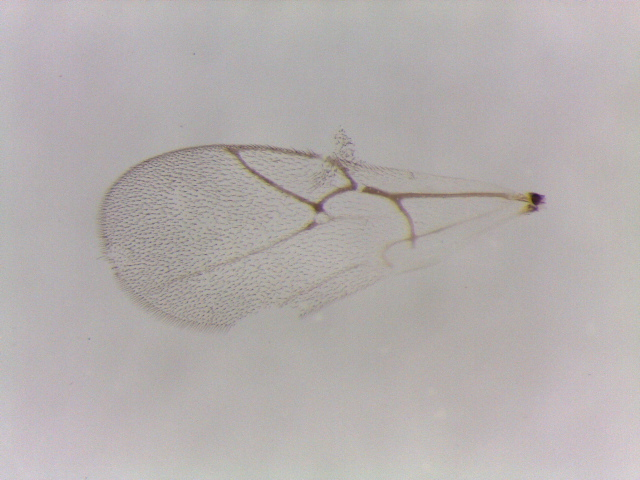

Figure S25. *Ceroptres* sp. 6a – 583-016-4a – female from *Melikaiella ostensackeni* on *Quercus palustris* in Walton, KY.

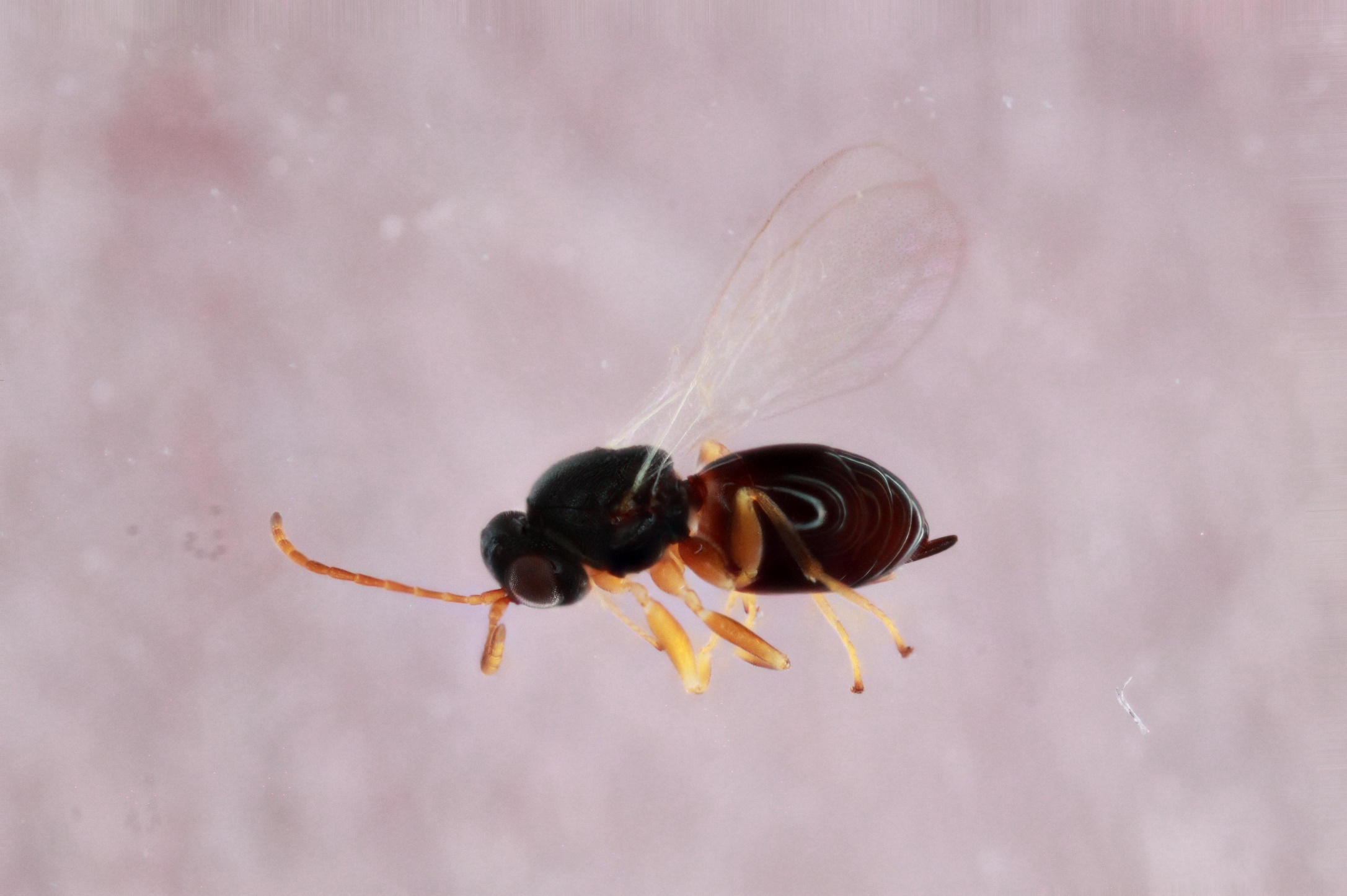

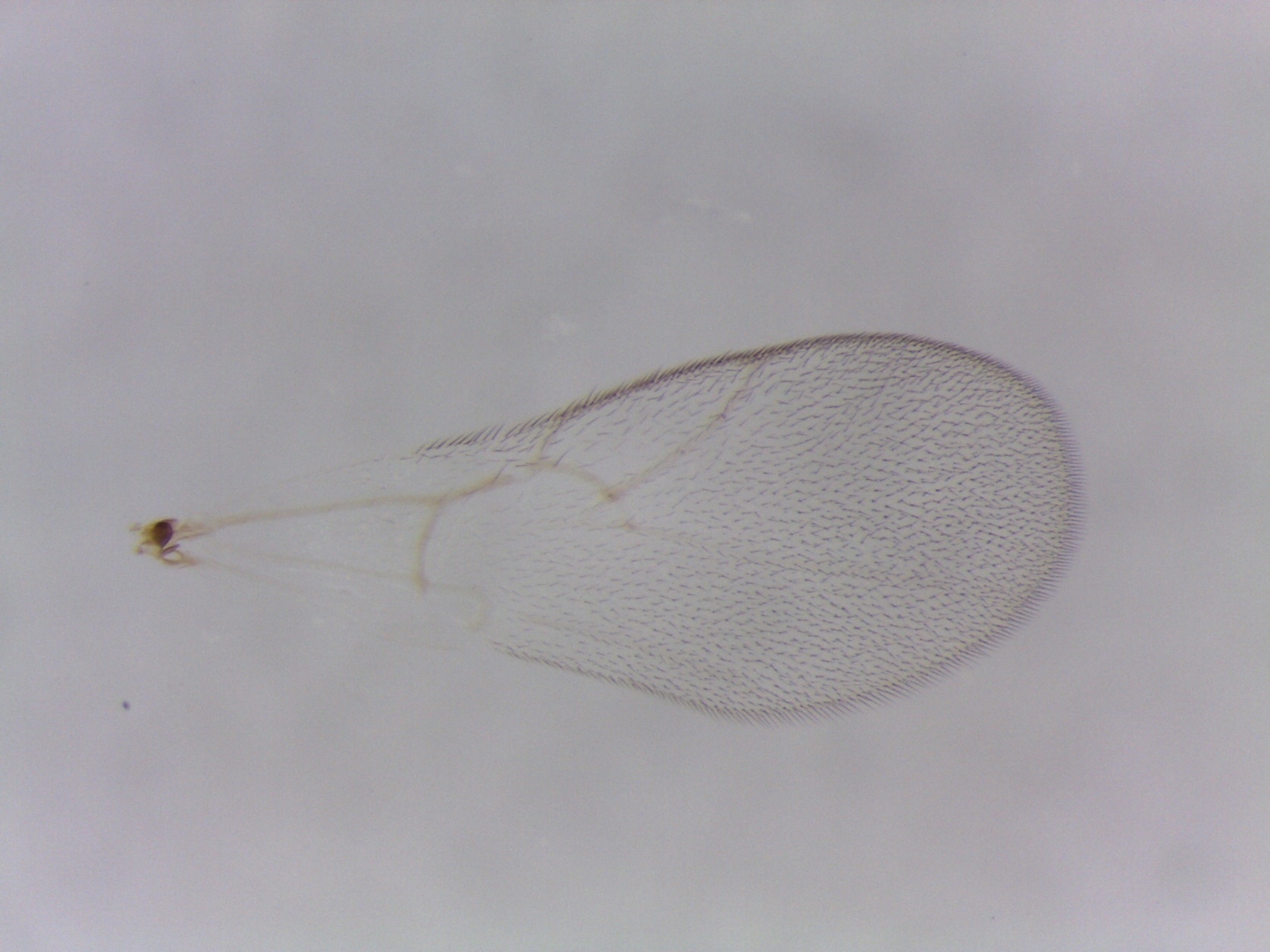

Figure S26. *Ceroptres* sp. 6b – 1076-101-17 – female from *Callirhytis quercusgemmaria* on *Quercus rubra* in Traverse City, MI.

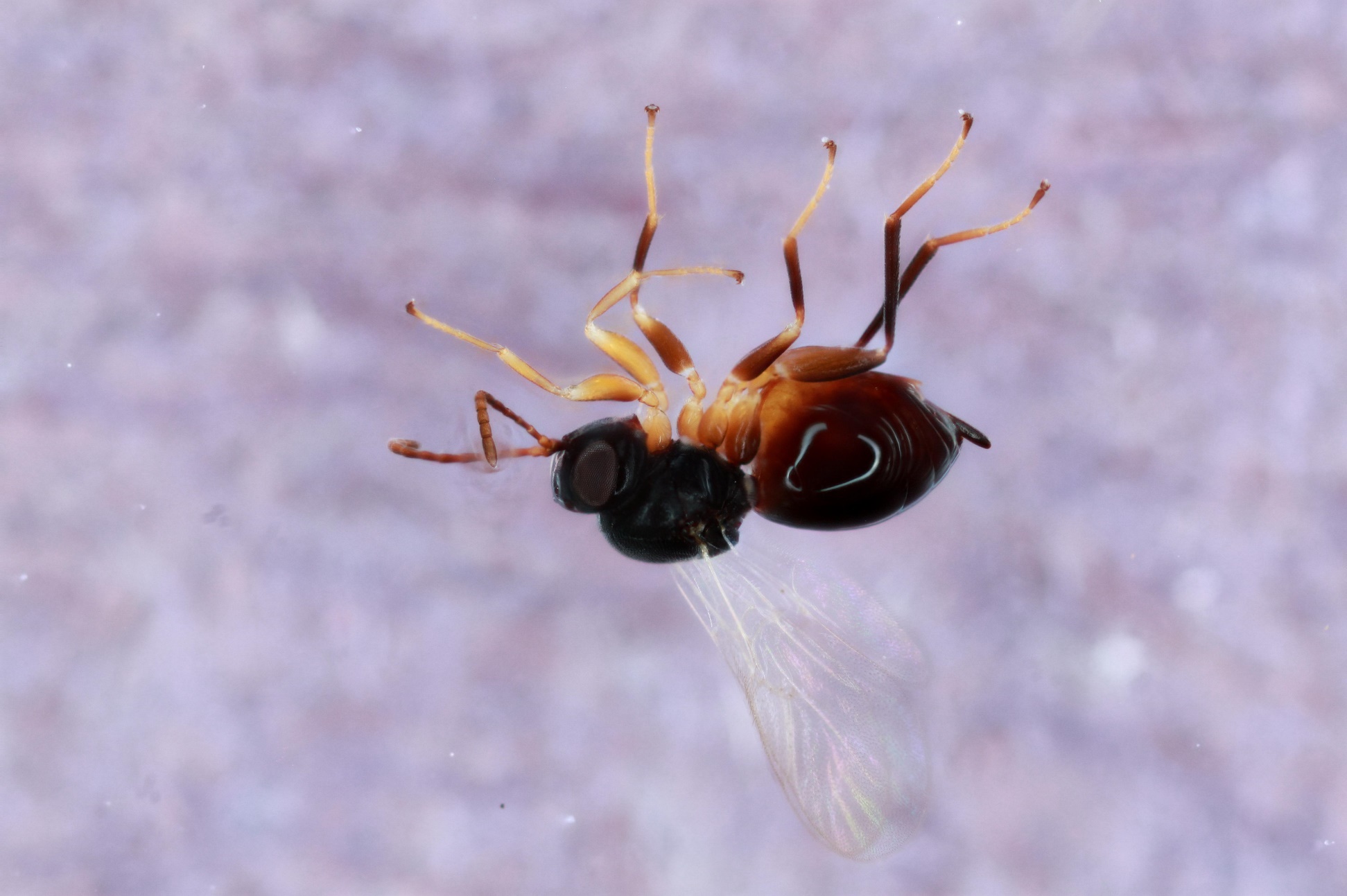

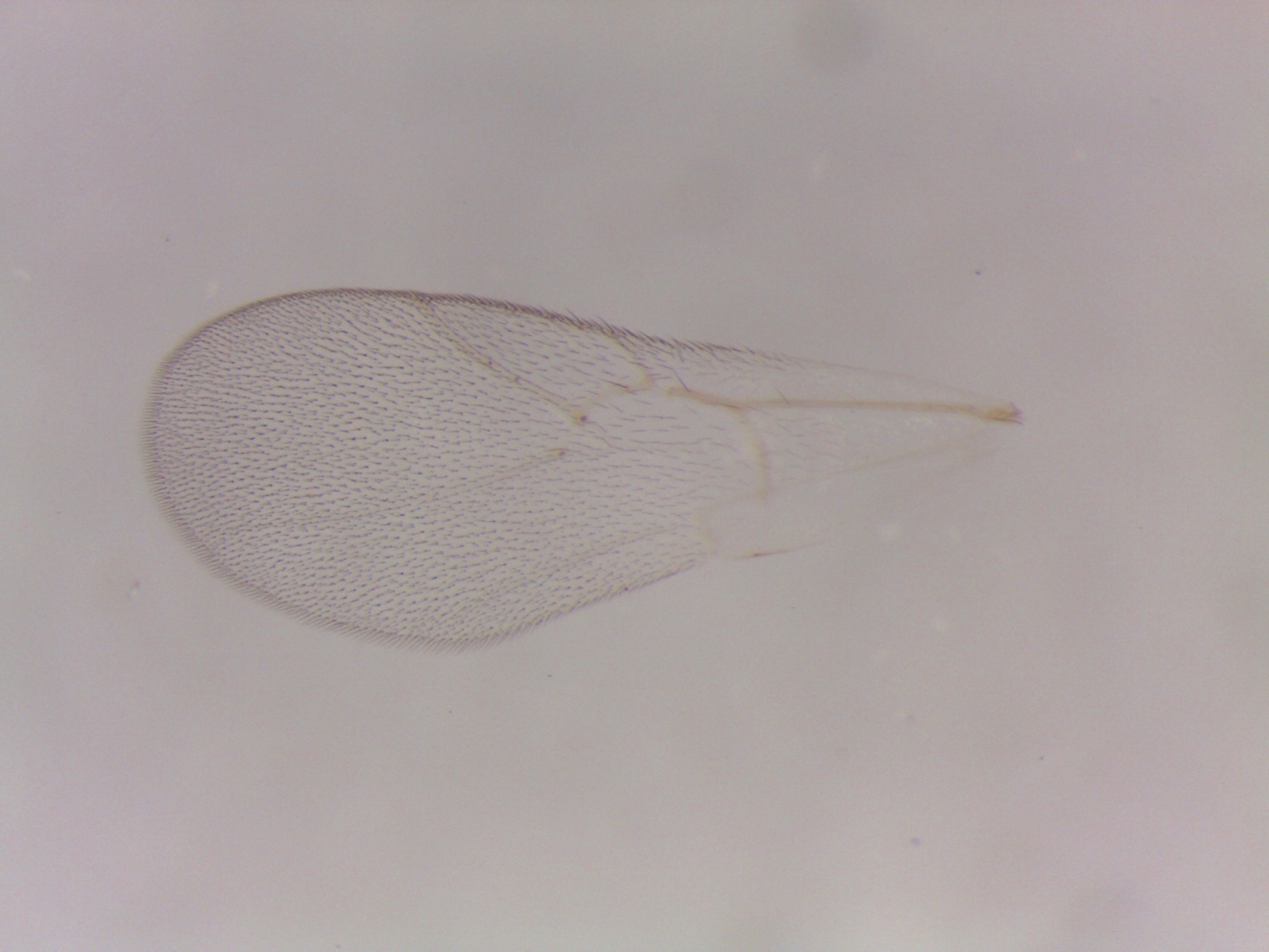

Figure S27. *Ceroptres* sp. 6c – 887-041-3 – female from *Callirhytis quercusventricosa* on *Quercus palustris* in Iowa City, IA.

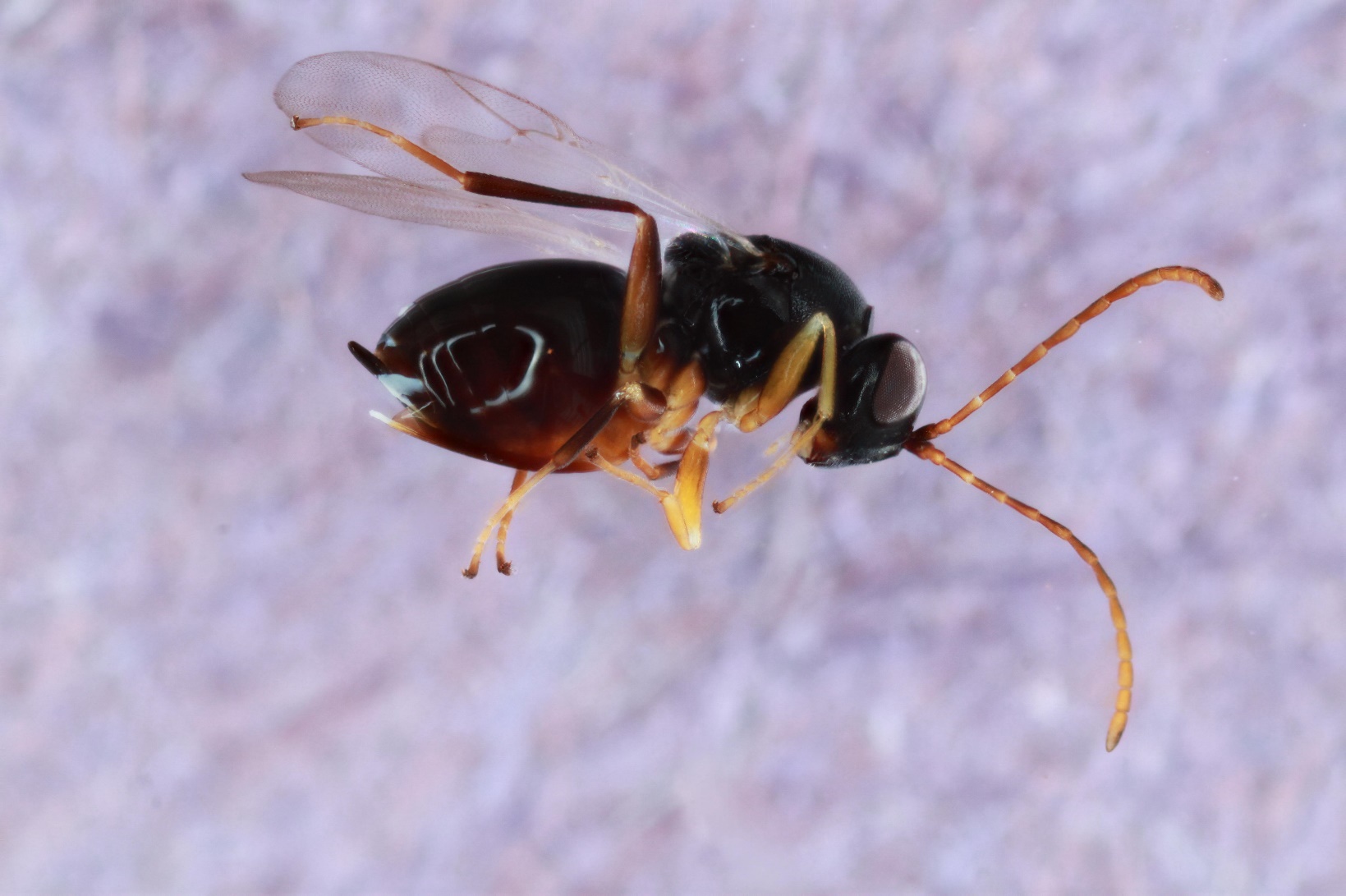

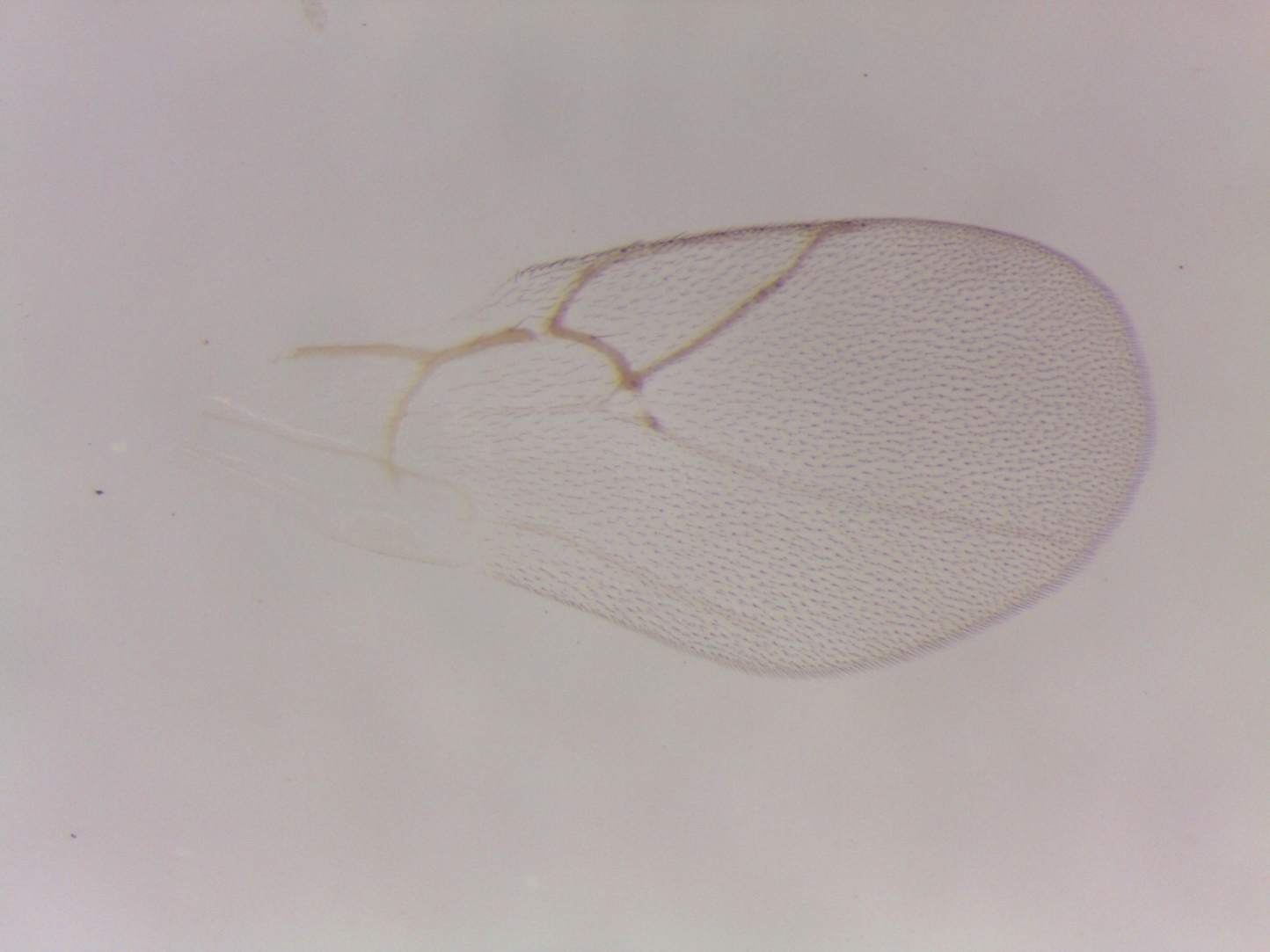

Figure S28. *Ceroptres* sp. 7-8-9 – 870-016-42 – female from *Melikaiella ostensackeni* on *Quercus rubra* in Iowa City, IA.

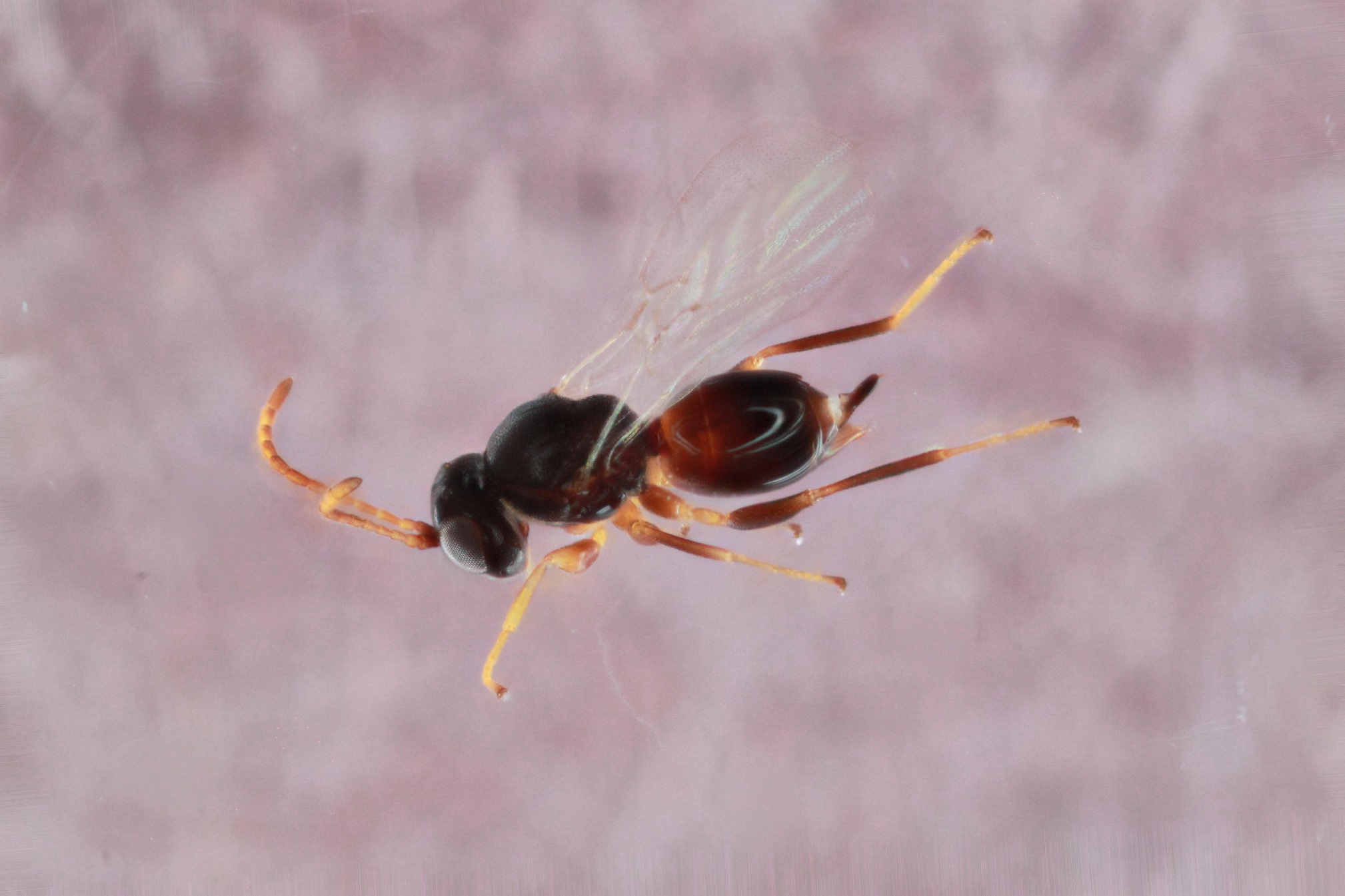

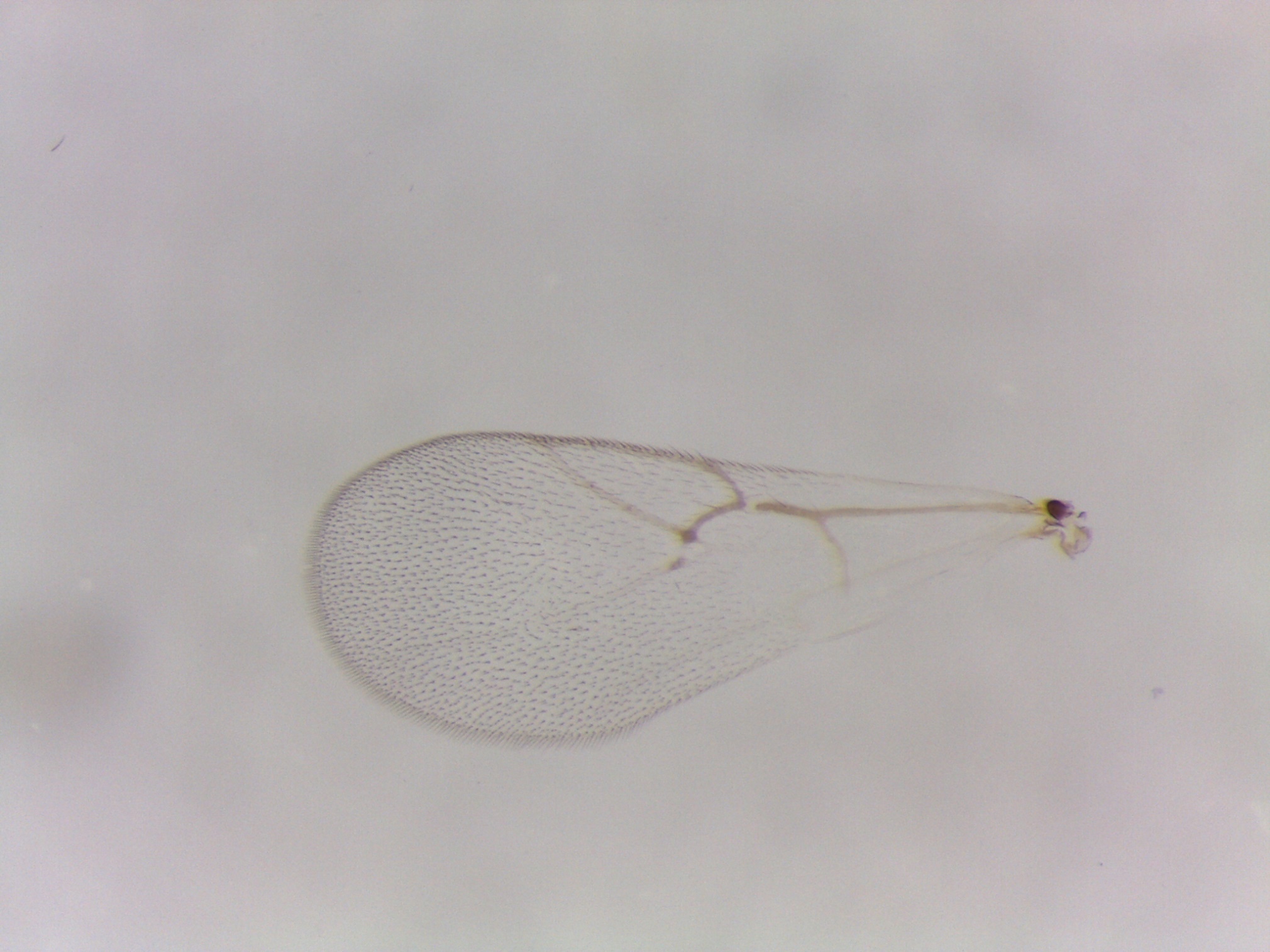

Figure S29. *Ceroptres* sp. 7-8-9 – 843-016-25 – female from *Melikaiella ostensackeni* on *Quercus rubra* in Hannibal, MO.

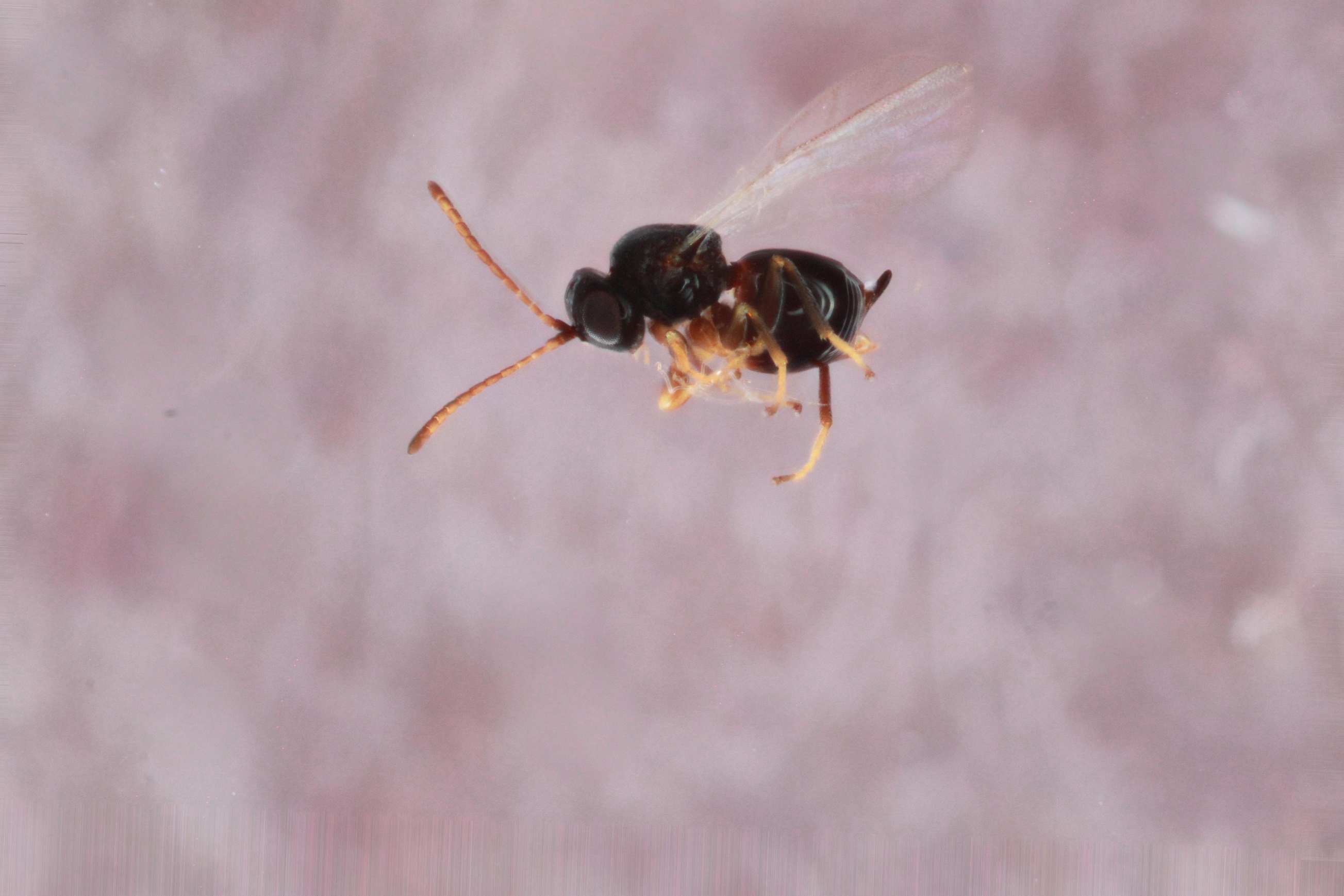

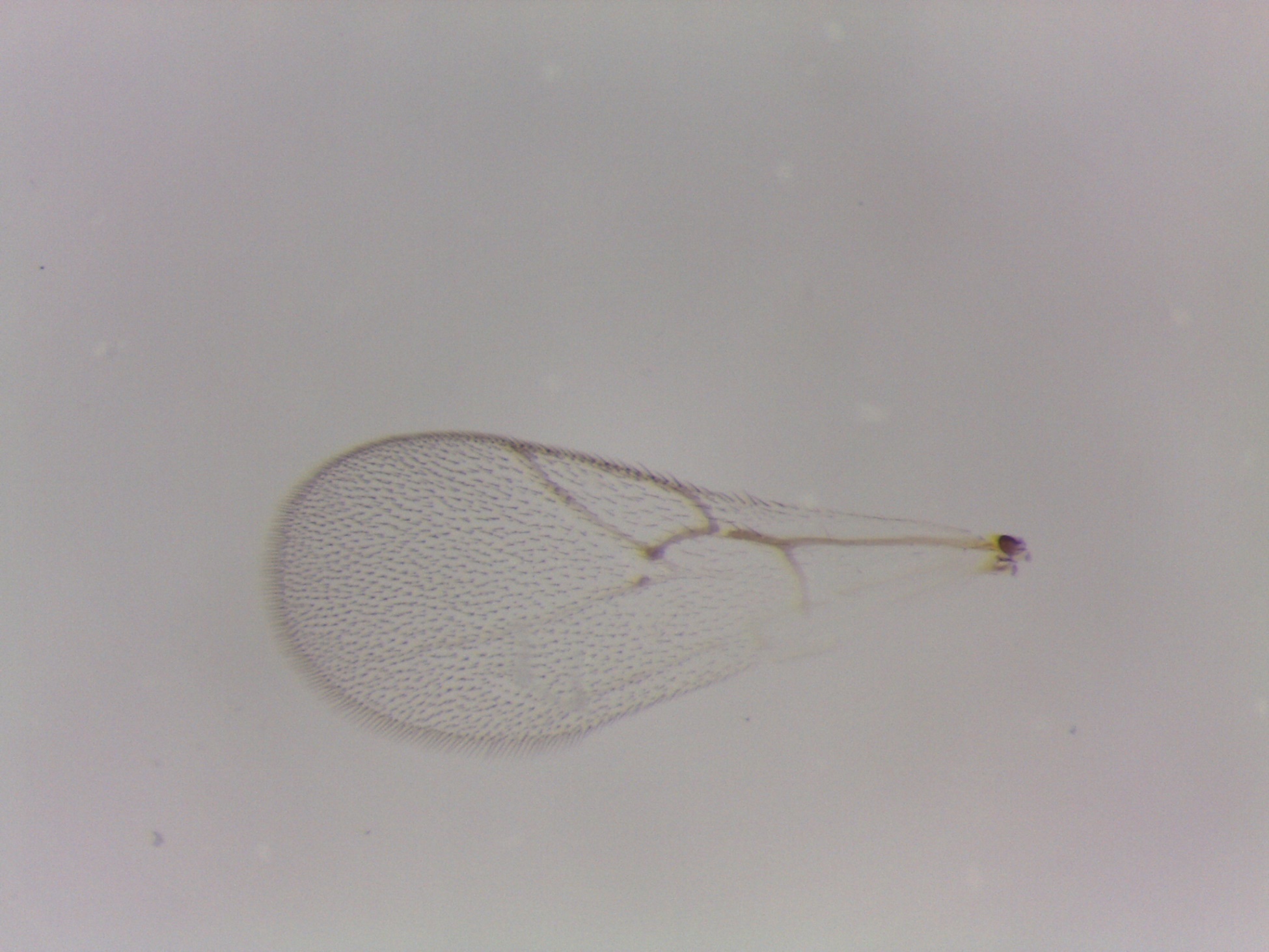

Figure S30. *Ceroptres* sp. 7-8-9 – 401-016-12 – female from *Melikaiella ostensackeni* on *Quercus rubra* in Traverse City, MI.

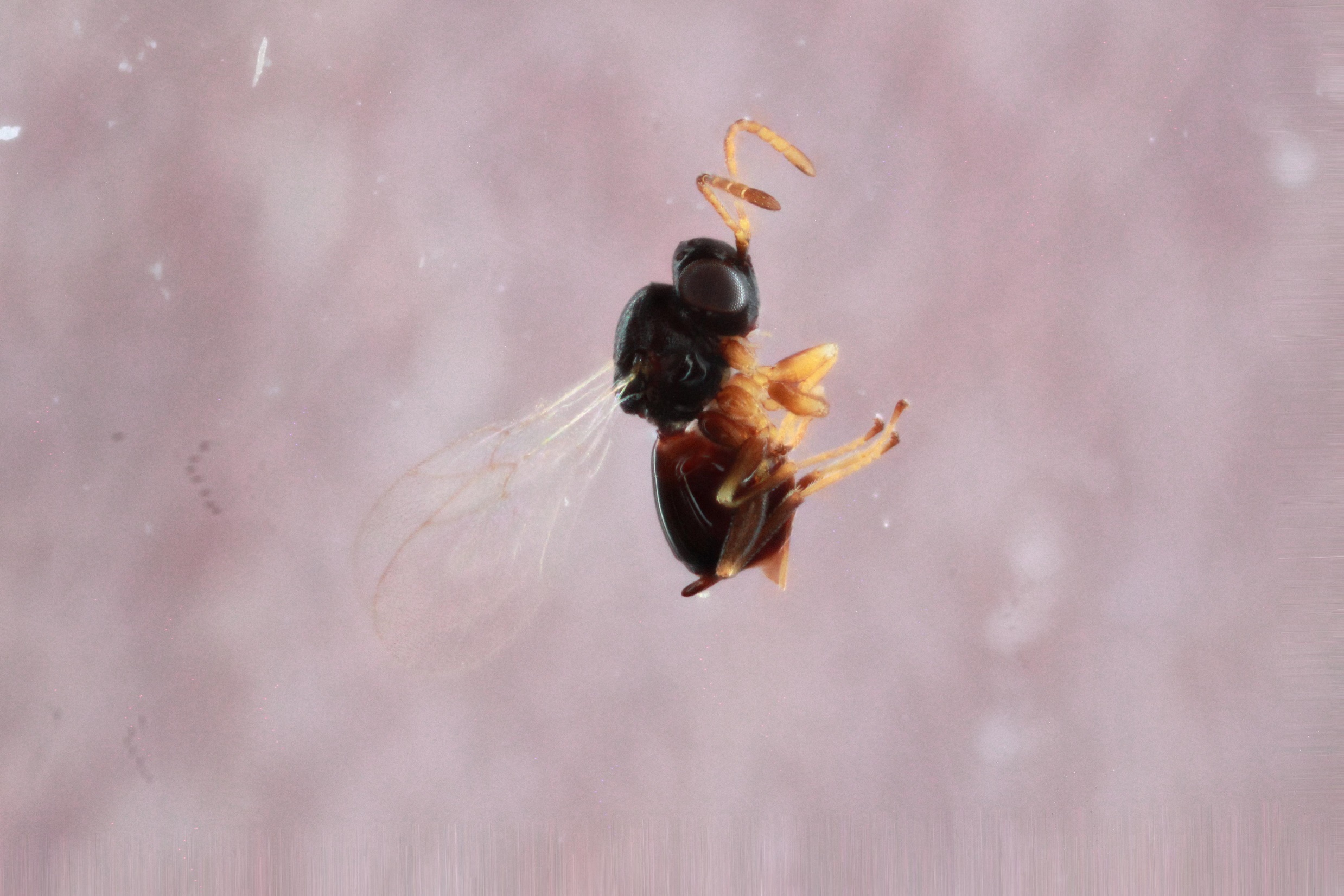

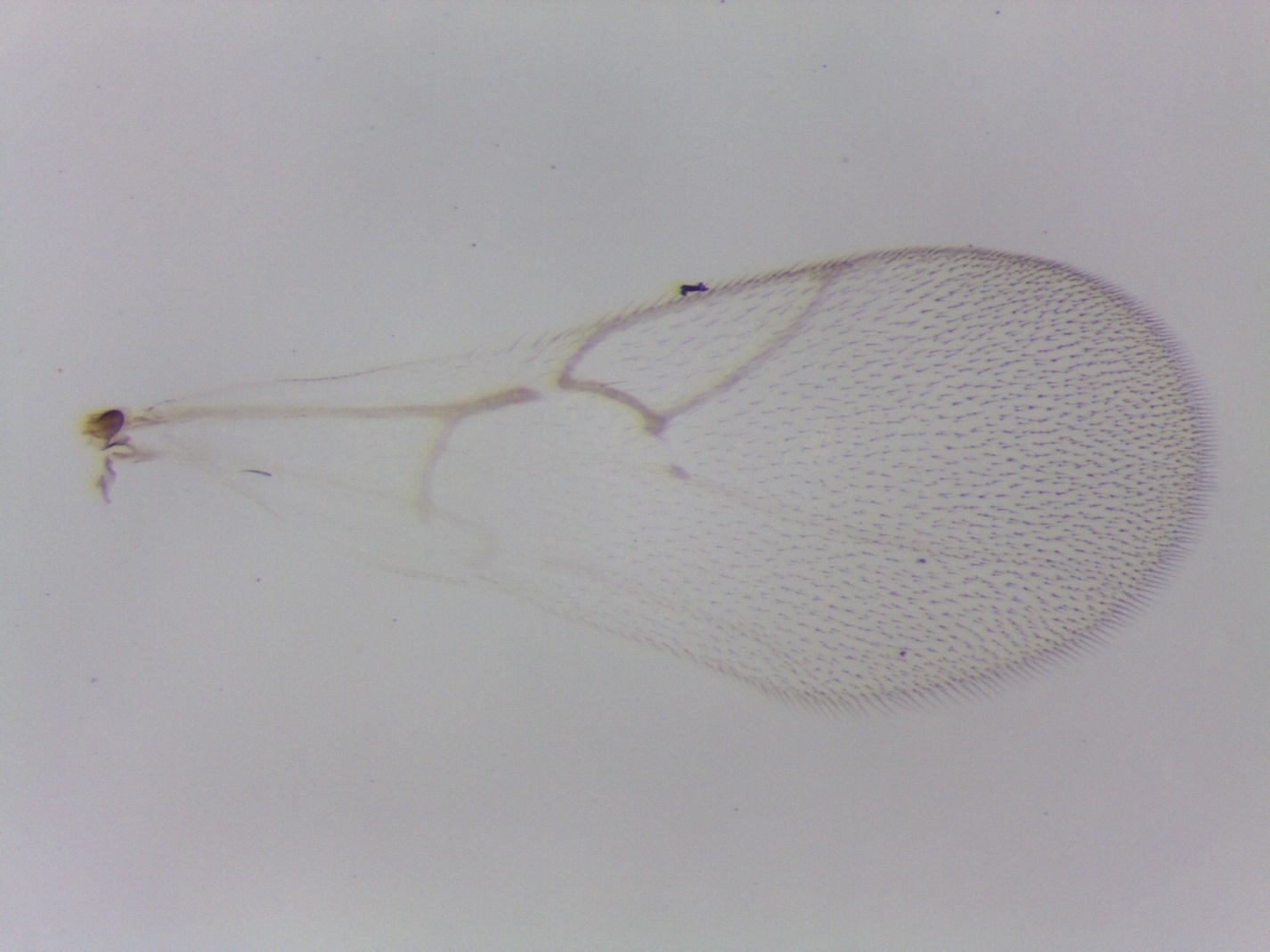

Figure S31. *Ceroptres* sp. 7-8-9 – 856-016-21 – female from *Melikaiella ostensackeni* on *Quercus palustris* in Iowa City, IA.

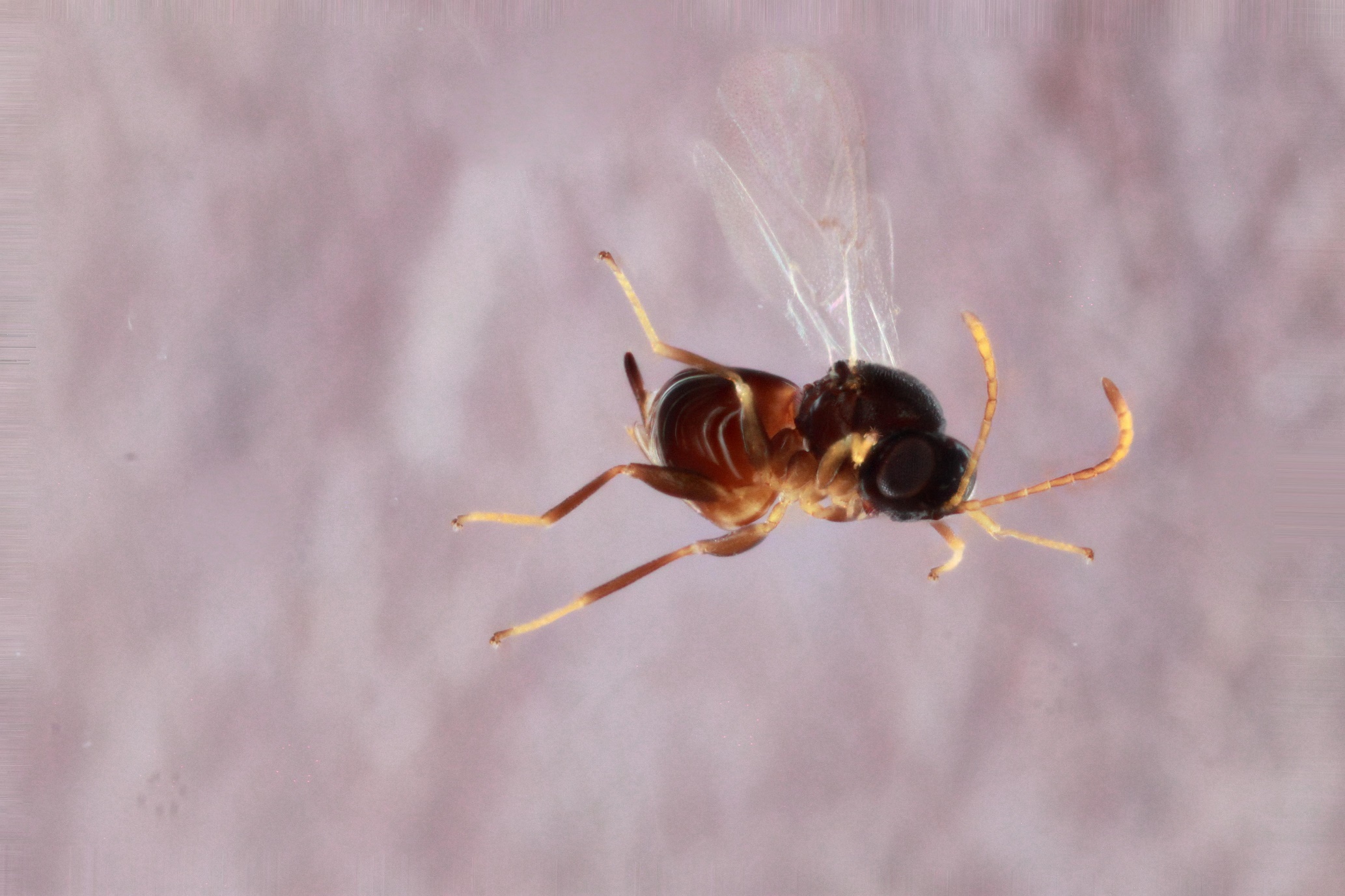

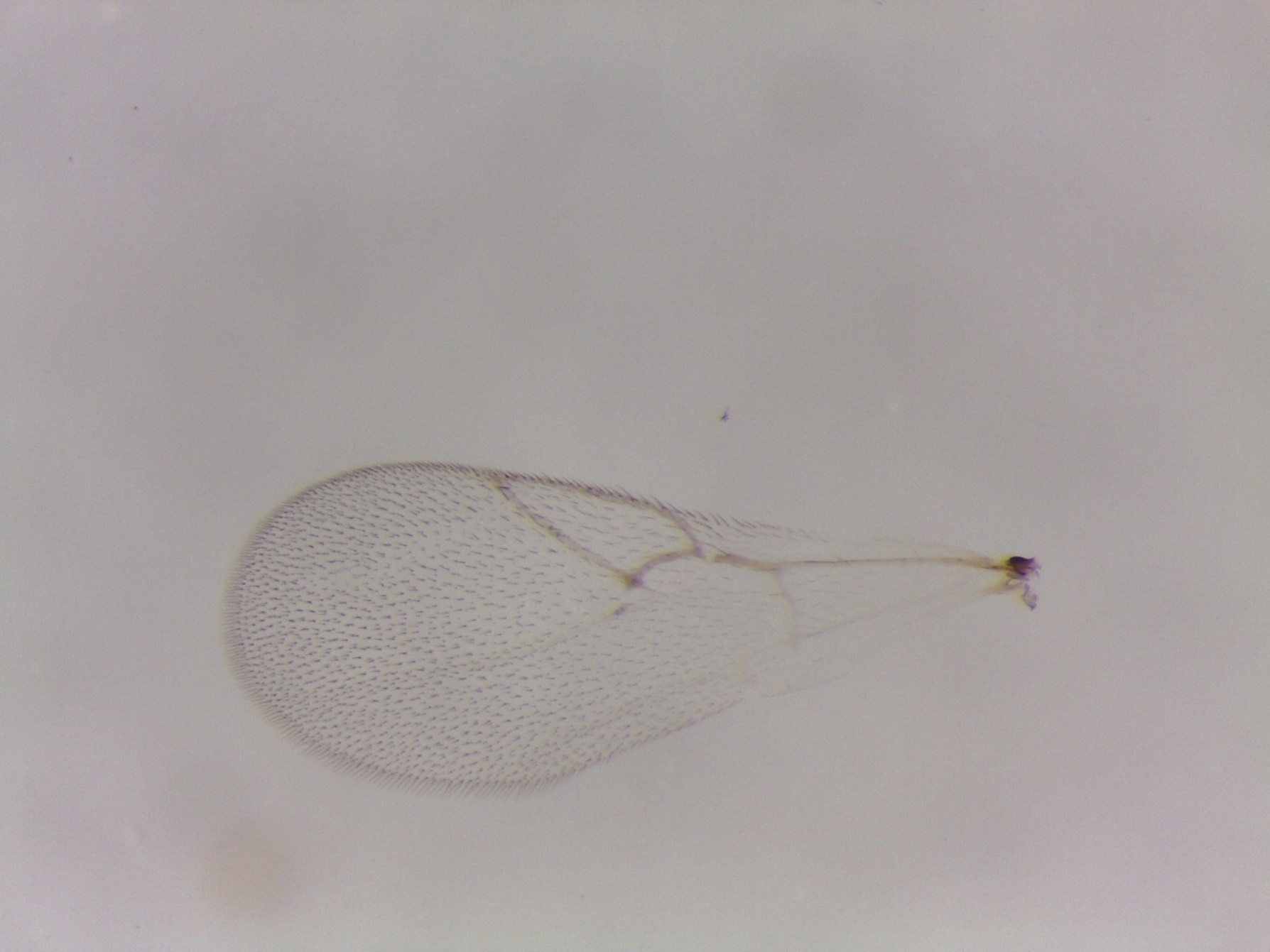

Figure S32. *Ceroptres* sp. 7-8-9 – 933-016-7A – female from *Melikaiella ostensackeni* on *Quercus palustris* in St. Louis, MO (no wing picture).

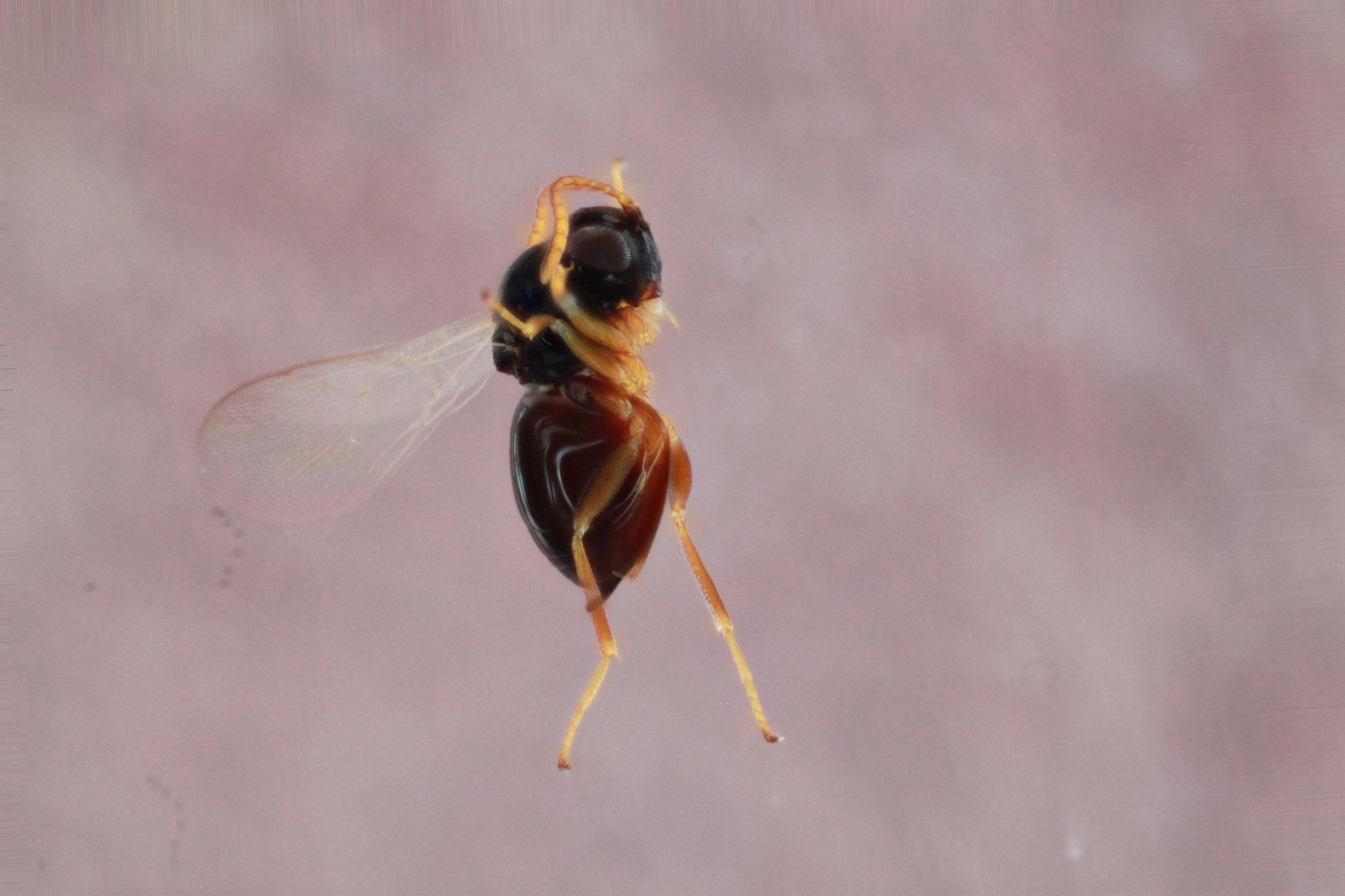

Figure S33. *Ceroptres* sp. 10 – 1416-043-16A – female from *Melikaiella tumifica* on *Quercus rubra* in Iowa City, IA.

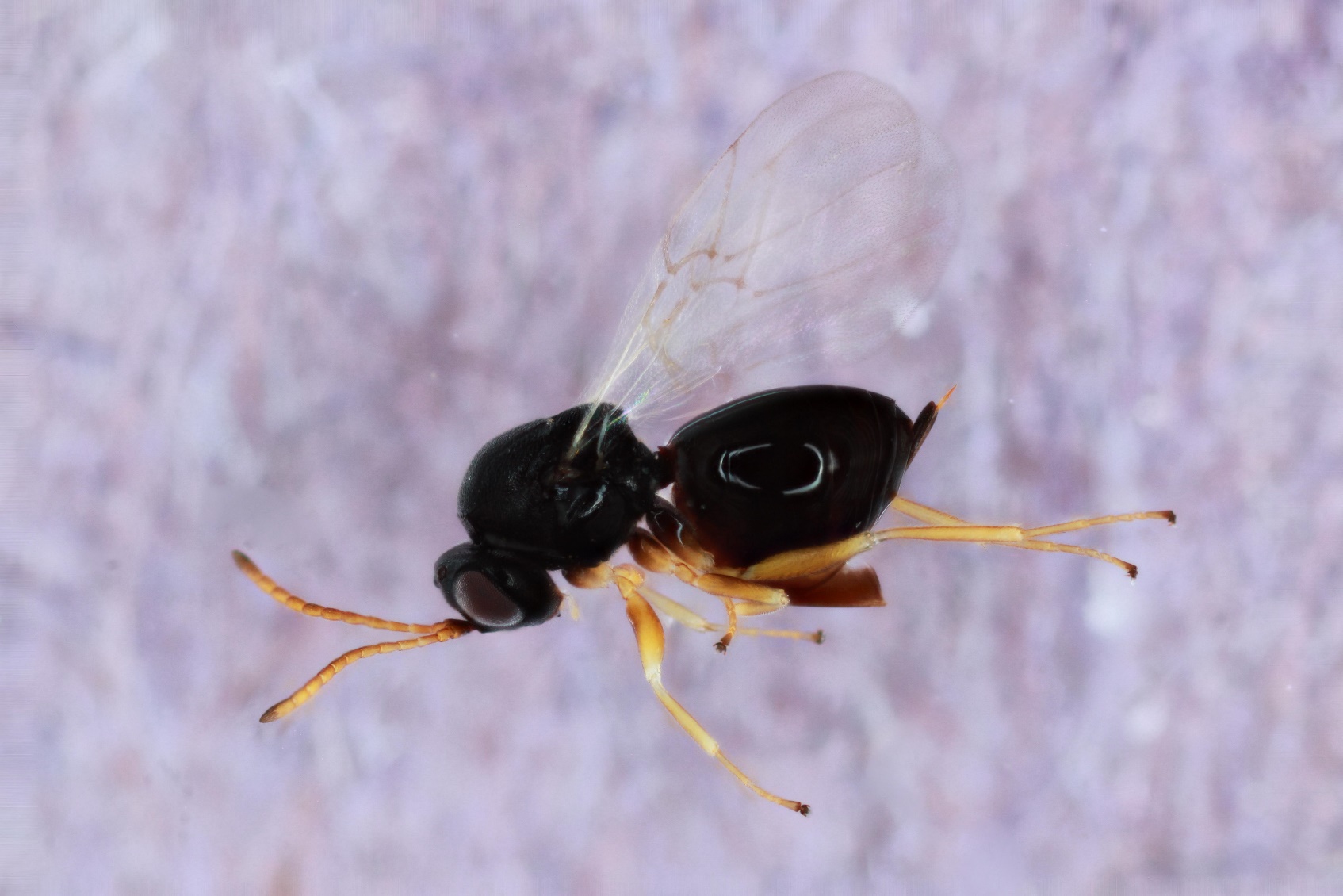

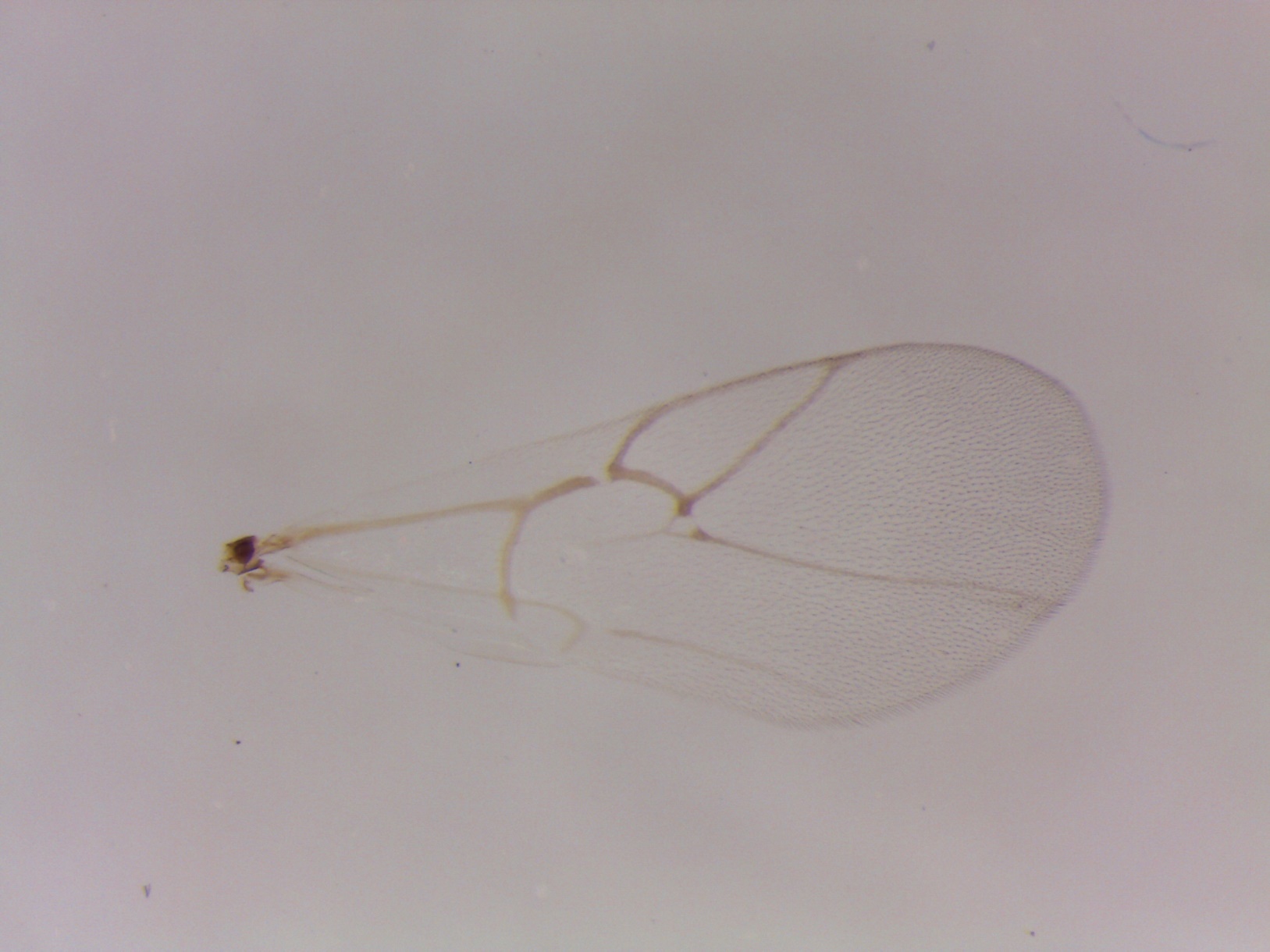

Figure S34. *Ceroptres* sp. 10 – 810-043-9A – female from *Melikaiella tumifica* on *Quercus velutina* in Tiffin, IA.

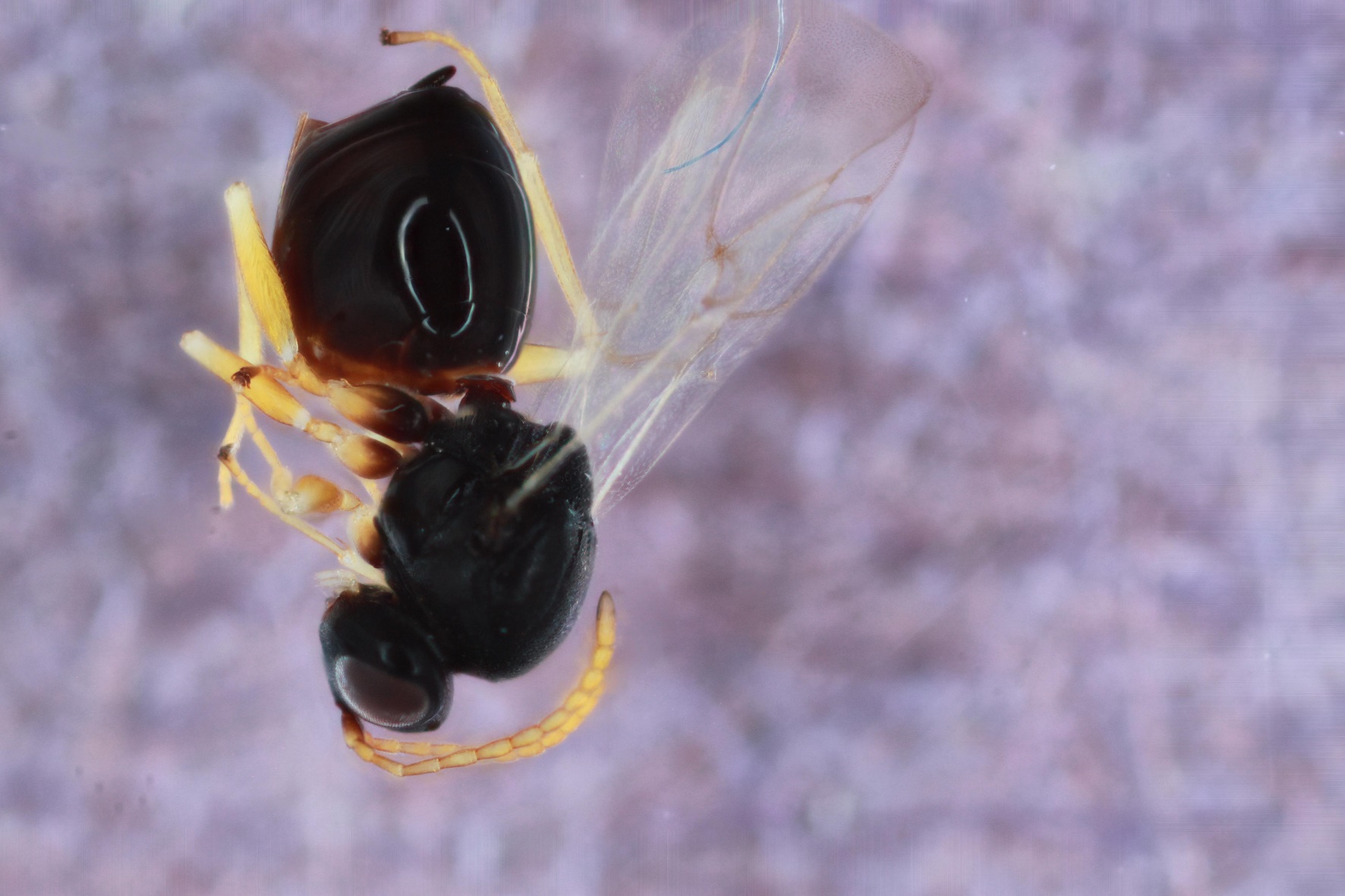

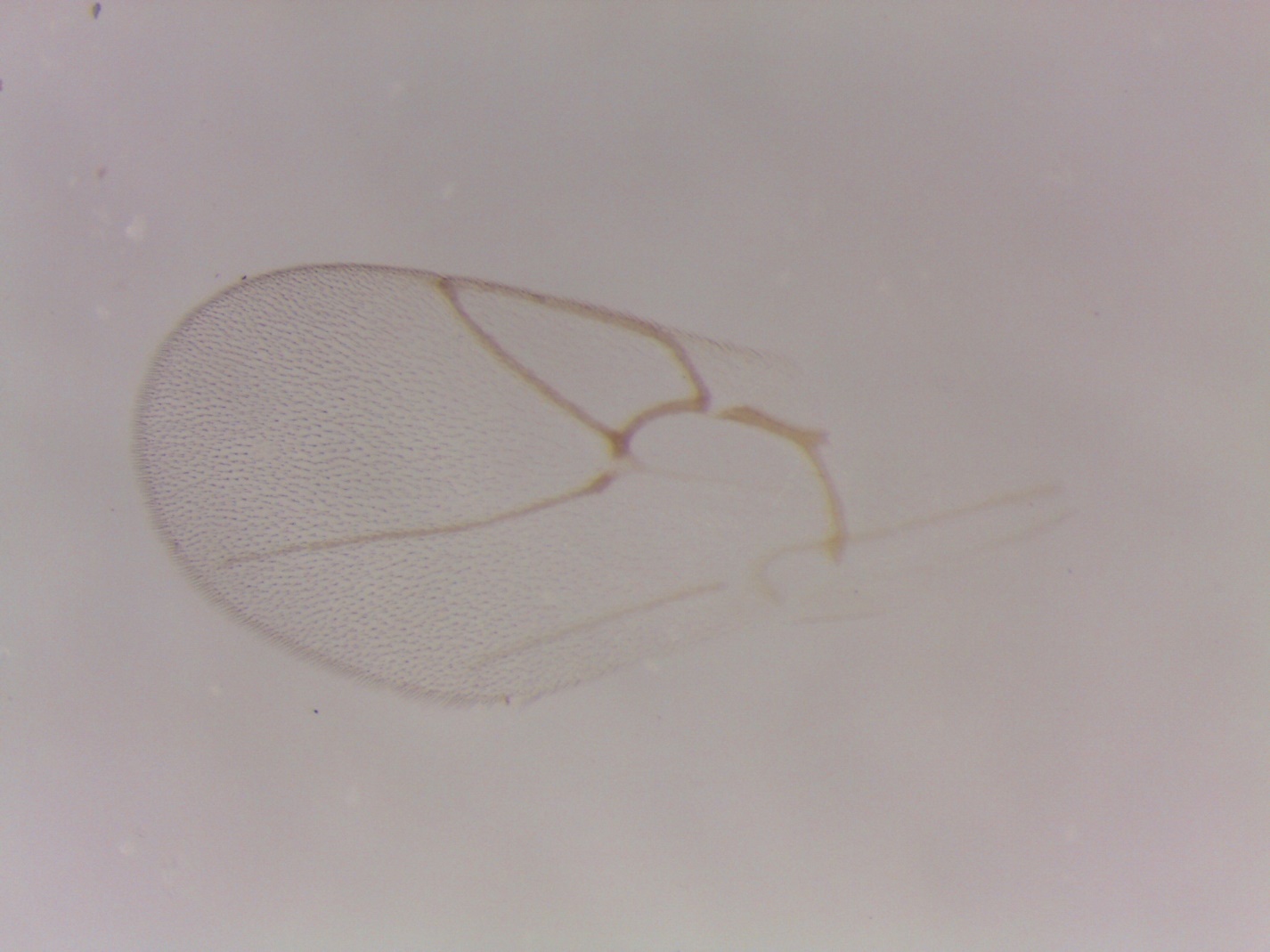

Figure S35. *Ceroptres* sp. 11-12 – 294-049-37A – female from *Callirhytis scitula* on *Quercus imbricaria* in Iowa City, IA.

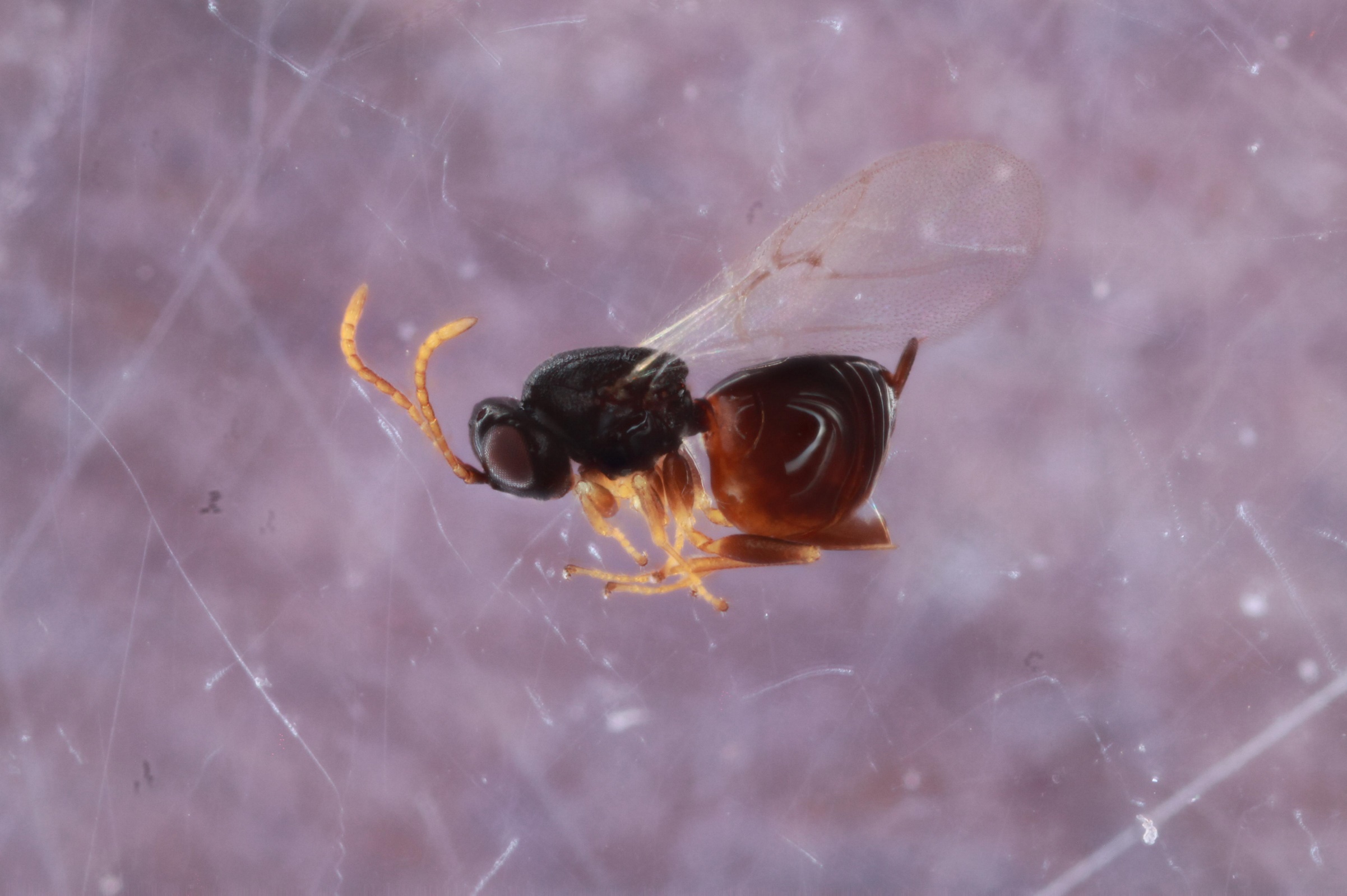

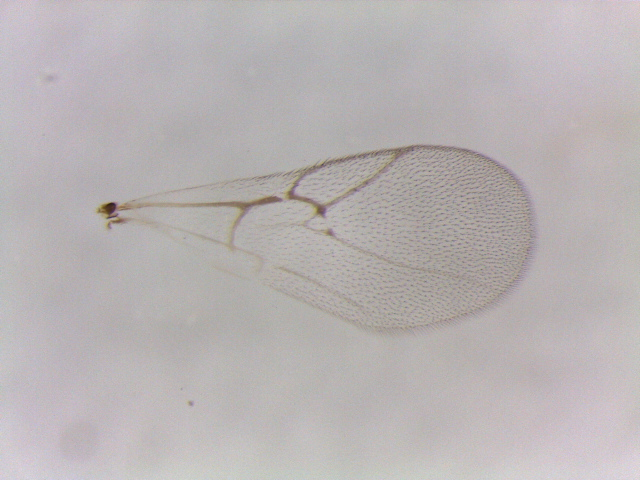

Figure S36. *Ceroptres* sp. 11-12 – 886-049-19A – female from *Callirhytis scitula* on *Quercus palustris* in Iowa City, IA.

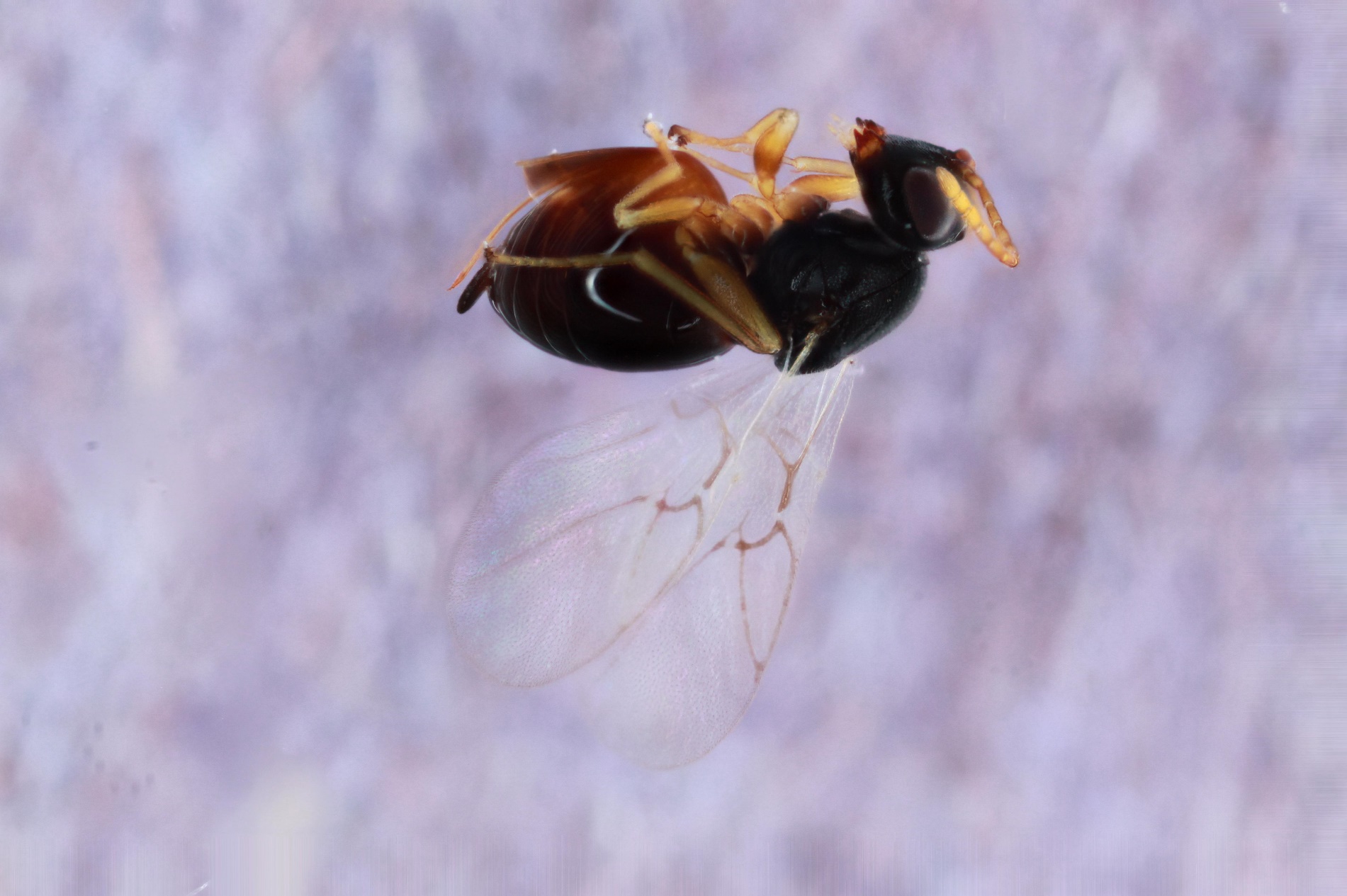

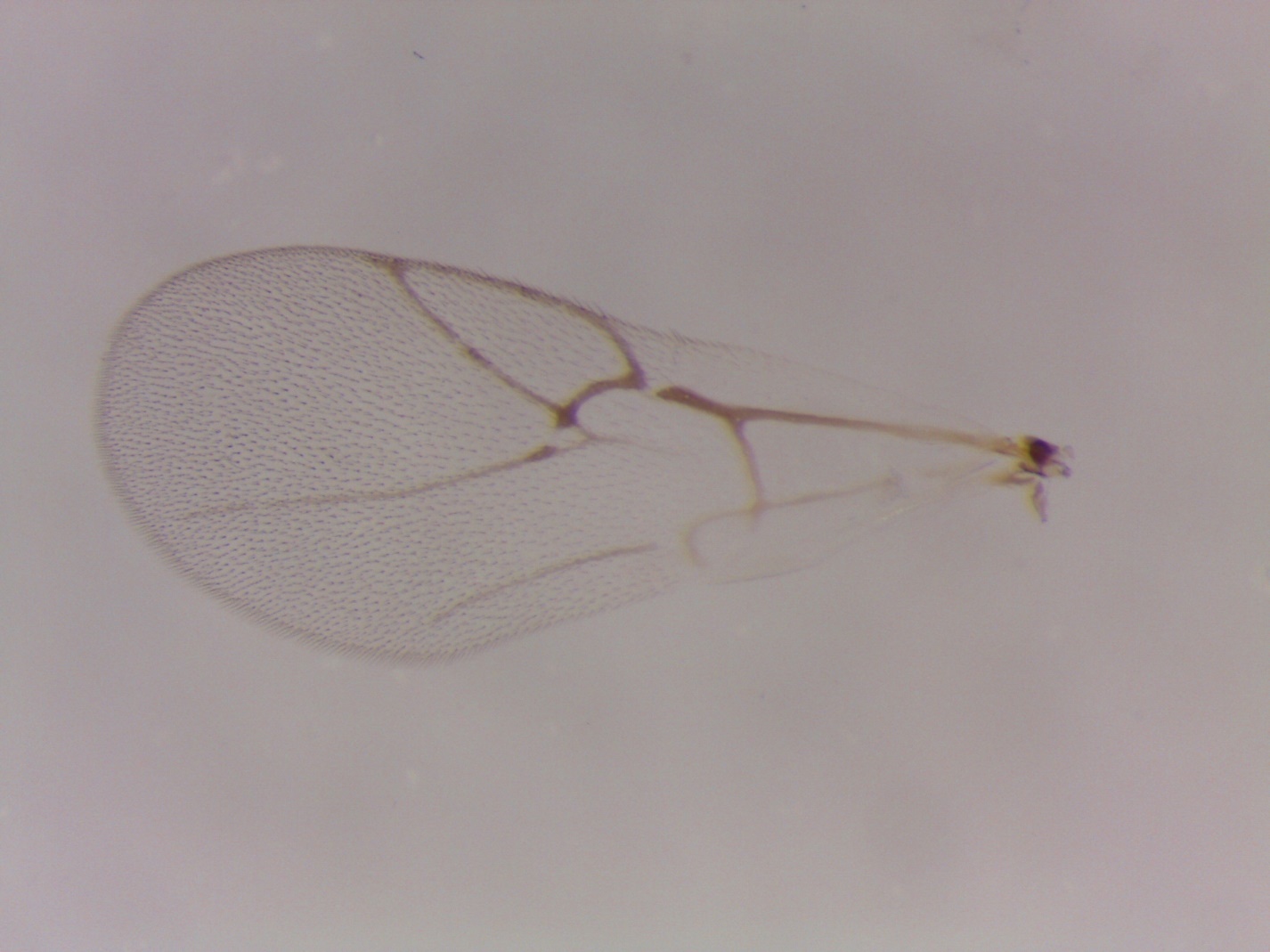

Figure S37. *Ceroptres* sp. 11-12 –528-049-4 – female from *Callirhytis scitula* on *Quercus velutina* in St. Peters, MO.

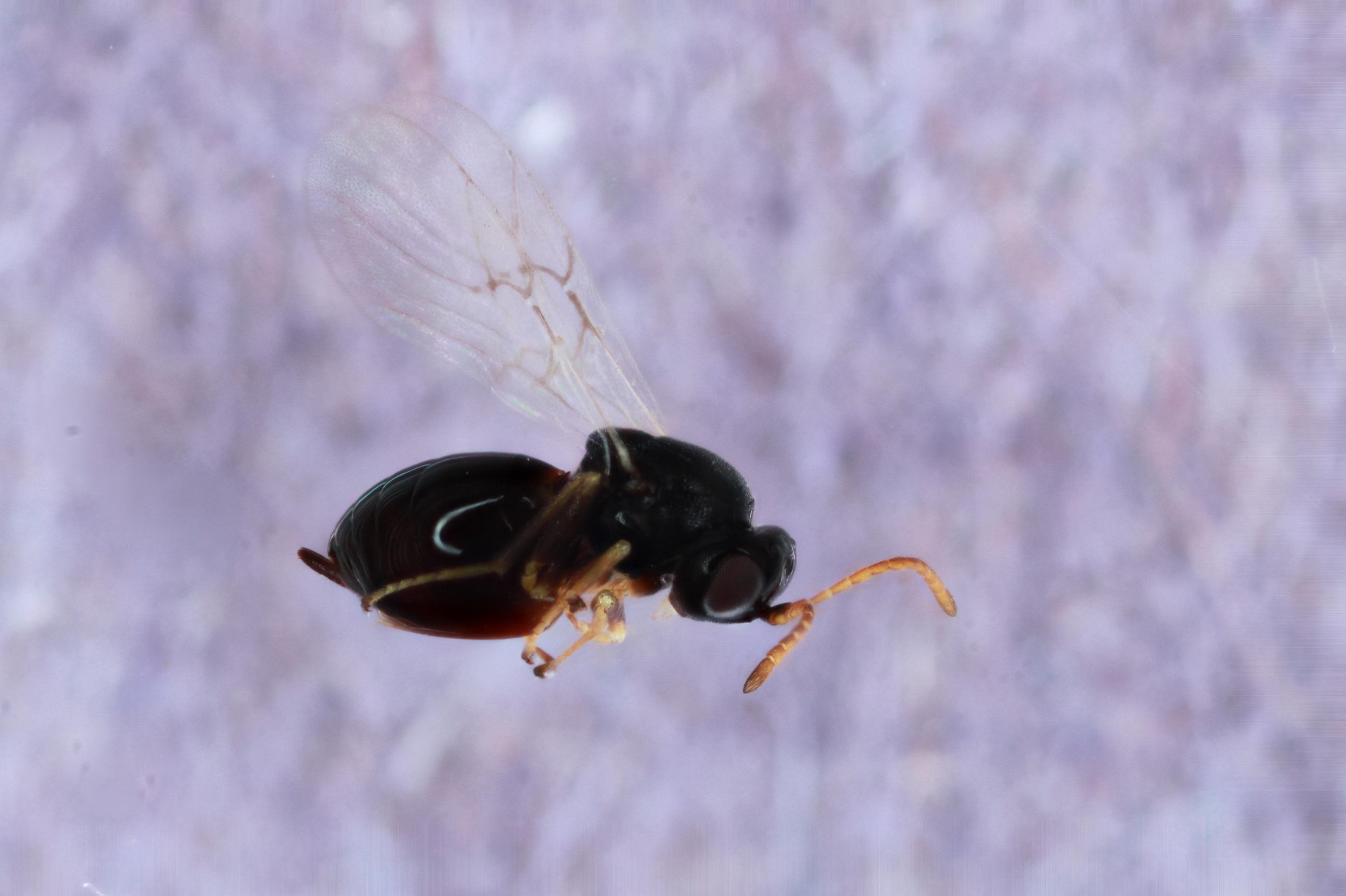

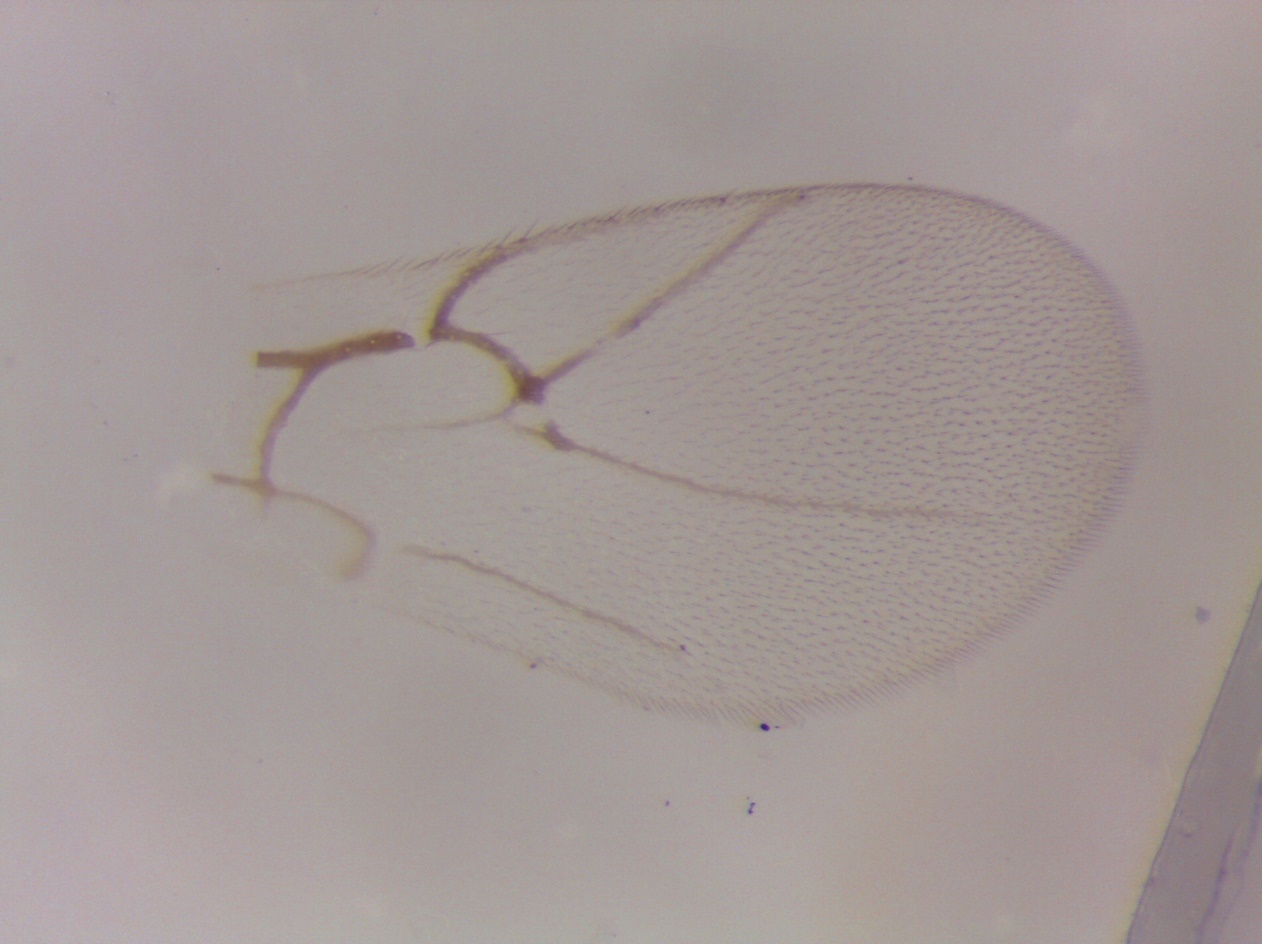

Figure S38. *Ceroptres* sp. 13 – 866-035-22 – female from *Loxaulus quercusmammula* on *Quercus alba* in Tiffin, IA.

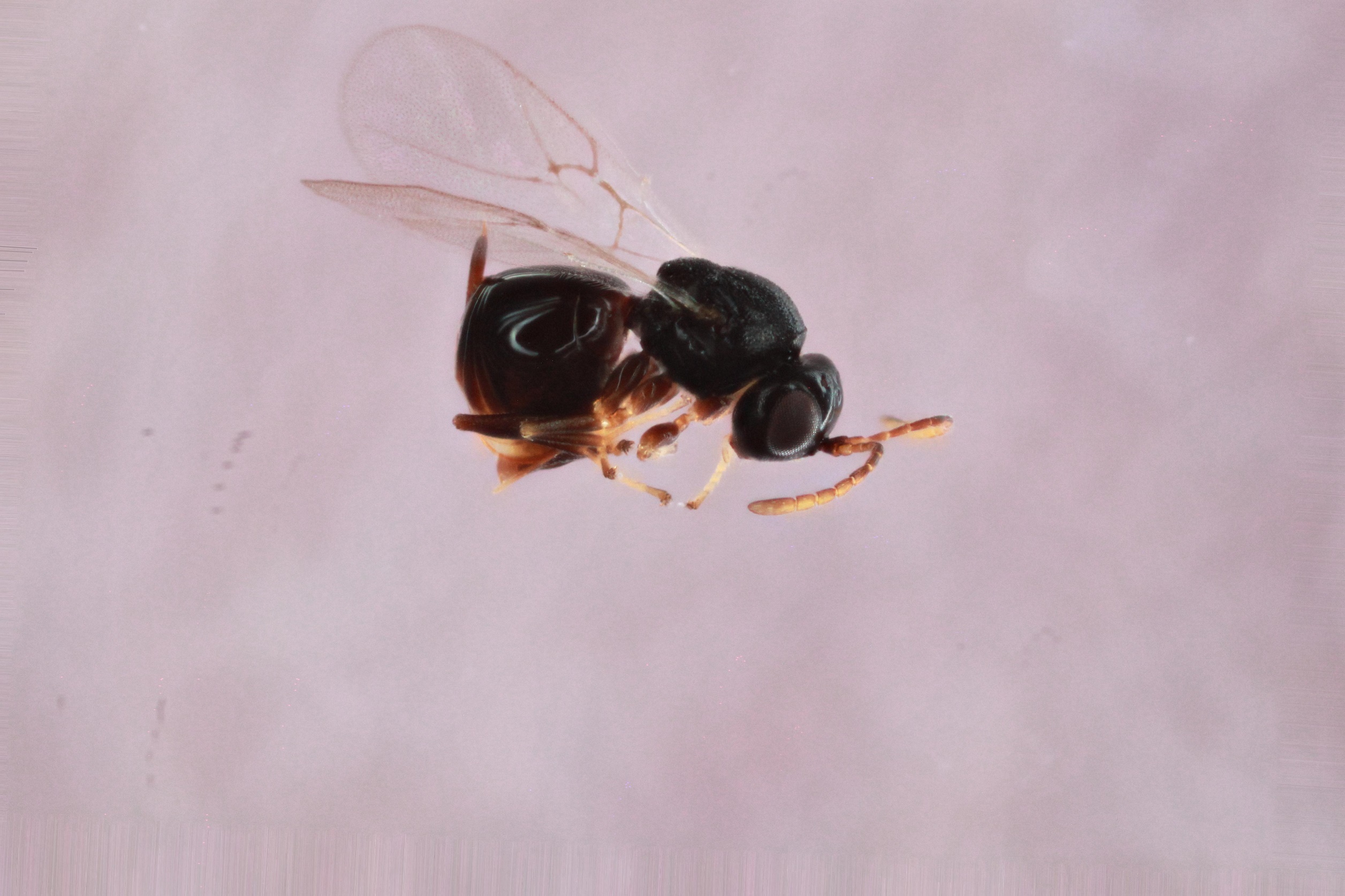

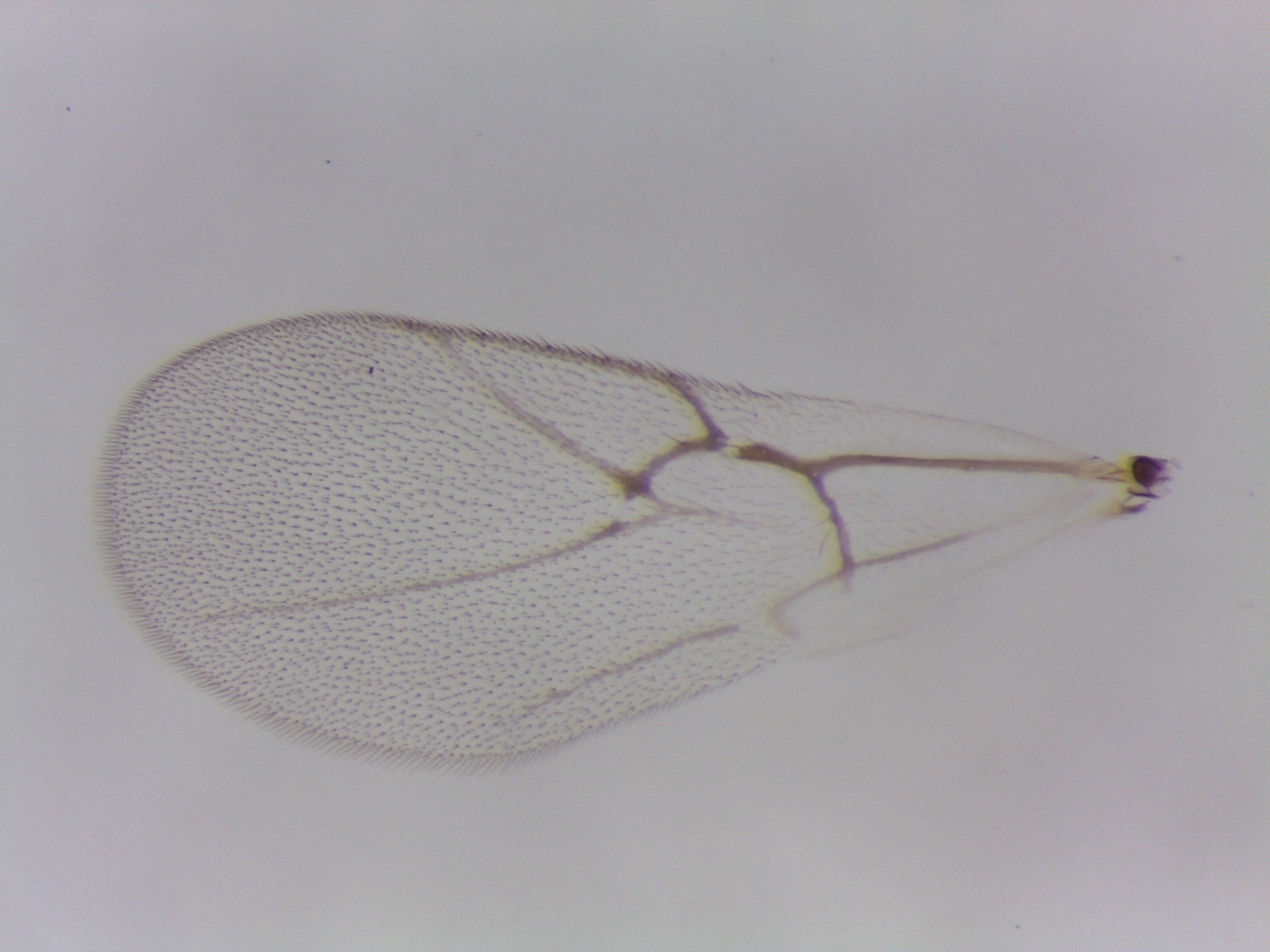

Figure S39. *Ceroptres* sp. 14 – 671-083-2 – female from an unknown stem gall on *Quercus macrocarpa* in Urbana, IL.

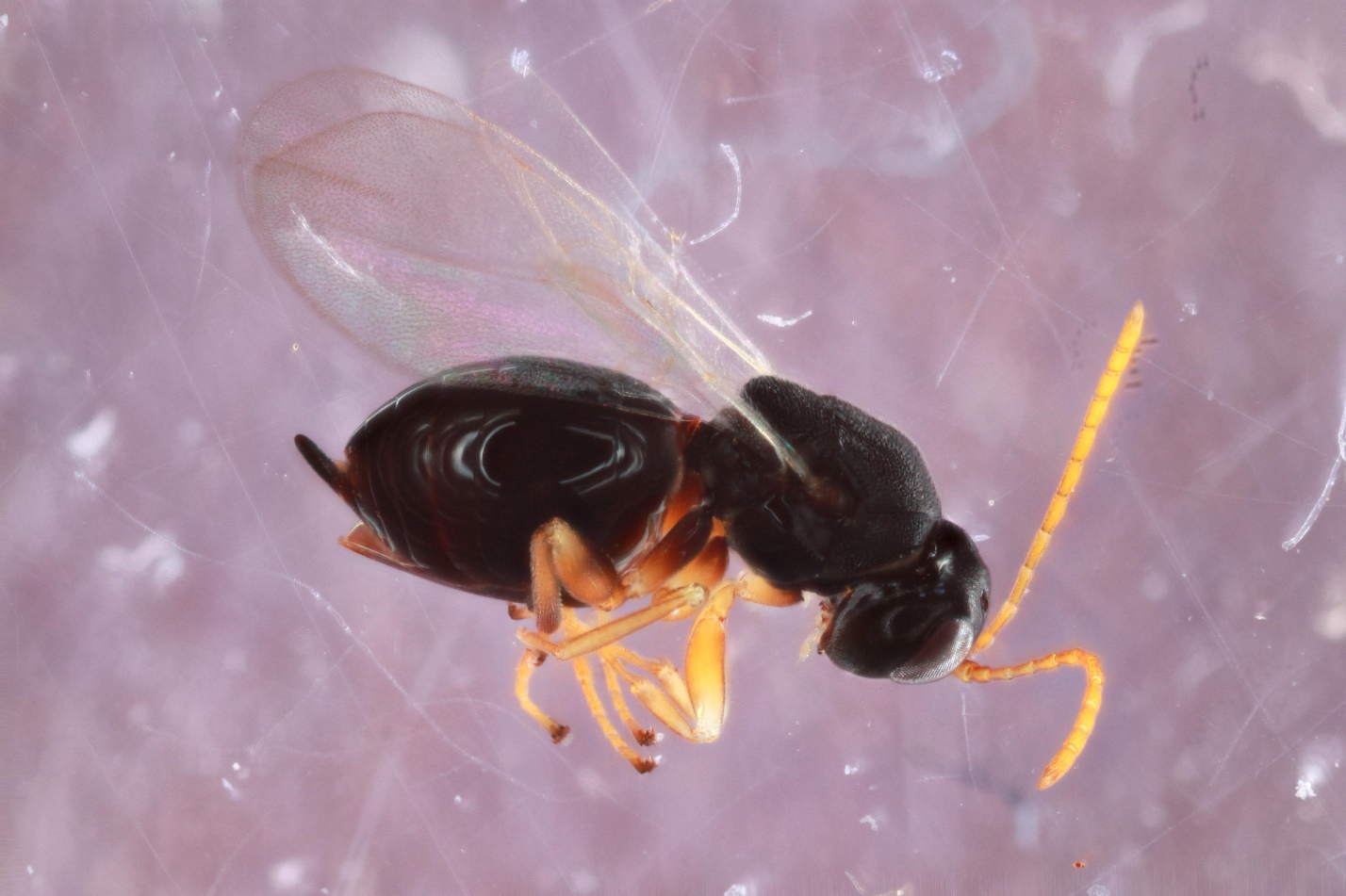

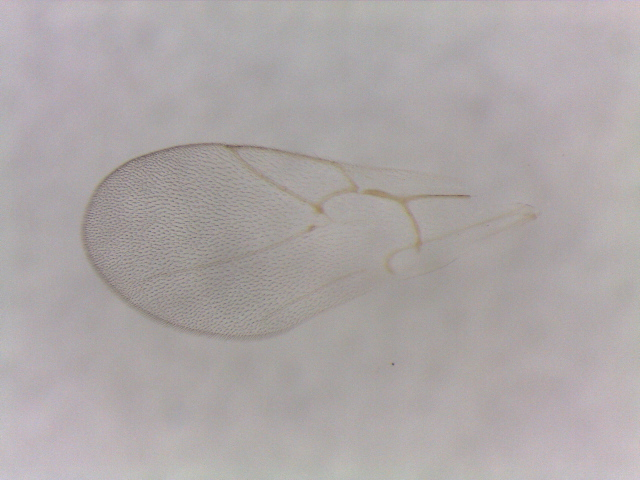

Figure S40. *Ceroptres* sp. 15-16 – 865-051-11 – male from *Callirhytis quercusfutilis* on *Quercus alba* in Tiffin, IA.

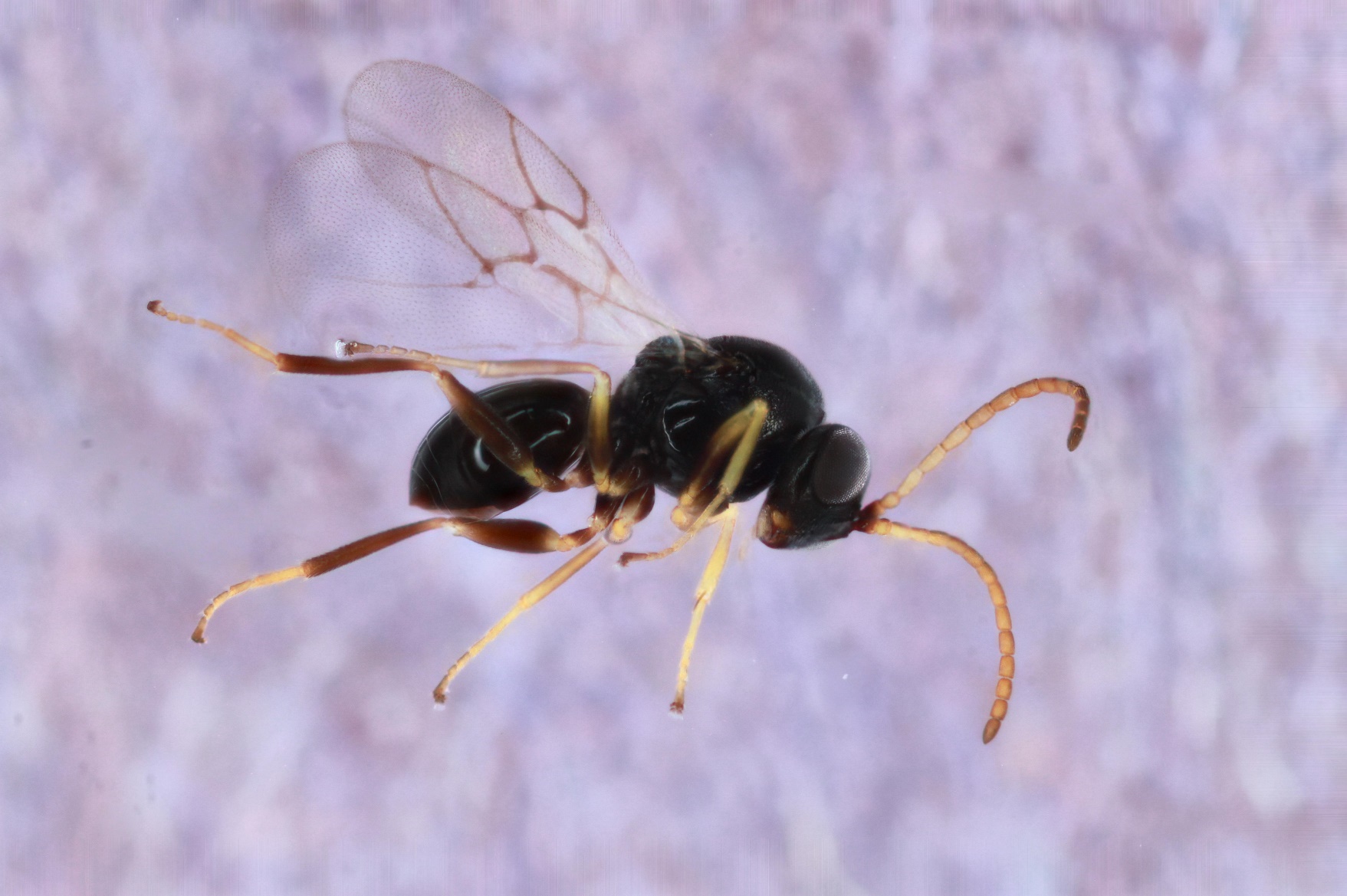

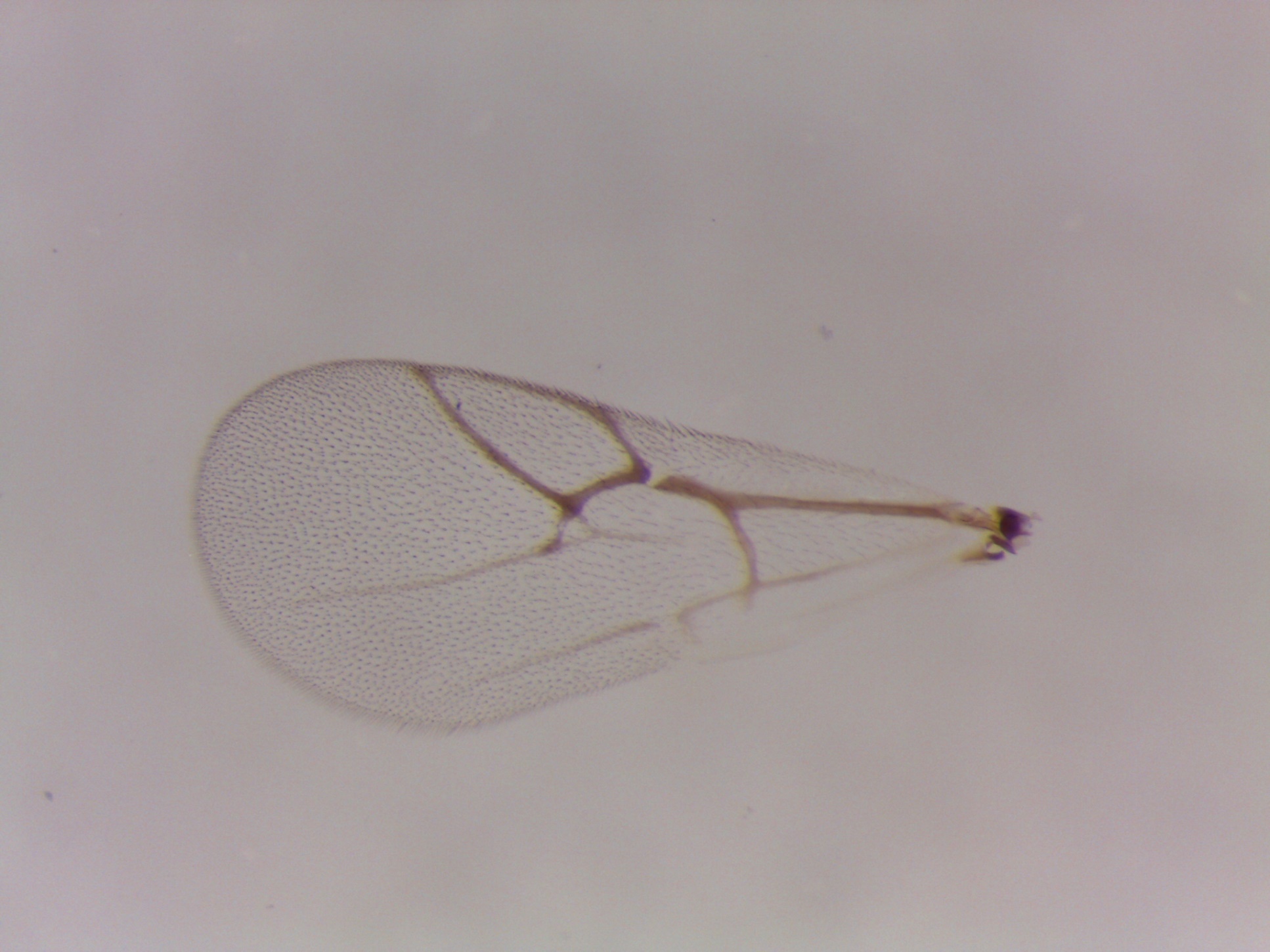

Figure S41. *Ceroptres* sp. 15-16 – 369-053-26 – female from *Neuroterus quercusbatatus* on *Quercus bicolor* in Iowa City, IA (no wing picture).

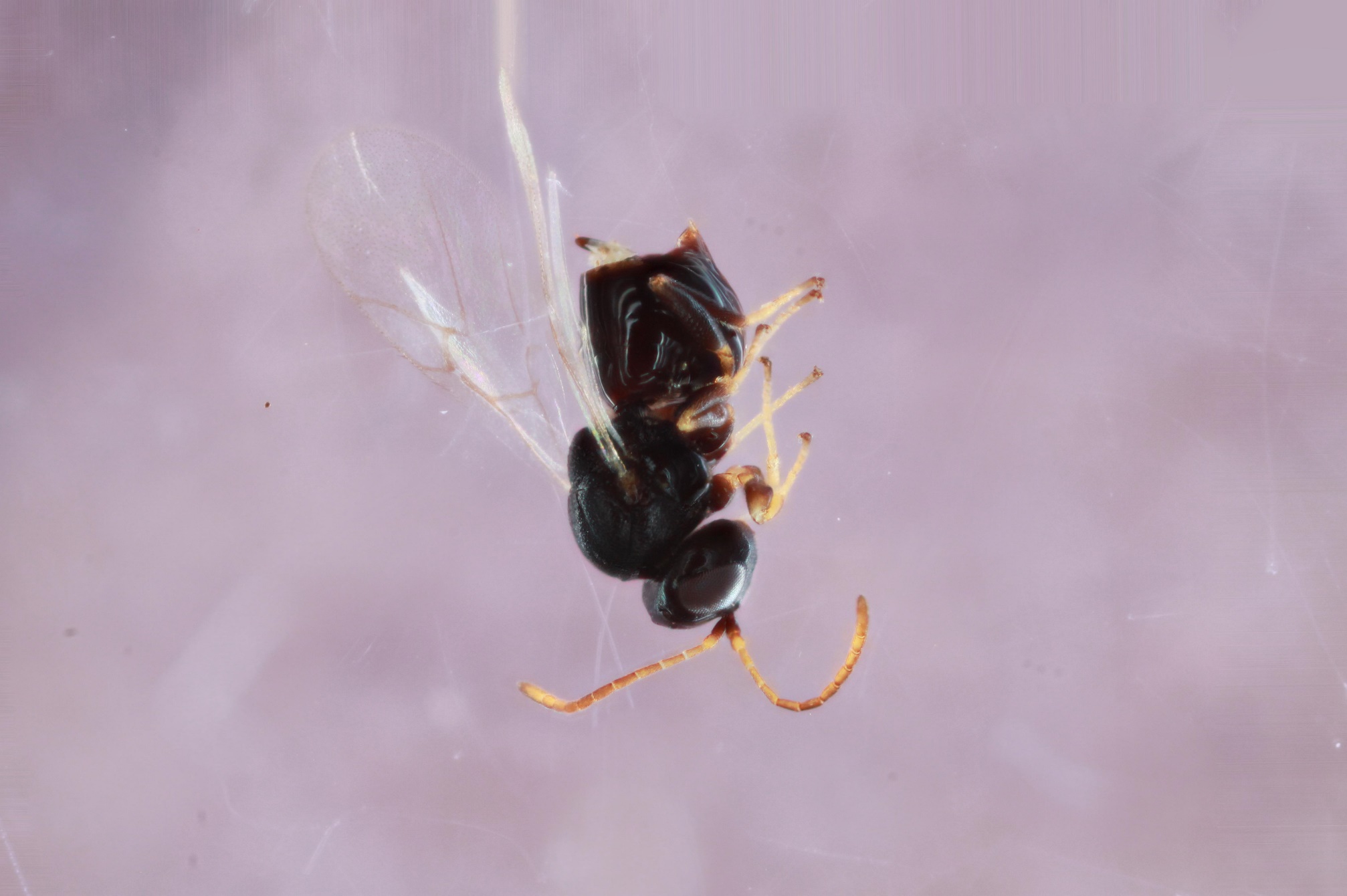

Figure S42. *Ceroptres* sp. 15-16 – 2-22-2 – female from an unidentified integral leaf gall on *Quercus macrocarpa* in Spirit Lake, IA (no body picture).

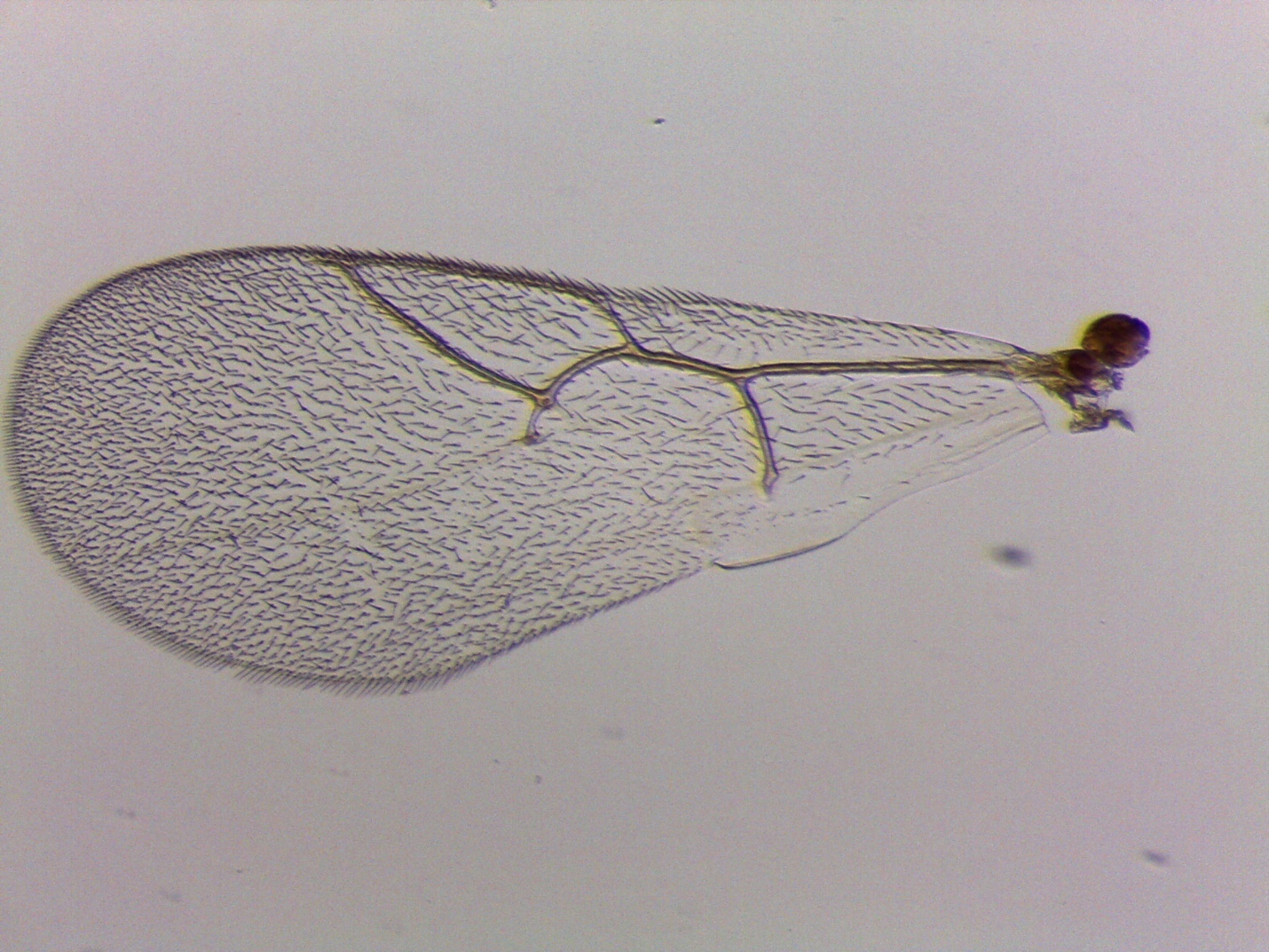

Figure S43. *Ceroptres* sp. 17 – 1239-118-7 – female from *Neuroterus quercusirregularis* on *Quercus stellata* in Austin, TX.

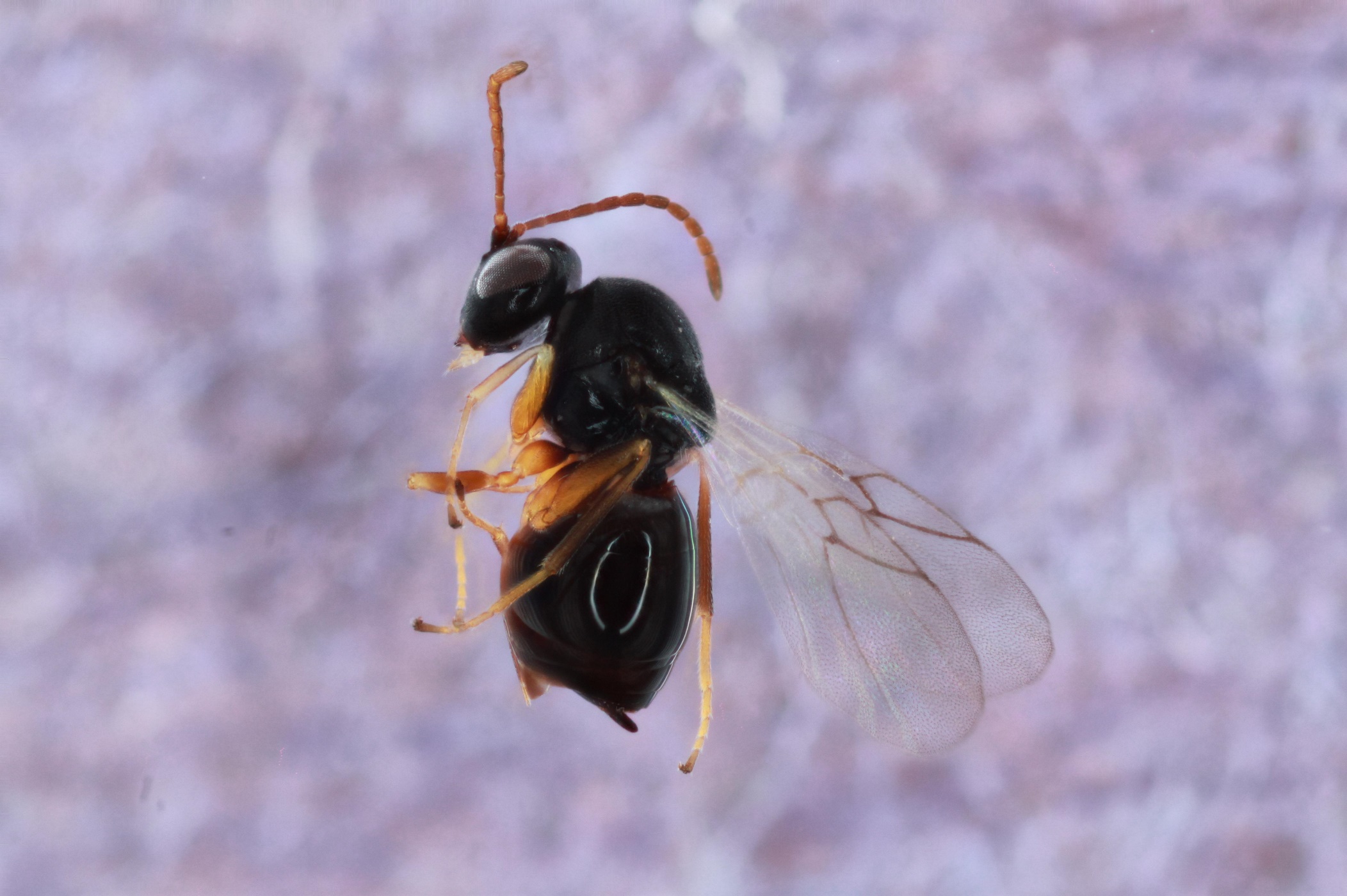

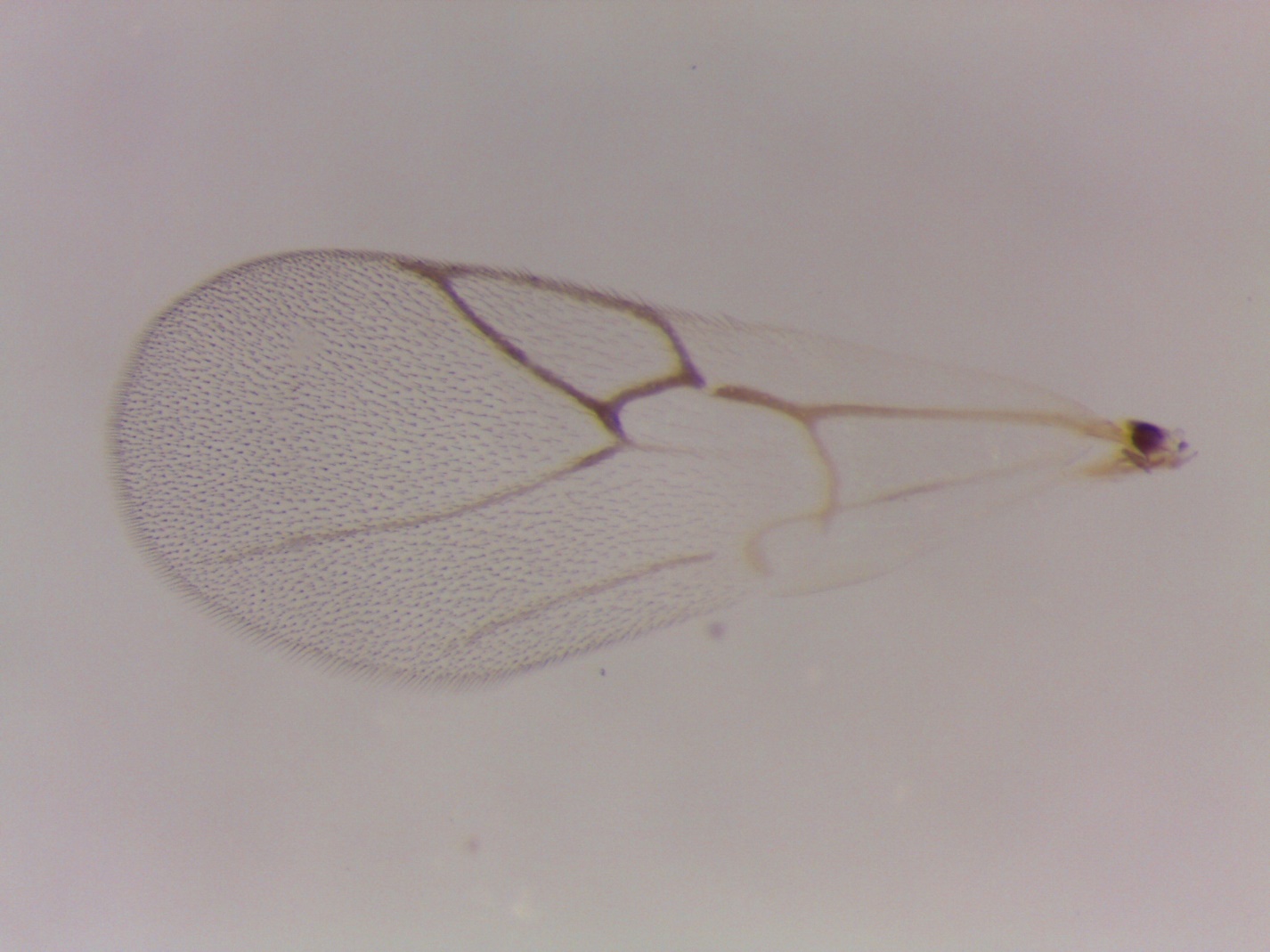

Figure S44. *Ceroptres* sp. 18-19 – 1184-051-5 – female from *Callirhytis quercusfutilis* on *Quercus alba* in White Oak, PA.

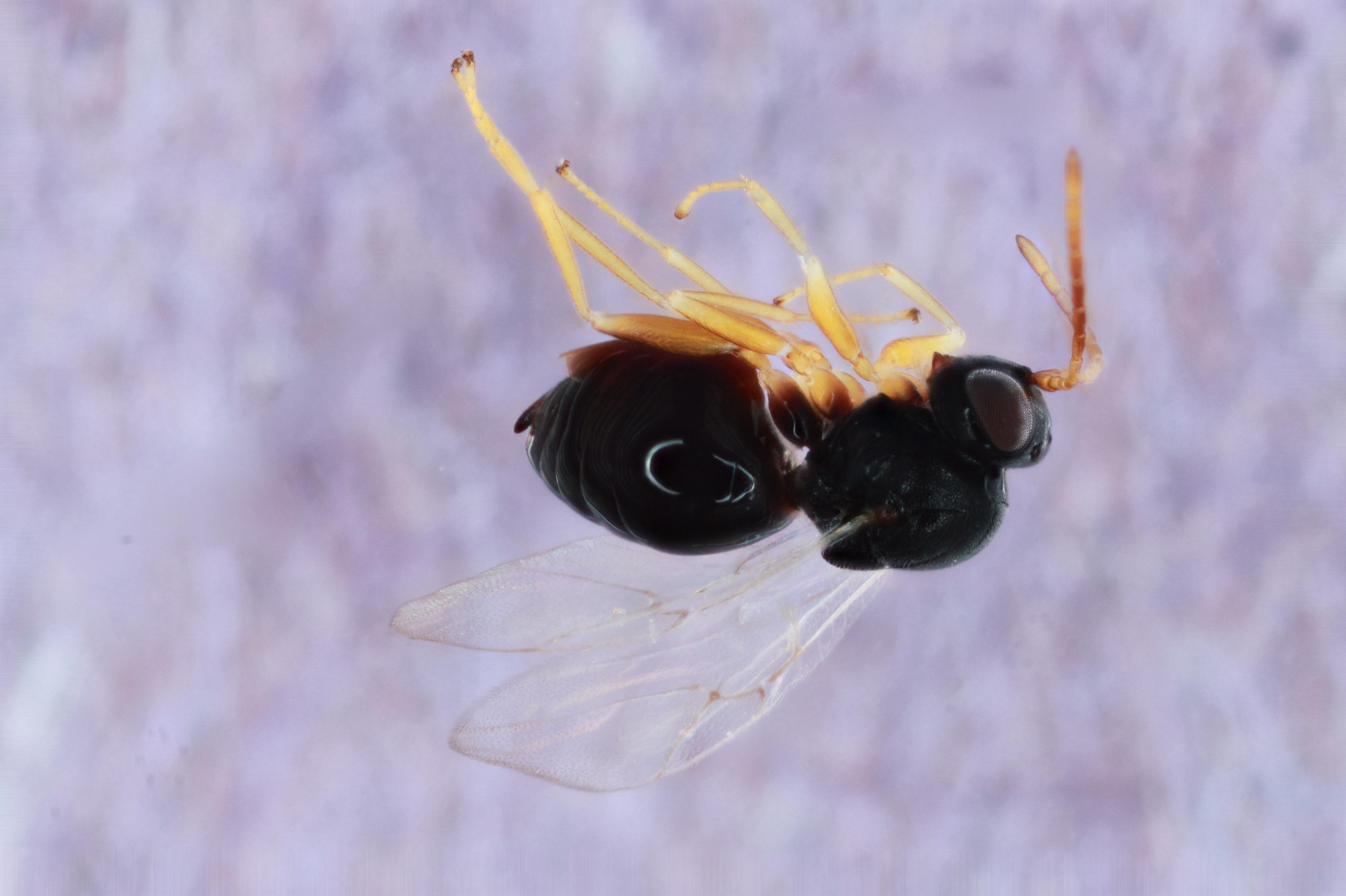

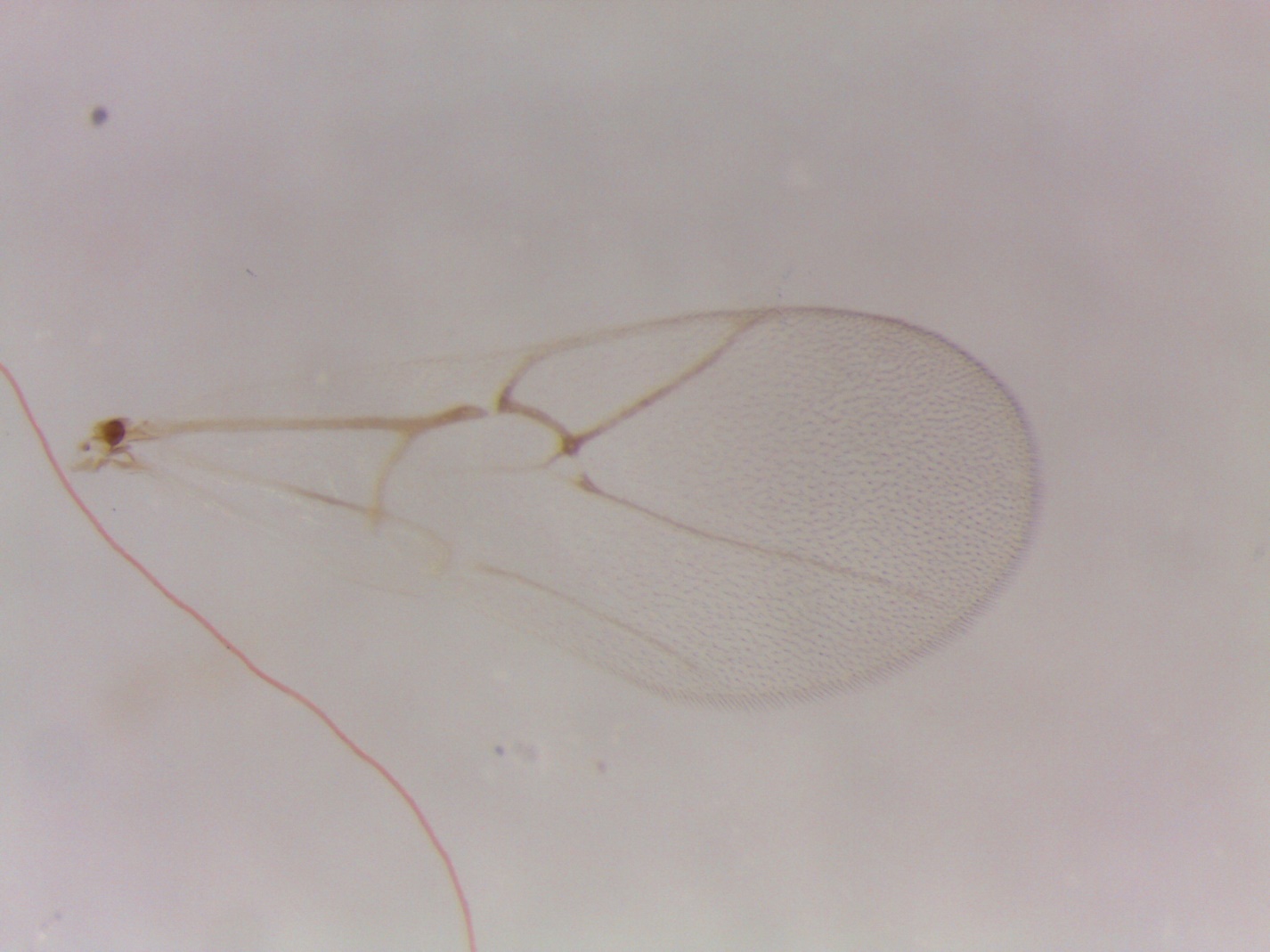

Figure S45. *Ceroptres* sp. 18-19 – 963-051-15 – female from *Callirhytis quercusfutilis* on *Quercus macrocarpa* in Iowa City, IA.

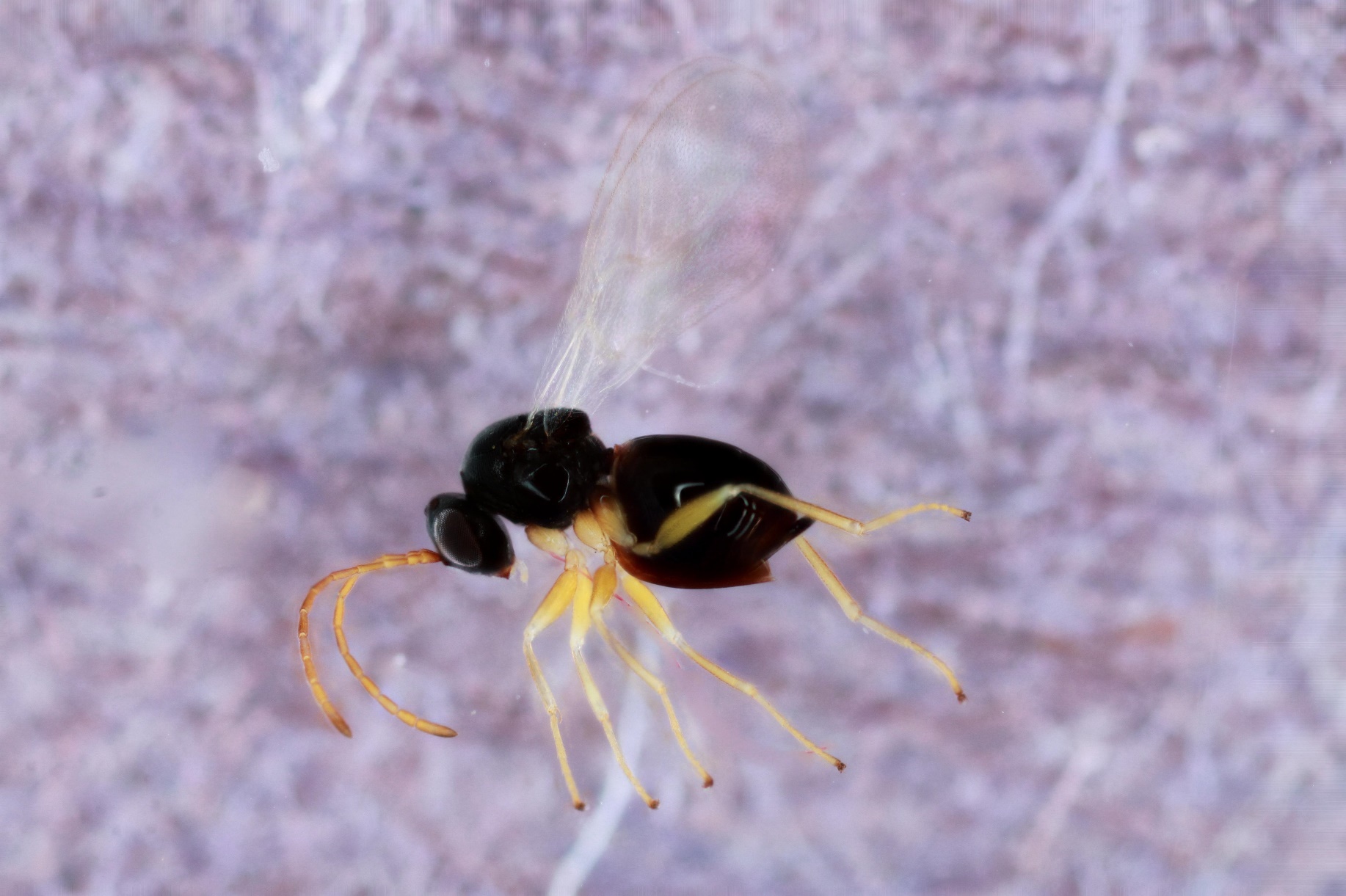

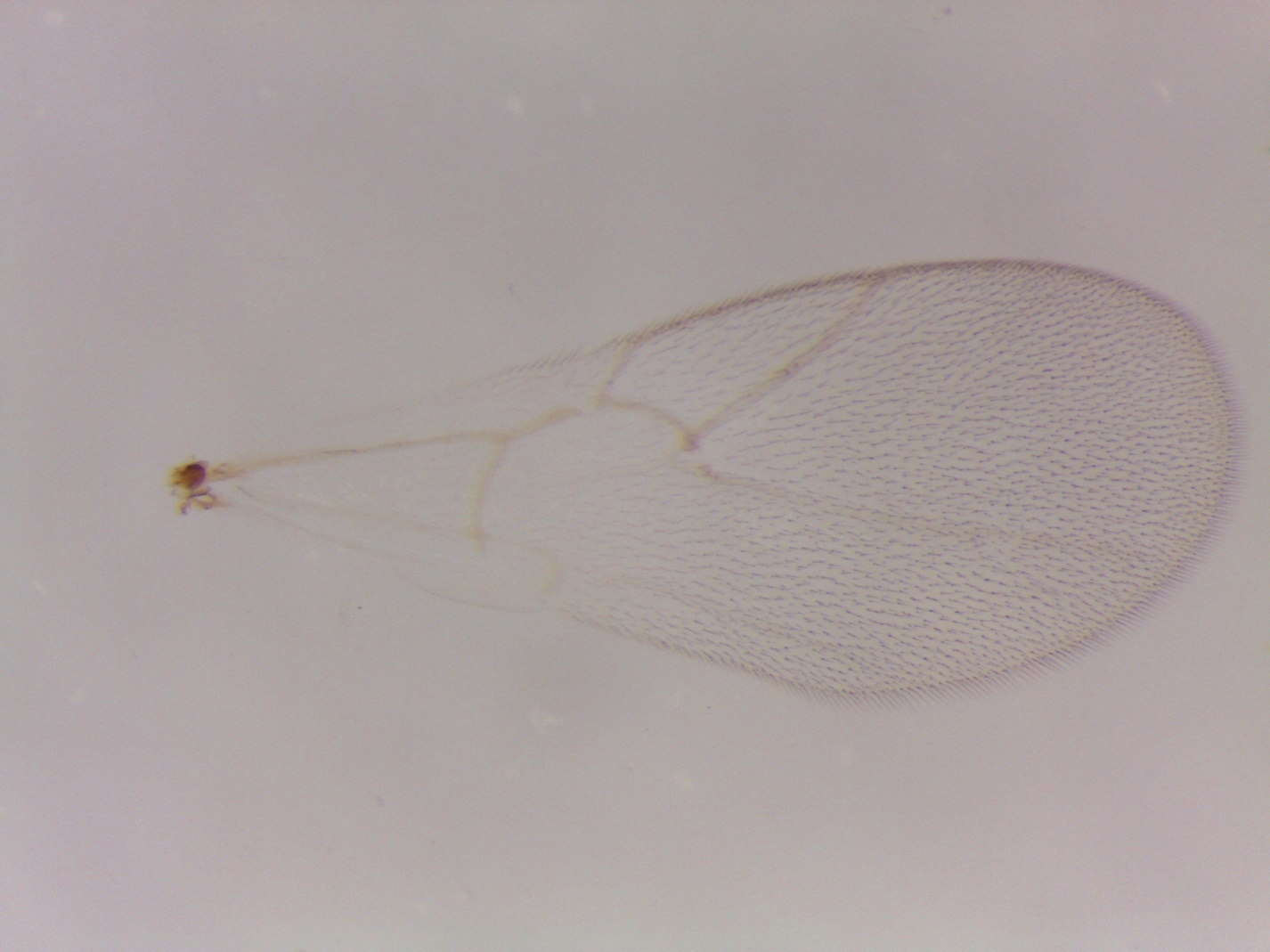

Figure S46. *Ceroptres* sp. 18-19 – 963-051-13A – female from *Callirhytis quercusfutilis* on *Quercus macrocarpa* in Iowa City, IA.

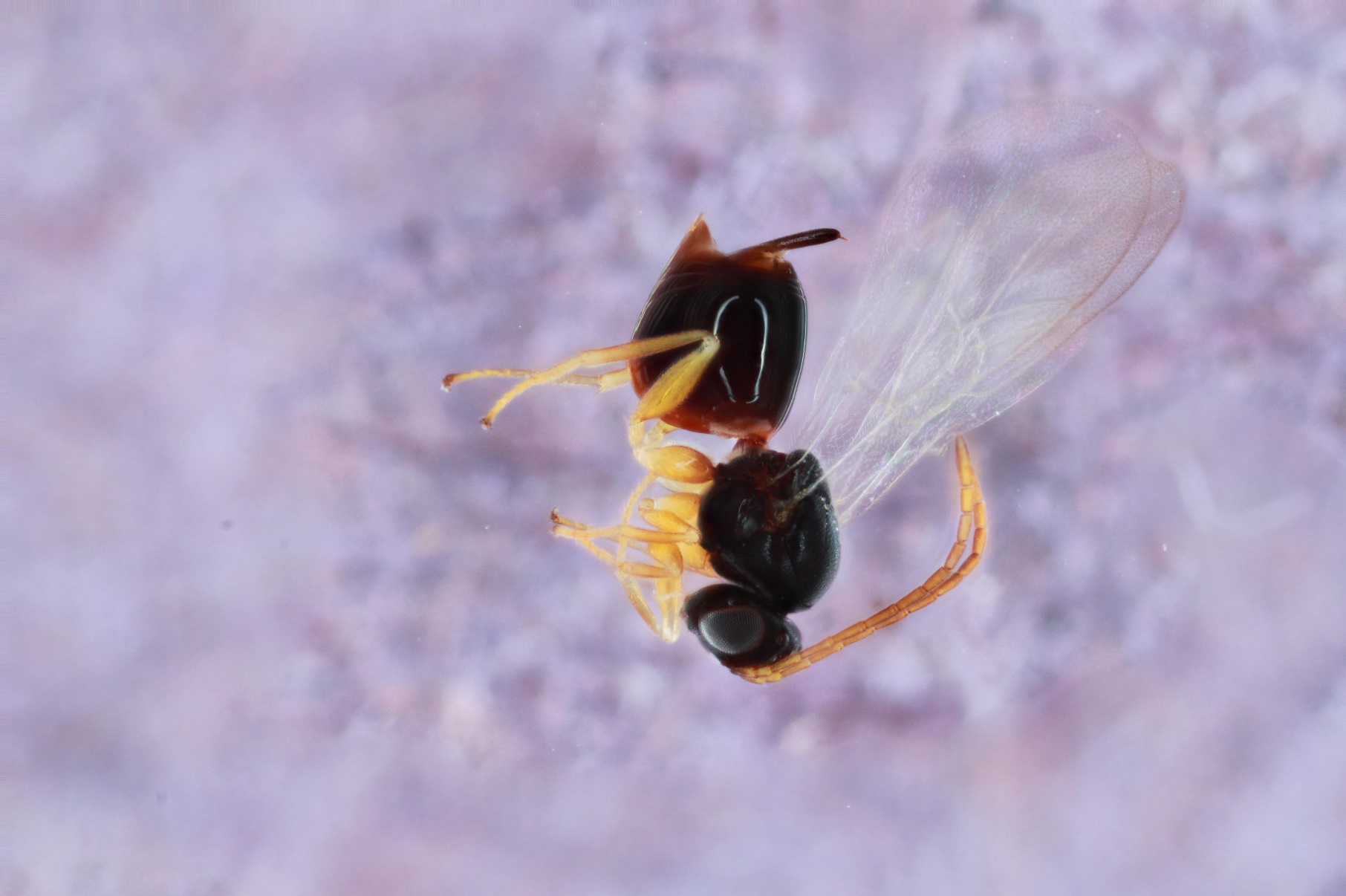

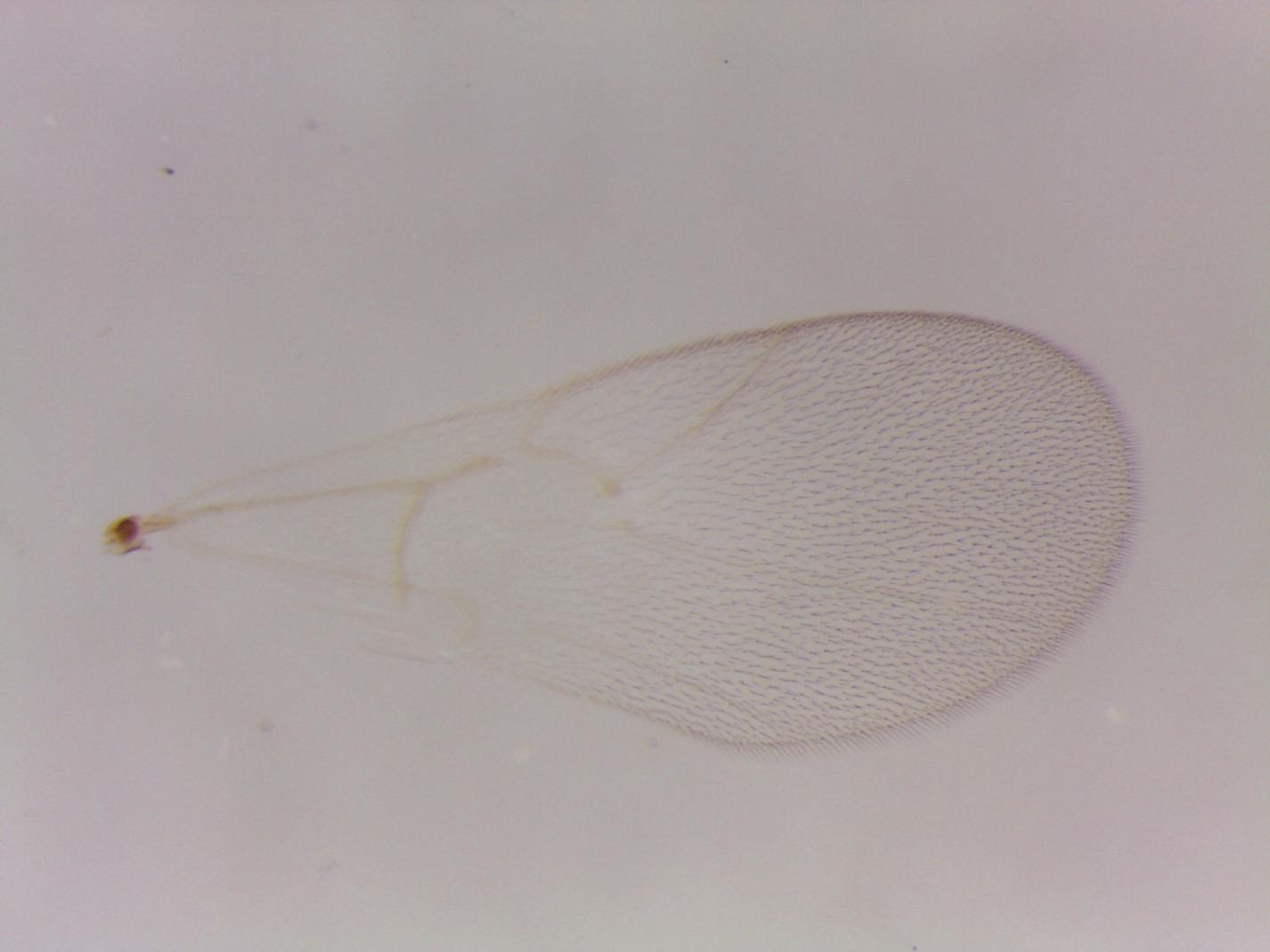

Figure S47. *Ceroptres* sp. 20 – 1551-005-8 – female from *Andricus dimorphus* on *Quercus macrocarpa* in Iowa City, IA.

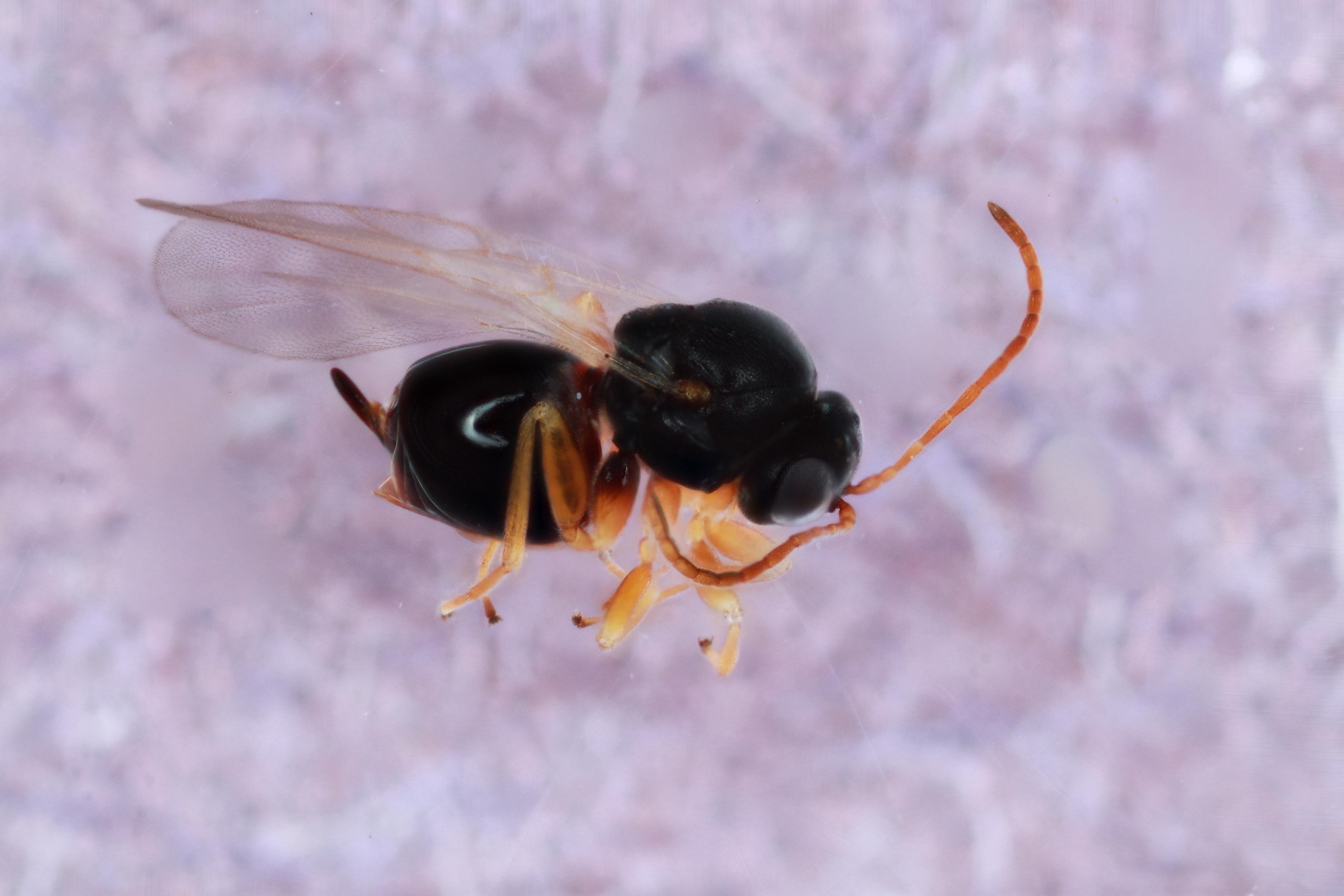

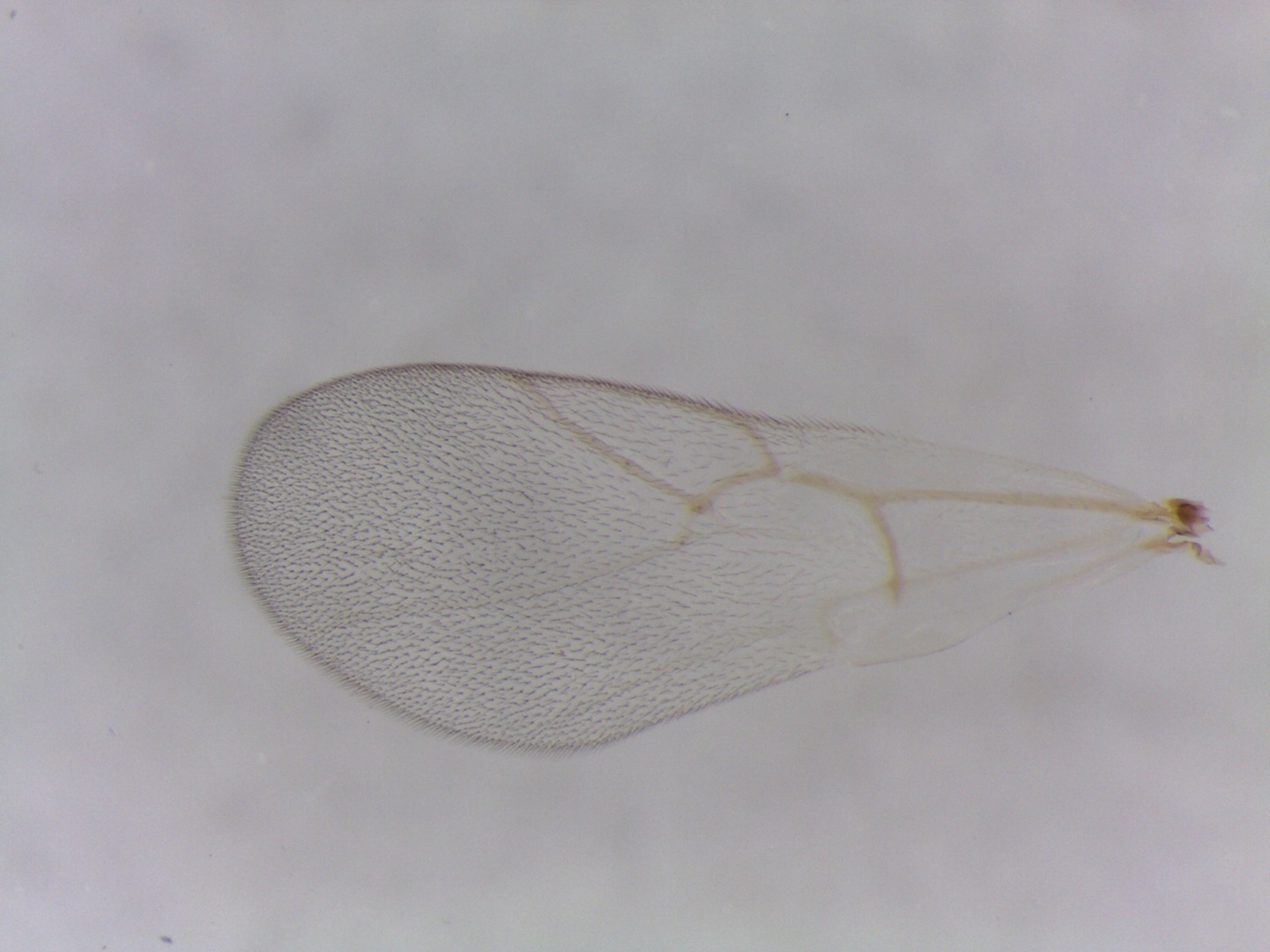

Figure S48. *Ceroptres* sp. 20 – 645-005-3 – female from *Andricus dimorphus* on *Quercus macrocarpa* in Iowa City, IA.

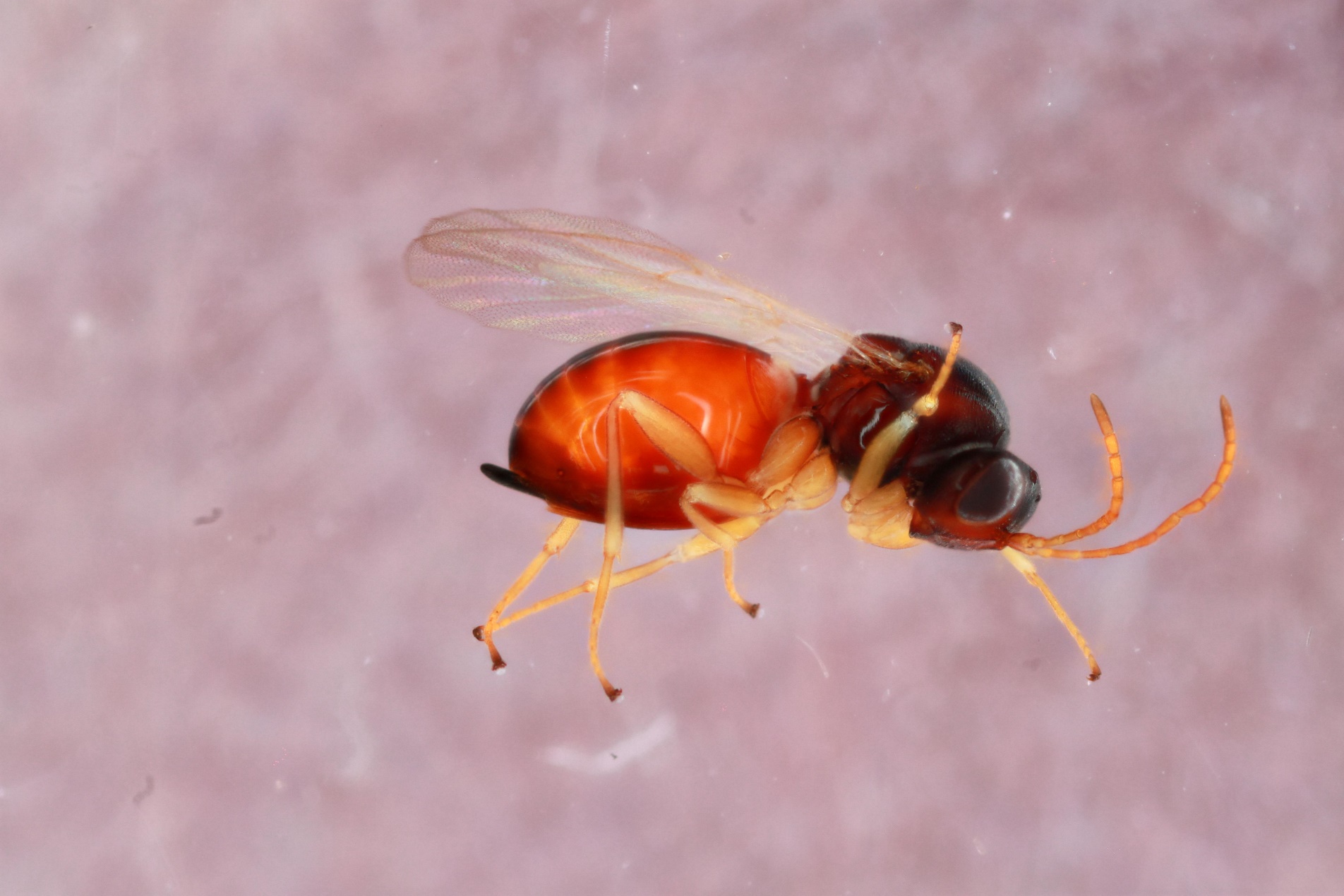

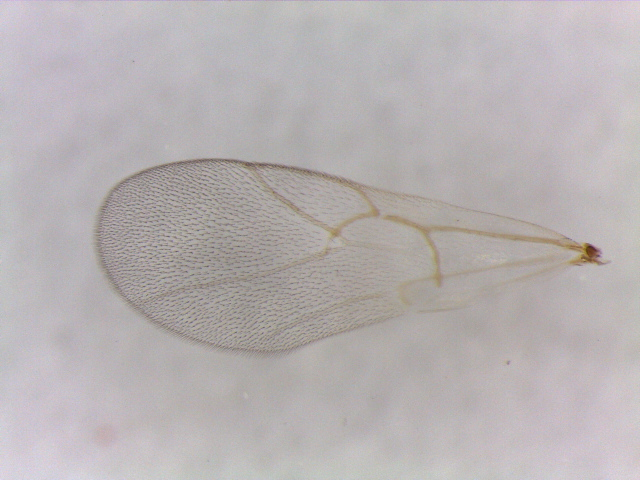

Figure S49. *Ceroptres* sp. 20 – 166-008-1 –male from *Druon ignotum* on *Quercus alba* in Iowa City, IA.

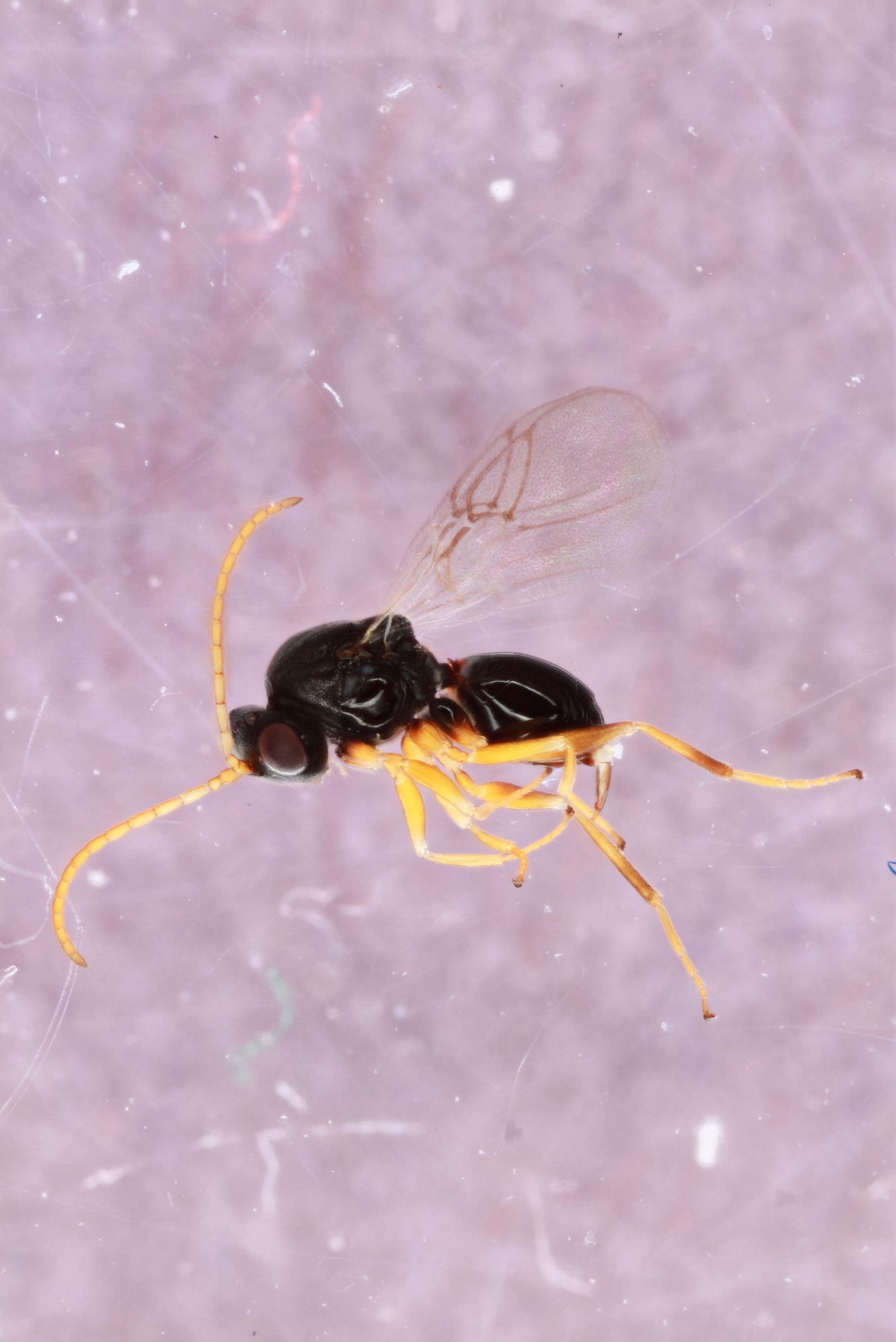

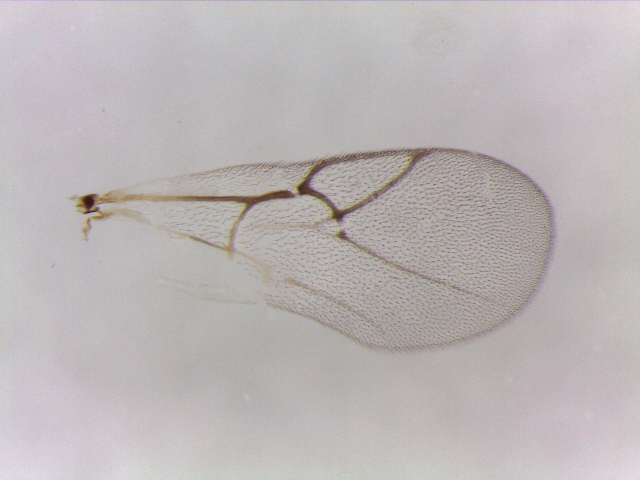

Figure S50. *Ceroptres* sp. 21 – 1578-153-1A – female from *Neuroterus laurifoliae* on *Quercus imbricariae* in Iowa City, IA.

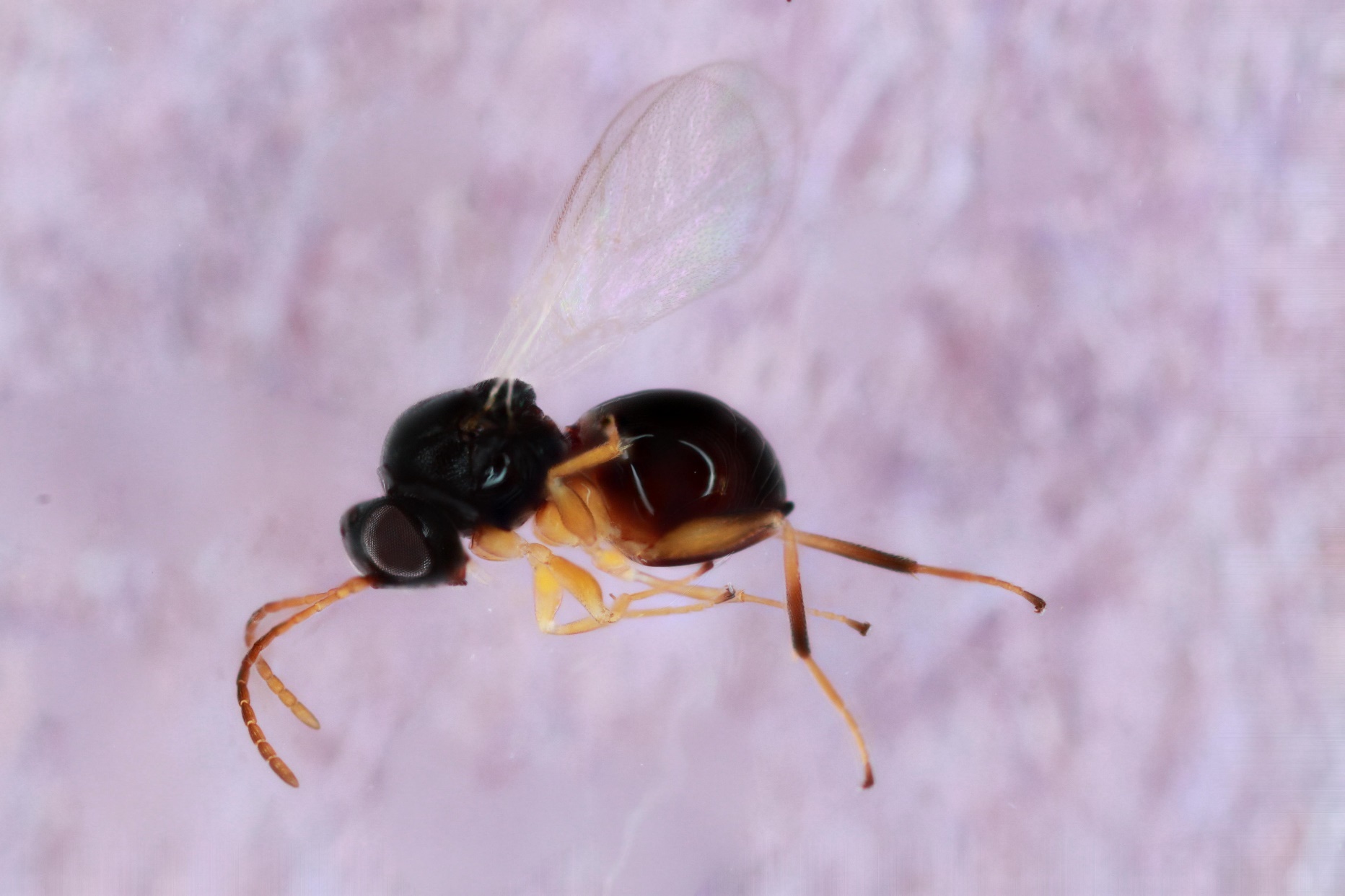

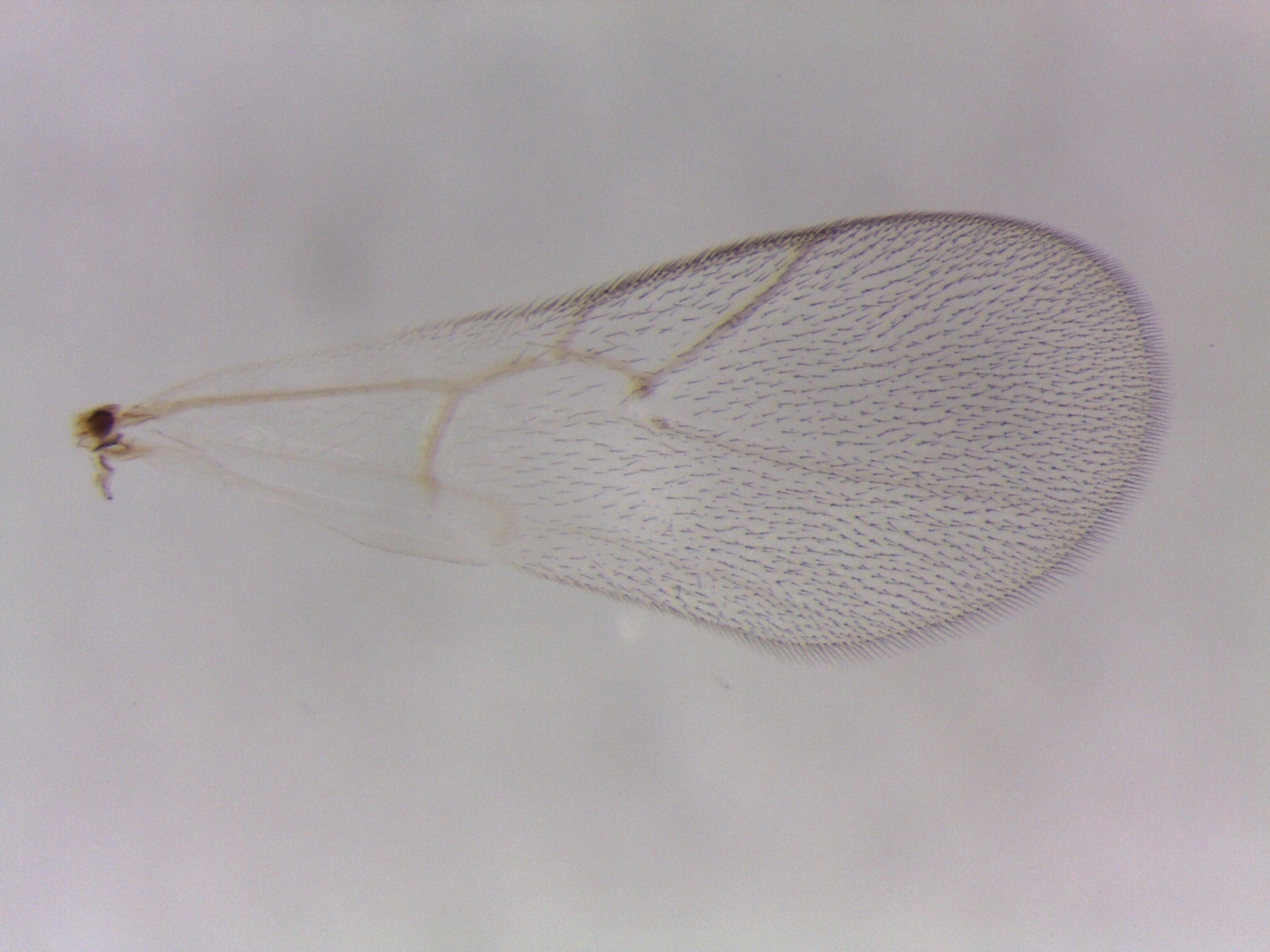

Figure S51. *Ceroptres* sp. 21 – 1578-153-3 – female from *Neuroterus laurifoliae* on *Quercus imbricariae* in Iowa City, IA.

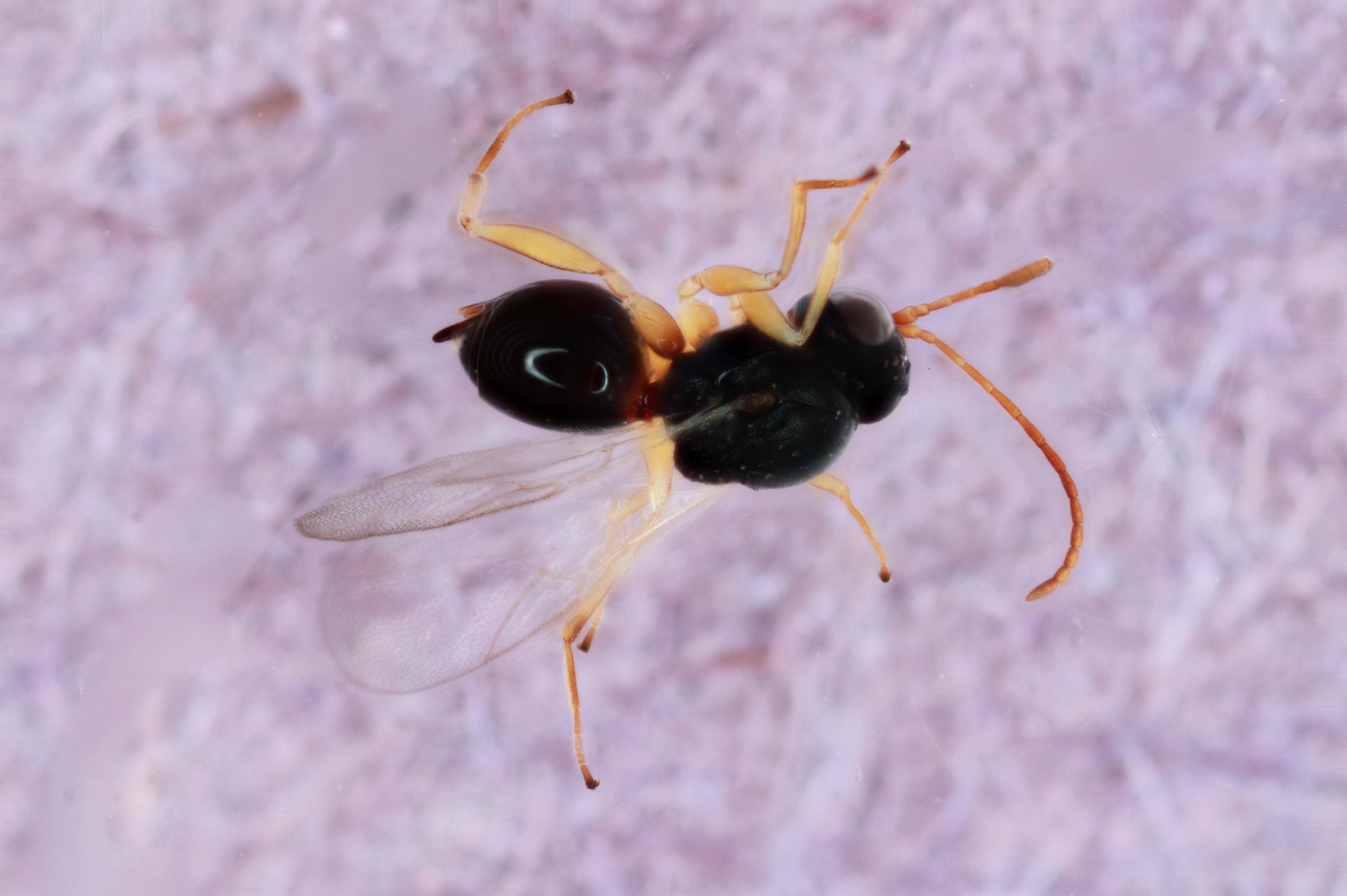

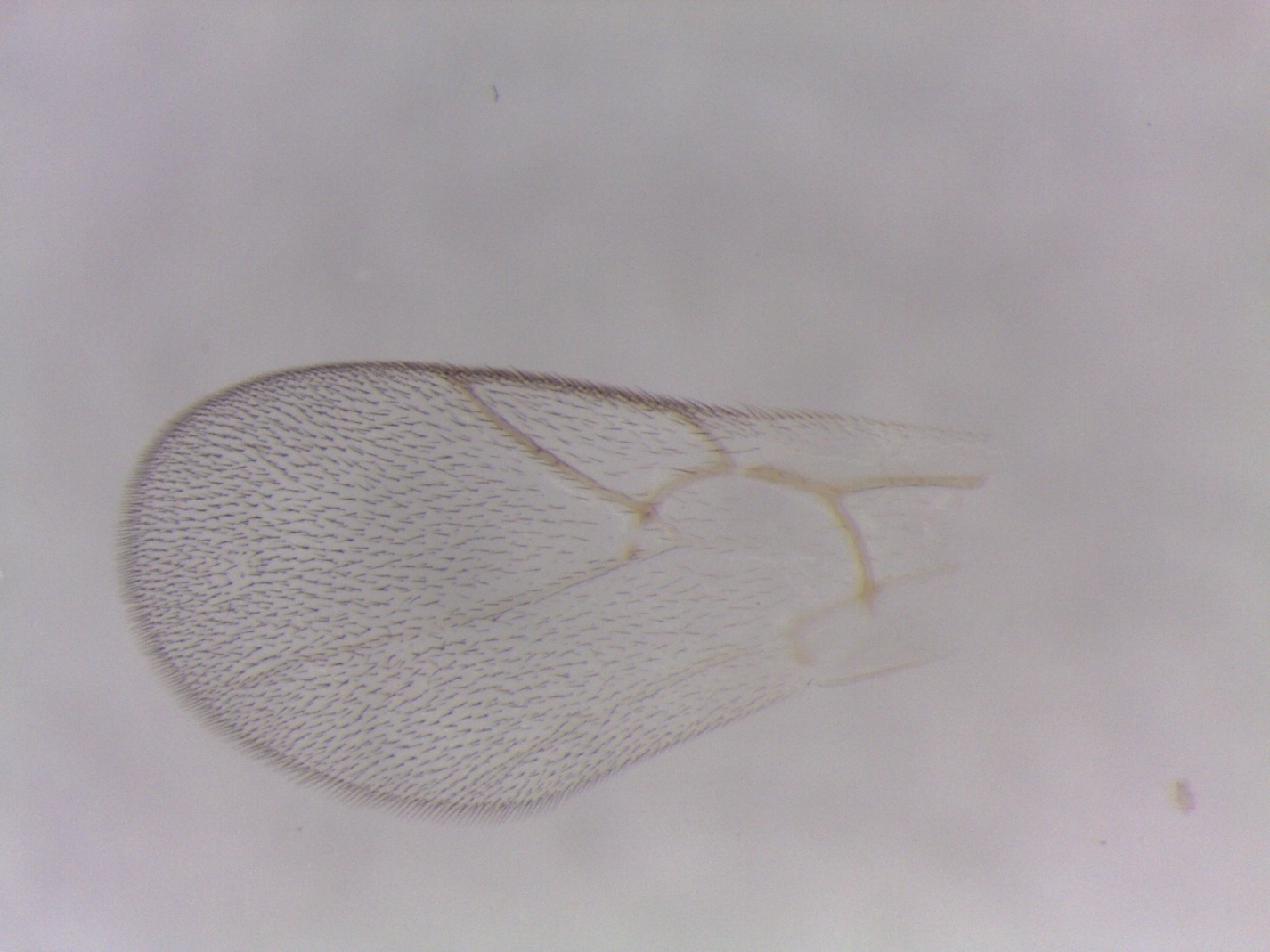

Figure S52. *Ceroptres* sp. 22 – 1219-014-2 – female from *Callirhytis furva* on *Quercus palustris* in Iowa City, IA.

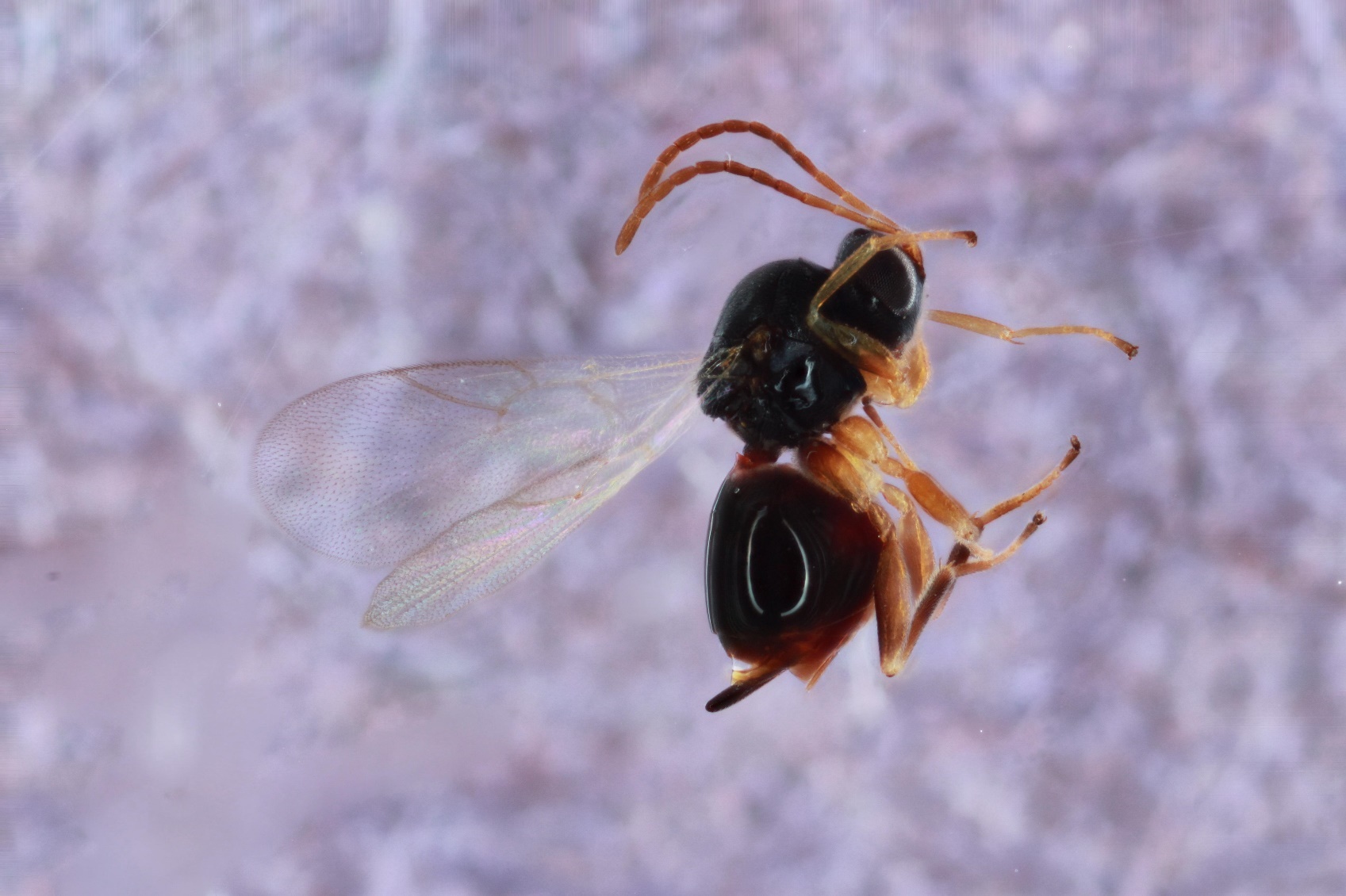

Figure S53. *Ceroptres* sp. 23 – 360-021-82A – female from *Neuroterus saltarius* on *Quercus bicolor* in Iowa City, IA.

Figure S54. *Ceroptres* sp. 23 – 652-008-4 – female from *Druon ignotum* on *Quercus bicolor* in Iowa City, IA.

Figure S55. *Ceroptres* sp. 24 – 667-062-1 – male from *Zopheroteras sphaerula* on *Quercus rubra* in Tiffin, IA.

Figure S56. *Ceroptres* sp. 24 – 668-062-1 – male from *Zopheroteras sphaerula* on *Quercus rubra* in Tiffin, IA.

Figure S57. *Ceroptres* sp. 25 – 662-082-2 – female from *Phylloteras rubinum* on *Quercus alba* in Iowa City, IA.

Figure S58. *Ceroptres* sp. 25 – 899-051-19 – female from *Callirhytis quercusfutilis* on *Quercus alba* in Iowa City, IA.

Figure S59. *Ceroptres* sp. 26-27-28 – 1568-023-4 –male from *Philonix fulvicollis* on *Quercus alba* in Vestal, NY.

Figure S60. *Ceroptres* sp. 26-27-28 – 646-001-6 – male from *Acraspis quercushirta* on *Quercus macrocarpa* in Iowa City, IA.

Figure S61. *Ceroptres* sp. 26-27-28 – 1456-001-2 – female from *Acraspis quercushirta* on *Quercus macrocarpa* in Spring Green, WI.

Figure S62. *Ceroptres* sp. 26-27-28 – 431-004-1 – male from *Acraspis villosa* on *Quercus macrocarpa* in Iowa City, IA.

Figure S63. *Ceroptres* sp. 26-27-28 – 1566-002-4 – female from *Acraspis erinacei* on *Quercus alba* in Vestal, NY (no wing picture).

Figure S64. *Ceroptres* sp. 26-27-28 – 591-023-1 – male from *Philonix fulvicollis* on *Quercus alba* in Knoxville, TN.

Figure S65. *Ceroptres* sp. 26-27-28 – 648-023-1 – female from *Philonix fulvicollis* on *Quercus alba* in Iowa City, IA.

Figure S66. *Ceroptres* sp. 30a – 558-081-16 – male from *Druon pattoni* on *Quercus stellata* in Peducah, KY.

Figure S67. *Ceroptres* sp. 30b – 1014-055-2A – female from *Neuroterus quercusverrucarum* on *Quercus alba* in Iowa City, IA.

Figure S68. *Ceroptres* sp. 31 – 1243-117-22B – female from *Andricus quercuspetiolicola* on *Quercus stellata* in Austin, TX.

Figure S69. *Ceroptres* sp. 35 – 860-007-28A – female from *Andricus foliaformis* on *Quercus macrocarpa* in Iowa City, IA.
