## Supplemental Tables S4-S6, Figures S70-S85 for "Speciation in kleptoparasites of oak gall wasps often correlates with a shift into a new tree habitat, tree organ, or gall morphospace"

**Figures S70-S85**

**for Ward et al. (*****Speciation in kleptoparasitic oak gall wasps often correlates with shifts into new tree habitats, tree organs, and gall morphospace*)**

Table S4. Gall traits, ranked from most to least influential on the clustering of kleptoparasite species in gall trait space based on regression analysis.

| **Trait** | ***R^2^*** | **F value** | ***P* value** |
| --- | --- | --- | --- |
| Polythalamous | 0.65 | 124.40 | <<<0.001 |
| Detachable | 0.65 | 122.80 | <<<0.001 |
| Fleshy-woody | 0.30 | 29.03 | <<0.001 |
| Clustered | 0.19 | 15.72 | <0.001 |
| Size | 0.15 | 11.95 | <0.001 |
| Fleshy | 0.11 | 8.07 | 0.006 |
| Wooly | 0.07 | 5.43 | 0.023 |
| Woody | 0.05 | 3.84 | 0.054 |
| Hairy | 0.05 | 3.54 | 0.064 |
| Nectar | 0.04 | 3.35 | 0.072 |
| Hollow | 0.02 | 1.56 | 0.216 |
| Textured | 0.01 | 0.64 | 0.427 |
| Fiber | 0.01 | 0.56 | 0.455 |
| Bract | 0.01 | 0.52 | 0.474 |
| Spiny | 0.004 | 0.28 | 0.601 |

Table S5. ParaFitLink1 and ParaFitLink2 analysis results for *Synergus* wasps and their gall wasp hosts. Interactions with P ≤ 0.05 for both link analyses are bolded and in red type.

| Host | Parasite | F1.stat | p.F1 | F2.stat | p.F2 |
| --- | --- | --- | --- | --- | --- |
| Acraspis erinacei | S. erinacei A | 1.12E-03 | 0.098819012 | 0.033968045 | 0.062459375 |
| Acraspis erinacei | S. oneratus C | 1.29E-03 | 0.105928941 | 0.039167688 | 0.064539355 |
| Acraspis erinacei | S. punctatus A | 8.98E-04 | 0.275587244 | 0.027273104 | 0.194928051 |
| Acraspis pezomachoides | S. erinacei A | 1.13E-03 | 0.094589054 | 0.034292358 | 0.058949411 |
| Acraspis pezomachoides | S. oneratus C | 1.23E-03 | 0.120128799 | 0.037299654 | 0.074689253 |
| Acraspis quercushirta | S. erinacei B | 1.07E-03 | 0.111938881 | 0.032491699 | 0.070849292 |
| Acraspis quercushirta | S. oneratus B | -9.49E-05 | 0.784472155 | -0.002882615 | 0.799272007 |
| Acraspis villosa | S. villosus A | 7.33E-04 | 0.638143619 | 0.022271365 | 0.525784742 |
| **Amphibolips cookii** | **S. laeviventris B** | **1.76E-03** | **0.01898981** | **0.053488357** | **0.01204988** |
| **Amphibolips quercusinanis** | **S. laeviventris B** | **1.77E-03** | **0.01897981** | **0.053651586** | **0.012079879** |
| **Amphibolips quercusjuglans** | **S. laeviventris B** | **1.71E-03** | **0.02099979** | **0.051954577** | **0.013529865** |
| Amphibolips quercusjuglans | S. magnus | 8.09E-04 | 0.17203828 | 0.024563859 | 0.098729013 |
| **Amphibolips quercusostensackenii** | **S. laeviventris B** | **1.74E-03** | **0.020649794** | **0.052897117** | **0.012939871** |
| Andricus kingi | Synergus sp.7 | 1.29E-03 | 0.638373616 | 0.039035788 | 0.535124649 |
| Andricus pisiformis | S. laeviventris A | 5.83E-04 | 0.540294597 | 0.017694157 | 0.444895551 |
| Andricus quercusstrobilanus | S. oneratus A | 1.42E-04 | 0.710492895 | 0.004316055 | 0.680893191 |
| Atrusca quercuscentricola | S. oneratus C | 2.31E-04 | 0.609103909 | 0.007016555 | 0.573914261 |
| Belonocnema kinseyi | Synergus sp.6 | 2.67E-03 | 0.231457685 | 0.081130201 | 0.150088499 |
| Callirhytis quercuspunctata | S. lignicola | -4.97E-06 | 0.853921461 | -0.000150802 | 0.854631454 |
| Callirhytis quercusventricosa | S. coniferae | 1.56E-04 | 0.694853051 | 0.004745962 | 0.671533285 |
| Cynips conspicuus | S. ochreus | 6.60E-04 | 0.298767012 | 0.020028739 | 0.207327927 |
| Disholcaspis quercusglobulus | S. campanula A | -2.30E-04 | 0.818051819 | -0.006990019 | 0.841891581 |
| Disholcaspis quercusglobulus | S. laeviventris C | -2.73E-04 | 0.901920981 | -0.008287575 | 0.918760812 |
| Disholcaspis quercusglobulus | S. oneratus B | 2.16E-04 | 0.523154768 | 0.006566371 | 0.475235248 |
| Disholcaspis quercusglobulus | Synergus sp.3 | 1.60E-03 | 0.323656763 | 0.0485745 | 0.228517715 |
| Disholcaspis quercusmamma | S. campanula A | -1.70E-04 | 0.792162078 | -0.005174602 | 0.811141889 |
| Disholcaspis quercusmamma | S. oneratus B | 3.73E-04 | 0.380126199 | 0.011319279 | 0.3099569 |
| Druon ignotum | S. walshii A | 2.70E-04 | 0.590884091 | 0.008208503 | 0.543574564 |
| Druon quercuslanigerum | S. walshii A | 2.11E-04 | 0.6299837 | 0.006409917 | 0.593504065 |
| Dryocosmus minusculus | Synergus sp.8 | 2.79E-03 | 0.242757572 | 0.08464426 | 0.156118439 |
| Philonix fulvicollis | S. oneratus B | -6.11E-04 | 0.968190318 | -0.018563549 | 0.981540185 |
| Philonix fulvicollis | S. oneratus C | -7.73E-04 | 0.969740303 | -0.023461624 | 0.982640174 |
| **Philonix fulvicollis** | **Synergus sp.2** | **5.38E-03** | **0.003459965** | **0.163518217** | **0.00295997** |
| **Philonix fulvicollis** | **Synergus sp.3** | **5.30E-03** | **0.003809962** | **0.161011096** | **0.003199968** |

Table S6. ParaFitLink1 and ParaFitLink2 analysis results for *Ceroptres* wasps and their gall wasp hosts. Interactions with P ≤ 0.05 for both link analyses are bolded and in red type.

| Host | Parasite | Fl.stat | p.F1 | F2.stat | p.F2 |
| --- | --- | --- | --- | --- | --- |
| Andricus foliaformis | Cer sp. 35 | 1.63E-04 | 0.79526205 | 3.64E-03 | 0.80679193 |
| Andricus foliaformis | Cer sp 5b | 5.75E-04 | 0.42337577 | 0.012853292 | 0.44754552 |
| Andricus quercuspetiolicola | Cer sp. 1-32-33-34 | 2.88E-04 | 0.64755352 | 0.006426976 | 0.66821332 |
| Andricus quercuspetiolicola | Cer sp. 31 | 2.61E-04 | 0.72386276 | 0.005827594 | 0.74136259 |
| Callirhytis pigra | Cer sp. 2 | 2.00E-04 | 0.76068239 | 0.004476141 | 0.7701823 |
| Callirhytis quercusfutilis | Cer sp. 15-16 | 1.37E-04 | 0.8596714 | 0.86209138 | 0.003056011 |
| Callirhytis quercusfutilis | Cer sp. 18-19 | 2.09E-04 | 0.7604124 | 0.004681603 | 0.7704423 |
| Callirhytis quercusfutilis | Cer sp. 25 | 1.04E-04 | 0.90485095 | 0.002317863 | 0.90789092 |
| Callirhytis quercusfutilis | Cer sp. 5a | 8.62E-05 | 0.89617104 | 0.001926333 | 0.89807102 |
| Callirhytis quercuspunctata | Cer sp. 3 | -3.18E-05 | 0.87600124 | -0.000711661 | 0.87540125 |
| Callirhytis quercusventricosa | Cer sp. 6C | 6.02E-04 | 0.50946491 | 0.013448537 | 0.53383466 |
| Callirhytis scitula | Cer sp. 11-1.2 | 4.43E-04 | 0.92474075 | 0.009907463 | 0.92837072 |
| Druon ignotum | Cer sp. 20 | 2.09E-04 | 0.9202408 | 0.004678311 | 9.24E-01 |
| Druon ignotum | Cer sp. 23 | 1.05E-04 | 0.92095079 | 0.002337239 | 0.92357076 |
| Loxaulus quercusmammula | Cer sp. 1.3 | 4.92E-04 | 0.56594434 | 0.010992173 | 0.59407406 |
| Melikaiela ostensackeni | Cer sp. 3 | -3.60E-05 | 0.88058119 | -0. 000805546 | 0.8800212 |
| Melikaiela ostensackeni | Cer sp. 6a | 5.28E-04 | 0.20891791 | 0.011809592 | 0.22922771 |
| Melikaiela ostensackeni | Cer sp. 7-8-9 | 7.86E-04 | 0.22334777 | 0.01757161 | 0.24900751 |
| Melikaiela tumifica | Cer sp. 10 | 5.07E-04 | 0.34525655 | 0.011341678 | 0.37717623 |
| Neuroterus quercusbatatus | Cer sp. 15-16 | 3.89E-04 | 0.60638394 | 0.008699246 | 0.6200238 |
| Neuroterus quercusirregularis | Cer sp. 17 | 5.78804 | 0.21111789 | 0.0129114 | 0.24125759 |
| **Neuroterus quercusverrucarum** | **Cer sp. 30b** | **6.10E-04** | **0.03876961** | **0.013638276** | **0.04411956** |
| Neuroterus vesicula | Cer sp. 4 | 6.18E-04 | 0.09433906 | 0.013810412 | 0.10452895 |
| Zopheroteras sphaerula | Cer sp. 24 | 6.70E-04 | 0.0799592 | 0.014970699 | 8.77E-02 |
| Zopheroteras sphaerula | Cer sp. 3 | 3.56E-04 | 0.39930601 | 0.007951628 | 0.41220588 |

Figure S70. Maximum likelihood phylogeny built in IQ-TREE based on the *Synergus* 90% complete data matrix (993 loci).

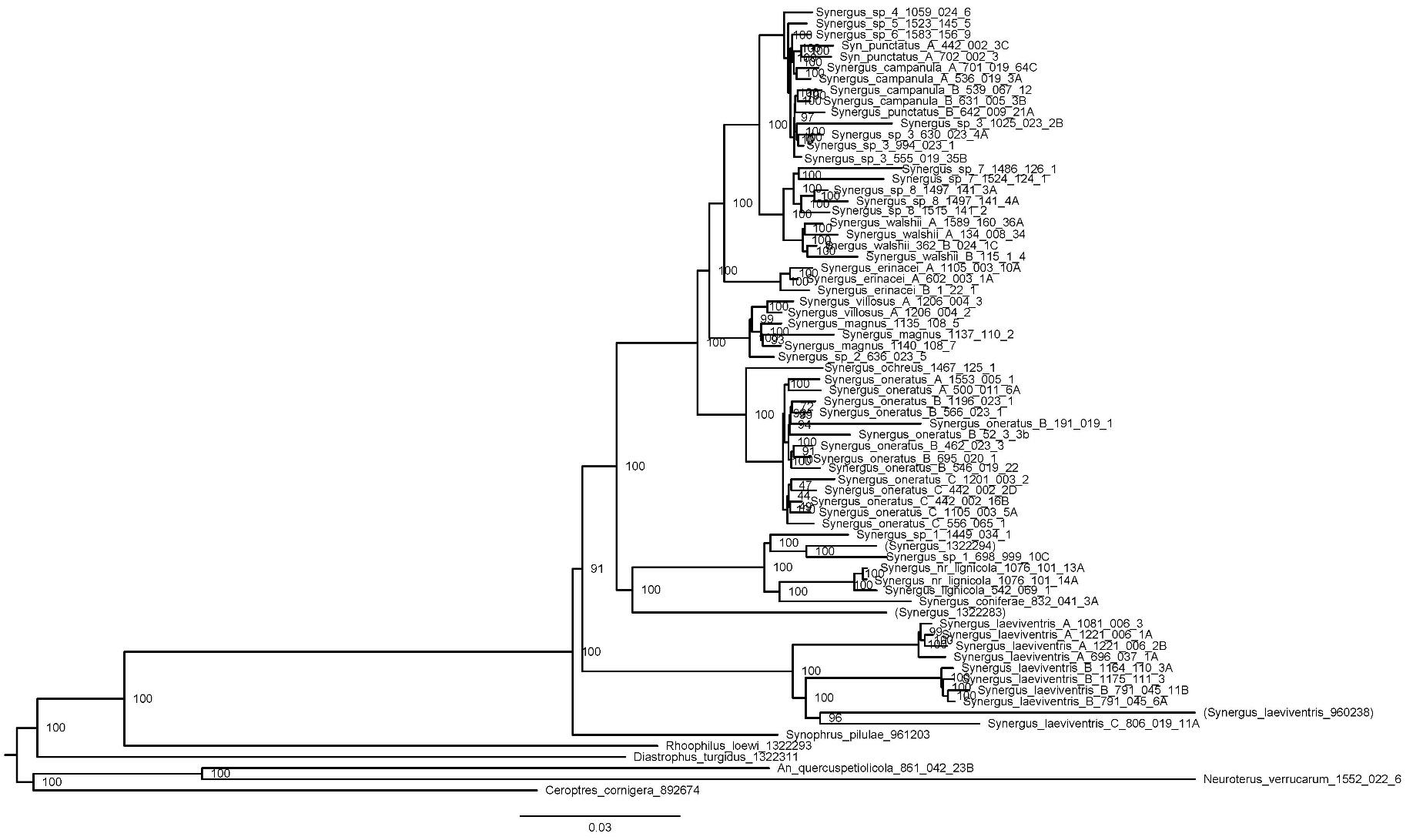

Figure S71. Maximum likelihood phylogeny built in IQ-TREE based on the *Ceroptres* 90% complete data matrix (1019 loci).

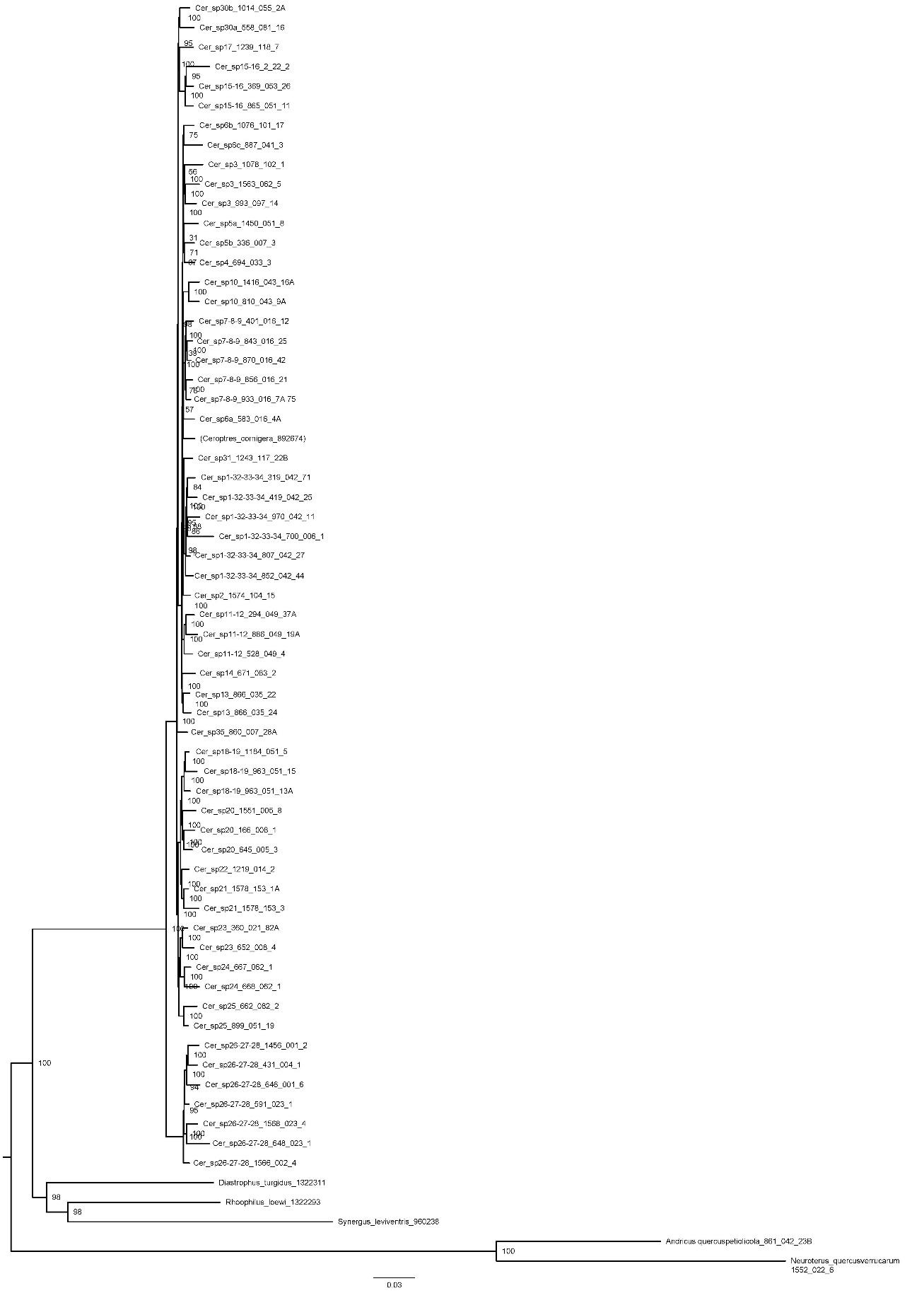

Figure S72. Dendrogram of all kleptoparasite species (*Synergus* and *Ceroptres*) visualizing similarity with respect to traits of host galls. The two primary clusters, “A” and “B” are shown in black boxes, with sub-clusters denoted with grey boxes. A key to species names abbreviations can be found in Table S3. Columns of circles, from left to right, denote host gall external defenses, gall sizes, gall internal defenses, and gall spatial defenses. See figure S73 for color legend.

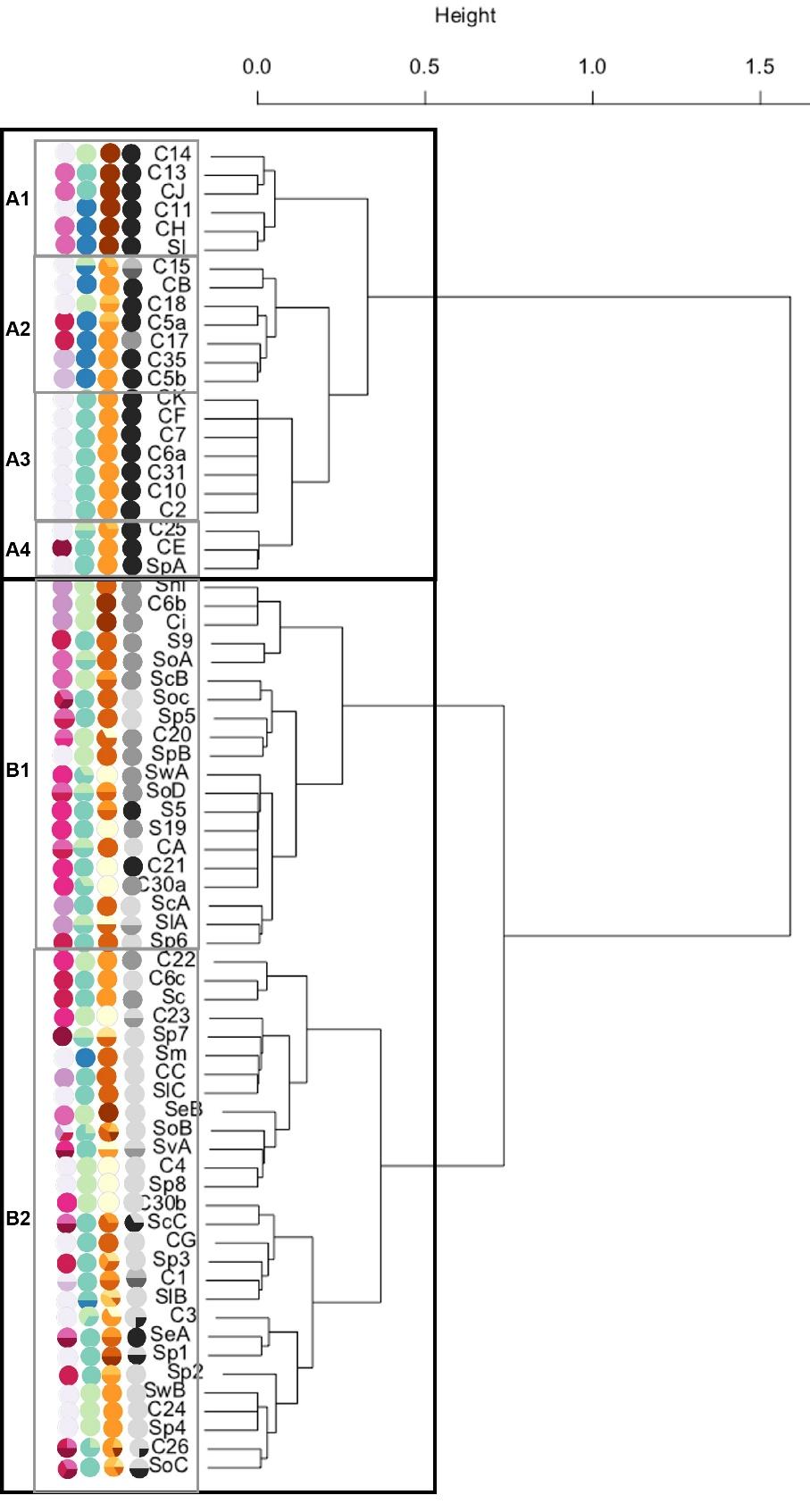

Figure S73. Centroids of host gall trait combinations for each kleptoparasite species in gall trait space. A) with gall spatial defenses; B) showing gall internal defenses; C) showing gall external defenses; D) showing gall sizes.

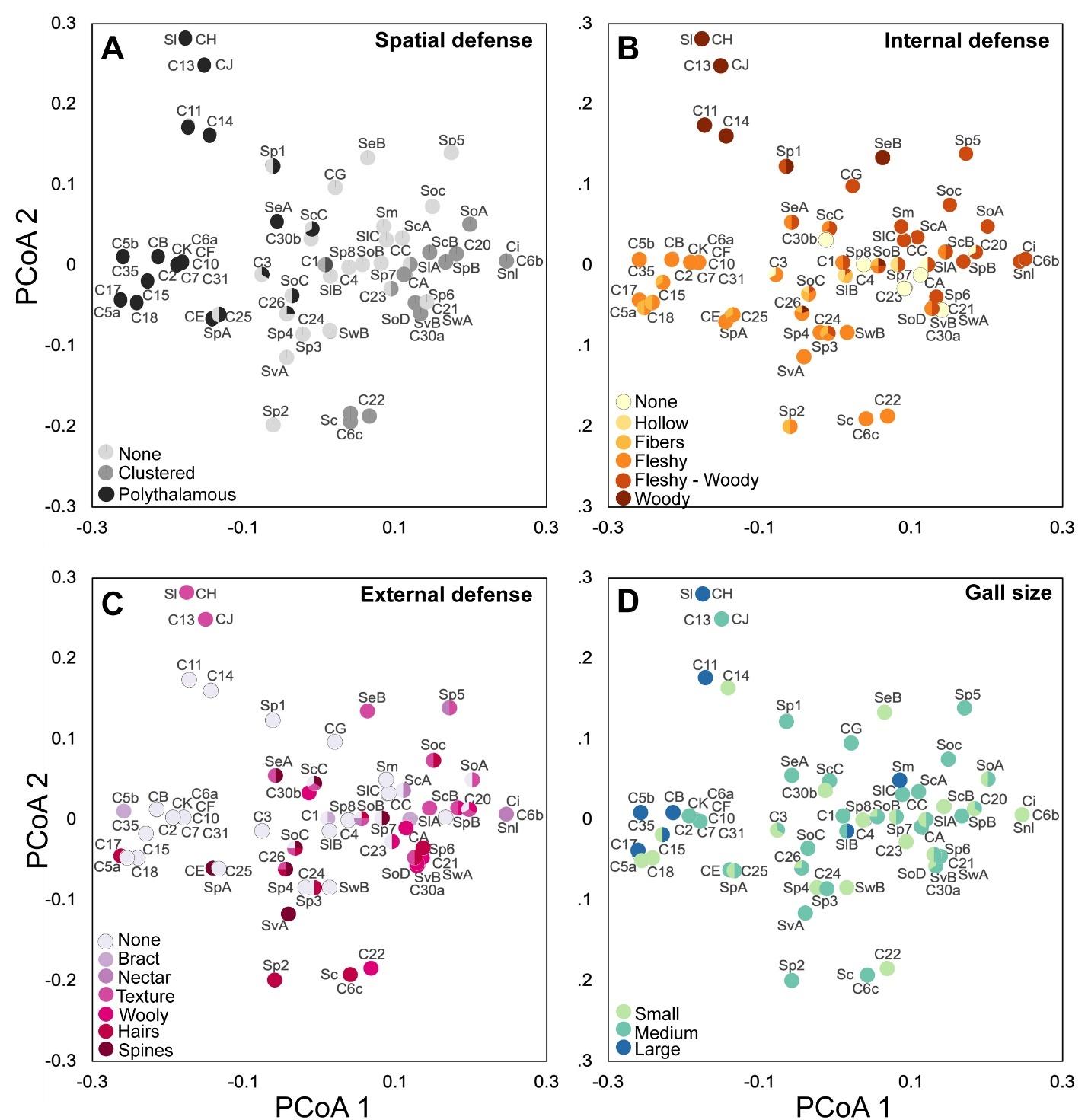

Figure S74. Input tanglegram for *Synergus* PACo analyses.

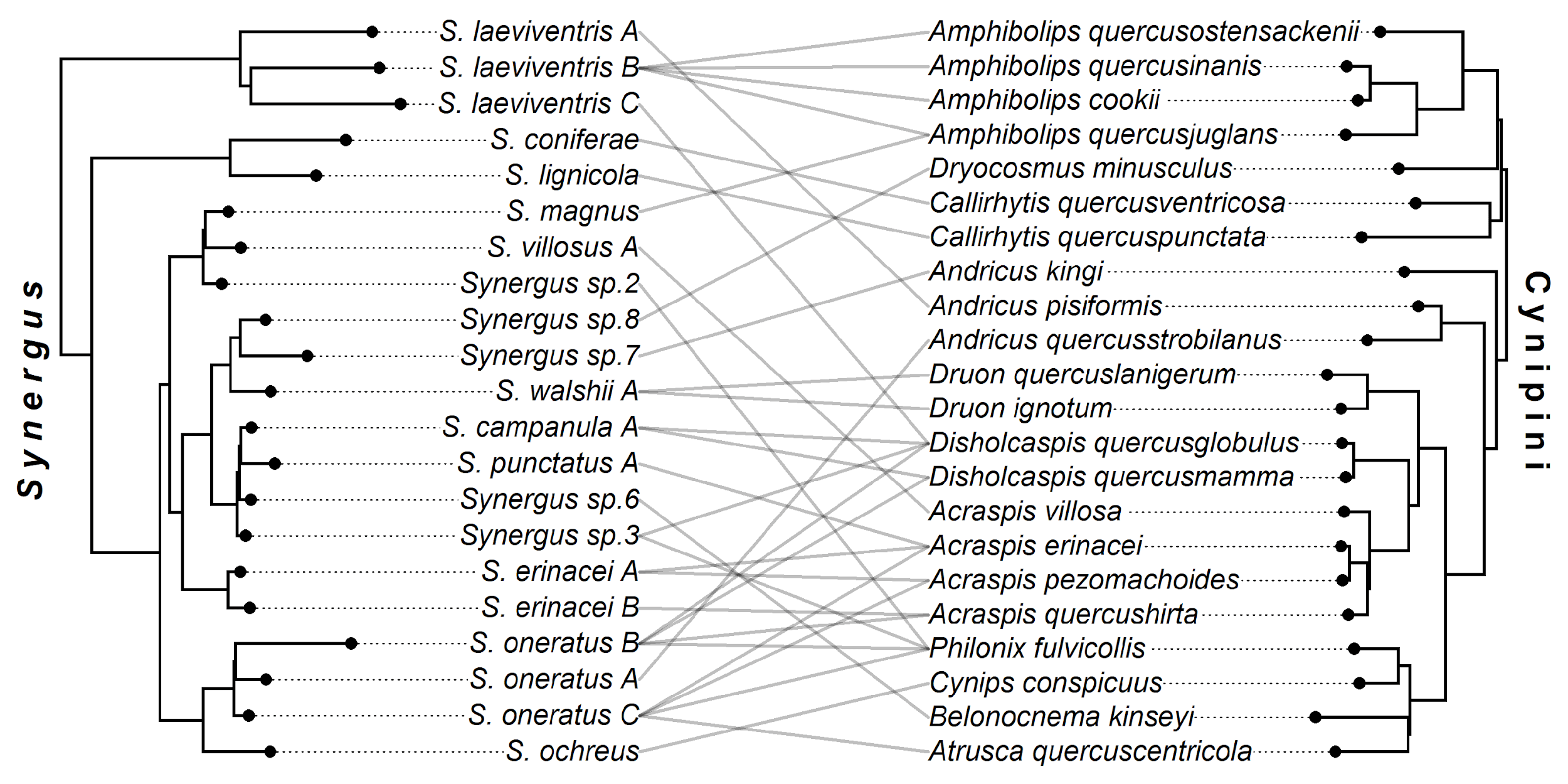

Figure S75. Input tanglegram for PACo Ceroptres analyses

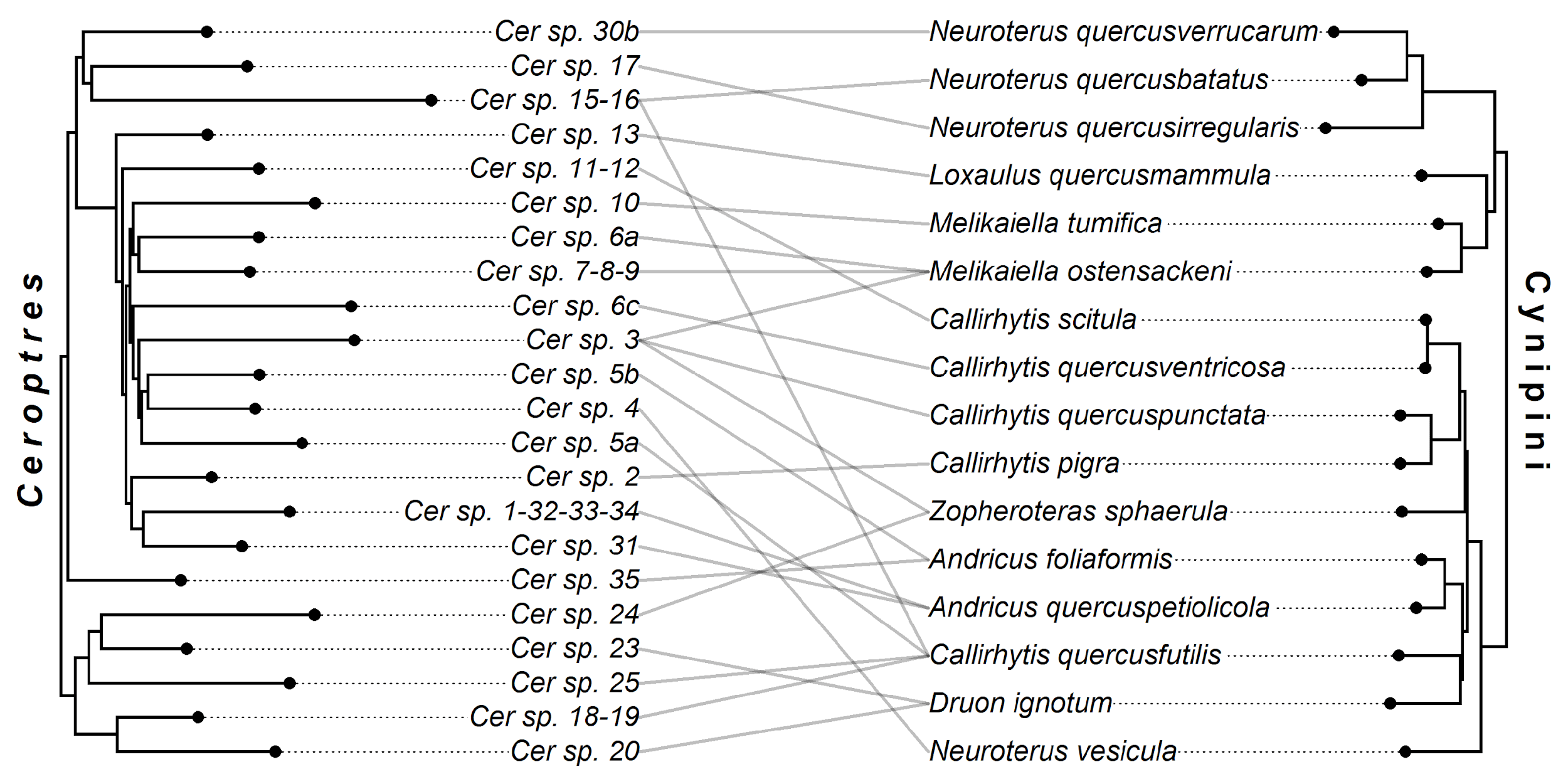

Figure S76. Alternate median eMPRess solution for the cophylogeny of *Synergus* parasites with their gall wasp hosts. Event costs for this solution space were as follows: cospeciation (0), duplication (>2), transfer (1-2), loss (1). With intraspecific events subtracted, this solution produced five cospeciation events, 15 host shifting events, and one loss.

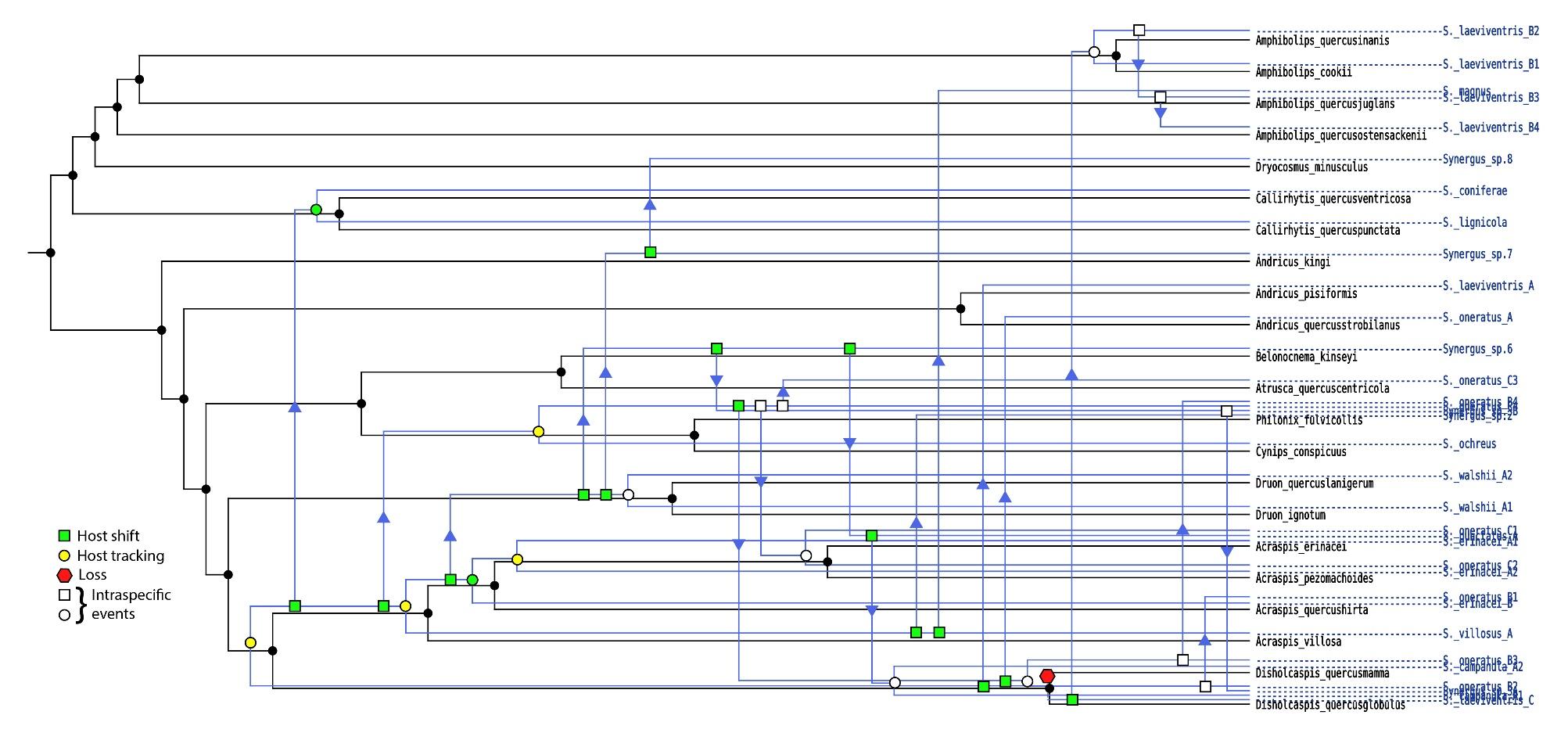

Figure S77. Alternate median eMPRess solution for the cophylogeny of *Ceroptres* parasites with their gall wasp hosts. Event costs for this solution space were as follows: cospeciation (0), duplication (>2), transfer (1-2), loss (1). With intraspecific events subtracted, this solution produced five cospeciation events, two duplication events, 15 host shifting events, and one loss.

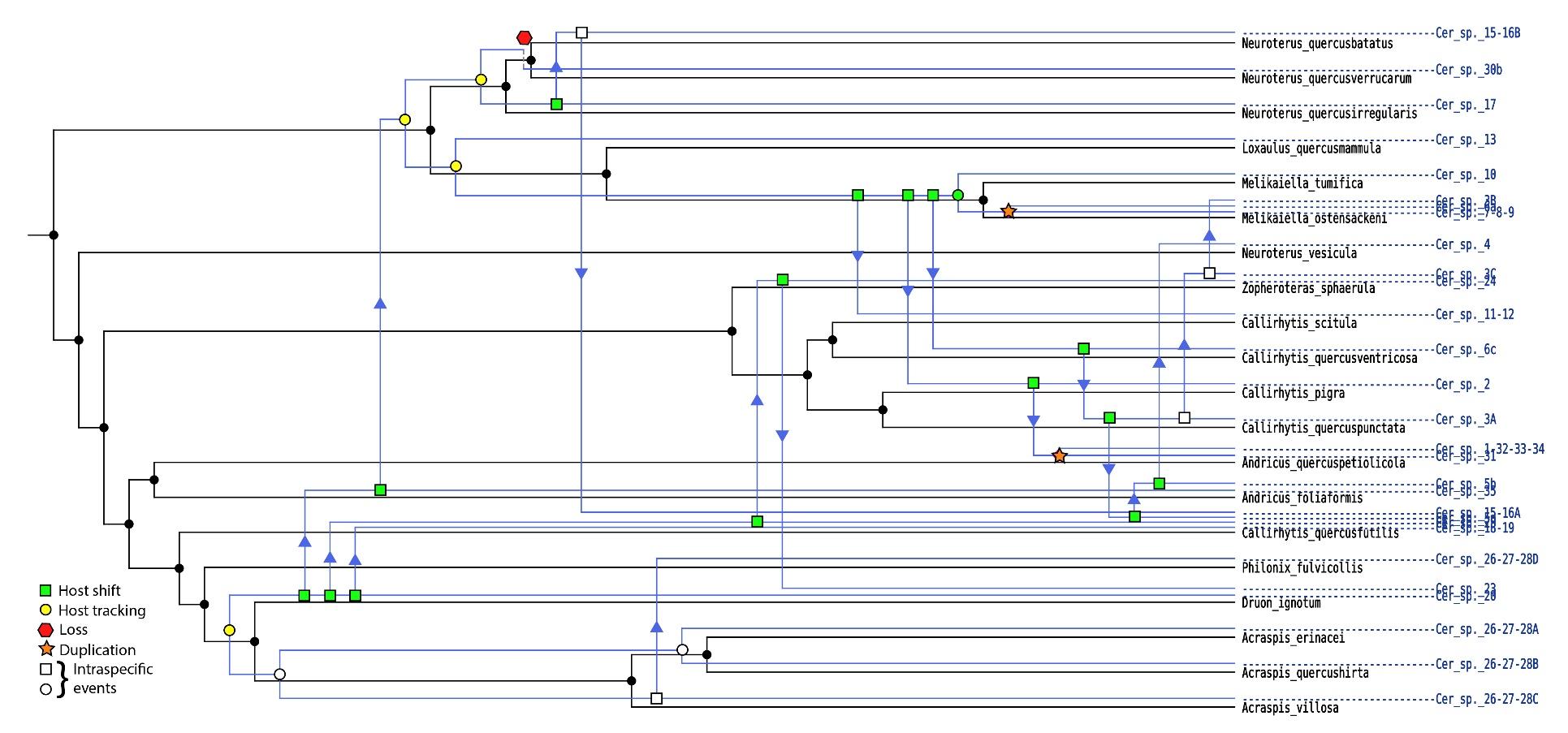

Figure S78. Ancestral state reconstruction of *Synergus* species association with oak subsections.
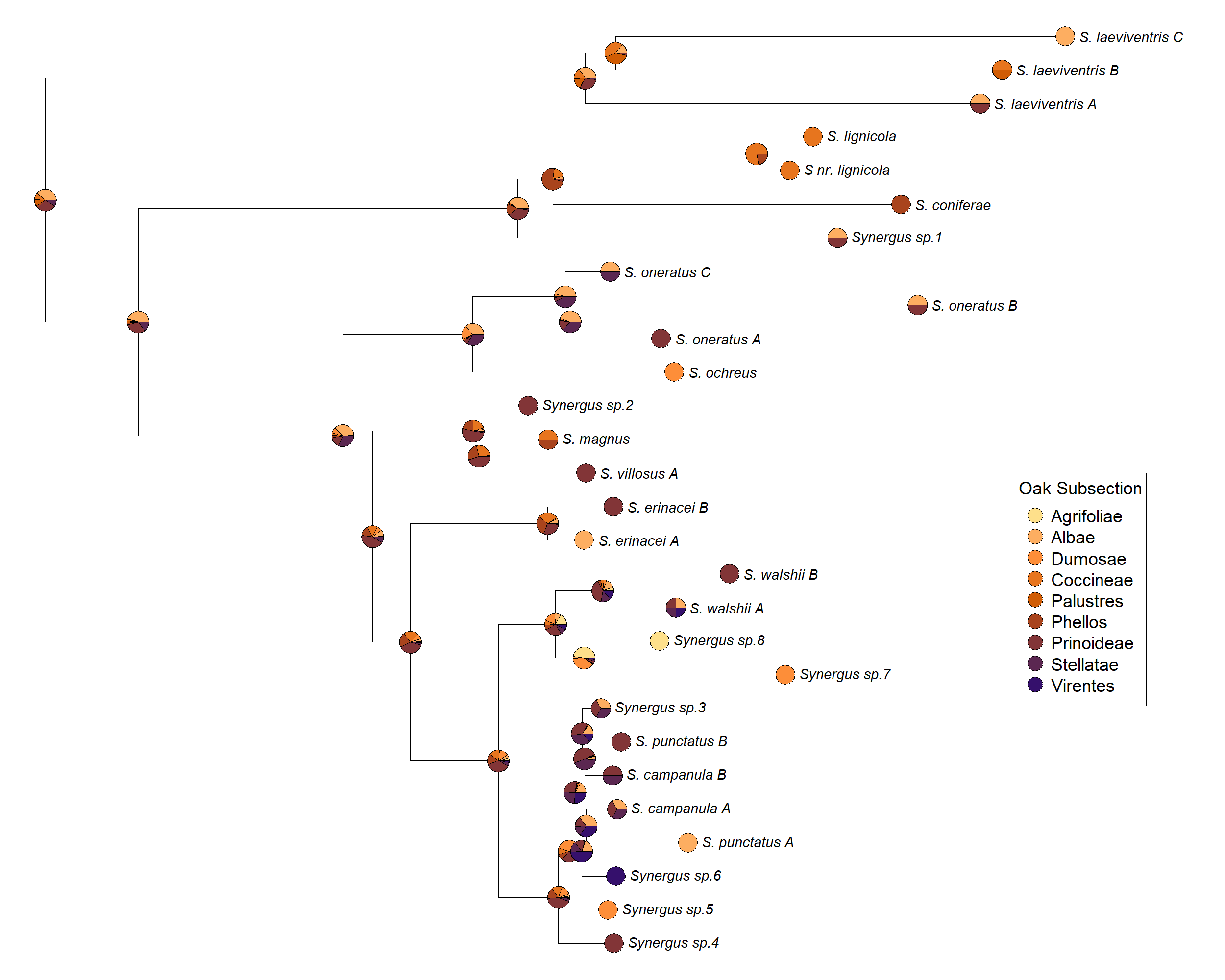

Figure S79. Ancestral state reconstruction of *Synergus* species association with tree organs.
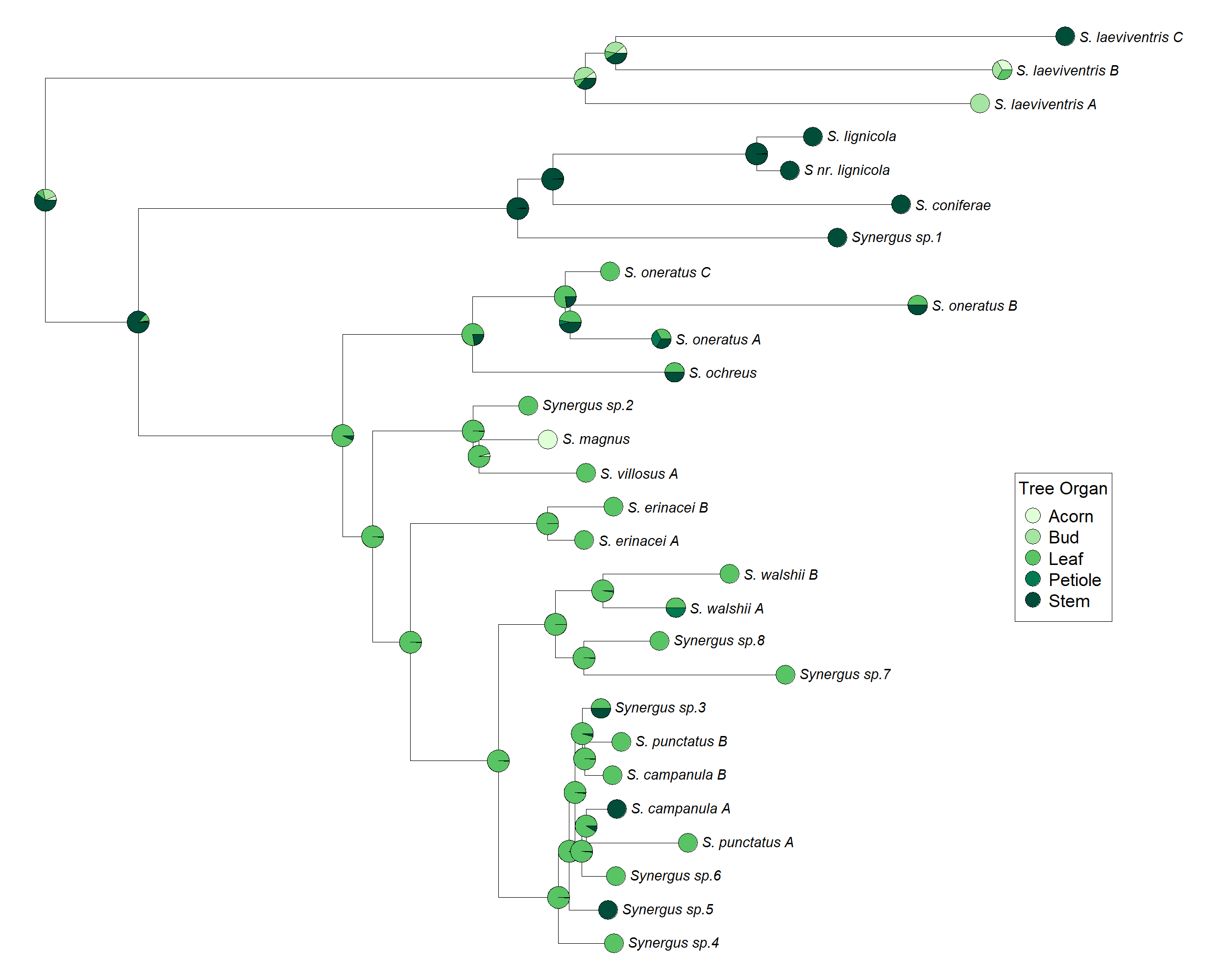
.

Figure S80. Ancestral state reconstruction of *Synergus* species association with morphology. Clusters refer to the dendrogram in Figure S72.

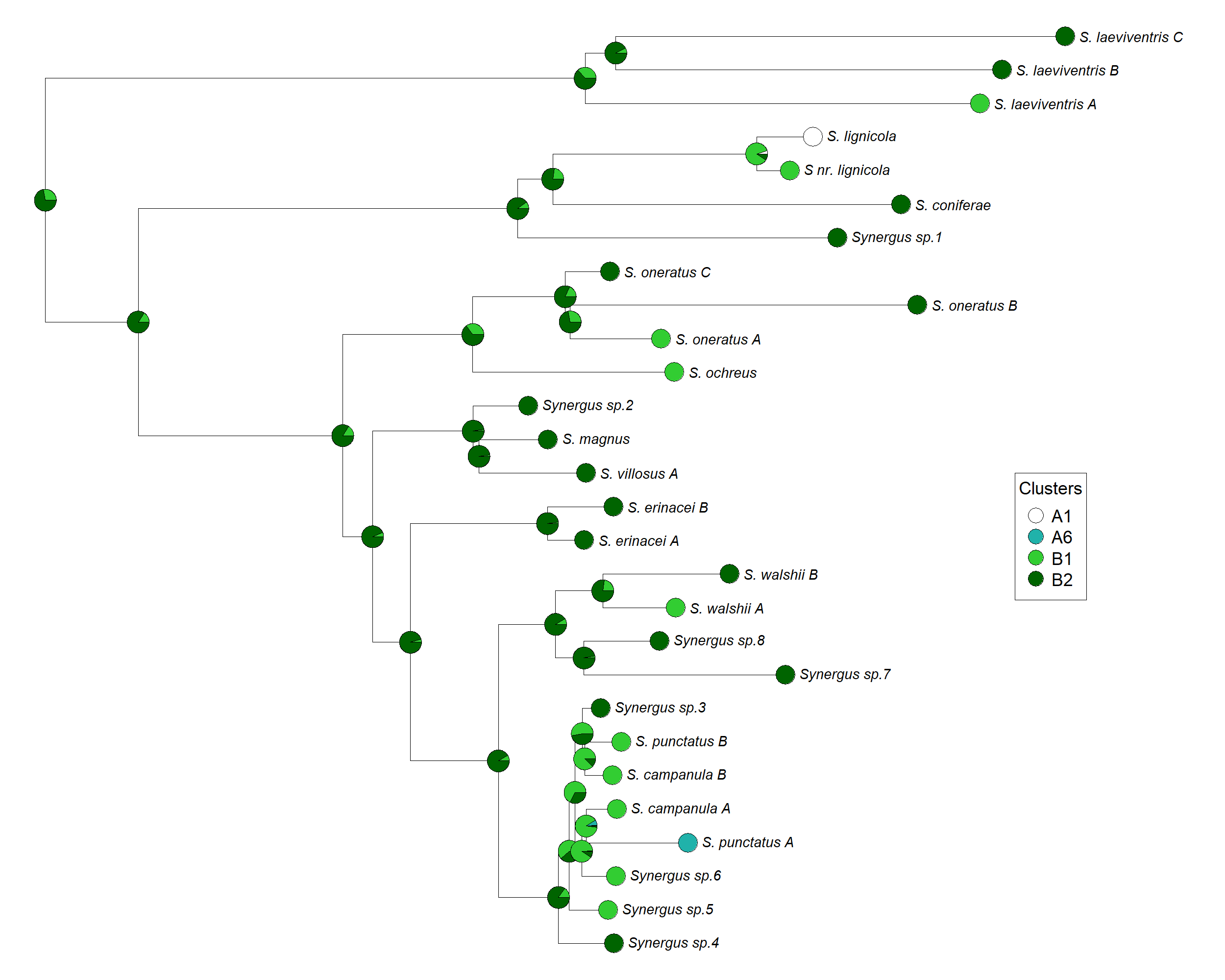

Figure S81. Ancestral state reconstruction of *Synergus* species association with galls of the asexual or sexual generation of gall wasps.

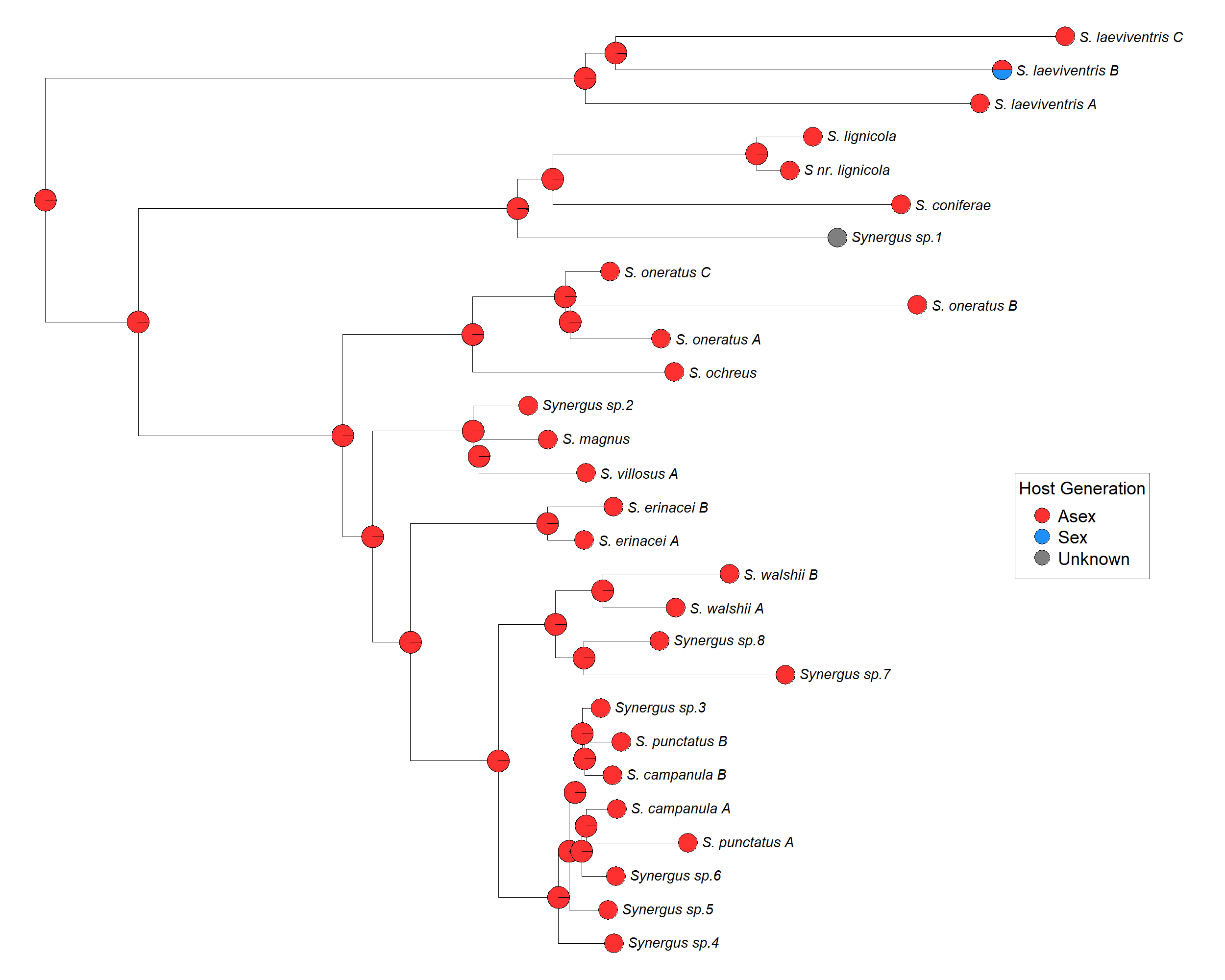

Figure S82. Ancestral state reconstruction of *Ceroptres* species association with oak subsections.
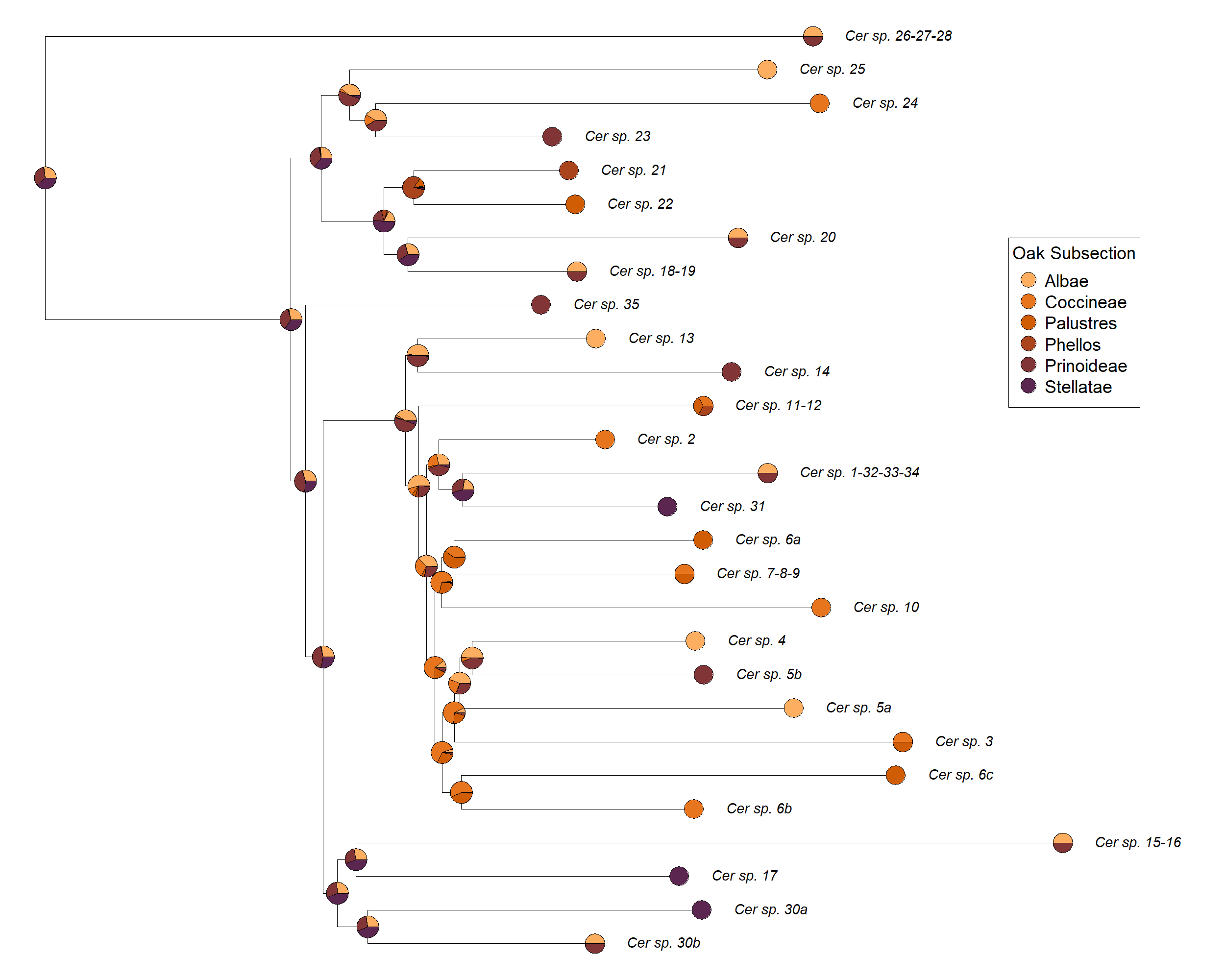

Figure S83. Ancestral state reconstruction of *Ceroptres* species association with tree organs.

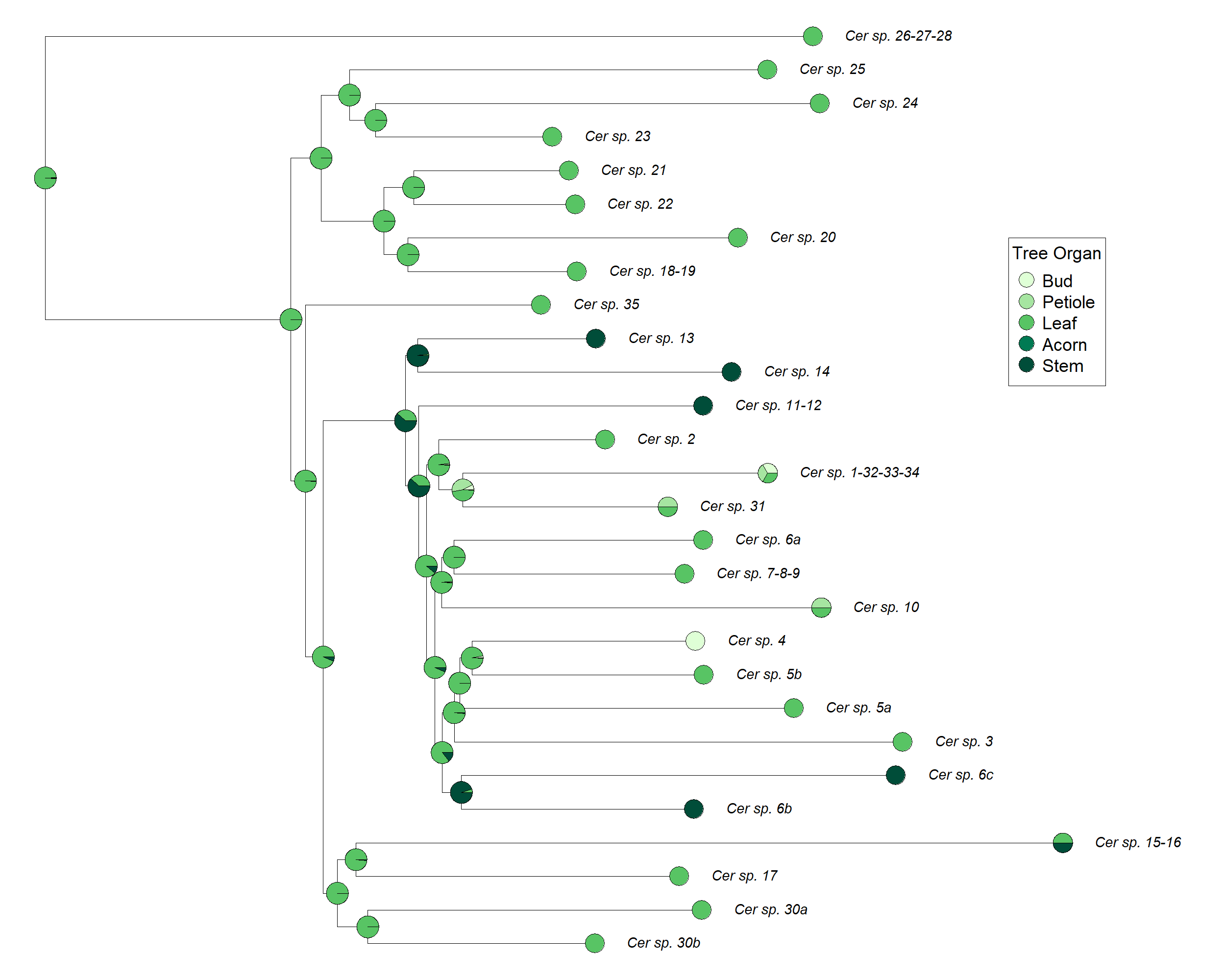

Figure S84. Ancestral state reconstruction of *Ceroptres* species association with morphology. Clusters refer to the dendrogram in Figure S72.

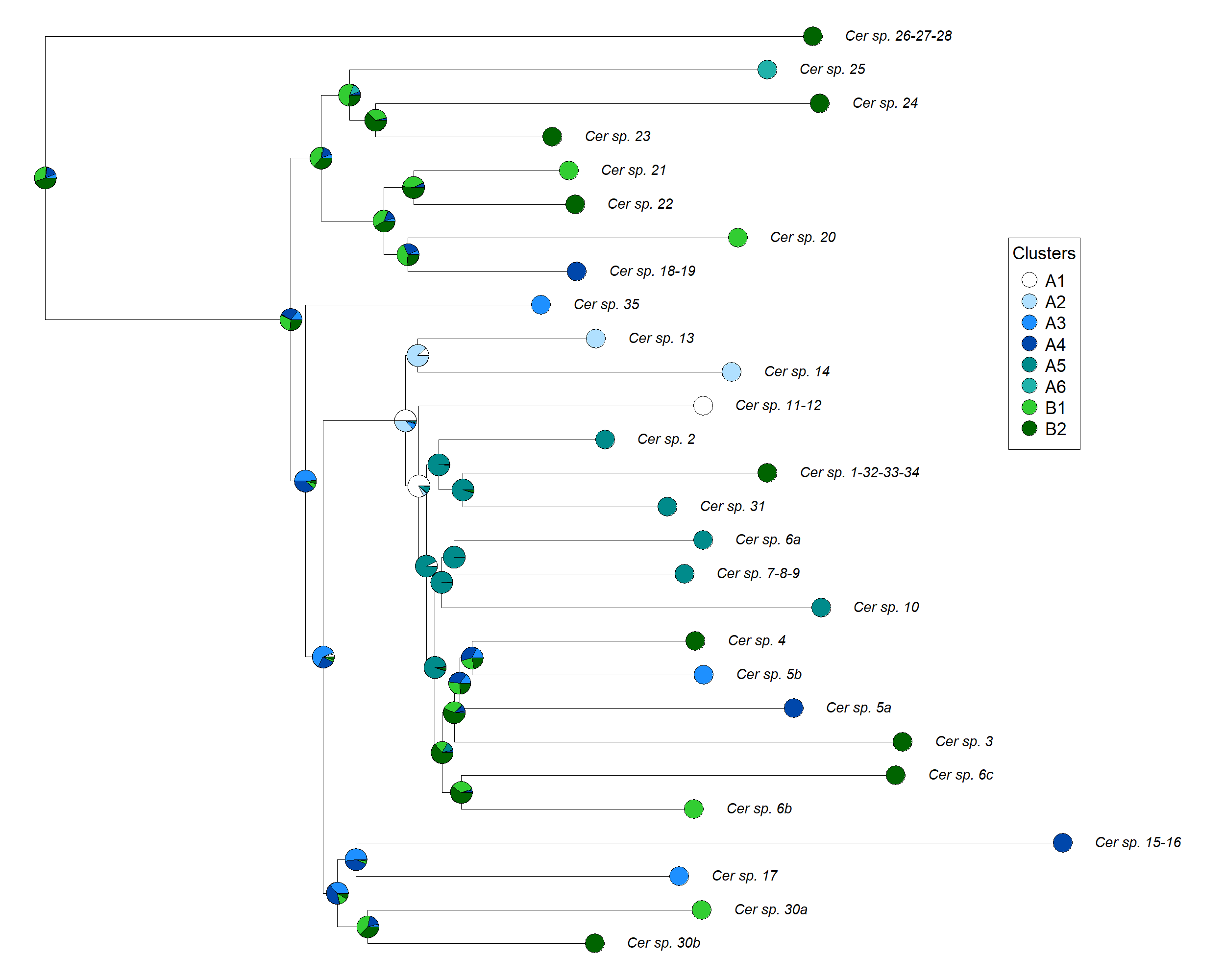

Figure S85. Ancestral state reconstruction of *Ceroptres* species association with galls of the asexual or sexual generation of gall wasps.

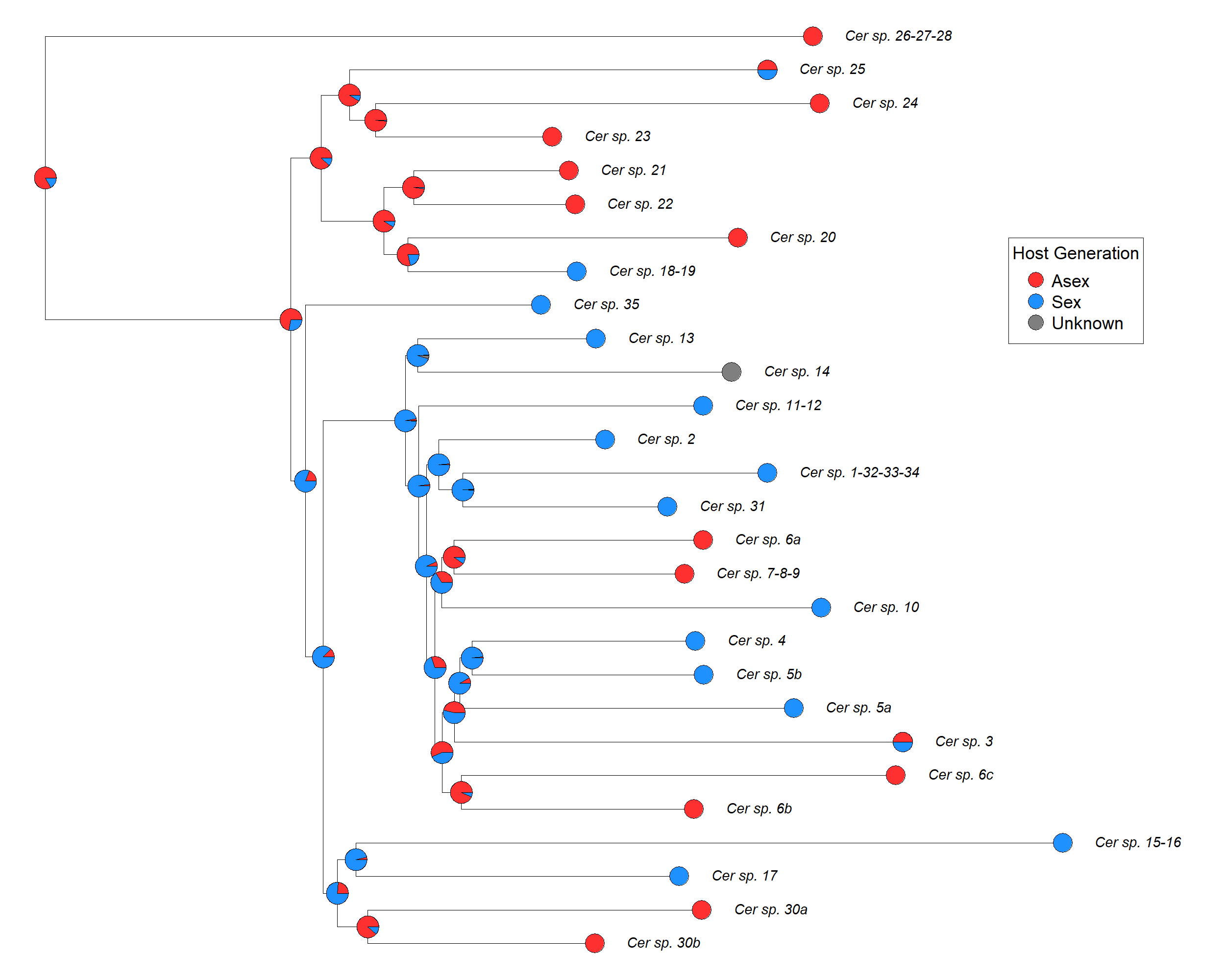
